# Sapogenin based self-assembly structures activating a non-apoptotic cell death via multiple pathways

**DOI:** 10.1101/2021.09.07.459231

**Authors:** Göklem Üner, Erdal Bedir, Onur Serçinoğlu, Petek Ballar Kırmızıbayrak

## Abstract

Induction of distinct cell death pathways is critical to deal with tumor heterogeneity and therapeutic resistance. In our previous study, we reported a promising saponin analog (AG-08) for cancer therapy inducing non-canonical necrotic cell death. Here, we describe that AG-08 forms unique supramolecular structures responsible for its biological activity. After internalization via non-canonical endocytosis pathway, these structures affect several cell signaling pathways including unfolded protein response, immune response, oxidative stress and heat stress. Moreover, we prepared 18 analogs to reveal the role of residues on the formation of supramolecular structures and biological activities. The results have demonstrated that unique structural features are required for particulate structures and unprecedented cell death mechanism. Although small molecule based supramolecular assemblies have widely been accepted as nuisance for drug discovery studies, our results indicate that they may provide a new research field for anti-cancer drug development studies.

## 1. Introduction

After the discovery of regulated necrosis, the induction of necrotic cell death has attracted huge attention and numerous studies state that necrotic death inducers possess high potential for cancer treatment. In a recent study, we reported a new semi-synthetic saponin analog (AG-08) triggering necrotic cell death along with unprecedented pathways including enhancement of global proteolysis and several alterations in lysosomal function and physiology[1]. The subsequent studies revealed an unexpected property of AG-08: After one freeze thaw cycle of its solution, AG-08 lost its cytotoxicity without any change in its chemical structure. Thus, we readily suspected formation of self-assembly supramolecular structures. A number of studies reported that small molecules could form colloidal aggregates in aqueous environments. As they non-specifically bind and inhibit enzymes, these colloidal structures are described as one of the reasons for the false positive results in bioactivity screenings[2-4]. Interestingly, the formation of colloidal aggregates by some FDA-approved drugs such as fulvestrant, lapatinib and sorafenib have also been reported. While the colloids of these drugs non-specifically inhibit enzymes in cell free condition, their activities were lost in the course of cytotoxicity screening studies. On the other hand, several small compounds of natural origin (adenine, phenylalanine, cysteine, tyrosine, adenine, uracil and orotic acids) were stated to gain cytotoxic properties after forming amyloid-like ultrastructures at high concentrations. These supramolecular structures are suggested to cause various metabolic disorders such as phenylketonuria[5]. Concomitantly, the side effects of the artificial sweetener, viz. aspartame (a small peptide), were reported to originate due to the formation of amyloid-like structures at high doses[6].

Considering the false positive results in cell-free enzyme inhibition tests, the unwanted toxicities, and the loss of biological activities, the small molecules forming aggregates are generally excluded from further drug development studies. Thus, there is huge gap in the literature regarding the physicochemical properties, and biological activities of small molecule-based supramolecular assemblies. On the other hand, limited number of studies indicate that some of these structures may have unique properties. The protein binding properties or incorporation of other molecules into the aggregate structure may reflect their potential as being candidates for protein and drug transport[3, 7]. Also, there are studies indicating that some small molecules forming aggregates may have fascinating biological activities. For instance, supramolecular structures formed by a molecule called 1543 activates caspases and induces apoptosis along with general proteolysis[8]. Similarly, a compound, which contains a naphthyl group and two phenylalanine residues, self-assembles into nanofiber structure at relatively high concentration (320–340 µm) and induces apoptosis via disruption of the dynamics of microtubules[9]. Betulinic acid nano tubes formed in ethanol and water mixture, induce reactive oxygen species (ROS) mediated apoptosis accompanied with increase in the levels of pro-inflammatory cytokines[10]. Additionally, ursolic acid forms aggregates and induces IL-1ß expression by binding to the CD36 receptor. Interestingly, seven derivatives of ursolic acid were also determined to form aggregates but they did not cause any change in IL-1ß synthesis[11]. These results indicate that small molecule-based aggregates have unique characters and activities.

Here, we report that a saponin analog AG-08 self-assembles into supramolecular structures responsible for its intriguing cell death inducing activity. After being internalized via non-canonical endocytosis, AG-08 particles mainly affect the expression of genes taking role in unfolded protein response, heat stress and immune response. Moreover, in order to see the initial structure activity relationships (SAR), we prepared 18 analogs, four of which exhibited similar bioactivity profiles as in AG-08. Collectively, our results indicate that both the biophysical properties and unique bioactivity through a distinct cell death pathway make AG-08 an exciting molecule for future research. In general perspective, this study has provided new evidence that small molecule-based supramolecular assemblies should not be neglected with prejudgment as they may hold great opportunities for drug development studies.

## 2. Results

### 2.1. AG-08 forms self-assembly particles

In our previous studies, we discovered that AG-08 (Figure 1) induces necrosis via enhanced global proteolysis involving calpains, cathepsins and caspases. Strikingly, cytotoxic activity of AG-08 was found to be lost after even one freeze thaw cycle without any change in its chemical structure. Since freeze thaw cycle was reported to affect molecular clusters [12], we suspected formation of supramolecular structures. Therefore, to ascertain that AG-08 formed particles in neutral buffer/medium, we performed a STEM microscopy analysis and revealed that circular particles were merged to form micro-sized supramolecular structures (Figure 2A). Next, the minimum concentration of AG-08 required for the particle formation was determined by using Nile red staining. Nile red is widely used to monitor supramolecular structures such as micelles or aggregates since it is almost non-fluorescent in aqueous condition but tends to associate with hydrophobic domain within the assemblies and intensely fluoresces[13, 14]. The critical assembly concentration of AG-08 was found to be approximately 2 µM (Figure 2B). Moreover, when the particle formation of parent compound (AG) was investigated via Nile red, no fluorescence intensity even at 30 µM was observed (Figure 2B). This finding indicates that AG is not capable to form ordinary supramolecular structures interacting with Nile red. Subsequently, diameters of the particles were measured using zeta sizer (DLS–Particulate Systems) and a uniform particle size distribution was observed ranging from 0.5 to 1.3 micrometer depending on AG-08 concentration (Table 1).

**Table 1.** Diameter of 4, 8 and 16 µM AG-08 in PBS was detected via Zeta Sizer.

| DMSO | 4 μM | 8 μM | 16 μM |
| --- | --- | --- | --- |
| 0 | 443.9±2.33 | 724.83±10.22 | 1353.67±44.34 |

**Figure 1.**
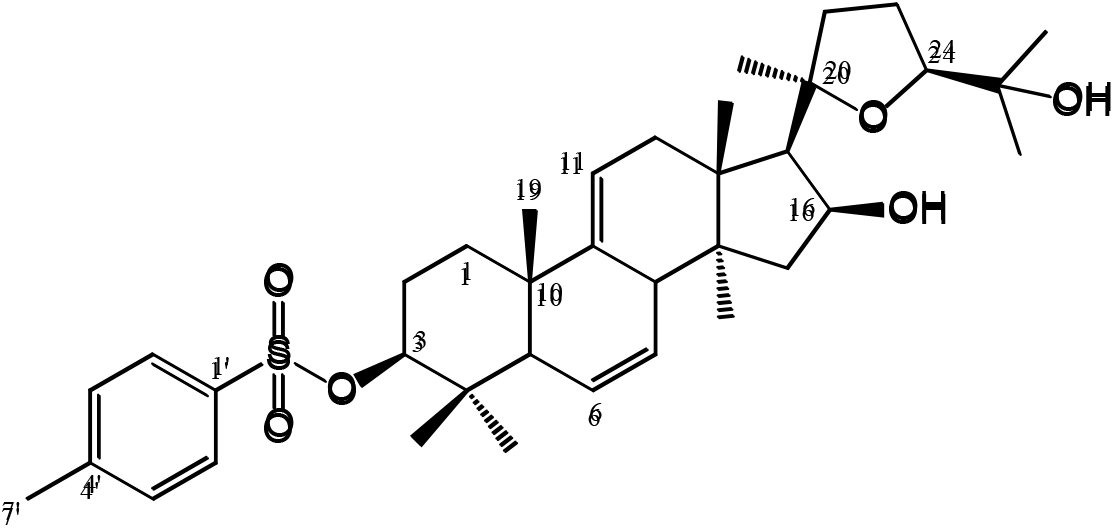
Structure of AG-08.

**Figure 2.**
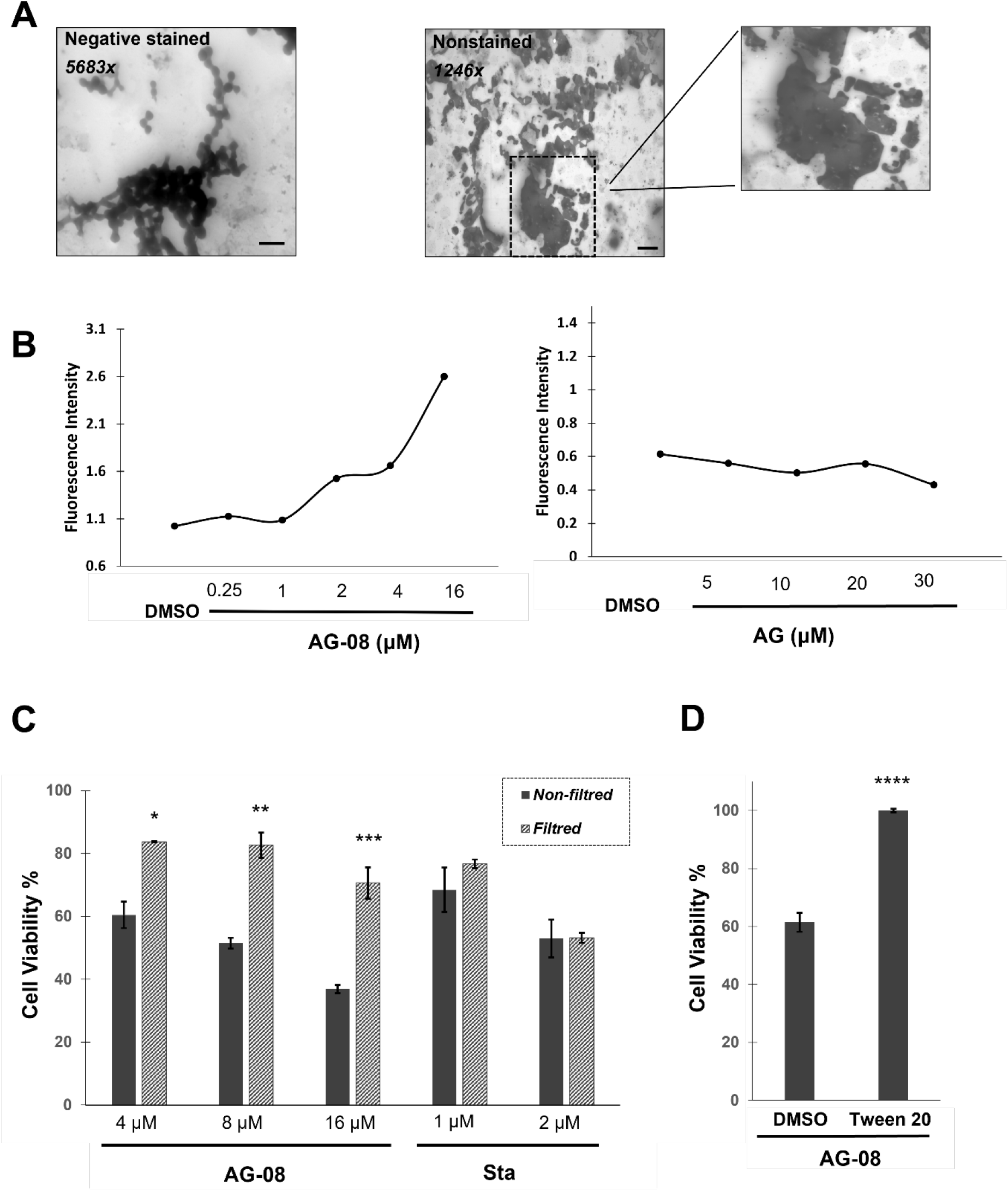
AG-08 mediated cytotoxicity relies on the supramolecular structures. **(A)** The scanning transmission electron micrograph of 16 µM AG-08 in PBS. Right picture shows non-stained image (1246× magnification; scale bar=2 µm), while left one is a representative image of stained sample with uranyl acetate (5683× magnification; scale bar=600 nm). **(B)** After 3 h incubation of Nile red solution with AG-08 or AG, fluorescence intensity of Nile red was measured. **(C)** AG-08, Staurosporine (Sta) or vehicle samples prepared in the cell condition medium was filtrated by 0.22 µm filter and HCC1937 cells were exposed to the filtrates. After 24 h, the viability of cells was measured and represented as a percentage of that of the vehicle control. The error bars represent mean value ±s.d. (n=3). *p<0.05, **p<0.005, ***p<0.001. **(D)** Tween 20 (1:40000) was added to fresh medium before addition of 8 μM AG-08 or vehicle. Then cell medium was replaced with this fresh medium. Cell viability was detected after 24h incubation. Error bars are the standard deviations (n=3; ****p<0.0001)

To verify that the activity of AG-08 arises from its the micro-sized supramolecular structures, freshly prepared AG-08 solutions were filtered using 220 nm-filters to remove AG-08 particles from medium, and the filtrates were analyzed for their cytotoxicity. While filtrates of staurosporine solution, a well-known apoptosis inducer, still effectively caused cell death, the filtered AG-08 sample no longer exhibited any cytotoxicity (Figure 2C). Next, Tween 20 was utilized as non-ionic detergent to block particle formation, and AG-08 mediated cell death was inhibited as expected (Figure 2D). Collectively, our findings proved that AG-08 but not the parent compound forms self-assembly particles, and those particles are responsible for the previously reported distinct cell death signaling.

### 2.2. AG-08 shows a higher propensity to self-aggregate in molecular dynamics simulations

Next, we tested whether the observed difference in terms of the aggregation propensity of AG and AG-08 may be confirmed from a theoretical perspective as well. For this purpose, we used molecular dynamics simulations, which was used successfully before to distinguish between aggregators form non-aggregators[15]. The underlying principle of this method is that when placed randomly in a simulation box containing solvent molecules, a self-aggregator starts to make non-bonded contacts with copies of itself instead of the surrounding waters, whereas a non-aggregator will remain dispersed throughout the simulation box. Considering the fact that aggregation behavior of small molecules is also highly dependent on the concentration, the number of molecules as well as the size of the simulation box should be carefully selected in order to mimic experimentally obtained critical assembly concentrations. However it is not always possible to test desired concentration ranges with MD simulation, as lower concentrations require large box sizes with fewer molecules, which in turn require longer simulation times (and hence higher computational capacity) to allow copies of molecules to clash into each other so that aggregation behavior is tested[16]. The systems we could afford to construct for AG as well AG-08 thus included significantly higher concentrations than their observed critical assembly concentration ranges (Table 2). Although the results therefore may not necessarily be easily compared to experimental observations, it can be still reasoned that an analysis of intermolecular contacts even at these high concentrations may yield useful insight into the self-aggregation behavior differences between the molecules.

**Table 2.** Molecular systems used in MD simulations and number of each contact type observed at 250 ns of simulation time.

|  | **AG** | **AG-08** |
| --- | --- | --- |
| **Number of molecules** | 59 | 55 |
| **Concentration** | ~60 mM | ~60 mM |
| **Water molecules** | 41704 | 41388 |
| **Box size** | 1331 nm<sup>3</sup> | 1331 nm<sup>3</sup> |
| **Contact Type\* (at 250 ns)** |  |  |
| **vdw_clash** | 20 | 207 |
| **vdw** | 38 | 86 |
| **proximal** | 4499 | 6516 |
| **hbond** | 3 | 31 |
| **w_hbond** | 125 | 119 |
| **aromatic** | 0 | 98 |
| **hydrophobic** | 991 | 1627 |
| **polar** | 24 | 108 |
| **w_polar** | 95 | 161 |
| **total** | 4557 | 6809 |

MD simulations were allowed to run to cover 250 ns of simulation time for each system. Our results indicated that both molecules may undergo self-aggregation at the studied concentrations and formed aggregates already at 5 ns of simulation time. As time progressed, the aggregates were observed grow larger, yet never fully reached a single large aggregate (Figure 3). Accordingly, AG may start to form aggregates at 60 nM. Despite the similarity in aggregation behavior of AG and AG-08 in MD simulations, an analysis of intermolecular contacts indicated that the aggregates of AG-08 may be more tightly connected to each other than those of AG. This is evident from the non-bonded interaction numbers within the formed aggregates, which were computed using Arpeggio based on the aggregate structures obtained at 250 ns[17]. Accordingly, higher number of non-bonded interactions were observed between copies of AG-08, including van-der Waals interactions, hydrogen bonds as well as hydrophobic and polar interactions (Table 2).

**Figure 3.**
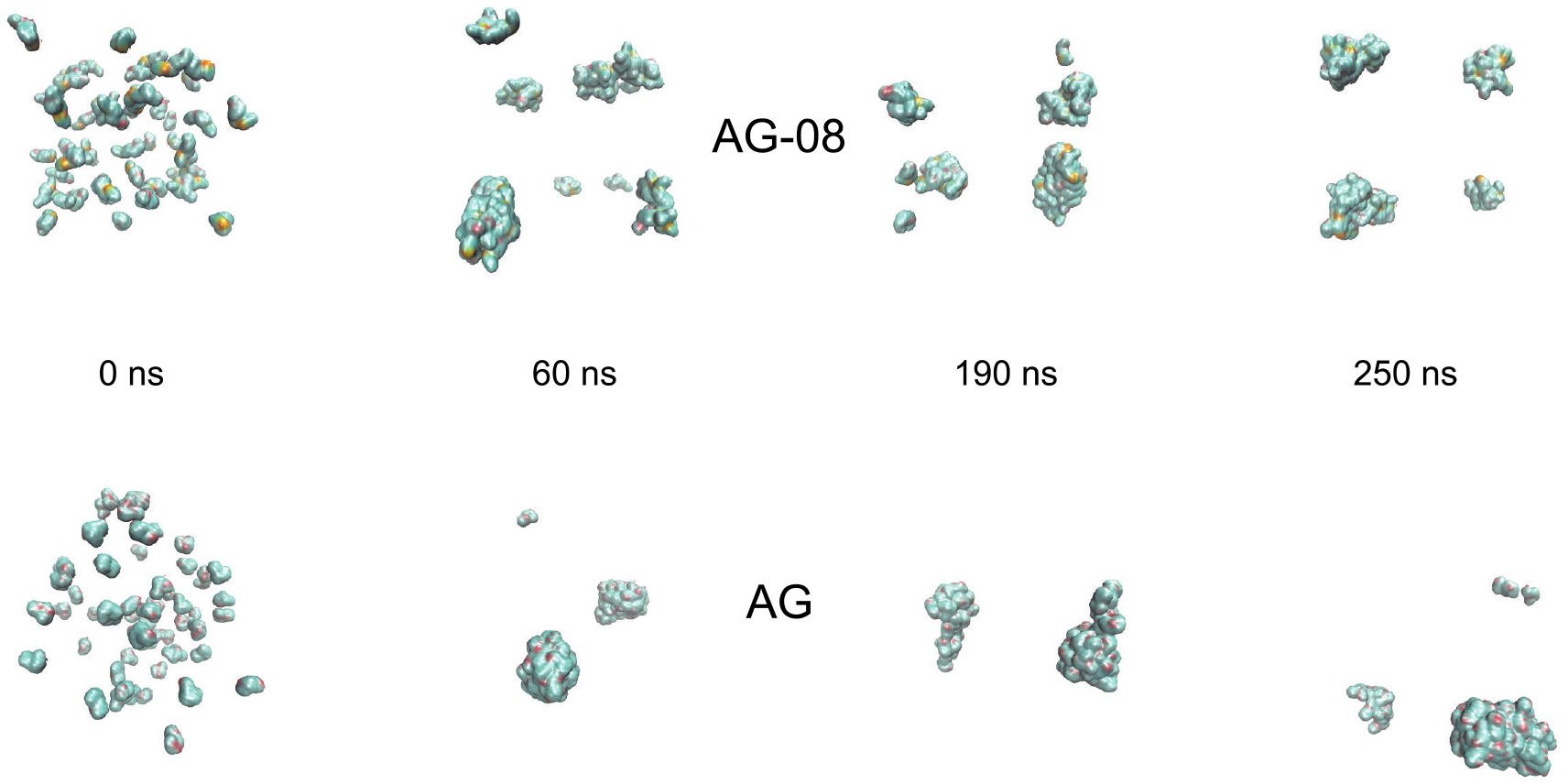
AG and AG-08 form aggregates at 60 nM concentration in MD simulations.

### 2.3. Interrelationship between AG-08 particles and endocytosis

Next, the relationship of AG-08 with endocytosis, which is a transport process responsible for the cellular uptake of particles, was assessed by utilizing several endocytosis inhibitors. Our results revealed that methyl β-cyclodextrin, targeting cholesterol related endocytosis through cholesterol depletion[18], and pitstop II, an inhibitor of both clathrin dependent and independent endocytosis[19, 20], effectively protected cells against AG-08 mediated cell death (Figure 4A). Intriguingly, dynasore, a dynamin inhibitor[21], cytochalasin D, an inhibitor of actin polymerization responsible for membrane ruffling[22], and chlorpromazine, a specific inhibitor of clathrin dependent endocytosis[23] that prevents coated pit assembly at the cell surface, failed to rescue cells against AG-08 mediated cell death (Figure 4A). Notably the protection of cells from AG-08 mediated cell death by methyl β-cyclodextrin and pitstop II was found to be dose dependent (Figure S1A-E). These results suggest that AG-08 utilizes a clathrin, dynamin and actin-independent but cholesterol dependent non-canonical endocytosis pathway. In addition, Brefeldin A, an inhibitor of vesicular trafficking, reduced AG-08 dependent cell death, suggesting AG-08 microparticles should be transported via vesicles in cells (Figure 4B).

**Figure 4.**
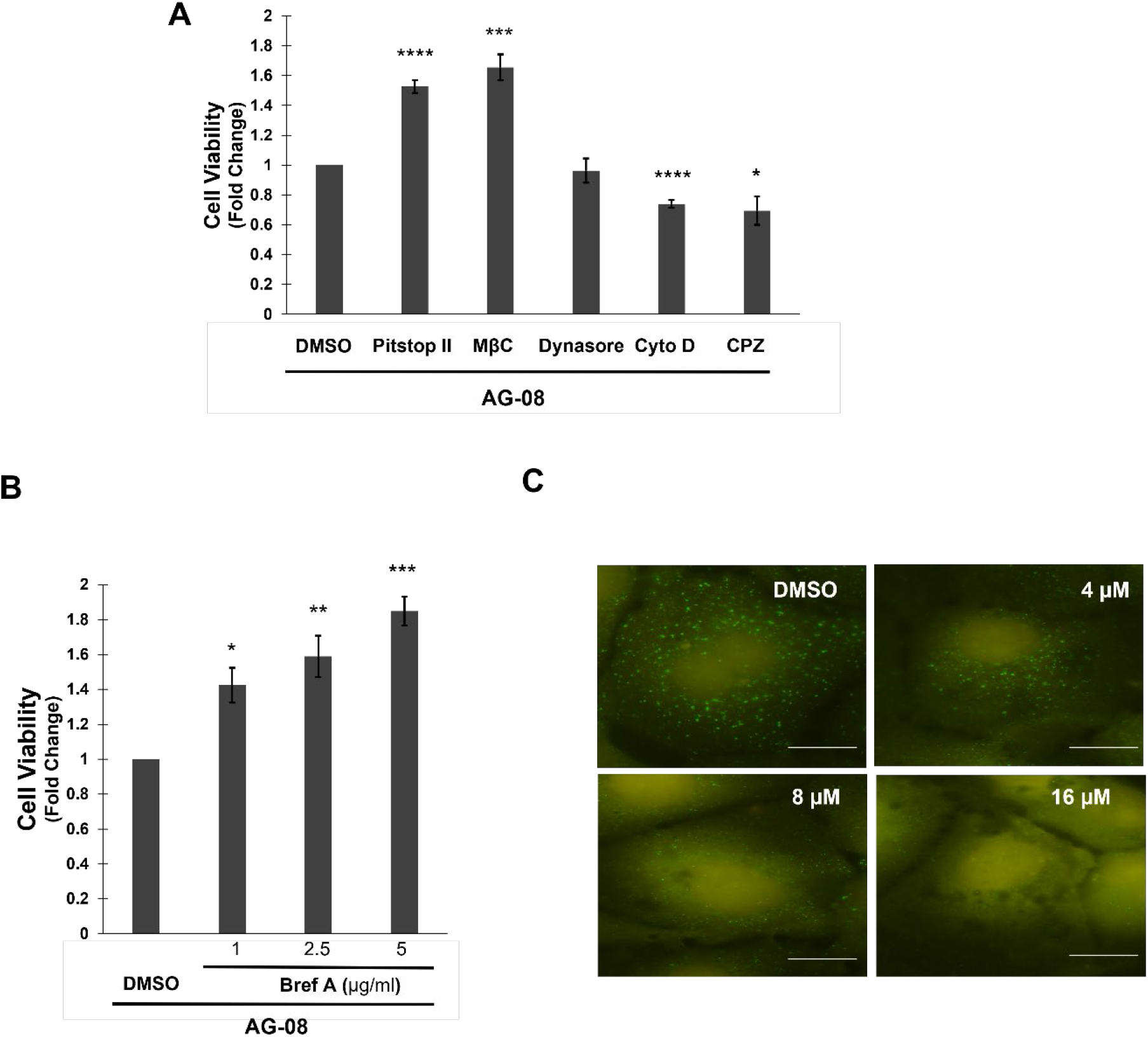
AG-08 is intriguingly related with endosomal pathway. **(A)** After pre-treatment with 20 µM pitstop II, 0.5 mM methyl-β-cyclodextrin (MβC), 80 µM dynasore, 20 µM cytochalasin d (Cyto D) or 20 µM chlorpromazine (CPZ) for 1 h, HCC1937 cells were treated with 8 µM AG-08 for 24 h. Then cell viability was determined by using WST-1. Reported values were normalized on cells treated with only AG-08. Error bars are the standard deviations (n=3). p-values were calculated with respect to AG-08 treated cells by two-tailed equal variance Student’s t-test (*p<0.05, ***p<0.001, ****p<0.0001) **(B)** HCC1937 cells were treated with brefeldin A (Bref A) for 1 h and then treated with 8 µM AG-08 for 24 h. Error bars represent standard deviations (n=3). p-values were calculated with respect to AG-08-treated cells (*p<0.05, **p<0.005, ***p<0.001). **(C)** Following AG-08 treatment for 16 h, EEA1 proteins were stained using anti-EEA1 antibody in HCC1937 cells. (Scale bar=25 µm).

To further understand the interrelationship between AG-08 and endocytosis, EAA1 (early endosome antigen 1), an early endosome marker, was stained in control and AG-08 treated cells. A dramatic decrease in EEA1 positive vesicles was observed by AG-08 treatment (Figure 4C and S1F).

### 2.4. General response to AG-08 particles

To explore the general cellular response to AG-08 treatment, microarray gene expression analysis of cells treated with vehicle and AG-08 treated groups was performed. The genes with a fold change were ≥3.0 and p-value ≤ 0.05 (unpaired t-test) were considered significantly regulated via AG-08 treatment. The expression of 193 genes (78 were upregulated and 115 were downregulated) were significantly affected by AG-08 treatment (Figure 5A., Table S1). Next, protein-protein interaction (PPI) network analysis was evaluated by the STRING (Search Tool for the Retrieval of Interacting Genes/Proteins) network. The constructed PPI network is shown in Figure 5B. There were 61 nodes and 145 edges in the network (PPI enrichment p-value:< 1×10^−16^). The number of interactions was significantly higher than expected for a random set of proteins. Then, enrichment was performed using GO and KEGG pathway analyses. Selected six GO and KEGG terms with the smallest p-values were shown in Figure 5C. The results of enrichment analysis revealed that noticeable variation in the expression of proteins in pathways associated with unfolded proteins, immune system, and oxidative response. Also, the same finding was confirmed by Reactome analysis which showed significant enrichment in heat stress related pathways.

**Figure 5.**
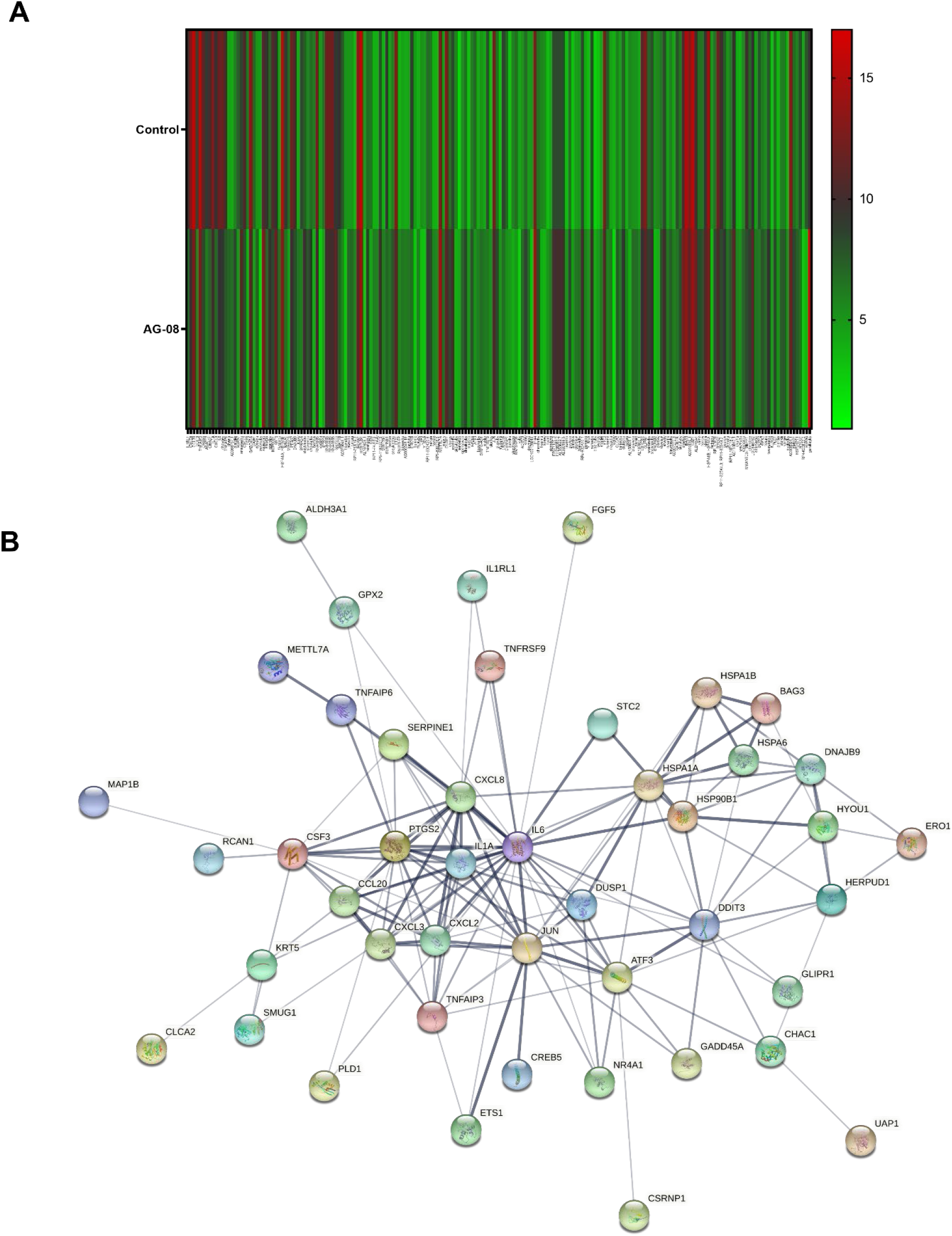
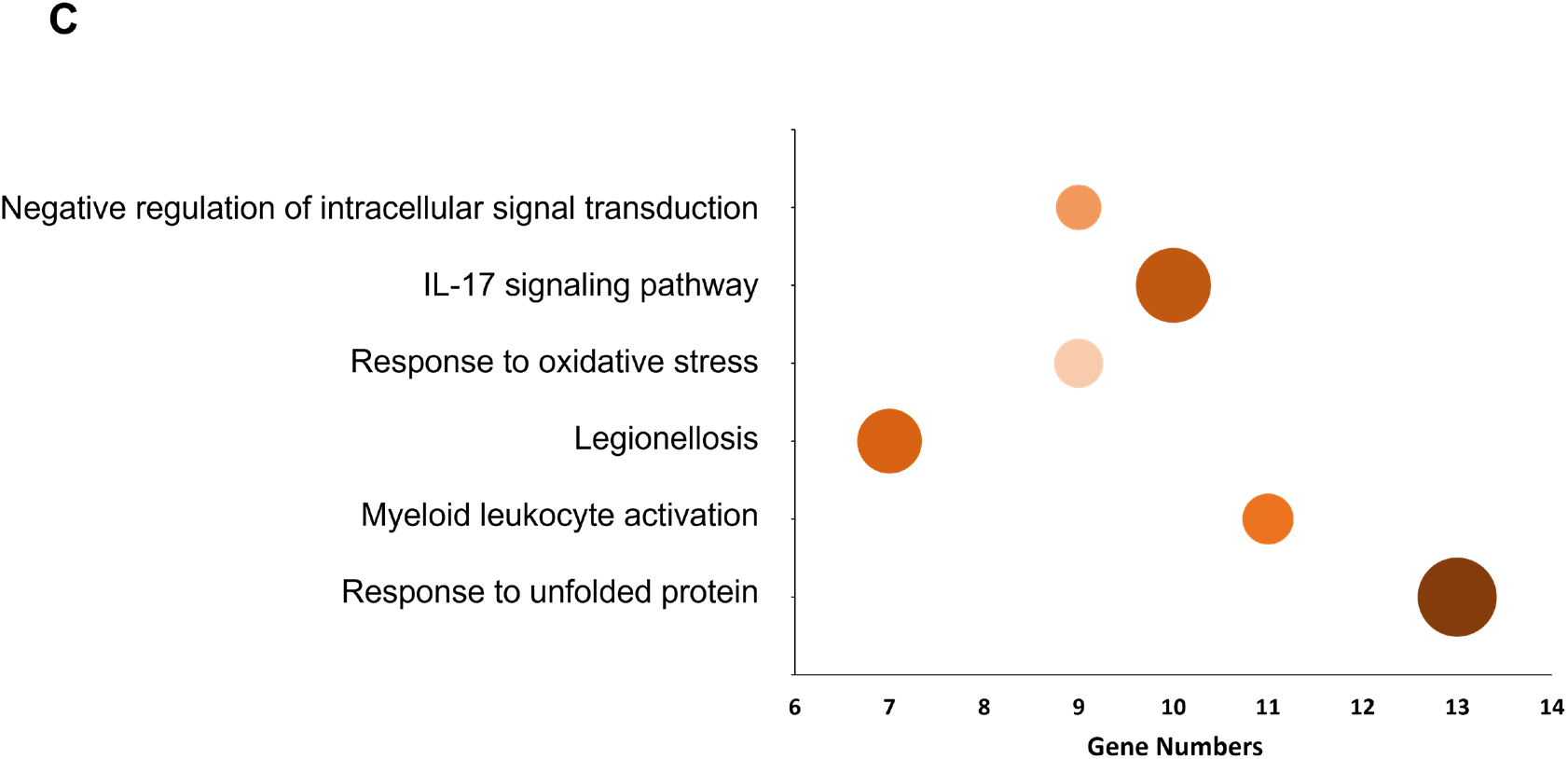
mRNA microarray analysis was used to identify differentially expression genes via AG-08 treatment. **(A)** Heatmap representing the expressions of 193 genes with significant different expression in AG-08 treated cells compared with control. **(B)** Protein-protein network analysis was formed via STRING. The thickness of the connecting lines indicates the strength of data support for physical and/or functional associations. **(C)** Pathway enrichment was determined using GO, KEGG and Reactome pathway analysis. Figure displays six terms with the smallest p-values. Size of circle represents the significance of enrichment.

### 2.5. AG-08 particles induce UPR

Microarray results showed that one of the pathways induced by AG-08 particles is Unfolded Protein Response (UPR) which is known to be activated after prolonged endoplasmic reticulum (ER) stress [24]. Thus, we aimed to investigate the induction of ER stress upon AG-08 treatment. In consistent with microarray results\ our data verified that CHOP (DDIT3), a known ER stress marker, was upregulated in AG-08 treated cells in a dose dependent manner (Figure 6A) (Table 3). Interestingly, CHOP protein level was enhanced by 8 µM AG-08 treatment but surprisingly decreased by 16 µM (Figure 6B). Since similar results was observed in the p62 protein levels in the previous study[1], we suggested that the CHOP levels were induced due to ER stress, but the enhancement of proteolysis by the prolonged or higher dose treatments of AG-08 might lead to degradation of full-length CHOP protein. Concomitantly, the expression level of calnexin, a chaperone localized in the ER upregulated during ER stress[25] was analyzed and the intensity of calnexin staining was found to be significantly increased in AG-08 treated cells (Figure 6C). Collectively, all these data confirmed that AG-08 particles induce ER stress.

**Table 3.** Upregulation of UPR genes with AG-08 treatment.

| Genes | Fold Changes |
| --- | --- |
| CXCL8 | 9.38 |
| DDIT3 (CHOP) | 7.57 |
| DAW1 | 5.06 |
| HSP90B1 | 4.89 |
| DNAJB9 | 4.59 |
| HERPUD1 | 4.06 |
| ATF3 | 3.61 |
| HYOU1 | 3.14 |

**Figure 6.**
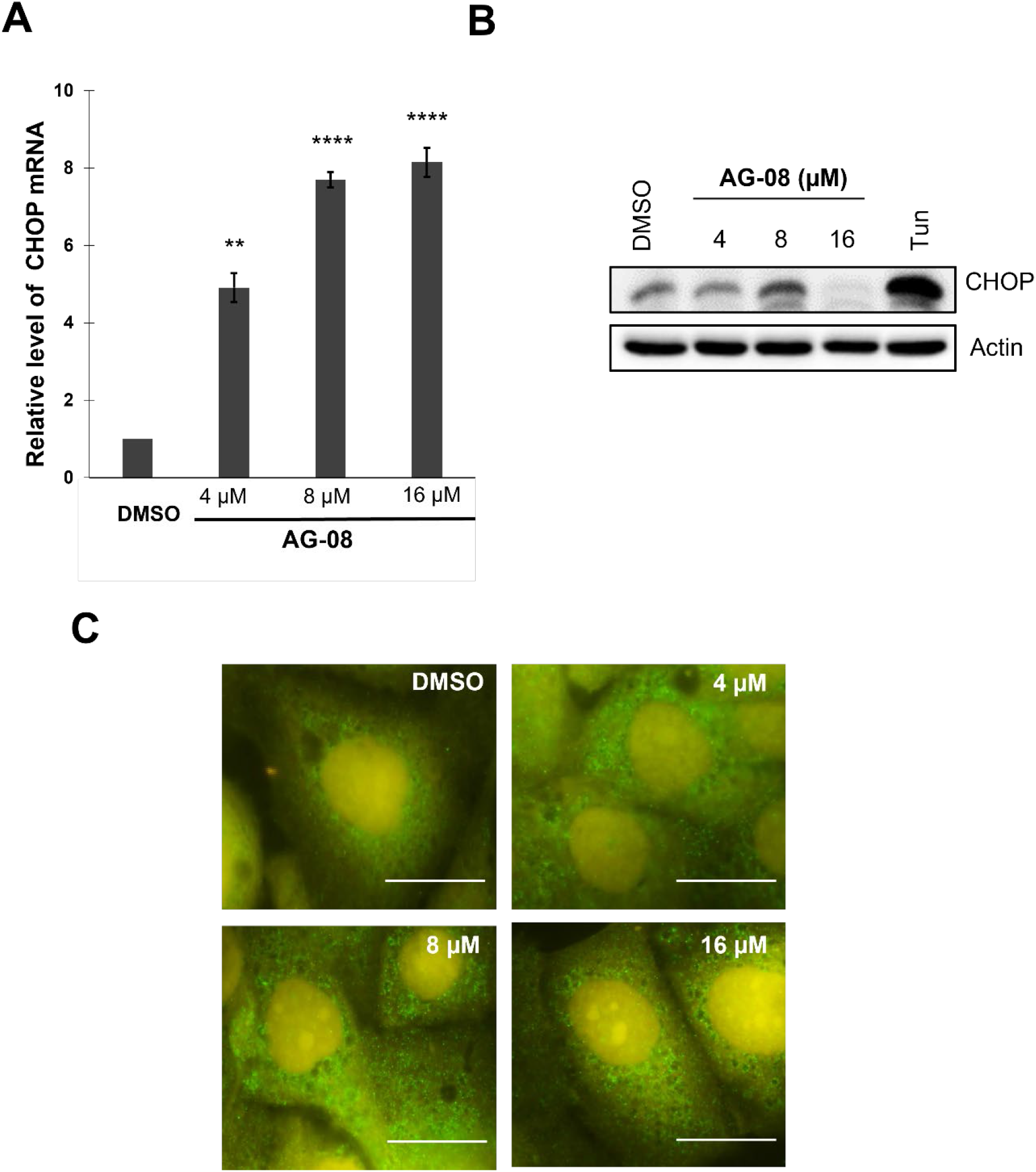
ER stress is induced by AG-08 treatment. HCC1937 cells were treated with AG-08 or vehicle for 16 h. **(A)** The protein levels of CHOP were determined via IB. 0.5 µg/ml tunicamycin (Tun) was used as positive control. The experiments were repeated three times independently; with one representative result shown. **(B)** mRNA level of the CHOP was investigated using quantitative PCR (q-RTPCR). Error bars represent standard error (s.e.). p-values were calculated with respect to vehicle-treated cells by one-way Anova test. (**p<0.005, ****p<0.0001). **(C)** The expression and localization of Calnexin protein was evaluated using immunofluorescence via anti-Calnexin antibody. (Scale bar=25 µm).

### 2.6. Synthesis of analogs

We next aimed to further characterize the relation between the chemical structure of AG-08 and its capacity of supramolecular assembly formation together with its biological activities by preparing further derivatives. In this regard, 18 analogs (Figure 7) were synthesized from 20(27)-octanor-cycloastragenol (SCG), cycloastregenol (CG) and astragenol (AG) by the methods shown in the method section. The details and characterization of which are shown in Supplementary Information. The cytotoxic activities of the semi-synthetic analogs were evaluated against four human cancer cell lines, namely HeLa, HCC1937, MCF7, and A549 as well as against MRC-5 as a normal cell line. Molecules having lower IC_50_ values than 20 µM were considered as potent cytotoxic molecules and included into subsequent studies. The compounds except CG-03, CG-06, AG-05, and CG-05 did not exhibit promising cytotoxicity (IC_50_ >20 µM) (Table 4). While AG-05 (between 1.55 and 5.9 µM) and CG-05 (between 1.85 and 6.1 µM) showed relatively higher cytotoxic activity, the potency of AG-06 (between 5.7 and 19.07 µM) and CG-03 (between 3.65 and 12.87 µM) were notably lower than AG-08.

**Table 4.** IC_50_ values (µM) of the analogs and their parent molecules.

|  | **HeLa** | **HCC1937** | **MRC-5** | **MCF7** | **A549** |
| --- | --- | --- | --- | --- | --- |
| **AG-01** | >30 | >30 | >30 | >30 | >30 |
| **AG-02** | 42.6 | 46.08 | 39.9 | 32.1 | 30.5 |
| **AG-03** | 33.42 | 20.31 | 24.4 | 21.76 | 25.53 |
| **AG-04** | >50 | >50 | 29.55 | 22.05 | 27.5 |
| **AG-05** | 5.9 | 4.28 | 1.55 | 2.55 | 2.02 |
| **AG-06** | 10.81 | 10.02 | 19.07 | 9.34 | 5.7 |
| **AG-07** | 34.7 | 38.3 | 7.4 | ND | 38.2 |
| **AG-08** | 5.7 | 3.81 [1] | 2.58 | 3.78 | 2.3 [1] |
| **CG-01** | >30 | >30 | >30 | >30 | >30 |
| **CG-02** | 43.94 | 32.48 | 33.37 | 21.2 | 21.08 |
| **CG-03** | 12.87 | 6.13 | 5.29 | 5.5 | 3.65 |
| **CG-04** | >50 | >50 | >50 | >50 | >50 |
| **CG-05** | 6.1 | 2.66 | 2.58 | 3.85 | 1.85 |
| **CG-06** | >50 | 38.1 | >50 | 41.3 | >50 |
| **SCG-01** | >50 | >50 | >50 | >50 | >50 |
| **SCG-02** | >50 | >50 | >50 | >50 | >50 |
| **SCG-03** | >50 | >50 | >50 | >50 | >50 |
| **SCG-04** | >50 | >50 | >50 | >50 | >50 |
| **SCG-05** | >50 | >50 | >50 | >50 | >50 |
| **SCG-06** | >50 | >50 | >50 | >50 | >50 |
| **SCG-07** | >50 | >50 | >50 | >50 | >50 |
| **AG** | >30 | >30 | >30 | >30 | >30 |
| **CG** | >50 | >50 | >50 | >50 | >50 |
| **SCG** | >50 | >50 | >50 | >50 | >50 |

ND: not determined

**Figure 7.**
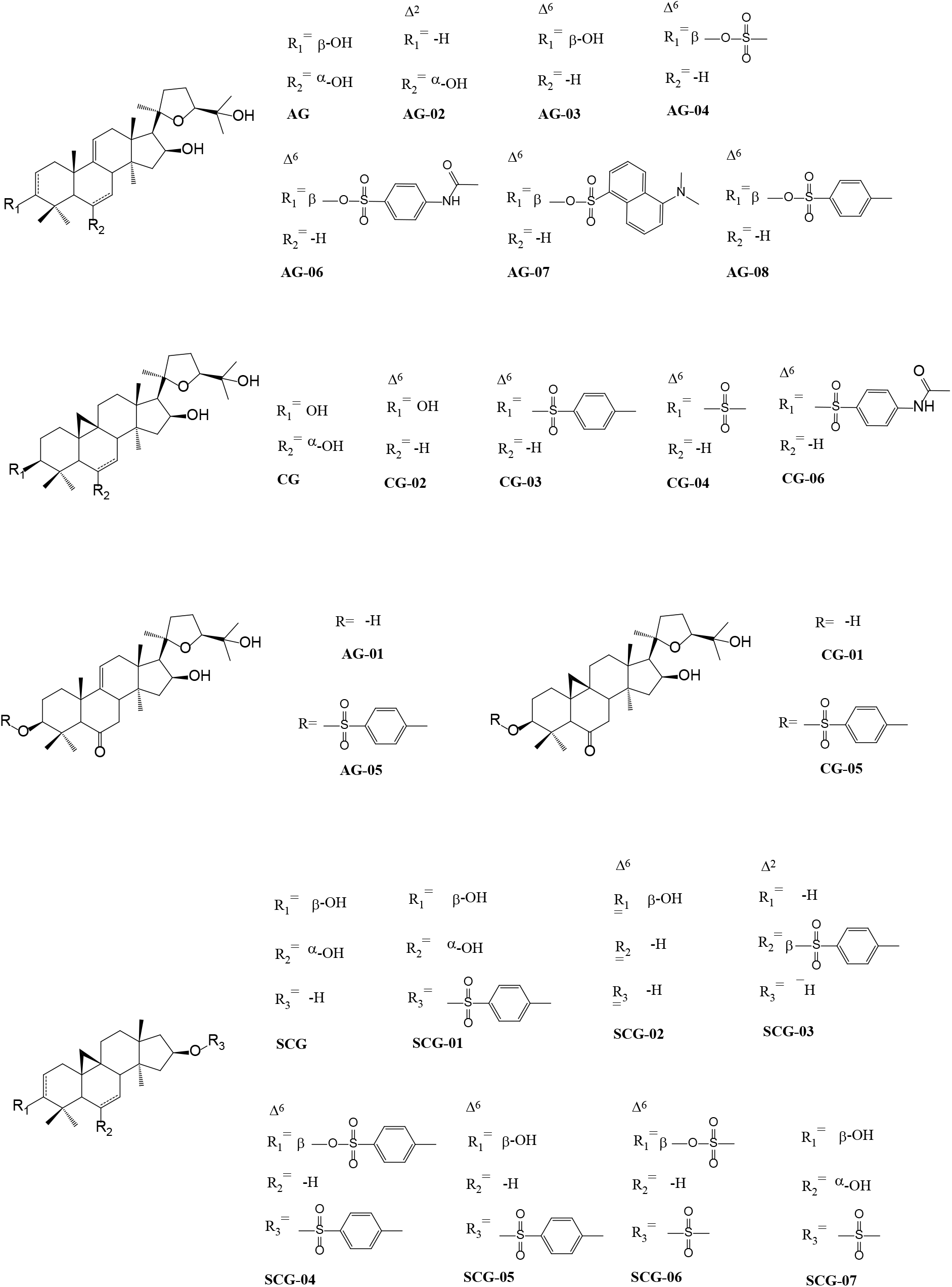
Structures of the AG-08 derivatives.

Next, the Lactate Dehydrogenase (LDH) release assay, which was previously used to demonstrate necrotic cell death for AG-08[1] was performed, and as shown in Fig 8A, all the tested compounds caused substantial release of LDH in a dose dependent manner. After LDH assay, immunoblotting was used to determine whether bioactive AG-08 analogs affect the expression of proteins known as autophagy and cell death markers in a similar manner of AG-08. Indeed, it was found that the effects of these compounds on the protein profiles of LC3, Atg-7 and caspase 3 were similar with those of AG-08 (Figure S2). Consequently, our results showed that AG-08, CG-03, CG-06, AG-05, and CG-05 induced necrotic cell death through similar mechanism of action.

**Figure 8.**
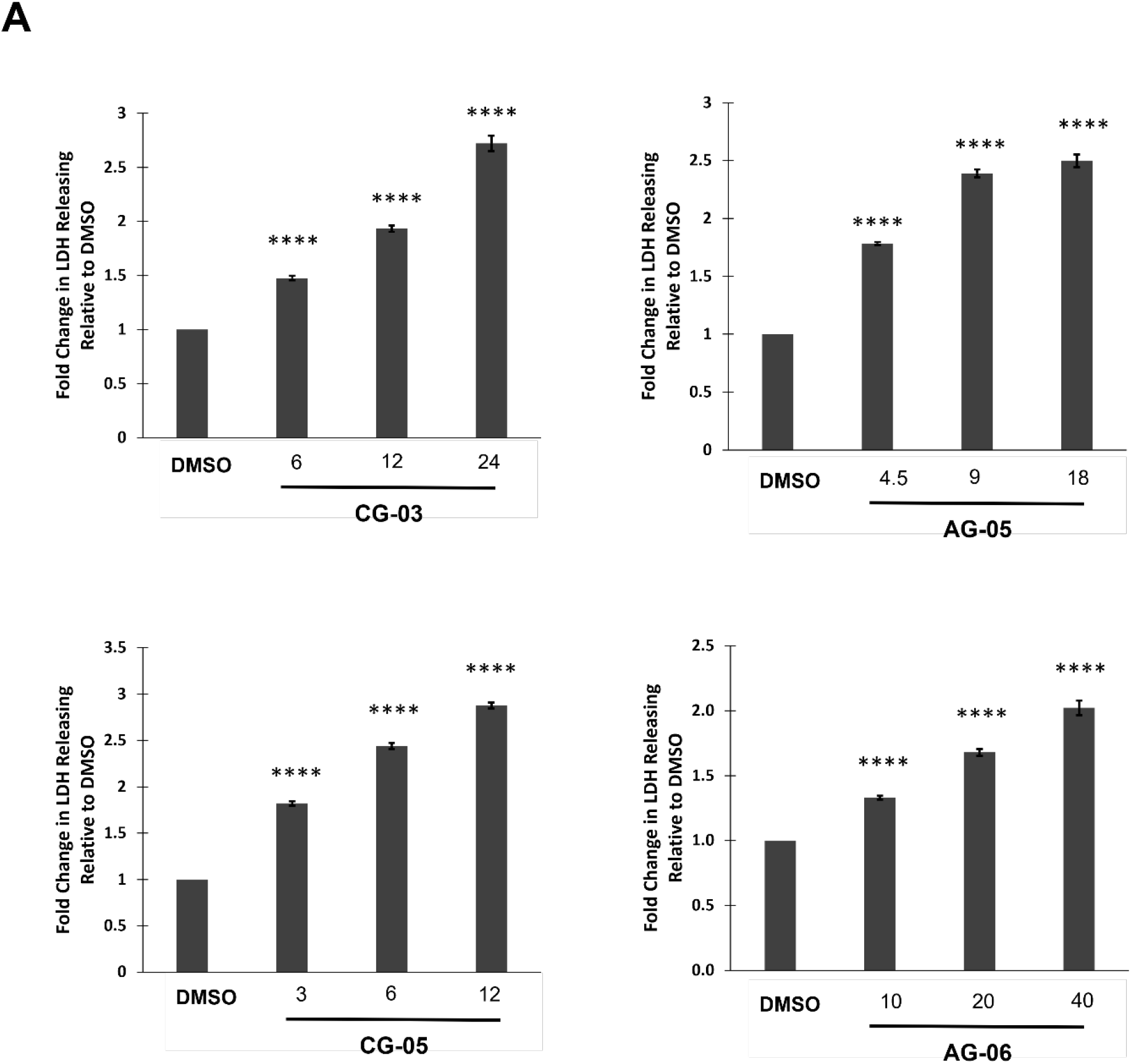
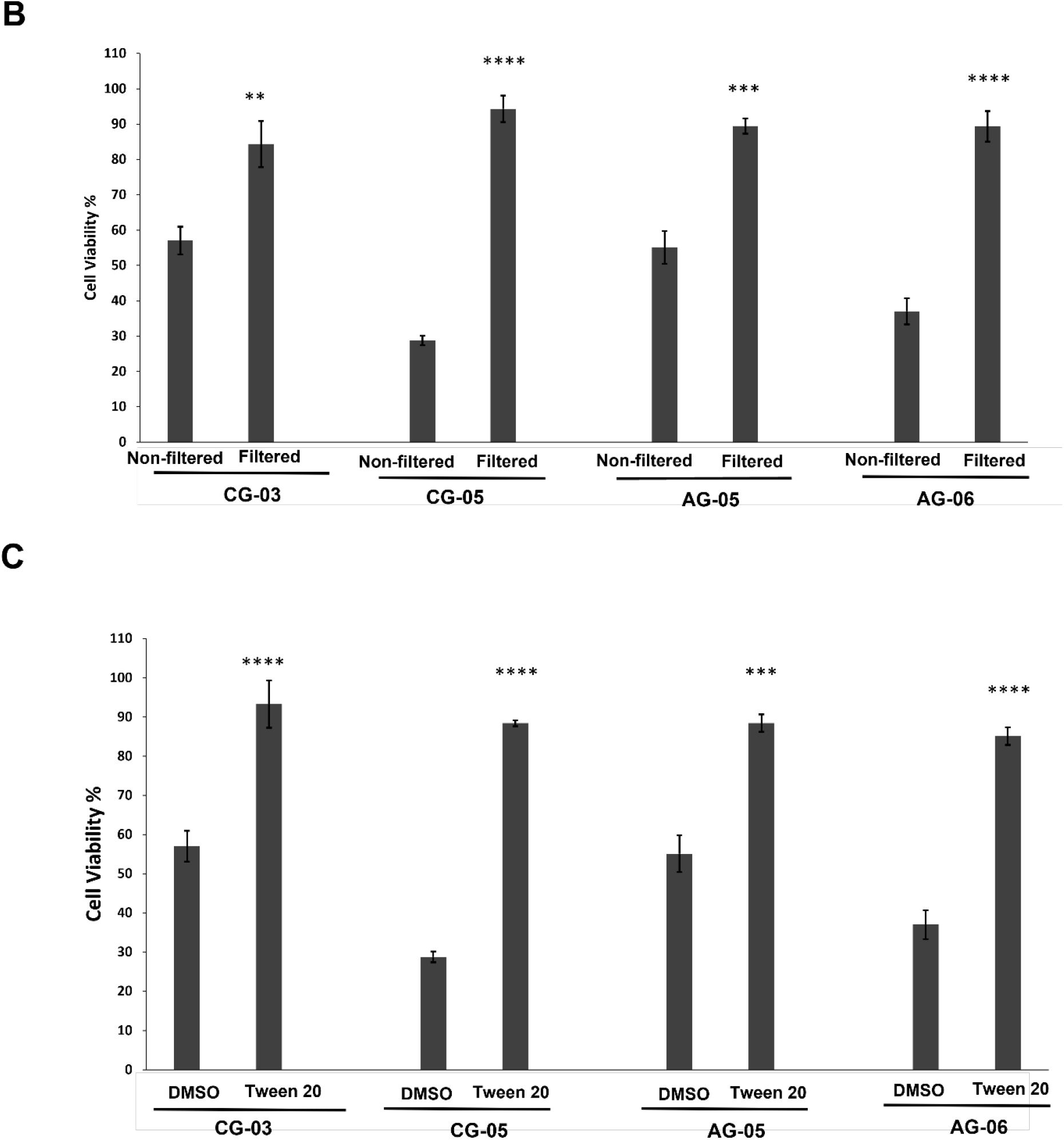
The cytotoxic activity of analogs depends on particle formation similar to AG-08. **(A)** LDH release was determined following the treatment of HCC1937 cells with 1-, 2- and 4-fold of IC_50_ values of cytotoxic compounds or vehicle (DMSO) for X h. The level of released LDH was calculated as fold change relative to vehicle. **(B)** Compounds (2-fold of IC_50_ values; 12 µM CG-03, 6 µM CG-05, 9 µM AG-05 and 20 µM AG-06) or vehicle solutions prepared in cell condition medium was filtered by 0.22 µm filter and replaced with normal medium. After 24 h, cell viability was measured. **(C)** The cell medium was replaced with fresh medium containing Tween 20 (1:40000) and compounds (2-fold of IC_50_ values; 12 µM CG-03, 6 µM CG-05, 9 µM AG-05 and 20 µM AG-06) or vehicle. Cell viability after 24 h was measured. The reported values correspond to mean values ±standard deviation (±s.d) of three experiments. ***p<0.001, ****p<0.0001.

Lastly, we aimed to evaluate whether the cytotoxicity of novel analogs also relies on the particle formation. As expected, filtration of samples via 220 nm-filters and utilizing Tween 20 to block particle formation disrupted the cytotoxic activity of all the analogs (Figure 8B and 8C). Thus, the results substantiate that the new analogs, viz. CG-03, CG-06, AG-05, and CG-05, also form supramolecular assemblies that are responsible for their actions.

## 3. Discussion

It has long been known that small molecules can form self-assembly higher order structures via noncovalent associations. However, when detected they are mostly excluded from drug development studies since self-assembly structures are reported as a reason of false positive results in enzyme inhibition assays. Moreover, their unwanted toxicities and the loss of existing biological activity strengthen their acceptance as nuisance for drug discovery. On contrary, several reports have also suggested that independent of their soluble forms these structures can provide unique biological activities; thereof, have potential to form novel drug development platforms. We have previously reported that AG-08, a semi-synthetic sapogenin, has a unique mechanism of activity including autophagy/lysosome impairment and enhancement of proteolysis leading to a necrotic cell death. In this recent study, we characterized the supramolecular structure of AG-08 and revealed that these structures but not its soluble form are responsible for its biological activity. Furthermore, we identified that AG-08 particles are internalized via non-canonical endocytosis depending on cholesterol, but not actin and dynamin. The internalization pattern of AG-08 is not typical because most of the characterized endocytosis pathways require dynamin and/or actin. On the other hand, similar to AG-08, some viruses like LCMV have been reported to enter the cells using a non-canonical pathway independent of essential endocytosis elements such as clathrin, caveolin, flotillin, Arf6, dynamin-2, and actin[26] but dependent on cholesterol[27]. AG-08 treatment also diminished the number of EEA1 positive vesicles indicating the inhibition of EEA1 recruitment to the early endosomes. Considering the inhibition of early endosome function and the role of EEA1 in tethering and/or docking of early endosome, AG-08 might be inhibiting EEA1 recruitment following its internalization through early endosome or might bypass early endosomes and go directly to late endosomes, as reported for LCMV[27]. In line with that, the protection of cells against AG-08 mediated cell death by brefeldin A, which is the inhibitor of intracellular vesicular trafficking, further suggests that AG-08 particles are transported via vesicles.

The microarray analysis and further validation studies have revealed that AG-08 particles induce a strong unfolded protein response (UPR). In our previous study, we reported that AG-08 triggers the activation of calpains and caspases as well as an increase in the activity of cathepsins in the cytoplasm. Both induction of UPR and protease activity by AG-08 particles are highly resembling to those induced by protein aggregates. For instance, the activation of calpains and caspases was reported during as a one of the cellular changes in Huntington’s disease, where proteases are possibly activated to cope with aggregates, although they cleave the huntingtin aggregates into more toxic smaller fragments[28-30]. Hence, based on the results mentioned above, one may suggest that the proteases activated by AG-08 particles contribute to the cell death via the cleavage of essential cellular components, and the enhanced proteasomal activity is a similar feature of protein aggregates and AG-08 particles.

Since the interpatient variability together with the inter- and intra-tumoral heterogeneity limit the success of cancer treatments, the discovery of different treatment approaches and/or alternative drugs such as small molecule-based anticancer drugs are highly critical especially for expanding the current chemotherapeutic repertoire and increasing options for combination therapies. The results of AG-08 signify once again that supramolecular structures formed by small molecules may have paved the way for their use for anticancer therapy. Moreover, the induction of necrotic cell death via AG-08 particles is also intriguing for cancer research since non-apoptotic cell death has attracted huge attention to cope with the resistance to apoptotic cell death, which is one of the main problems of current chemotherapeutic agents. The necrotic cell death is highly specified for intra-tumoral therapy, as it is able to activate antitumor immune response. Strikingly, PV-10, a necrosis triggering agent that was designated as an orphan drug in neuroblastoma, melanoma and hepatocellular carcinoma, underwent clinical trials for the treatment of different cancers and has been reported to cause a reduction in both injected and untreated tumors and metastasis sites[31, 32]. Tigilanol tiglate (EBC-46), which is a diterpenoid with necrotic properties, has also recently been approved by the FDA-CVM for veterinary use for intra-tumoral therapy, Furthermore, positive results were reported from phase I studies of EBC-46 with patients having accessible cutaneous, subcutaneous or nodal tumors refractory to conventional therapy[33]. Consistent with the signature of necrotic cell death, our microarray results verified that the immune system related genes were significantly altered by AG-08 treatment. Therefore, we speculate that AG-08 particles have high potential for the treatment of cancer via intra-tumoral injection.

Our computer-based study has showed that AG-08 could form much stronger non-covalent bonds than AG, the parent compound. In parallel, AG-08 forms supramolecular structures that encapsulate Nile red dye while AG cannot. Additionally, 18 analogs prepared from AG, CG, and SCG allowed us to realize the role of residues on the formation of these particles accounting for the biological activity. Among them, CG-03, AG-05, AG-06 and CG-05 were identified as cytotoxic compounds that formed supramolecular particles and induced necrotic cell death. The results also facilitated us to deduce the structure−activity relationships: Firstly, importance of the existence of a tosyl group at C-3 was evaluated: i) when the tosyl group was replaced with non-aromatic mesylate group (AG-04 and CG-04), the activity was lost; ii) AG-07 was prepared by addition of a bulky naphthalene sulfonyl chloride (dansyl chloride) to AG, which was again resulted in activity loss; iii) N-Acetylsulfanilyl substitution led to a decrease in activity with AG skeleton (AG-06) while resulting in complete activity loss in CG analog (CG-06). Thus, it is concluded that a specific sulfonyl ester (tosylate) is required for bioactivity towards cancer cells. The double bond at C-6, forming readily during tosylation reaction of AG, CG and SCG, was dispensable because the compounds with C-6 double bond and no tosyl group at C-3 did not exhibit antiproliferative effect (AG-02 and CG-02). On the other hand, the presence of a keto functionality at C-6 afforded higher activity in CG and AG analogs, viz. CG-05 and AG-05. Secondly, we prepared similar SCG analogs, missing the side chain extending from C-17 position. The SCG analogs showed no cytotoxicity suggesting that tosyl substitution by itself was not sufficient for activity alone, and intact sapogenin skeleton was required including 20,24-epoxy side chain. All these findings imply that certain chemical features must be met to lead the sapogenin skeleton to form stable particles and to show antiproliferative effects.

## 4. Conclusion

The work describes here reveals that AG-08 and four of its analogs form particles that trigger necrotic cell death with an unprecedented mechanism indicating that the small compound-based supramolecular structures have the capability to activate/suppress different cellular signals in the cell. Therefore, we demonstrate that distinct compounds can form unique supramolecular structures at low concentrations, and they have potential to become drug candidates with unique activities. We expect that AG-08 and its analogs may attract more attention in the field of saponin based drug discovery and revolutionize the drug development studies.

## 5. Methods

### 5.1. Synthesis

20(27)-octanor-cycloastragenol (SCG), CG, CG-01 and AG-01 was donated by Bionorm Natural Products (Turkey). AG was synthesized from cycloastragenol via performing a previously reported procedure[1].

#### Reaction of AG with p-TsCl

100 mg AG was dissolved in pyridine and 450 mg *p*-TsCl (p-Tosyl Chloride, Acros Organics) reagent was added. Reaction was continued for 6 h at room temperature and quenched by addition of distilled water. Following the extraction via 500 ml ethyl acetate (3 times), the extract was evaporated at 60°C in the rotary evaporator. Crude product was purified on a silica gel chromatography column (cyclohexane:EtOAc; 7:3) to give **AG-02** (3.8 mg lyophilized white-off powder), **AG-03** (6.5 mg lyophilized white-off powder) and **AG-08** (25 mg lyophilized white-off powder)[1]. For **AG-02**; ^1^H NMR (400 MHz, CDCl_3_): δ =5.5 (ddd J=10.1, 5.4, 3 Hz, 1H), 5.31 (m, 1H), 5.3 (m, 1H), 4.72 (ddd, J= 7.9, 7.9, 6.3 Hz, 1 H), 4.06 (ddd, J= 10.8, 10.8, 3.9 Hz, 1H), 3.75 (dd, J=8.1, 6.2 Hz, 1H), 2.59 (q, J= 10.4 Hz, 1H), 2.4 (m, 1H), 2.36 (d, J=7.8, 1H), 2.14 (m, 1H), 2.11 (m, 1H), 2.05 (m, 1H), 2.04 (m, 1H), 2 (m, 2H), 1.89 (m, 1H), 1.86 (m, 1H), 1.6 (m, 1H), 1.51 (dd, J=12.8, 6.2, 1H), 1.44 (m, 1H), 1.3 (s, 3H), 1.28 (m, 1H), 1.23 (s, 3H), 1.2 (s, 3H), 1.17 (s, 3H), 1.14 (s, 3H), 1.05 (s, 3H), 0.94 (s, 3H), 0.79 (s, 3H). ^13^C NMR (100 MHz, CDCl_3_): δ= 145.8, 139.6, 120.3, 116.3, 87.2, 81.5, 73.40, 72.0, 70.2, 56.2, 55.4, 45.1, 44.3, 43.9, 40.9, 40.3, 38.6, 37.9, 37.7, 36.0, 34.9, 34.6, 29.8, 28.0, 27.8, 26.7, 25.9, 23.7, 19.2, 18.1. HRMS (ESI): *m/z* 495.34667 [M+Na]^+^ calcd for C_30_H4_8_O_4_. For **AG-03**; ^1^H NMR (400 MHz, CDCl_3_): 5.71 (m, 1H), 5.57 (dt, J=10.2, 3.2 Hz, 1H), 5.16 (brs, 1H), 4.69 (m, 1H), 3.75 (t, J=7.2 Hz, 1H), 3.23 (dd, J=11.8, 4.8 Hz, 1H), 2.84 (brs, 1H), 2.56 (m, 1H), 2.24 (dd, J=11.8, 6.1 Hz 1H), 2.07 (m, 1H), 2.04 (m, 1H), 2.0 (m, 2H), 1.92 (m, 1H), 1.78 (m, 2H), 1.70 (d J=12.2, 1H), 1.59 (m, 2H), 1.58 (m, 1H), 1.57 (m, 1H), 1.29 (s, 3H), 1.23 (s, 3H), 1.14 (s, 3H), 1.01 (s, 6H), 0.98 (s, 3H), 0.83 (s, 3H), 0.64 (s, 3H). ^13^C NMR (100 MHz, CDCl_3_) δ= 145.9, 128.9, 127.3, 113.6, 87.3, 81.8, 79.3, 73.6, 72.2, 56.6, 52.2, 44.8, 44.3, 43.9, 43.8, 39.2, 38.8, 38.2, 34.8, 34.4, 28.2, 28.2, 28.2, 28.2, 26.9, 26.1, 20.4, 19.1, 18.9, 16.0. HRMS (ESI): *m/z* 495.34894 [M + Na]^+^ calcd for C_30_H_48_O_4._

#### Reaction of AG with MsCl

MsCl (140 µL) and TEA (150 µL) was added to a solution of AG (150 mg) in dichloromethane. The reaction mixture was stirred at 0 °C for 3 h. The reaction solvent was then removed under reduced pressure, and the residue was purified using silica gel column chromatography (*n*-hexane:EtOAc; 75:25) to give **AG-04** (10.5 mg lyophilized white-off powder). ^1^H NMR (400 MHz, CDCl_3_): δ =5.67 (dt, J= 10.1, 2.1 Hz, 1H), 5.63 (m, 1H), 5.17 (dt, J=5.3, 2.5 Hz, 1H), 4.71 (ddd J= 7.7, 7.7, 5.9 Hz, 1H), 4.35 (dd, J=1.6, 4.9 Hz, 1H), 3.76 (dd, J=7.2, 7.2 Hz, 1H), 3.03 (s, 3H), 2.85 (brs, 1H), 2.57 (q, J= 10.6 Hz, 1H), 2.24 (d, J= 7.6 Hz, 1H), 2.2 (m, 1H), 2.08 (m, 1H), 2.06 (m, 1H), 2.05 (m, 1H), 2 (m, 2H), 1.96 (m, 1H), 1.81 (brs, 1H), 1.65 (m, 2H), 1.64 (m, 1H), 1.57 (m, 1H), 1.3 (s, 3H), 1.24 (s, 3H), 1.15 (s, 3H), (1.05 s, 6H), 0.98 (s, 3H), 0.92 (s, 3H), 0.67 (s, 3H). ^13^C NMR (100 MHz, CDCl_3_): δ= 144.8, 129.42, 126.1, 113.9, 90.2, 87.0, 81.6, 73.3, 72.0, 56.4, 52.2, 44.6, 44.0, 43.7, 43.5, 39.0, 38.6, 38.3, 37.9, 34.6, 33.9, 28.1, 27.9, 27.8, 26.7, 25.8, 25.7, 20.1, 18.8, 18.7, 16.5. HRMS (ESI): *m/z* 573.32873 ([M + Na]^+^) calcd for C_31_H_50_O_6_S.

#### Reaction of AG-01 with p-TsCl

To a stirring solution of AG-01 (30 mg, in pyridine), *p*-TsCl solution (200 mg) in pyridine was added. The reaction mixture was stirred at room temperature for 12 h. Reaction mixture quenched with water and extracted three times with 100 mL EtOAc. Crude product was purified by silica gel column chromatography (cyclohexane:EtOAc; 8:2) to give **AG-05** (13.9 mg lyophilized white-off powder). ^1^H NMR (400 MHz, CDCl_3_): δ =7.77 (dd, J=8.2, 1.7 Hz, 2H), 7.32 (dd, J=8.2, 1.7 Hz, 2H), 5.44 (dd, J=6.3, 1.7 Hz, 1H), 4.68 (q, J=6.8 Hz, 1H), 4.07 (m, 1H), 3.75 (td, J=7.4, 1.6 Hz, 1H), 2.69 (brs, 1H), 2.56 (q, J=10.4 Hz, 1H), 2.43 (s, 3H), 2.34 (d, J=5 Hz, 1H), 2.32 (brs, 1H), 2.19 (m, 1H), 2.17 (brs, 1H), 2.15 (m, 1H), 1.96 (m, 1H), 1.95 (m, 1H), 1.91 (m, 2H), 1.88 (d, J=4.24 Hz, 1H), 1.85 (d, J=1.5 Hz, 1H), 1.68 (m, 1H), 1.56 (dt, J=12.6 Hz, 1H), 1.43 (dd, J=12.9, 6.3 Hz, 1H), 1.29 (s, 3H), 1.27 (d, J=1.3 Hz, 1H), 1.22 (s, 3H), 1.14 (s, 3H), 1.04 (s, 3H), 0.90 (s, 3H), 0.84 (s, 6H). ^13^C NMR (100 MHz, _CDCl3_): 210.4, 144.6, 144.5, 134.6, 129.8, 129.8, 127.8, 127.8, 118.1, 89.4, 86.9, 81.4, 73.0, 72.0, 61.7, 56.3, 44.3, 44.3, 44.0, 43.8, 44.2, 44.2, 37.7, 37.4, 35.9, 34.5, 28.0, 27.8, 27.8, 26.8, 25.8, 25.1, 23.8, 21.8, 18.9, 18.2, 16.1. HRMS (ESI): 665.35653 *m/z* ([M + Na]^+^) calcd for C_37_H_54_O_7_S.

#### Reaction of AG with N-Acetylsulfanilyl chloride (N-Ac-Sulfa)

40 mg AG was dissolved in pyridine and 80 mg N-Ac-Sulfa (Acros Organics) reagent was added. Reaction was continued for 6 h at room temperature and quenched by addition of distilled water. Following the extraction via 50 ml ethyl acetate (3 times), the extract was evaporated at 60°C in the rotary evaporator. The residue was purified by silica gel column chromatography (cyclohexane:EtOAc; 7:3) to give **AG-06** (8.3 mg lyophilized white-off powder). ^1^H NMR (400 MHz, CDCl_3_): δ = 7.8 (d, J=8.4 Hz, 2H), 7.7 (d, J=8.8 Hz, 2H), 5.58 (d, 10.4 Hz, 1H), 5.53 (dt, J=10.4, 3 Hz, 1H), 5.13 (t, J=5.3 Hz, 1H), 4.69 (ddd, J=6.5, 6.5, 6.0 Hz, 1 H), 4.15 (dd, J=9.8, 6.6, 1H), 3.74 (t, J=7.1 Hz, 1H), 2.8 (brs, 1H), 2.55 (q, J=10.5 Hz, 1H), 2.18 (s, 1H), 2.12 (d, J=7.7 Hz, 1H), 2.0 (m, 2H), 1.99 (m, 2H), 1.91 (m, 1H), 1.9 (m, 2H), 1.69 (d, J=4.2), 1.59 (m, 1H), 1.55 (m, 2H), 1.53 (m, 1H), 1.28 (s, 3H), 1.22 (s, 3H), 1.14 (s, 3H), 0.97 (s, 3H), 0.95 (s, 3H), 0.83 (s, 3H), 0.73 (s, 3H), 0.62 (s, 3H). ^13^C NMR (100 MHz, CDCl_3_): δ= 169.3, 144.8, 143. 2, 131.8, 129.2, 129.0, 129.0, 126.1, 119.3, 119.3, 113.8, 90.9, 86.9, 81.5, 73.3, 72.0, 56.3, 52.1, 44.5, 43.8, 43.6, 43.4, 38.6, 38.2, 37.8, 34.5, 33.9, 28.0, 27.7, 27.6, 26.7, 25.8, 25.4, 24.8, 20.0, 18.8, 18.6, 16.5. HRMS (ESI): 692.36893 *m/z* ([M + Na]^+^) calcd for C_38_H_55_O_7_S.

#### Reaction of AG with Dansyl Chloride

100 mg AG was dissolved in pyridine and 450 mg dansyl chloride (400 mg) reagent was added. Reaction was continued for 6 h at 70 °C and quenched by addition of distilled water. Following the extraction via 500 ml ethyl acetate (3 times), the extract was evaporated at 60°C in the rotary evaporator. Crude product was purified by silica gel column chromatography (*n*-hexane:EtOAc; 7:3) to give **AG-07** (3.5 mg lyophilized white-off powder). ^1^H NMR (400 MHz, CDCl_3_): *δ*= 8.56 (d, J=8.5 Hz, 1H), 8.32 (d, J=8.6 Hz, 1H), 8.25 (dd, J=7.3, 1.3 Hz, 1H), 7.59 (dd, J= 8.7, 7.6 Hz, 1H), 7.52 (dd, J= 8.6, 7.3 Hz, 1H), 7.19 (dd, J= 7.6, 0.9 Hz, 1H), 5.56 (m, 2H), 5.10 (m, 1H), 4.69 (ddd, J= 7.9, 7,9, 6.0 Hz, 1H), 4.28 (dd, J= 11.3, 5.2 Hz, 1H), 3.75 (dd, J= 8.7, 6.5 Hz, 1H), 2.88 (d, J= 1.0 Hz, 6H), 2.79 (s, 1H), 2.55 (q, J= 10.5 Hz, 1H), 2.21 (d, J= 7.6 Hz, 1H), 2.04 (m, 2H), 2.0 (m, 1H), 1.97 (m, 1H), 1.93 (m, 1H), 1.82 (m, 1H),1.79 (m, 1H),1.7 (brs, 1H), 1.59 (m, 1H), 1.52 (m, 1H),1.51 (m, 2H), 1.29 (s, 3H), 1.25 (s, 3H), 1.23 (s, 3H), 1.14 (s, 3H), 0.95 (s, 3H), 0.86 (s, 3H), 0.71 (s, 3H), 0.62 (s, 3H). ^13^C NMR (100 MHz, CDCl_3_): δ= 151.8, 144.9, 133.7, 131.2, 130.2, 129.89, 129.4, 129.2, 128.6, 126.2, 123.2, 120.0, 115.5, 113.7, 91.5, 87.0, 81.5, 73.3, 72.0, 56.3, 52.1, 45.6, 45.6, 44.5, 44.0, 43.6, 43.5, 38.6, 38.1, 37.8, 34.6, 33.9, 27.9, 27.8, 27.6, 26.7, 25.7, 25.3, 20.0, 18.8, 18.6, 16.6. HRMS (ESI): 728.40520 *m/z* ([M + Na]^+^) calcd for C_42_H_59_NO_6_S.

#### Reaction of CG with p-TsCl

To a stirring solution of CG (200 mg, in pyridine) and 5 mg DMAP, *p*-TsCl solution (450 mg) in pyridine was added. The reaction mixture was stirred at reflux for 6 h. Reaction mixture quenched with water and extracted three times with 100 mL EtOAc. Crude product was purified on a silica gel chromatography column (cyclohexane:EtOAc; 8:2) to give **CG-02** (8.6 mg lyophilized white-off powder) and **CG-03** (95.1 mg lyophilized white-off powder). For **CG-02**; ^1^H NMR (400 MHz, CDCl_3_): *δ*=5.63 (d, J=10.6, 1H), 5.48 (brs, 1H), 4.69 (q, J=7.2, 1H), 3.74 (dd, J=7.1, 7.1, 1H), 3.32 (dd, J=11.4, 4.6, 1H), 2.71 (d, J=7.9, 1H), 2.59 (q, J=10.6, 1H), 2.27 (d, J=7.6, 1H), 2.0 (m, 2H), 1.88 (m, 1H), 1.85 (m, 1H), 1.84 (m, 1H), 1.82 (m, 1H), 1.62 (m, 1H), 1.6 (m, 1H), 1.59 (m, 1H), 1.52 (m, 1H), 1.42 (m, 1H), 1.41 (m, 1H), 1.40 (m, 1H), 1.31 (s, 3H), 1.25 (s, 1H), 1.23 (s, 6H), 1.22 (s, 3H), 1.15 (s, 3H), 1.05 (s, 2H), 0.78 (s, 3H), 0.74 (m, 1H), −0.16 (d, J=4.3, 1H). ^13^C NMR (100 MHz, CDCl_3_): δ=129.2, 126.6, 87.3, 81.4, 78.5, 73.5, 72.1, 56.4, 48.5, 46.6, 45.2, 43.6, 43.4, 40.3, 34.6, 33.4, 30.1, 29.8, 28.4, 28.1, 27.9, 26.7, 25.9, 25.6, 25.2, 20.6, 18.7, 18.2, 18.1, 14.5. HRMS (ESI): *m/z* 573.32873 ([M + Na]^+^) calcd for C_30_H_48_O_4._ For **CG-03**; ^1^H NMR (400 MHz, CDCl_3_): δ= 7.79 (d, J= 8.2 Hz, 2H), 7.32 (d, J= 8.2 Hz, 2H), 5.51 (d, J= 10.7 Hz, 1H), 5.46 (dd, J= 10.7, 6, 3 Hz, 1H), 4.68 (ddd, J= 7.7, 7.7, 6.1 Hz, 1H), 4.31 (dd, J=11.8, 4.6 Hz, 1H), 3.74 (t, J= 7.1 Hz, 1H), 2.68 (dd, J= 6.0, 2.5 Hz, 1H), 2.58 (q, J=10.5 Hz, 1H), 2.43 (s, 3H), 2.26 (d, J= 7.7 Hz, 1H), 2.0 (td, J= 10.5, 9.5 Hz, 2H), 1.89 (m, 1H), 1.86 (m, 1H), 1.85 (m, 1H), 1.79 (m, 1H), 1.77 (m, 1H), 1.58 (m, 1H), 1.57 (m, 1H), 1.56 (m, 1H), 1.48 (m, 1H), 1.41 (m, 1H), 1.38 (m, 1H), 1.34 (m, 1H), 1.29 (s, 3H), 1.21 (s, 3H), 1.19 (s, 3H), 1.13 (s, 3H), 0.81 (s, 3H), 0.8 (s, 3H), 0.72 (d, J= 4.2 Hz, 1H), 0.71 (s, 3H), −0.17 (d, J= 4.2 Hz, 1H). ^13^C NMR (100 MHz, CDCl_3_): δ= 144.5, 134.9, 129.8, 127.8, 125.7, 90.2, 87.2, 81.3, 73.4, 72.0, 56.4, 48.4, 46.7, 45.1, 43.5, 43.3, 40.0, 34.5, 33.3, 29.5, 28.2, 27.9, 27.8, 26.7, 25.9, 25.4, 25.2, 21.8, 20.7, 18.6, 18.2, 18.1, 15.4. HRMS (ESI): *m/z* 649.36046 [M + Na]^+^ calcd for C_37_H_54_O_6_S.

#### Reaction of CG with MsCl

MsCl (140 µL) and TEA (294 µL) was added to a solution of 350 mg CG in dichloromethane. The reaction mixture was stirred at 0 °C for 3 h. The reaction solvent was evaporated under reduced pressure and the residue was precipitated in methanol to give **CG-04** (193 mg lyophilized white-off powder). ^1^H NMR (400 MHz, CDCl_3_): δ= 5.58 (d, J= 10.6 Hz, 1H), 5.49 (ddd, J= 10.6, 6.1, 3.1 Hz, 1H), 4.7 (ddd, J=7.7, 7.7, 6.0 Hz, 1H), 4.43 (dd, J= 11.9, 4.6 Hz, 1H), 3.73 (t, J= 7.1 Hz, 1H), 3.01 (s, 3H), 2.71 (dd, J= 6.2, 2.6 Hz, 1H), 2.58 (d, J=10.6 Hz, 1H), 2.27 (d, J=7.6 Hz, 1H), 2.12 (dd, J=12.5, 3.9 Hz, 1H), 2 (m, 2H), 1.96 (m, 1H), 1.9 (m, 1H), 1.89 (m, 1H), 1. 81 (m, 1H), 1.68 (m, 1H), 1.61 (m, 1H), 1.6 (m, 1H), 1.51 (m, 1H), 1.47 (m, 1H), 1.43 (m, 1H), 1.38 (m, 1H), 1.29 (s, 3H), 1.22 (s, 3H), 1.20 (s, 3H), 1.14 (s, 3H), 1.05 (s, 3H), 0.86 (s, 3H), 0.77 (d, J= 3.6 Hz, 1H), 0.74 (s, 3H), - 0.12 (d, J=4.3 Hz, 1H). ^13^C NMR (100 MHz, CDCl_3_): δ= 129.9, 125.6, 89.7, 87.2, 81.3, 73.4, 72.0, 56.4, 48.4, 46.7, 45.1, 43.5, 43.3, 40.0, 38.9, 34.5, 33.3, 29.5, 28.2, 28.2, 27.9, 27.8, 26.7, 25.9, 25.8, 25.2, 20.8, 18.6, 18.2, 18.1, 15.2. HRMS (ESI): *m/z* 573.32624 [M+Na]^+^ calcd for C_31_H_50_O_6_S.

#### Reaction of CG-01 with p-TsCl

100 mg CG was dissolved in pyridine and 300 mg *p-*TsCl reagent was added. Reaction was continued for 12 h at room temperature and quenched by addition of distilled water. Following the extraction via 500 ml ethyl acetate (3 times), the extract was evaporated at 60°C in the rotary evaporator. Crude product was purified on a silica gel chromatography column (cyclohexane:EtOAc; 7:3) to give **CG-05** (33 mg lyophilized white-off powder). ^1^H NMR (400 MHz, CDCl_3_): δ= 7.79 (d, J= 8.3 Hz, 2H), 7.3 (d, J= 8.3 Hz, 2H), 4.69 (ddd, J= 7.8, 7.8, 6.1 Hz, 1H), 4.21 (m, 1H), 3.75 (dd, J= 8.3, 6.1 Hz, 1H), 2.66 (dd, J= 8.5, 4.0 Hz, 1H), 2.57 (q, J= 10.8 Hz, 1H), 2.43 (s, 3H), 2.32 (d, J= 7.6 Hz, 1H), 2.3 (brs, 1H), 2.17 (m, 1H), 2.12 (m, 1H), 1.98 (m, 1H),1.94 (m, 1H), 1.85 (m, 1H), 1.84 (m, 1H), 1.79 (m, 1H), 1.76 (m, 1H), 1.6 (m, 1H), 1.47 (m, 1H), 1.46 (m, 1H), 1.37 (m, 1H), 1.21 (m, 1H), 1.21 (s, 3H), 1.57 (m, 1H), 1.42 (m, 1H), 1.3 (s, 3H), 1.14 (s, 3H), 1.02 (s, 3H), 1.0 (s, 3H), 0.89 (s, 3H), 0.21 (d, J= 5.5 Hz, 1H), 0.6 (d, J= 5.5 Hz, 1H). ^13^C NMR (100 MHz, CDCl_3_): δ= 210, 144.6, 134.7, 129.8, 129.8, 127.8, 127.8, 89.3, 87.1, 81.3, 72.9, 72.1, 57.2, 56.9, 47.1, 45.3, 43.8, 42.4, 41.2, 40.0, 34.5, 33.0, 30.0, 29.7, 28.1, 27.8, 27.3, 26.8, 26.5, 26.3, 25.9, 22.2, 21.8, 21.7,19.1, 18.4, 14.8. HRMS (ESI): *m/z* 665.35109 [M+Na]^+^ calcd for C_37_H_54_O_7_S.

#### Reaction of CG with N-Ac-Sulfa

50 mg CG was dissolved in pyridine and 150 mg N-Ac-Sulfa (Acros Organics) reagent was added. Reaction was continued for 11 h at room temperature and quenched by addition of distilled water. Following the extraction via 50 ml ethyl acetate (3 times), the extract was evaporated at 60 °C in the rotary evaporator. The residue was purified by silica gel column chromatography (CHCl_3_:MeOH; 95:5) to give **CG-06** (12.1 mg). ^1^H NMR (400 MHz, CDCl_3_): δ= 7.82 (d, J= 8.5 Hz, 2H), 7.3 (d, J= 8.5 Hz, 2H), 5.50 (d, J= 10.7 Hz, 1H), 5.43 (ddd, J= 0.2, 6.2, 2.9 Hz, 1H), 4.67 (q, J= 4.2 Hz, 1H)), 4.26 (dd, J= 11.8, 4.7 Hz, 1H), 3.73 (dd, J= 6.9 Hz, 1H), 2.66 (m, 1H), 2.57 (q, J= 10.8 Hz, 1H), 2.26 (d, J= 7.5, Hz, 1H), 2.17 (s, 3H), 1.98 (m, 2H), 1.87 (m, 1H), 1.85 (m, 1H), 1.78 (m, 1H), 1.77 (m, 1H), 1.57 (m, 1H), 1.56 (m, 1H), 1.56 (m, 1H), 1.37 (m, 1H), 1.33 (m, 1H), 1.85 (m, 1H), 1.48 (m, 1H), 1.40 (m, 1H), 1.27 (s, 3H), 1.18 (s, 3H), 1.18 (s, 3H), 1.13 (s, 3H), 0.8 (s, 3H), 0.78 (s, 3H), 0.69 (s, 3H), 0.69 (d, J= 4.1 Hz, 1H), −0.18 (d, J= 4.1 Hz, 1H). ^13^C NMR (100 MHz, CDCl_3_): δ=169.3, 142.2, 131.8, 129.7, 129.0, 129.0, 125.6, 119.3, 119.3, 90.4, 87.1, 81.3, 73.4, 72.0, 56.3, 48.3, 46.6, 45.0, 43.4, 43.2, 40.0, 34.5, 33.2, 29.5, 28.2, 27.8, 27.7, 27.7, 26.7, 25.9, 25.4, 25.1, 24.7, 20.7, 18.6, 18.1, 18.1, 15.3. HRMS (ESI): *m/z* 692.36865 [M + Na]^+^ calcd for C_38_H_57_NO_8_S.

#### Reaction of SCG with TsCl

500 mg SCG was dissolved in pyridine and 1000 mg *p*-TsCl reagent was added. The reaction mixture was stirred at reflux for 6 h and quenched by addition of distilled water. Following the extraction via 500 ml ethyl acetate (3 times), the extract was evaporated at 60 °C in the rotary evaporator. Crude product was purified on a silica gel chromatography column (*n*-hexane:EtOAc; 90:10) to give **SCG-01** (15.4 mg lyophilized white-off powder), **SCG-02** (8.6 mg lyophilized white-off powder), **SCG-03** (13 mg lyophilized white-off powder), **SCG-04** (19.2 mg lyophilized white-off powder), **SCG-05** (15.2 mg lyophilized white-off powder). For **SCG-01**; ^1^H NMR (400 MHz, CDCl_3_): δ=7.75 (d, J= 8.2 Hz, 2H), 7.31 (d, J= 8.2 Hz, 2H), 1.2 (d, J= 3.2 Hz, 1H), 5.06 (ddd, J= 15.4, 7.9, 1.4 Hz, 1H), 3.51 (ddd, J= 9.1, 9.1, 4.2 Hz, 1H), 3.29 (dd, J= 11.3, 4.6 Hz, 1H), 2.44 (s, 3H), 1.92 (m, 1H), 1.88 (m, 2H), 1.86 (m, 1H), 1.78 (m, 1H), 1.64 (d, J= 1.4 Hz, 1H), 1.6 (m, 2H), 1.57 (m, 1H), 1.56 (m, 1H), 1.44 (m, 1H), 1.4 (m, 1H), 1.34 (d, J= 1.97 Hz, 1H), 1.3 (m, 1H), 0.95 (s, 3H), 1.25 (m, 1H), 1.22 (s, 3H), 1.04 (s, 3H), 0.92 (s, 3H), 0.45 (d, J= 4.6 Hz, 1H), 0.28 (d, J= 4.6 Hz, 1H). ^13^C NMR (100 MHz, CDCl_3_): δ= 144.6, 134.4, 129.9, 129.9, 127.8, 127.8, 82.9, 78.4, 68.5, 53.5, 46, 45.9, 45.8, 44.5, 44.4, 41.6, 37.5, 32.0, 30.3, 30.3, 30.0, 29.8, 28.0, 26.1, 25.0, 21.8, 20.8, 19.8, 15.3. HRMS (ESI): *m/z* 525.2699 [M + Na]^+^ calcd for C_29_H_42_O_5_S. For **SCG-02;** ^1^H NMR (400 MHz, CDCl_3_): δ=5.62 (d, J= 10.5 Hz, 1H), 5.44 (ddd, J= 10.6, 6.1, 3.2 Hz, 1H), 4.55 (ddd, J= 14.5, 7.7, 1.4 Hz, 1H), 3.3 (dd, J= 11.2, 4.4 Hz, 1H), 2.48 (dd, J= 6.2, 2.6 Hz, 1H), 2.05 (dd, J= 13.7, 8.2 Hz, 1H), 1.89 (m, 1H), 1.86 (m, 1H), 1.85 (m, 1H), 1.82 (m, 1H), 1.69 (m, 1H), 1.62 (m, 1H), 1.6 (m, 1H), 1.46 (m, 1H), 1.42 (m, 1H), 1.27 (m, 1H), 1.25 (m, 1H), 1.22 (m, 1H), 1.05 (s, 3H), 0.96 (s, 3H), 0.92 (s, 3H), 0.77 (s, 3H), 0.73 (d, J= 4.5 Hz, 1H), −0.15 (d, J= 4.1 Hz, 1H). ^13^C NMR (100 MHz, CDCl_3_): δ= 129.1, 126.3, 78.5, 72.2, 48.5, 47.8, 46.5, 45.4, 45.3, 43.7, 40.3, 31.4, 30.1, 29.8, 28.3, 25.6, 25.2, 22.2, 21.2, 18.5, 18.4, 14.5. HRMS (ESI): *m/z* 331.26321 [M+H]^+^ calcd for C_22_H_34_O_2_. For **SCG-03**; ^1^H NMR (400 MHz, CDCl_3_): δ= 7.74 (dd, J= 8.2, 3.6 Hz, 2H), 7.31 (dd, J= 8.2, 3.6 Hz, 2H), 5.49 (ddd, J= 9.9, 5.8, 2.0 Hz, 1H), 5.3 (dd, J= 9.8, 2.7 Hz, 1H), 5.04 (q, J= 7.4 Hz, 1H), 3.45 (td, J=9.8, 4.7 Hz, 1H), 2.44 (s, 3H), 2.30 (m, 1H), 2.07 (m, 1H), 1.90 (m, 1H), 1.88 (m, 1H), 1.87 (m, 1H), 1.65 (m, 1H), 1.62 (m, 1H),1.58 (m, 1H), 1.54 (m, 1H), 1.47 (m, 1H), 1.39 (m, 1H), 1.34 (m, 1H), 1.23 (s, 3H), 1.1 (m, 1H), 1.05 (s, 3H), 1.03 (s, 3H), 0.99 (s, 3H), 0.51 (d, J= 5.0 Hz, 1H), 0.34 (d, J= 4.5 Hz, 1H). ^13^C NMR (100 MHz, CDCl_3_): δ= 144.6, 140.7, 134.2, 129.9, 129.9, 127.8, 127.8, 122.8, 83.0, 70.7, 52.6, 47.9, 46.1, 45.9, 44.9, 44.4, 38.0, 34.9, 33.1, 31.7, 30.2, 28.4, 28.2, 25.8, 25.7, 23.5, 21.8, 20.2, 19.2. HRMS (ESI): *m/z* 507.2578 [M + Na]^+^ calcd for C_29_H_40_O_4_S. For **SCG-04**; ^1^H NMR (400 MHz, CDCl_3_): δ= 7.8 (d, J=7.7 Hz, 2H), 7.75 (d, J= 7.8 Hz, 2H), 7.32 (d, J= 7.8 Hz, 4H), 5.52 (d, J= 10.7 Hz, 1H), 5.36 (ddd, J= 9.6, 5.9, 2.9 Hz, 1H), 5.06 (q, J= 7.5 Hz, 1H), 4.30 (dd, J= 11.6, 4.5 Hz, 1H), 2.44 (s, 6H), 2.41 (m, 1H), 2.0 (m, 1H), 1.9 (m, 1H), 1.89 (m, 1H), 1.81 (m, 2H), 1.8 (m, 1H), 1.8 (m, 1H), 1.61 (m, 1H), 1.6 (m, 1H), 1.5 (m, 1H), 1.4 (m, 1H), 1.39 (m, 1H), 1.16 (m, 1H), (4.1), 0.88 (s, 3H), 0.83 (s, 3H), 0.82 (s, 3H), 0.79 (s, 3H), 0.71 (d, J= 4.1 Hz, 1H), −0.16 (d, J= 4.1 Hz, 1H). ^13^C NMR (100 MHz, CDCl_3_): δ= 144.7, 144.4, 134.7, 134.2, 128.9, 129.7, 129.7, 129.7, 129.7, 127.7, 127.7, 127.7, 127.7, 125.7, 89.9, 82.4, 48.1, 46.5, 43.1, 44.9, 44.4, 41.9, 39.8, 30.7, 29.4, 27.7, 27.6, 25.3, 24.8, 21.8, 21.6, 21.6, 20.9, 18.3, 17.6, 15.2. HRMS (ESI): *m/z*.656.2970 [M + NH_4_]^+^ calcd for C_36_H_46_O_6_S_2._ For **SCG-05**; ^1^H NMR (400 MHz, CDCl_3_): δ= 7.76 (d, J= 8.1 Hz, 2H), 7.32 (d, J= 7.9 Hz, 2H), 5.61 (d, J= 10.5 Hz, 1H), 5.37 (m, 1H), 5.07 (dd, J= 14.7, 7.2 Hz, 1H), 3.33 (dd, J= 10.8, 3.9 Hz, 1H), 2.44 (m, 4H), 2.01 (m, 1H), 1.86 (d, J= 2.4 Hz, 1H), 1.83 (m, 2H), 1.82 (m, 1H), 1.81 (d, J= 5.6 Hz, 1H), 1.63 (d, J= 12.9 Hz, 1H), 1.62 (m, 1H), 1.53 (d, J= 14.7 Hz, 1H), 1.43 (m, 1H), 1.42 (m, 1H), 1.23 (s, 3H), 1.2 (dd, J= 12.7, 4.3 Hz, 1H), 1.05 (s, 3H), 0.9 (s, 3H), 0.85 (s, 3H), 0.77 (s, 3H), 0.72 (d, J= 3.1 Hz, 1H), −0.15 (d, J= 3.9 Hz, 1H). ^13^C NMR (100 MHz, CDCl_3_): δ= 144.5, 134.5, 129.9, 129.9, 128.5, 127.8, 127.8, 126.9, 82.7, 78.5, 48.4, 46.5, 45.2, 44.6, 43.3, 42.1, 40.4, 31.0, 30.1, 29.8, 28.3, 25.6, 25.0, 21.8, 21.6, 21.1, 18.5, 17.9, 14.5. HRMS (ESI): *m/z* 502.2940 ([M + NH_4_]^+^) calcd for C_29_H_40_O_4_S.

#### Reaction of SCG with MsCl

130 µL MsCl was added to a solution of SCG (150 mg) in pyridine. The reaction mixture was stirred at reflux for 6 h. The reaction solvent was then removed under reduced pressure, and the residue was purified by silica gel column chromatography (n-hexane:EtOAc; 90:10) to give **SCG-06** (13 mg lyophilized white-off powder), and **SCG-07** (4.3 mg lyophilized white-off powder). For **SCG-06**, ^1^H NMR (400 MHz, CDCl_3_): δ=5.61 (d, J=10.5 Hz, 1H), 5.45 (ddd, J= 10.3, 6.1, 3.0 Hz, 1H), 5.29 (q, J= 7.6 Hz, 1H), 4.45 (dd, J= 12.0, 4.6 Hz, 1H), 3.03 (s, 3H), 2.97 (s, 3H), 2.51 (dd, J= 6.0, 2.6 Hz, 1H), 2.2 (m, 1H), 2.14 (m, 1H), 2.04 (m, 1H), 2.0 (m, 1H), 1.93 (m, 1H), 1.92 (m, 1H), 1.92 (m, 1H),1.89 (m, 1H), 1.70 (m, 1H), 1.69 (m, 1H), 1.63 (m, 1H), 1.43 (m, 1H), 1.27 (dd, J= 13, 5.1 Hz, 1H), 1.10 (s, 3H), 0.98 (s, 3H), 0.87 (s, 3H), 0.86 (s, 3H), 0.77 (d, J= 4.3 Hz, 1H), −0.07 (d, J= 4.3 Hz, 1H). ^13^C NMR (100 MHz, CDCl_3_): *δ*= 129.2, 125.9, 89.5, 82.0, 48.4, 46.7, 45.2, 44.7, 43.2, 42.3, 40.0, 39.0, 38.5, 30.9, 29.5, 28.1, 27.8, 25.8, 25.0, 21.2, 22.0, 18.4, 17.9, 15.3. HRMS (ESI): *m/z* 995.3974 [2M + Na]^+^ calcd for C_24_H_38_O_6_S_2_. **For SCG-07**, ^1^H NMR (400 MHz, CDCl_3_): δ= 5.26 (q, J= 8.0 Hz, 1H), 3.56 (ddd, J= 9.2, 9.2, 4.1 Hz, 1H), 3.22 (dd, J= 11.2, 4.2 Hz, 1H), 2.97 (s, 3H,) 2.11 (m, 1H), 2.06 (m, 1H), 2.01 (m, 1H), 1.97 (m, 1H), 1.8 (m, 1H), 1.79 (m, 1H), 1.71 (m, 1H), 1.63 (m, 1H), 1.58 (m, 1H), 1.55 (m, 1H), 1.48 (m, 1H), 1.38 (m, 1H), 1.37 (m, 1H), 1.31 (m, 1H), 1.26 (m, 1H), 1.23 (s, 3H), 1.10 (s, 3H), 1.03 (s, 3H), 0.94 (s, 3H), 0.5 (d, J= 4.7 Hz, 1H), 0.31 (d, J= 4.7 Hz, 1H). ^13^C NMR (100 MHz, CDCl_3_): δ=82.4, 78.5, 68.4, 53.4, 46.2, 46.0, 45.6, 44.7, 44.6, 41.6, 37.6, 31.3, 30.3, 30.2, 29.0, 27.9, 26.7, 26.1, 24.9, 20.7,19.8, 38.5, 15.3. HRMS (ESI): *m/z* 449.2366 ([M + NH_4_]^+^) calcd for C_23_H_38_O_5_S.

### 5.2 Biological Studies

#### 5.2.1. Chemicals

Pitstop II (Sigma; SML1169), methyl β cyclodextrin (Cayman; 21633), chlorpromazine (Cayman; 16129), dynasore (Cayman,14062) and cytochalasin D (Cayman; 11330) were used as endocytosis inhibitors.

Derivatives dissolved in anhydride DMSO as a 1000-fold concentrated stock. The dissolved compounds immediately used for the treatment of cells.

#### 5.2.2. Cell culture

HCC1937 (Human breast cancer line), HeLa (Human endometrial carcinoma), A549 (Human lung adenocarcinoma), MRC-5 (Human lung fibroblasts), MCF7 (Human breast cancer cells), A549 (human lung carcinoma) were obtained from American Type Culture Collection and maintained as exponentially growing monolayers by culturing according to the supplier’s instructions. MCF7, HeLa, MRC-5 and A549 cell lines were cultured and routinely passaged in DMEM media containing 10% FBS, while HCC1937 cell lines were propagated in RPMI 1640 containing 10% FBS.

#### 5.2.3. Cytotoxicity analysis

Following the treatment with compounds or vehicle for 48 h, the 10% WST-1 (Roche, Switzerland) in medium was replaced in each 96 well. After 4 h incubation with WST-1 reagent at 37 °C and 5% CO_2_, thw absorbance was measured by using a microplate reader at 440 nm (Varioscan, Thermo Fisher Scientific, US). Graph Pad Prism 5 (San Diego, CA, US.) was used to calculate the IC_50_, which represents the concentration of compounds that is required for 50% inhibition in comparison to the vehicle-treated controls. The experiments were repeated three times independently.

#### 5.2.4. LDH (Lactate dehydrogenase) releasing assay

LDH-Cytotoxicity Colorimetric Assay Kit II (Biovision) was used to detect LDH releasing. Briefly, HCC1937 were seeded on 96 well plate. After treatment with compounds, cells were centrifuged at 600xg for 10 min. 10 µl supernatant were transferred into clean 96 well plate and added into ?? 100 µl LDH Reaction Mix. Following 30 min incubation at room temperature, 10 µl stop solution were added and absorbance were measured at 450 nm (Varioscan, Thermo Fisher Scientific, US).

#### 5.2.5. Immunofluorescence studies

Cells were grown on glass coverslips and treated with AG-08. At the end of treatments, cells were washed twice with ice cold PBS and fixed with 4% paraformaldehyde in PBS for 30 min at 4 °C. After washing 6 times with PBS, cells were permeabilized and blocked with 0,01 % saponin and 0,01% BSA in PBS. Then fixed cells were incubated with primary antibodies for 1 h at 37 °C. Antibodies were used at the following dilutions: Calnexin (Sigma Aldrich-C7617, UK), 1:200; EEA1 (CST-3288, US), 1:100. Cells were then incubated with secondary antibodies (1:400) for 1 h at 37 °C. Mounted samples were analyzed by fluorescence microscopy (Olympus IX70, Japan).

#### 5.2.6. Total RNA isolation and Expression Analysis by Quantitative RT-PCR

The total RNA was isolated using Total RNA Isolation Kit (Bio-Rad, US) following the manufacturer’s instructions. cDNAs were synthesized using The iScript cDNA synthesis kit (Bio-Rad, US) according to the manufacturer’s instructions. To gene expression analysis, specific primers were designed against CHOP (forward: AGTCTAAGGCACTGAGCGTATCAT, reverse: CTTTCAGGTGTGGTGATGTATGAA). Quantitative RT-PCR (qRT-PCR) was performed using The SYBR Green I Mastermix (Bio-Rad, US) and LightCycler480 thermocycler (Roche). Fold change for the transcripts were normalized to the housekeeping gene TBP1 (TATA-Box Binding Protein1) (forward: GAGTTCTGGGATTGTACCGCA, reverse: CGTGGTTCGTGGCTCTCTT). For relative quantification, reaction efficiency incorporated ΔΔCq formula was used. Six independent biological replicates with two technical replicates per experiment were used for each PCR.

#### 5.2.7. Western blot analysis

Cell lysates were prepared by RIPA buffer (1XPBS, 1% nonidet P-40, 0.5% sodium deoxycholate, and 0.1% SDS, pH 8.0). Protein concentrations were determined by bicinchoninic acid (BCA) protein assay (Thermo Fisher Scientific, US). After equal amounts of proteins were loaded to the gels, proteins were separated by SDS-PAGE electrophoresis and transferred to PVDF membranes (EMD Millipore, Thermo Fisher Scientific, US).

Following the classic immunoblotting steps (blocking, incubating with primary and secondary antibodies), chemiluminescence signals were detected using Clarity ECL substrate solution (Bio-Rad, US) by Fusion-FX7 (Vilber Lourmat, Thermo Fisher Scientific, US). Monoclonal antibodies used in this study were anti-LC3 (CST-12741, USA), anti-Atg-5/12 (CST-12994, USA), anti-Atg-7 (CST-8558, US), anti-actin (Sigma-Aldrich-A5316, UK), anti-caspase-3 (CST-9665, US), Anti-CHOP (CST-2895, UK), The experiments were repeated three times independently; with one representative result shown.

#### 5.2.8. Microaaray analysis

Microarray analyses were conducted using the human Clariom D Affymetrix platform following the manufacturer’s instructions (902915, Thermo Fisher Scientific). Briefly, the total RNA was isolated using Total RNA Isolation Kit (Bio-Rad, US). Isolated total RNA was amplified, labeled, and hybridized on Clariom D Affymetrix platform.The arrays were stained by using GeneChip Fluidics Station 450. The cartridge was scanned with the GeneChip Scanner 3000 7G System (Affymetrix CA, USA)

#### 5.2.9. Protein-protein interaction network and enrichment

Protein network analysis was performed using with Search Tool for the Retrieval of Interacting Genes/Proteins (STRING 11; https://string-db.org/). The confidence score for selection was ≥0.4. Disconnected nodes in the network were hidden to simplify the display.

The Metascape online tool was used to thoroughly analyze enrichment information (Metascape; www.metascape.org/). DE gene lists were supplied to the Metascape tool to produce enriched GO terms, KEGG pathways. Terms with P < 0.01 were considered significantly enriched. Additionally, Reactome database was used for additional enrichment analysis (https://reactome.org/)^[34]^.

### 5.3. Self-aggregation molecular dynamics simulations

All simulations were conducted using Gromacs version 2018.1[35]. AG and AG-08 molecules were first defined in mol2 file format, and submitted for ligand parametrization at the Automated Topology Builder (ATB) server[36]. For each simulation, a pre-defined amount of molecules were first randomly placed within a pre-defined box, and the system was solvated. For each simulation, a minimization was for a maximum of 50,000 steps until the maximum force was below 10 kJ/mol. Here, cutoff values of 1.2, 1.4 and 1.4 nm were used for neighbor lists, and as Coulomb and van-der Waals interactions, respectively. Particle Mesh Ewald (PME) method was used for treatment of long-range electrostatic interactions. Following energy minimization, the systems were subjected to NVT and NPT equilibration for 50,000 steps each at 300 K and 1 bar using the same cutoff values and settings as in minimization. Finally, NPT simulations were conducted for 250ns ns to test self-aggregation behavior.

### 5.4. Characterization of particles

#### 5.4.1. STEM Microscopy

Firstly, AG-08 was dissolved in DMSO a 1000-fold concentrated stock and immediately added in filtered PBS. Then solution was placed on grids (formvar/carbon-coated 400 mesh copper grids). After completely dry, images were taken via scanning transmission electron microscope (STEM-Zeiss Sigma 500). Additionally, 1% uranyl acetate (pH 7.4) was used to achieve picture with negative staining. After washing of uranyl acetate via water, images were recorded via STEM.

#### 5.4.2. Measurement of Particle Diameter via Zeta Sizer

AG-08 was dissolved in DMSO a 1000-fold concentrated stock and added into the filtered PBS. Then particle diameter was immediately measured via zeta-sizer (DLS– Particulate Systems). Each measurement was repeated in triplicate.

#### 5.4.3. Nile red staining

AG-08 and AG was dissolved in DMSO a 1000-fold concentrated stock added into the filtered PBS which contain 2.5 µM Nile red. After 3 h incubation at room temperature, the fluorescence intensity was measured with a microplate reader (550 nm for excitation and 635 nm for the emission wavelength) (Varioscan, Thermo Fisher Scientific, US).

### 5.5. Statistical analysis

The statistical significance of differences between groups was assessed by two-tailed equal variance Student’s t-test or one-way ANOVA using GraphPad Prism software.

## Acknowledgement

We are very grateful to Bionorm Natural Products for donating CG, AG-01, CG-01 and SCG, and Doç. Dr. Rükan Genç Altürk for donating Nile red. Also, we thank to the Pharmaceutical Sciences Research Centre (FABAL, Ege University, Faculty of Pharmacy) and Biotechnology and Bioengineering Application and Research Centre (BIYOMER, İzmir Institute of Technology) for equipmental support. Additionally, we thank to Izmir Biomedicine and Genome Center (IBG) for STEM microscope analysis.

This project was supported by The Scientific and Technological Research Council of Turkey (TUBITAK, Project No: 118S709) and Scientific Research Projects Coordination Unit of Izmir Institute of Technology (IZTECH BAP, Project No: 2017IYTE71.

## Contributor Roles

G.Ü., P.B.K and E.B. formed the hypothesis and designed the study. G.Ü. participated in the experimental design and conducted the experiments and collected data. O.S. designed and performed molecular dynamics studies. All authors analyzed the results and contributed to the manuscript preparation. All authors read and approved the manuscript.

## References

1. Uner, G.; Tag, O.; Erzurumlu, Y.; Kirmizibayrak, P. B.; Bedir, E.; Chemical Research in Toxicology 2020, 33 (11), 2880–2891.

2. Coan, K. E.; Maltby, D. A.; Burlingame, A. L.; Shoichet, B. K.; Journal of medicinal chemistry 2009, 52 (7), 2067–2075.

3. Ganesh, A. N.; Donders, E. N.; Shoichet, B. K.; Shoichet, M. S.; Nano Today 2018, 19, 188-200. DOI 10.1016/j.nantod.2018.02.011.

4. Coan, K. E.; Shoichet, B. K.; Journal of the American Chemical Society 2008, 130 (29), 9606–9612.

5. Shaham-Niv, S.; Adler-Abramovich, L.; Schnaider, L.; Gazit, E.; Science advances 2015, 1 (7), e1500137.

6. Anand, B. G.; Prajapati, K. P.; Dubey, K.; Ahamad, N.; Shekhawat, D. S.; Rath, P. C.; Joseph, G. K.; Kar, K.; ACS Nano 2019, 13 (5), 6033–6049. DOI 10.1021/acsnano.9b02284.

7. McLaughlin, C. K.; Duan, D.; Ganesh, A. N.; Torosyan, H.; Shoichet, B. K.; Shoichet, M. S.; ACS Chem Biol 2016, 11 (4), 992–1000. DOI 10.1021/acschembio.5b00806.

8. Julien, O.; Kampmann, M.; Bassik, M. C.; Zorn, J. A.; Venditto, V. J.; Shimbo, K.; Agard, N. J.; Shimada, K.; Rheingold, A. L.; Stockwell, B. R.; Weissman, J. S.; Wells, J. A.; Nat Chem Biol 2014, 10 (11), 969–76. DOI 10.1038/nchembio.1639.

9. Kuang, Y.; Xu, B.; Angewandte Chemie 2013, 125 (27), 7082–7086.

10. Dash, S. K.; Chattopadhyay, S.; Dash, S. S.; Tripathy, S.; Das, B.; Mahapatra, S. K.; Bag, B. G.; Karmakar, P.; Roy, S.; Bioorg Chem 2015, 63, 85–100. DOI 10.1016/j.bioorg.2015.09.006.

11. Ikeda, Y.; Murakami, A.; Fujimura, Y.; Tachibana, H.; Yamada, K.; Masuda, D.; Hirano, K.; Yamashita, S.; Ohigashi, H.; J Immunol 2007, 178 (8), 4854–64. DOI 10.4049/jimmunol.178.8.4854.

12. Ceru, S.; Kokalj, S. J.; Rabzelj, S.; Skarabot, M.; Gutierrez-Aguirre, I.; Kopitar-Jerala, N.; Anderluh, G.; Turk, D.; Turk, V.; Zerovnik, E.; Amyloid 2008, 15 (3), 147–59. DOI 10.1080/13506120802193555.

13. Liang, Y.; Sun, Y.; Fu, X.; Lin, Y.; Meng, Z.; Meng, Y.; Niu, J.; Lai, Y.; Sun, Y. J. A. c.; nanomedicine,; biotechnology, 2020, 48 (1), 525–532.

14. Kunitake, T.; Okahata, Y.; Shimomura, M.; Yasunami, S.; Takarabe, K. J. J. o. t. A. C. S.; 1981, 103 (18), 5401–5413.

15. Ghattas, M. A.; Bryce, R. A.; Al Rawashdah, S.; Atatreh, N.; Zalloum, W. A.; Journal of medicinal chemistry 2018, 13 (6), 500–506.

16. Ghattas, M. A.; Al Rawashdeh, S.; Atatreh, N.; Bryce, R. A.; Modeling, Journal of Chemical Information 2020, 60 (8), 3901–3909.

17. Jubb, H. C.; Higueruelo, A. P.; Ochoa-Montaño, B.; Pitt, W. R.; Ascher, D. B.; Blundell, T. L.; Journal of molecular biology 2017, 429 (3), 365–371.

18. Dutta, D.; Donaldson, J. G.; Cellular logistics 2012, 2 (4), 203–208.

19. Willox, A. K.; Sahraoui, Y. M.; Royle, S. J.; Biology open 2014, 3 (5), 326–331.

20. Dutta, D.; Williamson, C. D.; Cole, N. B.; Donaldson, J. G.; PloS one 2012, 7 (9).

21. Macia, E.; Ehrlich, M.; Massol, R.; Boucrot, E.; Brunner, C.; Kirchhausen, T.; Developmental cell 2006, 10 (6), 839–850.

22. Lamaze, C.; Fujimoto, L. M.; Yin, H. L.; Schmid, S. L.; Journal of Biological Chemistry 1997, 272 (33), 20332–20335.

23. Wang, L.-H.; Rothberg, K. G.; Anderson, R.; The Journal of cell biology 1993, 123 (5), 1107–1117.

24. Corazzari, M.; Gagliardi, M.; Fimia, G. M.; Piacentini, M.; Frontiers in oncology 2017, 7, 78.

25. Huang, Y.; Hui, K.; Jin, M.; Yin, S.; Wang, W.; Ren, Q.; Scientific reports 2016, 6 (1), 1–11.

26. Mercer, J.; Schelhaas, M.; Helenius, A.; Annu Rev Biochem 2010, 79, 803–33. DOI 10.1146/annurev-biochem-060208-104626.

27. Quirin, K.; Eschli, B.; Scheu, I.; Poort, L.; Kartenbeck, J.; Helenius, A.; Virology 2008, 378 (1), 21–33.

28. Wellington, C. L.; Ellerby, L. M.; Gutekunst, C.-A.; Rogers, D.; Warby, S.; Graham, R. K.; Loubser, O.; van Raamsdonk, J.; Singaraja, R.; Yang, Y.-Z.; Journal of Neuroscience 2002, 22 (18), 7862–7872.

29. Mejia, R. O. S.; Friedlander, R. M.; The Neuroscientist 2001, 7 (6), 480–489.

30. Gafni, J.; Ellerby, L. M.; Journal of Neuroscience 2002, 22 (12), 4842–4849.

31. Hamid, O.; Ismail, R.; Puzanov, I.; The oncologist 2020, 25 (3), e423.

32. Swift, L.; Zhang, C.; Trippett, T.; Narendran, A. J. O.; therapy, 2019, 12, 1293.

33. Panizza, B. J.; de Souza, P.; Cooper, A.; Roohullah, A.; Karapetis, C. S.; Lickliter, J. D. J. E.; 2019, 50, 433–441.

34. Jassal, B.; Matthews, L.; Viteri, G.; Gong, C.; Lorente, P.; Fabregat, A.; Sidiropoulos, K.; Cook, J.; Gillespie, M.; Haw, R.; Nucleic acids research 2020, 48 (D1), D498–D503.

35. Abraham, M. J.; Murtola, T.; Schulz, R.; Páll, S.; Smith, J. C.; Hess, B.; Lindahl, E. J. S.; 2015, 1, 19–25.

36. Malde, A. K.; Zuo, L.; Breeze, M.; Stroet, M.; Poger, D.; Nair, P. C.; Oostenbrink, C.; Mark, A. E. J. J. o. c. t.; computation, 2011, 7 (12), 4026–4037.

