## Supplemental Data for "Sapogenin based self-assembly structures activating a non-apoptotic cell death via multiple pathways"

Supplementary Table 5. The $^{13}C$ and $^{1}H$ NMR data of AG-05 (100/400 MHz, δ ppm, in CDCl3). 31
Spectrum 22. HR-ESI-MS Spectrum of AG-05 (positive mode). 32
Spectrum 23. $^{1}H$ NMR Spectrum of AG-05. 33
Spectrum 24. $^{13}C$ NMR Spectrum of AG-05. 33
Spectrum 25. COSY spectrum of AG-05 34
Spectrum 26. HSQC spectrum of AG-05. 34
Spectrum 27. HMBC spectrum of AG-05. 35
Spectrum 28. HMBC spectrum of AG-05. 35
Supplementary Figure 7. Chemical Structure of AG-06 36
Supplementary Table 6. The $^{13}C$ and $^{1}H$ NMR data of AG-06 (100/400 MHz, δ ppm, in CDCl3). 36
Spectrum 29. HR-ESI-MS Spectrum of AG-06 (positive mode). 37
Spectrum 30. $^{1}H$ NMR Spectrum of AG-06. 38
Spectrum 31. $^{13}C$ NMR Spectrum of AG-06. 38
Spectrum 32. DEPT135 spectrum of AG-06 39
Spectrum 33. COSY spectrum of AG-06 39
Spectrum 34. HSQC spectrum of AG-06. 40
Spectrum 36. HMBC spectrum of AG-06 41
Supplementary Figure 8. Chemical Structure of AG-07 41
Spectrum 37. HR-ESI-MS Spectrum of AG-07 (positive mode). 42
Spectrum 38. $^{1}H$ NMR Spectrum of AG-07. 42
Spectrum 39. $^{13}C$ NMR Spectrum of AG-07. 43
Supplementary Figure 9. Chemical Structure of CG-02 43
Supplementary Table 7. The $^{13}C$ and $^{1}H$ NMR data of CG-02 (100/400 MHz, δ ppm, in CDCl3). 43
Spectrum 40. HR-ESI-MS Spectrum of CG-02 (positive mode). 45
Spectrum 41. $^{1}H$ NMR Spectrum of CG-02. 45
Spectrum 42. $^{13}C$ NMR Spectrum of CG-02. 46
Spectrum 43. DEPT135 spectrum of CG-02. 46
Spectrum 44. COSY spectrum of CG-02. 47
Spectrum 45. HMQC spectrum of CG-02. 47
Spectrum 46. HMBC spectrum of CG-02. 48
Supplementary Figure 10. Chemical Structure of CG-03 48
Supplementary Table 8. The $^{13}C$ and $^{1}H$ NMR data of CG-03 (100/400 MHz, δ ppm, in CDCl3). 49
Spectrum 47. HR-ESI-MS Spectrum of CG-03 (positive mode). 50

Spectrum 48. $^{1}$H NMR Spectrum of CG-03. 50
Spectrum 49. $^{13}$C NMR Spectrum of CG-03. 51
Spectrum 50. DEPT135 spectrum of CG-03. 51
Spectrum 51. COSY spectrum of CG-03. 52
Spectrum 52. HMQC spectrum of CG-03. 52
Spectrum 53. HMBC spectrum of CG-03. 53
Supplementary Figure 11. Chemical Structure of CG-04 53
Supplementary Table 9. The $^{13}$C and $^{1}$H NMR data of CG-04 (100/400 MHz, δ ppm, in CDCl3). 54
Spectrum 54. HR-ESI-MS Spectrum of CG-04 (positive mode). 55
Spectrum 55. $^{1}$H NMR Spectrum of CG-04. 55
Spectrum 56. $^{13}$C NMR Spectrum of CG-04. 56
Spectrum 57. DEPT135 spectrum of CG-04. 56
Spectrum 58. COSY spectrum of CG-04. 57
Spectrum 59. HMQC spectrum of CG-04. 57
Spectrum 60. HMBC spectrum of CG-04. 58
Supplementary Figure 12. Chemical Structure of CG-05 58
Supplementary Table 10. The $^{13}$C and $^{1}$H NMR data of CG-05 (100/400 MHz, δ ppm, in CDCl3). 58
Spectrum 61. HR-ESI-MS Spectrum of CG-05 (positive mode). 59
Spectrum 62. $^{1}$H NMR Spectrum of CG-05. 60
Spectrum 63. $^{13}$C NMR Spectrum of CG-05. 60
Spectrum 64. DEPT135 spectrum of CG-05. 61
Spectrum 65. COSY spectrum of CG-05. 61
Spectrum 66. HMQC spectrum of CG-05. 62
Spectrum 67. HMBC spectrum of CG-05. 62
Supplementary Figure 13. Chemical Structure of CG-06 63
Supplementary Table 11. The $^{13}$C and $^{1}$H NMR data of CG-06 (100/400 MHz, δ ppm, in CDCl3). 63
Spectrum 68. HR-ESI-MS Spectrum of CG-06 (positive mode). 64
Spectrum 69. $^{1}$H NMR Spectrum of CG-06. 64
Spectrum 70. $^{13}$C NMR Spectrum of CG-06. 65
Spectrum 71. DEPT135 spectrum of CG-06 65
Spectrum 72. COSY spectrum of CG-06 66
Spectrum 73. HMQC spectrum of CG-06. 66
Spectrum 74. HMBC spectrum of CG-05. 67
Supplementary Figure 14. Chemical Structure of SCG-01 67

Supplementary Table 12. The $^{13}C$ and $^{1}H$ NMR data of SCG-01 (100/400 MHz, δ ppm, in CDCl3). 67

Spectrum 75.HR-ESI-MS Spectrum of SCG-01 (positive mode). 68

Spectrum 76. $^{1}H$ NMR Spectrum of SCG-01. 69

Spectrum 77. $^{13}C$ NMR Spectrum of SCG-01. 69

Spectrum 78. DEPT135 spectrum of SCG-01. 70

Spectrum 79. COSY spectrum of SCG-01. 70

Spectrum 80. HMQC spectrum of SCG-01. 71

Spectrum 81. HMBC spectrum of SCG-01. 71

Supplementary Figure 15. Chemical Structure of SCG-02 72

Supplementary Table 13. The $^{13}C$ and $^{1}H$ NMR data of SCG-02 (100/400 MHz, δ ppm, in CDCl3). 72

Spectrum 82. HR-ESI-MS Spectrum of SCG-02 (positive mode). 73

Spectrum 83. $^{1}H$ NMR Spectrum of SCG-02. 73

Spectrum 84. $^{13}C$ NMR Spectrum of SCG-02. 74

Spectrum 85. DEPT135 spectrum of SCG-02. 74

Spectrum 86. COSY spectrum of SCG-02. 75

Spectrum 87. HSQC spectrum of SCG-02. 75

Spectrum 88. HMBC spectrum of SCG-02. 76

Supplementary Figure 16. Chemical Structure of SCG-03 76

Supplementary Table 14. The $^{13}C$ and $^{1}H$ NMR data of SCG-03 (100/400 MHz, δ ppm, in CDCl3). 76

Spectrum 89. HR-ESI-MS Spectrum of SCG-03 (positive mode). 77

Spectrum 90. $^{1}H$ NMR Spectrum of SCG-03. 78

Spectrum 91. $^{13}C$ NMR Spectrum of SCG-03. 78

Spectrum 92. DEPT135 spectrum of SCG-03. 79

Spectrum 93. COSY spectrum of SCG-03. 79

Spectrum 94. HMQC spectrum of SCG-03. 80

Spectrum 95. HMBC spectrum of SCG-03. 80

Supplementary Figure 17. Chemical Structure of SCG-04 81

Supplementary Table 15. The $^{13}C$ and $^{1}H$ NMR data of SCG-04 (100/400 MHz, δ ppm, in CDCl3). 81

Spectrum 96. HR-ESI-MS Spectrum of SCG-04 (positive mode). 82

Spectrum 97. $^{1}H$ NMR Spectrum of SCG-04. 82

Spectrum 98. $^{13}C$ NMR Spectrum of SCG-04. 83

Spectrum 99. COSY spectrum of SCG-04. 83

Spectrum 100. HMQC spectrum of SCG-04. 84

Spectrum 101. HMBC spectrum of SCG-04. 84
Spectrum 102. HMBC spectrum of SCG-04. 85
Supplementary Figure 18. Chemical Structure of SCG-05 85
Supplementary Table 16. The $^{13}C$ and $^{1}H$ NMR data of SCG-05 (100/500 MHz, δ ppm, in CDCl3). 86
Spectrum 103. HR-ESI-MS Spectrum of SCG-05 (positive mode). 86
Spectrum 104. $^{1}H$ NMR Spectrum of SCG-05. 87
Spectrum 105. $^{13}C$ NMR Spectrum of SCG-05. 87
Spectrum 106. DEPT135 spectrum of SCG-05. 88
Spectrum 107. COSY spectrum of SCG-05. 88
Spectrum 108. HMQC spectrum of SCG-05. 89
Spectrum 109. HMBC spectrum of SCG-05. 89
Supplementary Figure 19. Chemical Structure of SCG-06 90
Supplementary Table 17. The $^{13}C$ and $^{1}H$ NMR data of SCG-06 (100/500 MHz, δ ppm, in CDCl3). 90
Spectrum 110. HR-ESI-MS Spectrum of SCG-06 (positive mode). 91
Spectrum 111. $^{1}H$ NMR Spectrum of SCG-06. 91
Spectrum 112. $^{13}C$ NMR Spectrum of SCG-06. 92
Spectrum 113. DEPT135 spectrum of SCG-06. 92
Spectrum 114. COSY spectrum of SCG-06. 93
Spectrum 115. HMQC spectrum of SCG-06. 93
Spectrum 116. HMBC spectrum of SCG-06. 94
Supplementary Figure 20. Chemical Structure of SCG-07 94
Supplementary Table 18. The $^{13}C$ and $^{1}H$ NMR data of SCG-07 (100/500 MHz, δ ppm, in CDCl3). 95
Spectrum 117. HR-ESI-MS Spectrum of SCG-07 (positive mode). 95
Spectrum 118. $^{1}H$ NMR Spectrum of SCG-07. 96
Spectrum 119. $^{13}C$ NMR Spectrum of SCG-07. 96
Spectrum 120. DEPT135 spectrum of SCG-07. 97
Spectrum 121. COSY spectrum of SCG-07. 97
Spectrum 122. HMQC spectrum of SCG-07. 98
Spectrum 123. HMBC spectrum of SCG-07. 98

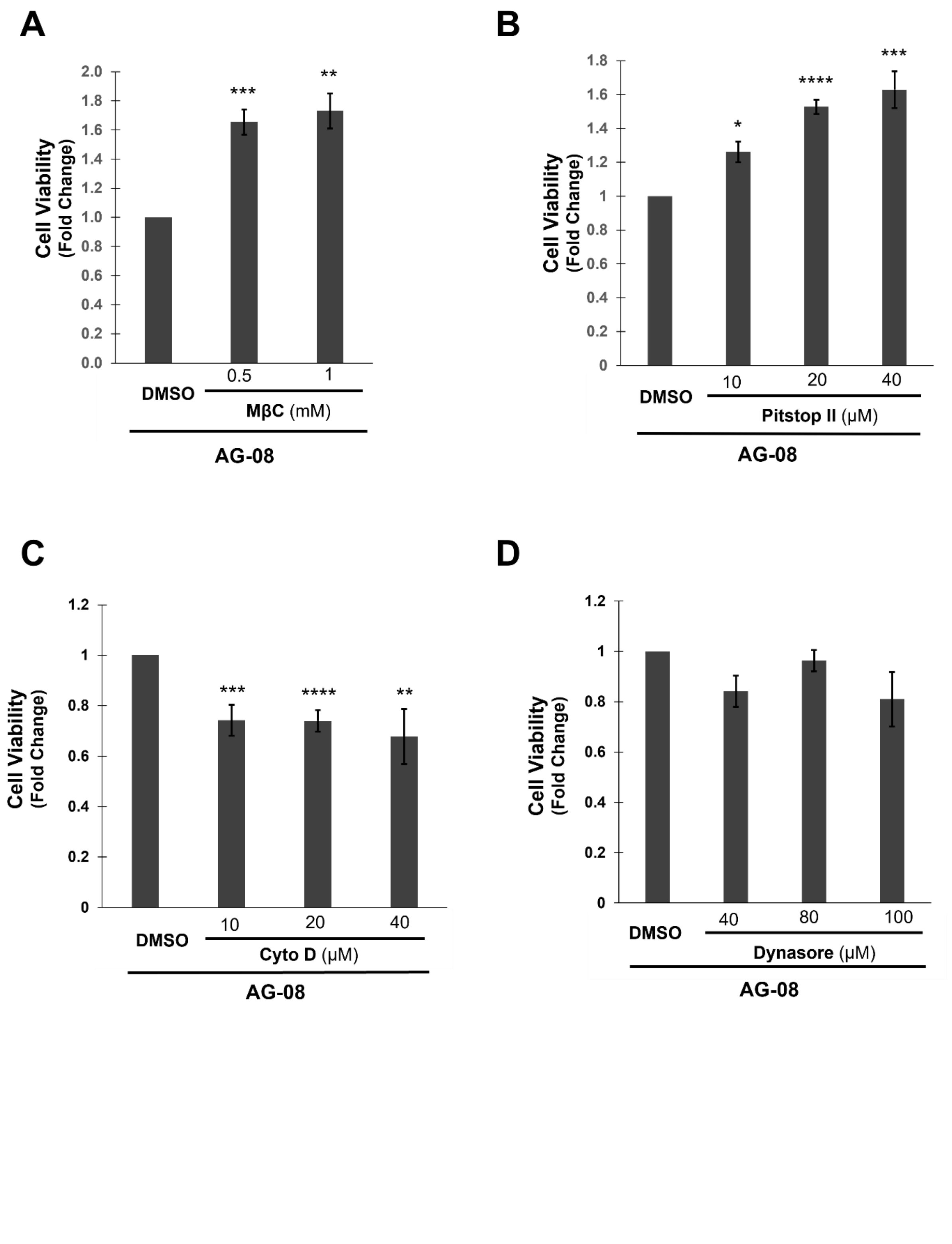
A
Cell Viability
(Fold Change)
2.0
1.8
1.6
1.4
1.2
1.0
0.8
0.6
0.4
0.2
0.0
***
**
0.5
1
DMSO
MβC (mM)
AG-08
B
Cell Viability
(Fold Change)
1.8
1.6
1.4
1.2
1
0.8
0.6
0.4
0.2
0
*
****
***
10
20
40
DMSO
Pitstop II (μM)
AG-08
C
Cell Viability
(Fold Change)
1.2
1
0.8
0.6
0.4
0.2
0
***
****
**
10
20
40
DMSO
Cyto D (μM)
AG-08
D
Cell Viability
(Fold Change)
1.2
1
0.8
0.6
0.4
0.2
0
40
80
100
DMSO
Dynasore (μM)
AG-08

**E**

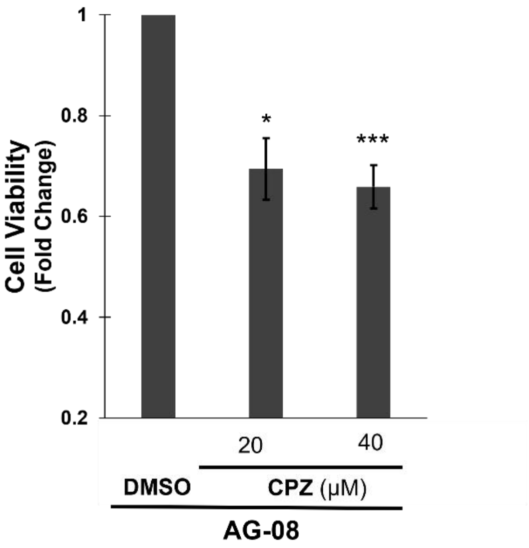

**F**

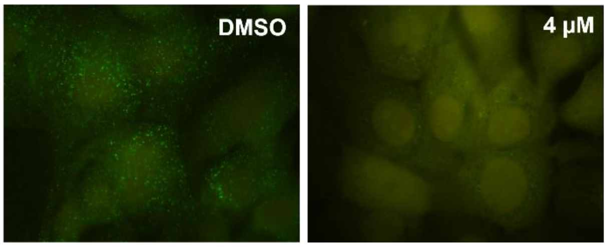

**Supplementary Figure 1. AG-08 particles affect endosomal pathway** (A-E) HCC1937 cells were pre-treated with different concentration of pitstop II, MβC, dynasore, Cyto D or CPZ for 1 h and 8 μM AG-08 for 24 h. Reported values was normalized on cells treated with only AG-08. Error bars are the standard deviations (n=3). p-values were calculated with respect to AG-08 treated cells by two-tailed equal variance Student's t-test (*$p<0.05$, **$p<0.005$, ***$p<0.001$, ****$p<0.0001$). (F) Following 4 μM AG-08 treatment for 40 h, EEA1 proteins of HCC1937 cells were stained using anti-EEA1 antibody

**Supplementary Table 1. Gene expression level, p value and FDR p value of AG-08 and control.**

| Gene Symbol | AG-08 (Log2) | Control (Log2) | P-val | FDR P-val |
|---|---|---|---|---|
| IL1RL1 | 10.06 | 6.82 | 0.013 | 0.8393 |
| CXCL8 | 12.86 | 9.63 | 8.63E-06 | 0.2667 |
| CCL20 | 15.79 | 12.58 | 6.31E-07 | 0.0857 |
| DDIT3 | 8.56 | 5.64 | 2.22E-05 | 0.335 |
| TNFAIP3 | 15.77 | 12.92 | 0.0009 | 0.8393 |
| CREB5 | 11.14 | 8.35 | 0.0181 | 0.8393 |
| AKAP12 | 9.04 | 6.27 | 0.0122 | 0.8393 |
| CHAC1 | 9.98 | 7.22 | 0.0002 | 0.8393 |
| IL1A | 12.9 | 10.2 | 1.96E-06 | 0.1329 |
| CXCL3 | 9.67 | 6.99 | 0.0152 | 0.8393 |
| IL6 | 11.97 | 9.32 | 0.0021 | 0.8393 |
| choygey | 11.85 | 9.26 | 0.0009 | 0.8393 |
| TNFAIP6 | 10.32 | 7.73 | 2.91E-05 | 0.3589 |
| cheeju | 4.53 | 7.11 | 0.0003 | 0.8393 |
| AC068797.1 | 4.17 | 6.72 | 0.0038 | 0.8393 |
| plergler | 5.85 | 8.38 | 0.0083 | 0.8393 |
| HSPA7 | 7.45 | 4.94 | 2.82E-05 | 0.3589 |
| snawperbu | 8.51 | 10.99 | 0.005 | 0.8393 |
| CXCL2 | 10.58 | 8.19 | 0.0125 | 0.8393 |
| FGF5 | 6.46 | 4.09 | 0.0036 | 0.8393 |
| GADD45A | 13.43 | 11.09 | 0.0001 | 0.8393 |
| DAW1 | 7.07 | 4.73 | 0.0045 | 0.8393 |
| dymabu | 4.94 | 7.28 | 0.0002 | 0.8393 |
| sworkaby | 4.69 | 2.37 | 0.0123 | 0.8393 |
| swawchu | 10.51 | 12.82 | 0.0272 | 0.8393 |
| FRMD6 | 9.43 | 7.12 | 0.0042 | 0.8393 |
| sparsloy | 7.2 | 9.51 | 0.0006 | 0.8393 |

| | | | | |
|---|---|---|---|---|
| HSP90B1 | 10.67 | 8.38 | 0.0007 | 0.8393 |
| KRT5 | 9.49 | 11.75 | 0.0017 | 0.8393 |
| flyspabu | 4.45 | 6.68 | 0.0001 | 0.8393 |
| HSPA1A | 15.75 | 13.53 | 9.73E-05 | 0.8393 |
| NR4A1 | 8.84 | 6.63 | 0.0295 | 0.8393 |
| MIR6515 | 6.96 | 4.75 | 0.0009 | 0.8393 |
| PTGS2 | 11.9 | 9.7 | 0.0005 | 0.8393 |
| DNAJB9 | 10.33 | 8.13 | 5.34E-05 | 0.558 |
| kluky | 4.18 | 6.37 | 0.0058 | 0.8393 |
| SPRR2A | 8.31 | 6.12 | 0.0024 | 0.8393 |
| steybleebu | 7.5 | 9.69 | 0.0032 | 0.8393 |
| slukeebu | 5.7 | 7.87 | 0.0385 | 0.8393 |
| chostarby | 6.86 | 9.01 | 0.0058 | 0.8393 |
| borflaw | 6.24 | 4.1 | 0.0471 | 0.8393 |
| vyglabu | 4.99 | 2.86 | 0.0099 | 0.8393 |
| SPRY4 | 6.6 | 4.5 | 0.0146 | 0.8393 |
| RNA18S5 | 12.08 | 10 | 0.0034 | 0.8393 |
| sportobo | 9.07 | 11.14 | 0.0166 | 0.8393 |
| MAP1B | 9.32 | 7.26 | 0.004 | 0.8393 |
| AC079949.1 | 7.8 | 5.75 | 0.0038 | 0.8393 |
| floynabo | 5.07 | 3.01 | 0.0245 | 0.8393 |
| jawrubu | 4.94 | 6.99 | 0.0289 | 0.8393 |
| runara | 4.96 | 7 | 0.0265 | 0.8393 |
| RP11-253I19.3 | 3.7 | 5.74 | 0.0188 | 0.8393 |
| RCAN1 | 13.43 | 11.4 | 0.0013 | 0.8393 |
| HERPUD1 | 15.89 | 13.87 | 0.0002 | 0.8393 |
| AL161626.1 | 4.86 | 2.85 | 0.0021 | 0.8393 |
| rersharbu | 6.82 | 8.82 | 0.0083 | 0.8393 |
| GEM | 6.45 | 4.45 | 0.0079 | 0.8393 |
| RP11-74J13.9 | 3.93 | 5.92 | 0.0026 | 0.8393 |

| | | | | |
|---|---|---|---|---|
| beyly | 3.88 | 5.86 | 0.0105 | 0.8393 |
| RP5-1180E21.5 | 7.94 | 9.91 | 0.0066 | 0.8393 |
| MIR1284 | 3.95 | 5.93 | 1.51E-05 | 0.335 |
| SERPINE1 | 9.03 | 7.07 | 0.0079 | 0.8393 |
| ferchor | 4.51 | 6.47 | 0.0031 | 0.8393 |
| sneykleebu | 5.94 | 7.89 | 0.02 | 0.8393 |
| ETS1 | 13.17 | 11.21 | 0.0054 | 0.8393 |
| RNU11-6P | 3.79 | 5.74 | 0.0073 | 0.8393 |
| bonawbu | 4.47 | 6.42 | 0.0096 | 0.8393 |
| AC006548.19 | 4.15 | 6.1 | 0.0083 | 0.8393 |
| keyfybu | 4.42 | 6.35 | 0.0053 | 0.8393 |
| ERO1B | 8.83 | 6.9 | 0.0032 | 0.8393 |
| hunimo | 4.9 | 2.97 | 0.0261 | 0.8393 |
| flyleyby | 3.87 | 5.79 | 0.0017 | 0.8393 |
| CSF3 | 5.21 | 3.29 | 0.0068 | 0.8393 |
| Y_RNA | 4.88 | 2.96 | 0.0148 | 0.8393 |
| RP11-632K20.8 | 6.86 | 8.77 | 0.001 | 0.8393 |
| nerure | 6.39 | 8.3 | 0.0013 | 0.8393 |
| temire | 3.91 | 5.81 | 0.0215 | 0.8393 |
| RP5-890E16.5 | 3.64 | 5.54 | 0.0346 | 0.8393 |
| METTL7A | 12.34 | 14.23 | 0.0073 | 0.8393 |
| mawry | 4.91 | 6.8 | 0.0007 | 0.8393 |
| DUSP1 | 12.84 | 10.95 | 0.0014 | 0.8393 |
| chergy | 6.6 | 8.48 | 0.0057 | 0.8393 |
| spaforby | 8.45 | 10.31 | 0.0004 | 0.8393 |
| skoytoyby | 5.73 | 3.86 | 0.0028 | 0.8393 |
| leeklerbu | 2.34 | 4.21 | 0.0158 | 0.8393 |
| spernawbo | 5.32 | 7.18 | 0.0061 | 0.8393 |
| skaskoyby | 6.69 | 8.54 | 0.0084 | 0.8393 |
| mazybu | 3.22 | 5.07 | 0.0143 | 0.8393 |

| | | | | |
|---|---|---|---|---|
| swyjee | 6.02 | 7.88 | 0.0353 | 0.8393 |
| ATF3 | 7.04 | 5.19 | 0.0003 | 0.8393 |
| teesnarby | 4.07 | 5.92 | 0.0064 | 0.8393 |
| SIRPG-AS1 | 4.97 | 6.81 | 0.0181 | 0.8393 |
| myklar | 4.37 | 6.22 | 0.0223 | 0.8393 |
| TNFRSF9 | 8.07 | 6.22 | 2.00E-05 | 0.335 |
| slorfley | 3.4 | 5.24 | 0.0038 | 0.8393 |
| JUN | 12.32 | 10.47 | 0.0028 | 0.8393 |
| glupuby | 6.47 | 8.3 | 0.0118 | 0.8393 |
| muneme | 3.26 | 5.09 | 0.0154 | 0.8393 |
| LRRC49 | 8.59 | 6.77 | 4.06E-05 | 0.4595 |
| CSRNP1 | 7.09 | 5.26 | 0.0052 | 0.8393 |
| guzobu | 3.12 | 4.94 | 0.0124 | 0.8393 |
| skeymeybo | 5.88 | 4.08 | 0.0442 | 0.8393 |
| RN7SKP36 | 6.69 | 4.9 | 0.0009 | 0.8393 |
| vybu | 4.27 | 2.47 | 0.0043 | 0.8393 |
| ACSM3 | 4.97 | 6.76 | 0.0089 | 0.8393 |
| klobly | 6.7 | 8.49 | 0.0015 | 0.8393 |
| doymabu | 4.76 | 6.54 | 0.0007 | 0.8393 |
| LOC100128914 | 4.93 | 3.15 | 0.044 | 0.8393 |
| IFI44 | 12.67 | 14.45 | 0.0489 | 0.8393 |
| cherstarby | 6.42 | 8.19 | 0.0067 | 0.8393 |
| hosimu | 5.79 | 4.01 | 0.0018 | 0.8393 |
| kosey | 5.44 | 3.66 | 0.0124 | 0.8393 |
| riyare | 3.31 | 5.08 | 0.0374 | 0.8393 |
| skopoybo | 4.26 | 6.02 | 0.0001 | 0.8393 |
| FP671120.3 | 8.3 | 10.07 | 0.0051 | 0.8393 |
| FP236383.2 | 8.3 | 10.07 | 0.0051 | 0.8393 |
| AL353644.7 | 8.3 | 10.07 | 0.0051 | 0.8393 |
| AL592188.5 | 8.3 | 10.07 | 0.0051 | 0.8393 |

| | | | | |
|---|---|---|---|---|
| luwarbu | 3.42 | 5.19 | 0.007 | 0.8393 |
| goylobo | 6.56 | 8.33 | 0.0304 | 0.8393 |
| wardaw | 3.08 | 4.84 | 0.0067 | 0.8393 |
| starblaw | 4.63 | 6.39 | 0.0077 | 0.8393 |
| blerverby | 4.88 | 6.64 | 0.0187 | 0.8393 |
| RP4-802A10.1 | 8.32 | 10.07 | 0.019 | 0.8393 |
| gamabo | 4.18 | 5.94 | 0.0266 | 0.8393 |
| nureebo | 8.33 | 10.08 | 0.0112 | 0.8393 |
| blaployby | 2.32 | 4.06 | 0.0022 | 0.8393 |
| sworoy | 3.06 | 4.81 | 0.0069 | 0.8393 |
| sharjer | 4.32 | 6.06 | 0.0003 | 0.8393 |
| STC2 | 12.61 | 10.87 | 0.0002 | 0.8393 |
| tunemo | 7.64 | 5.91 | 0.0102 | 0.8393 |
| neyzubu | 4.44 | 6.17 | 0.0008 | 0.8393 |
| kloloy | 3.91 | 2.19 | 0.0138 | 0.8393 |
| CLCA2 | 5.92 | 7.65 | 0.0007 | 0.8393 |
| barjoybu | 4.66 | 6.38 | 0.0142 | 0.8393 |
| stuloybu | 7.4 | 9.12 | 0.0029 | 0.8393 |
| rekare | 4.03 | 5.75 | 0.0368 | 0.8393 |
| AL158839.1 | 5.7 | 3.98 | 0.0059 | 0.8393 |
| blonawbo | 5.21 | 6.92 | 0.0269 | 0.8393 |
| harero | 5.11 | 6.81 | 0.0002 | 0.8393 |
| AL137800.1 | 4.33 | 6.03 | 0.0048 | 0.8393 |
| LRRC8C | 11.05 | 9.35 | 0.0075 | 0.8393 |
| GLIPR1 | 12.79 | 11.09 | 0.0281 | 0.8393 |
| wosnawby | 7.09 | 8.78 | 0.0147 | 0.8393 |
| snawspuby | 7.99 | 9.68 | 0.0347 | 0.8393 |
| SNORD2 | 5.29 | 3.6 | 0.003 | 0.8393 |
| plarbey | 5.68 | 3.98 | 0.0165 | 0.8393 |
| klawlaw | 6.35 | 8.03 | 0.0089 | 0.8393 |

| | | | | |
|---|---|---|---|---|
| shagly | 8.29 | 9.98 | 0.0091 | 0.8393 |
| rusame | 4.71 | 6.4 | 0.0002 | 0.8393 |
| bleymawby | 5.86 | 7.54 | 0.0016 | 0.8393 |
| AC007272.3 | 4.36 | 6.04 | 0.0077 | 0.8393 |
| RABEPK | 8.03 | 6.35 | 0.0325 | 0.8393 |
| skoytyby | 6.43 | 8.1 | 0.0002 | 0.8393 |
| cherfoy | 10.56 | 12.23 | 0.0052 | 0.8393 |
| BAG3 | 15.02 | 13.35 | 0.0294 | 0.8393 |
| AC010139.1 | 12.83 | 11.17 | 0.0222 | 0.8393 |
| UAP1 | 15.84 | 14.17 | 0.0349 | 0.8393 |
| ALDH3A1 | 10.37 | 12.03 | 0.0004 | 0.8393 |
| vamo | 6.72 | 8.38 | 0.0034 | 0.8393 |
| ferraw | 6.36 | 8.02 | 0.0316 | 0.8393 |
| klorpla | 9.12 | 10.78 | 0.044 | 0.8393 |
| HSPA1B | 14 | 12.34 | 0.0008 | 0.8393 |
| terbabu | 5.12 | 3.46 | 0.0044 | 0.8393 |
| RPL17P28 | 6.52 | 8.17 | 0.0065 | 0.8393 |
| HYOU1 | 12.17 | 10.52 | 0.0003 | 0.8393 |
| RP11-325K4.3 | 9.91 | 8.27 | 0.0028 | 0.8393 |
| geeraw | 7.51 | 9.14 | 0.0041 | 0.8393 |
| jernobu | 5 | 6.63 | 0.0067 | 0.8393 |
| RP11-187C18.2 | 6.09 | 7.72 | 0.0393 | 0.8393 |
| AC110813.1 | 9.52 | 11.14 | 0.0105 | 0.8393 |
| sharflu | 4.05 | 5.67 | 0.001 | 0.8393 |
| weymubu | 3.7 | 5.32 | 0.0084 | 0.8393 |
| steynarbu | 5.53 | 7.14 | 0.0297 | 0.8393 |
| MSANTD3-TMEFF1 | 9.84 | 8.22 | 0.0398 | 0.8393 |
| chershee | 7.19 | 8.8 | 0.0058 | 0.8393 |
| LOC646762 | 9.37 | 10.98 | 0.0136 | 0.8393 |
| snarnerbu | 4.83 | 6.43 | 0.0471 | 0.8393 |

| | | | | |
|---|---|---|---|---|
| SGPP2 | 7.81 | 6.2 | 0.0152 | 0.8393 |
| GPX2 | 6.23 | 7.83 | 0.0012 | 0.8393 |
| tosuru | 5.42 | 7.02 | 0.004 | 0.8393 |
| tawsweebu | 7.72 | 9.32 | 0.0105 | 0.8393 |
| KLRC3 | 3.09 | 4.68 | 0.0062 | 0.8393 |
| PLD1 | 6.82 | 8.42 | 0.0108 | 0.8393 |
| plerder | 3.65 | 5.24 | 0.0114 | 0.8393 |
| sterda | 5.92 | 7.51 | 0.0045 | 0.8393 |
| varger | 6.16 | 7.75 | 0.0056 | 0.8393 |
| nygleeby | 3.73 | 5.31 | 0.0215 | 0.8393 |
| AC008391.1 | 13.25 | 11.66 | 0.0175 | 0.8393 |
| SMUG1 | 6.89 | 8.48 | 0.0037 | 0.8393 |
| HSPD1P11 | 5.88 | 7.47 | 0.006 | 0.8393 |
| RP4-620F22.2 | 4.44 | 2.85 | 0.0459 | 0.8393 |
| HSPA6 | 7.79 | 3.65 | 3.84E-06 | 0.1738 |
| garskeyby | 9.6 | 16.43 | 0.0483 | 0.8393 |

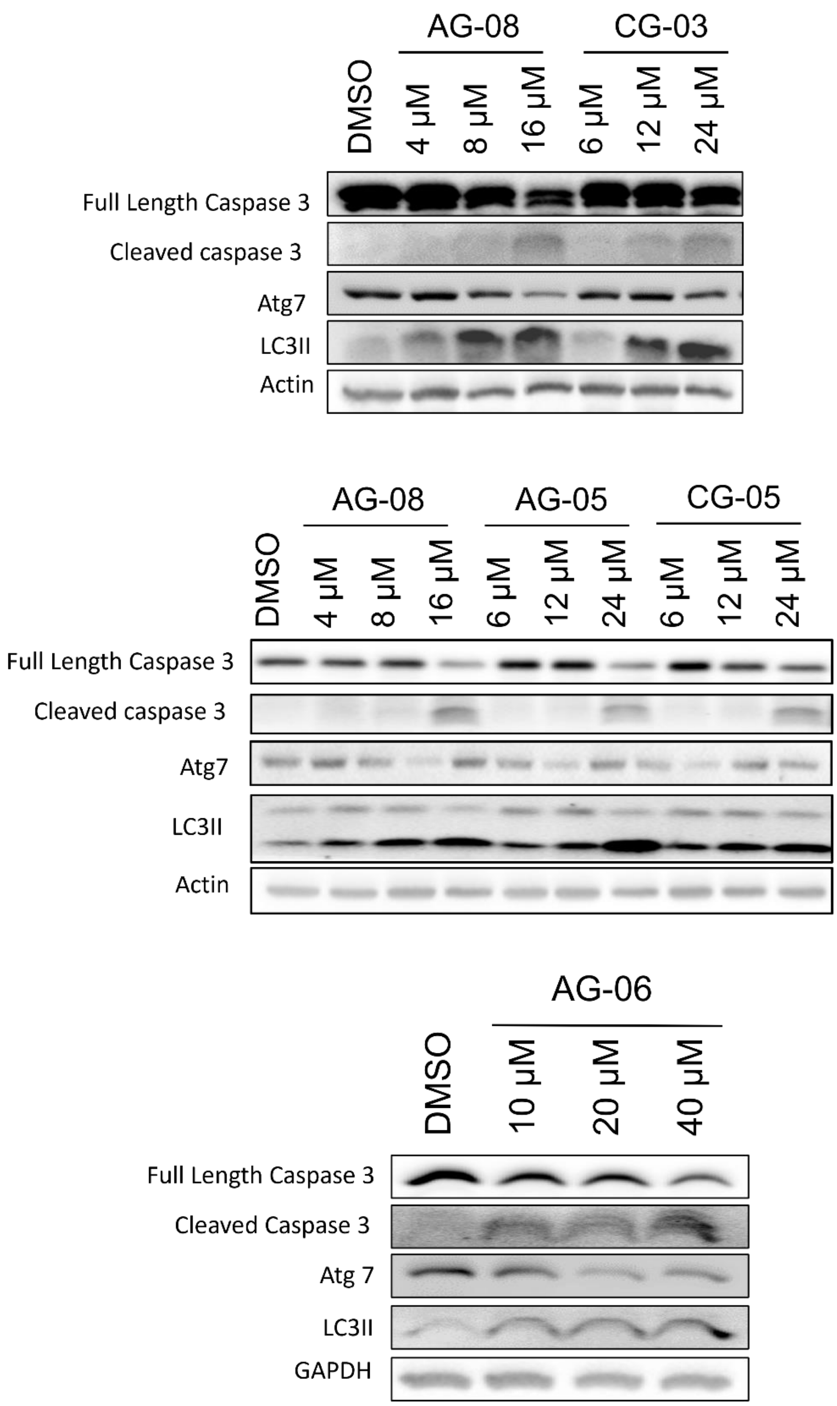

**Supplementary Figure 2. Cytotoxic compounds similarly affect LC3-II, caspase 3 and Atg7 proteins.** HCC1937 cells with cytotoxic compounds or vehicle (DMSO). The levels of LC3II, Atg-7, caspase 3 and cleaved caspase 3 were detected by immunoblotting using antibodies against them. β-Actin and GAPDH were used as the loading control.

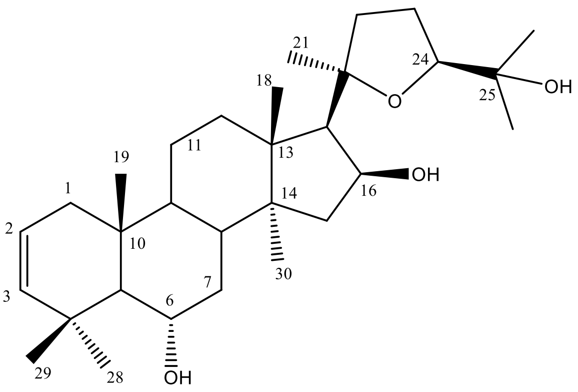

Supplementary Figure 3. Chemical Structure of AG-02

Supplementary Table 2. The $^{13}C$ and $^{1}H$ NMR data of AG-02 (100/400 MHz, δ ppm, in $CDCl_3$)

| **H/C** | **$\delta_C$ (ppm)** | **$\delta_H$ (ppm), *J* (Hz)** |
|---|---|---|
| **1** | 37.9 t | 2.04 m, 2.11 m |
| **2** | 120.3 d | 5.5 ddd (10.1, 5.4, 3) |
| **3** | 139.6 d | 5.31 m |
| **4** | 36 s | - |
| **5** | 55.4 d | 1.28 m |
| **6** | 70.2 d | 4.06 ddd (10.8, 10.8, 3.9) |
| **7** | 38.6 t | 1.44 m, 1.86 m |
| **8** | 40.9 d | 2.4 m |
| **9** | 145.8 s | - |
| **10** | 40.3 s | - |
| **11** | 116.3 d | 5.3 m |
| **12** | 37.8 t | 1.89 m, 2.14 m |
| **13** | 44.3 s | - |
| **14** | 43.9 s | - |
| **15** | 45.1 t | 1.51 dd (12.8, 6.2), 2.05 m |
| **16** | 73.4 d | 4.72 ddd (7.9, 7.9, 6.3) |
| **17** | 56.2 d | 2.36 d (7.8) |
| **18** | 18.1 q | 0.94 s |
| **19** | 23.75 q | 1.05 s |
| **20** | 87.2 s | - |

| | | |
|---|---|---|
| **21** | 28 q | 1.23 s |
| **22** | 34.6 t | 1.6 m, 2.59 q (10.4) |
| **23** | 25.9 t | 2 m |
| **24** | 81.51 d | 3.75 dd (8.1, 6.2) |
| **25** | 72 s | - |
| **26** | 26.7 q | 1.14 s |
| **27** | 27.81 q | 1.3 s |
| **28** | 34.9 | 1.2 s |
| **29** | 23.0 q | 1.17 s |
| **30** | 19.2 q | 0.79 s |

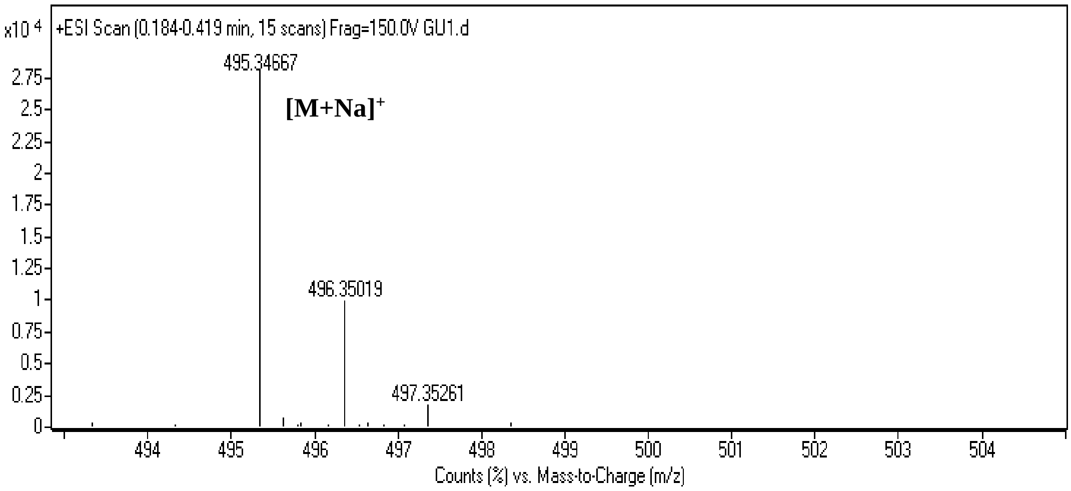

Spectrum 1. HR-ESI-MS Spectrum of AG-02 (positive mode)

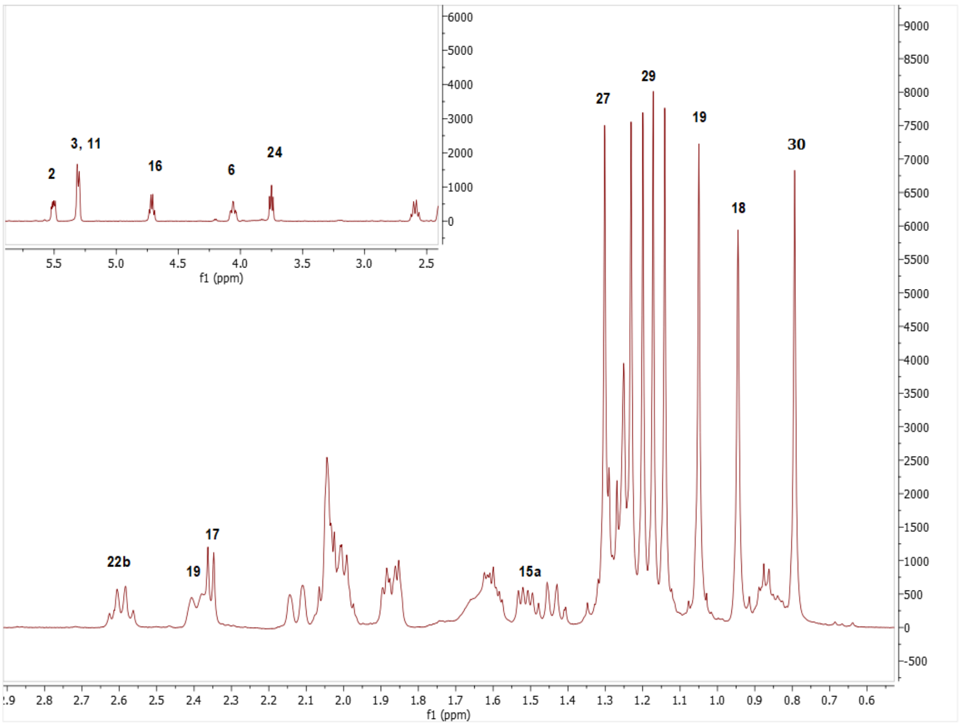

Spectrum 2. $^{1}H$ NMR Spectrum of AG-02.

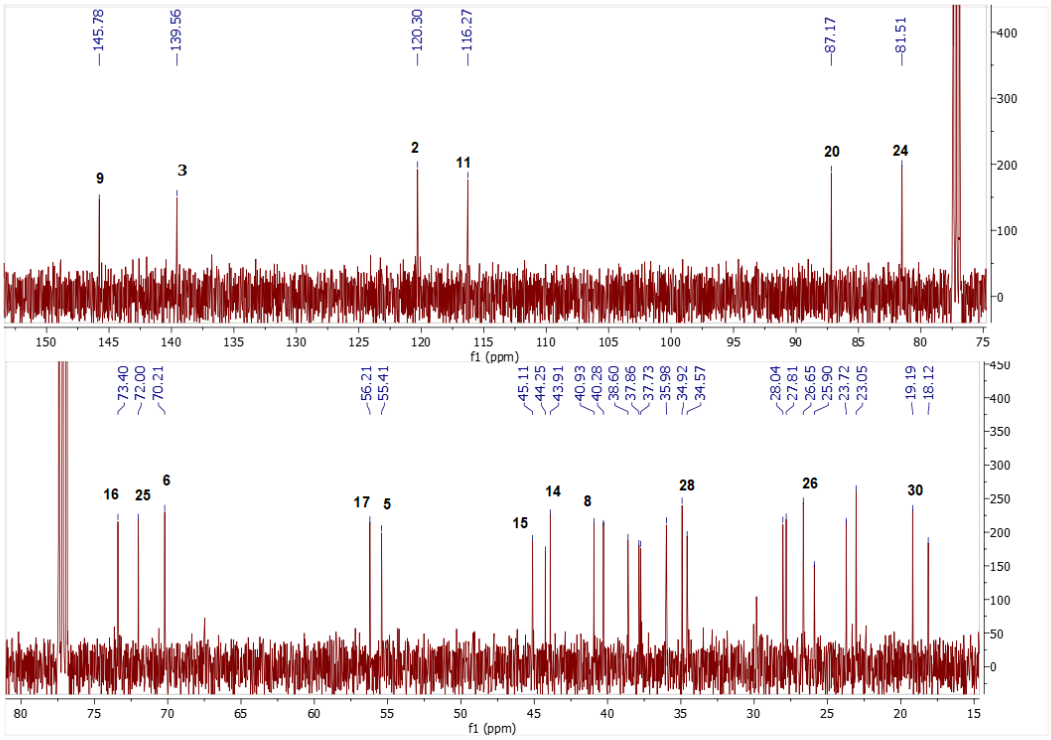

Spectrum 3. $^{13}C$ NMR Spectrum of AG-02.

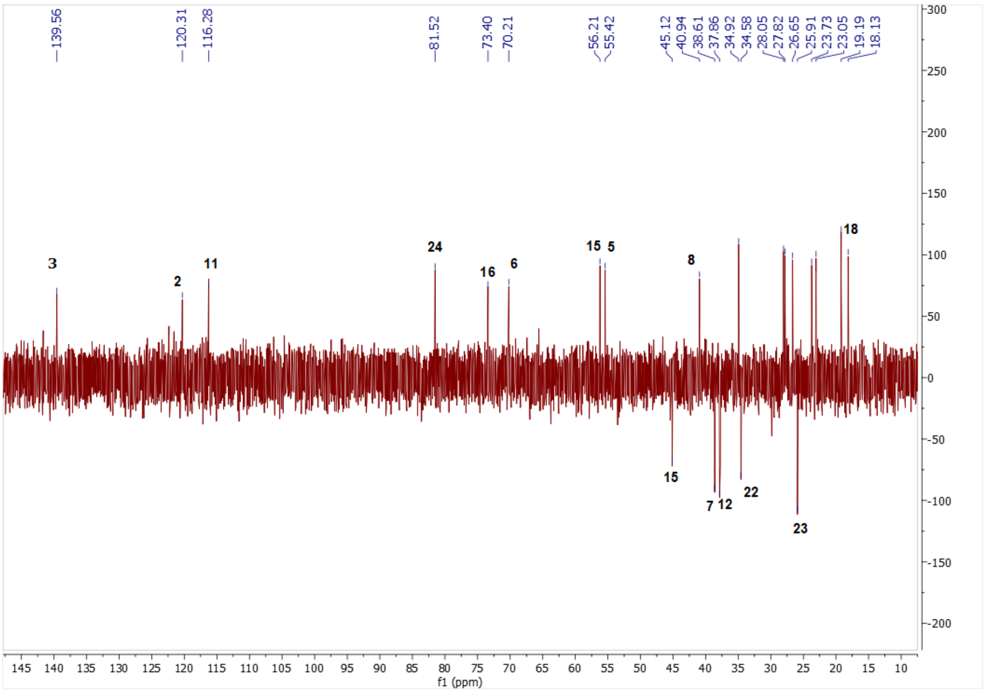

Spectrum 4. DEPT135 spectrum of AG-02

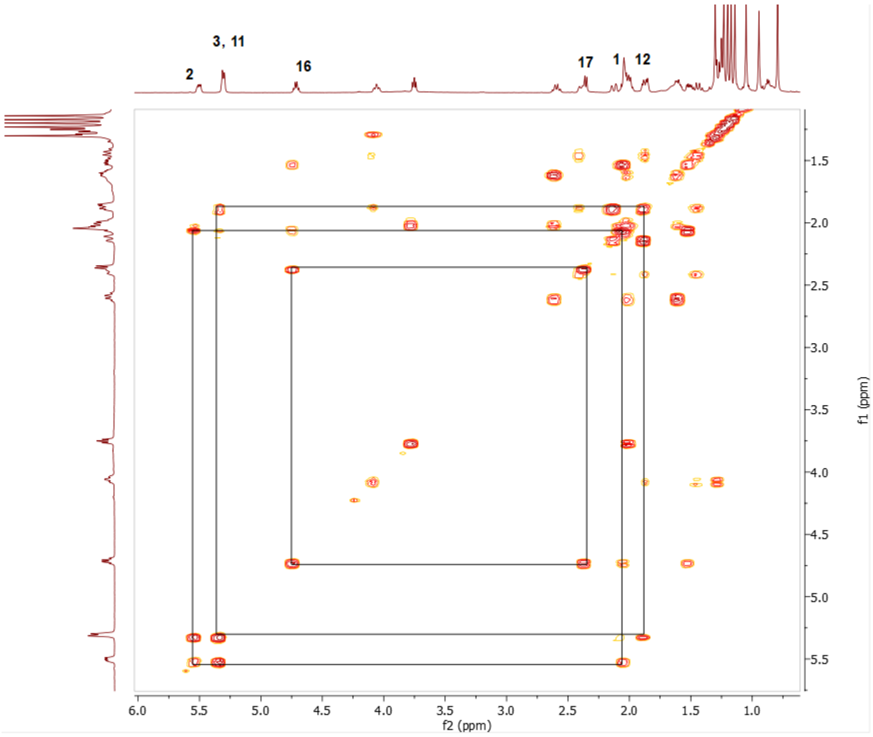

Spectrum 5. COSY spectrum of AG-02

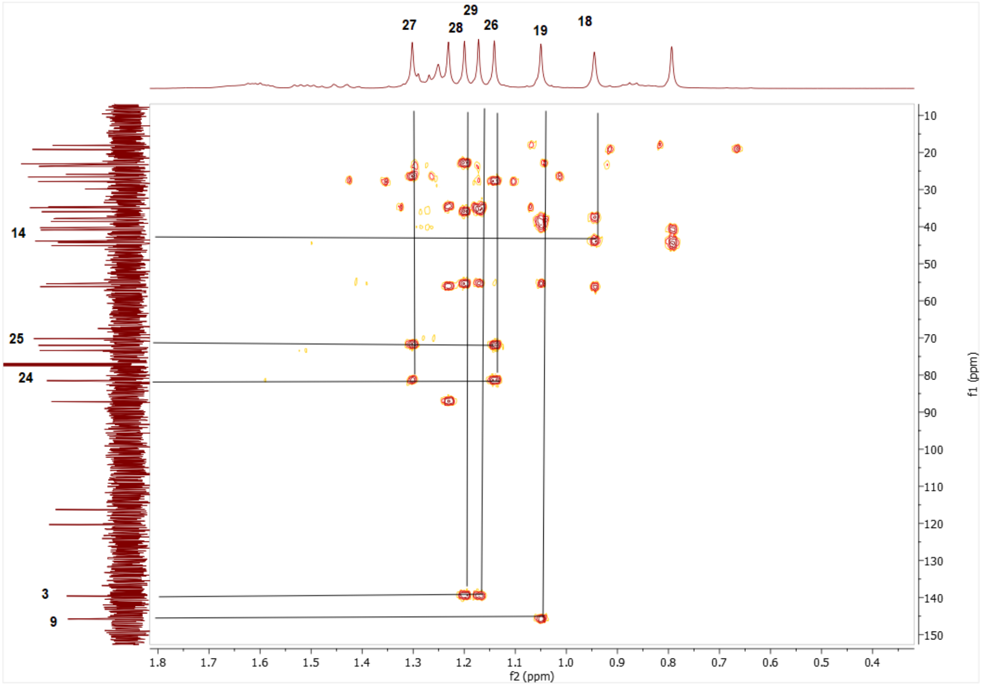

Spectrum 6. HMQC spectrum of AG-02.

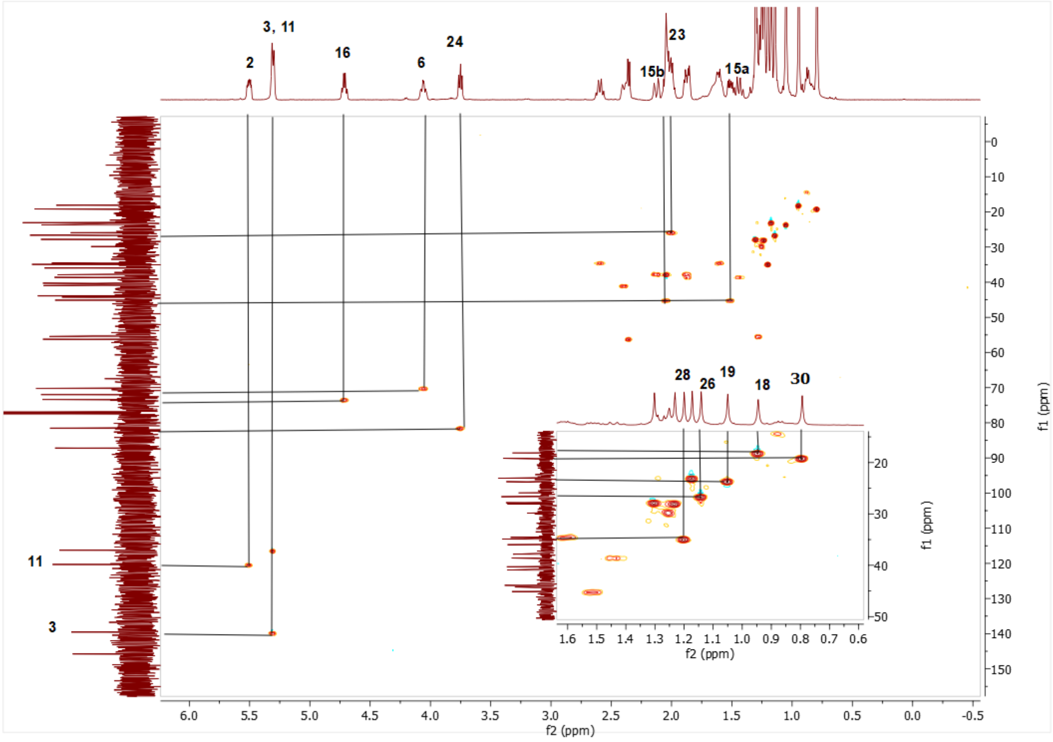

Spectrum 7. HMBC spectrum of AG-02.

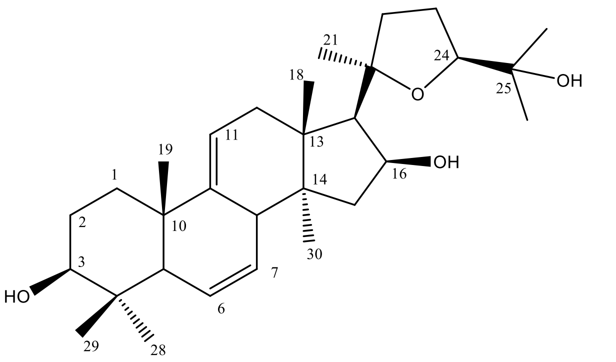

Supplementary Figure 4. Chemical Structure of AG-03

Supplementary Table 3. The $^{13}C$ and $^{1}H$ NMR data of AG-03 (100/400 MHz, δ ppm, in $CDCl_3$).

| **H/C** | **$\delta_C$ (ppm)** | **$\delta_H$ (ppm), *J* (Hz)** |
|---|---|---|
| **1** | 34.4 t | 1.59 m |
| **2** | 28.2 t | 1.78 m |
| **3** | 79.3 d | 3.23 dd (11.8, 4.8) |
| **4** | 38.8 s | - |
| **5** | 52.2 d | 1.70 d (12.2) |
| **6** | 127.3 d | 5.71 m |
| **7** | 128.9 d | 5.57 dt (10.2, 3.2) |
| **8** | 43.9 d | 2.84 brs |
| **9** | 145.9 s | - |
| **10** | 39.2 s | - |
| **11** | 113.6 d | 5.16 brs |
| **12** | 38.2 t | 2.04 m, 1.92 m |
| **13** | 43.8 s | - |
| **14** | 44.8 s | - |
| **15** | 44.3 t | 2.07 m, 1.57 m |
| **16** | 73.6 d | 4.69 m |
| **17** | 56.6 d | 2.24 dd (11.8, 6.1) |
| **18** | 19.1 q | 0.98 s |
| **19** | 20.4 q | 1.01 |
| **20** | 87.3 s | - |
| **21** | 28.2 d | 1.23 s |
| **22** | 34.8 d | 1.58 m, 2.56 m |
| **23** | 26.1 t | 2 m |
| **24** | 81.8 d | 3.75 t (7.2) |
| **25** | 72.2 s | - |
| **26** | 28.2 q | 1.29 s |
| **27** | 26.9 q | 1.14 s |
| **28** | 28.2 q | 1.01 s |
| **29** | 16 q | 0.83 s |
| **30** | 18.91 | 0.64 s |

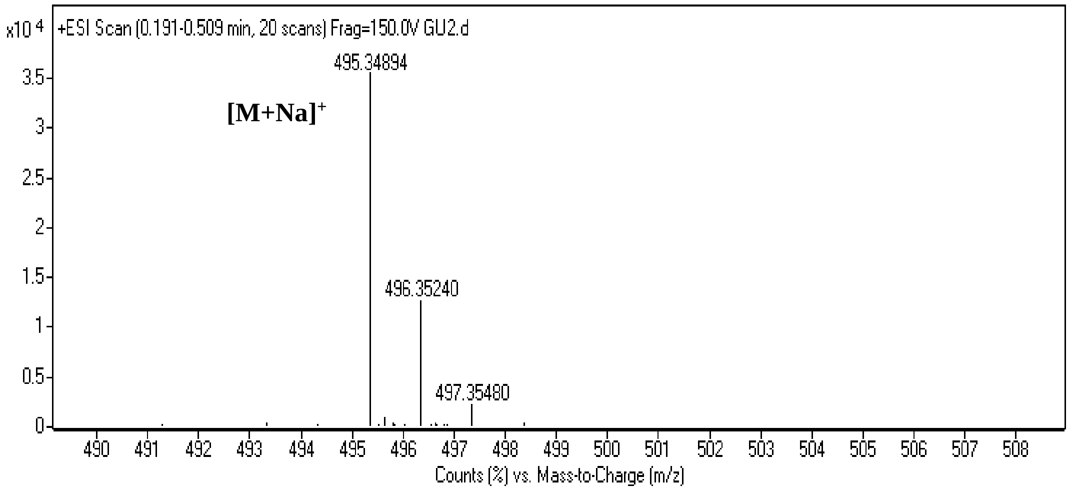

Spectrum 8. HR-ESI-MS Spectrum of AG-03 (positive mode).

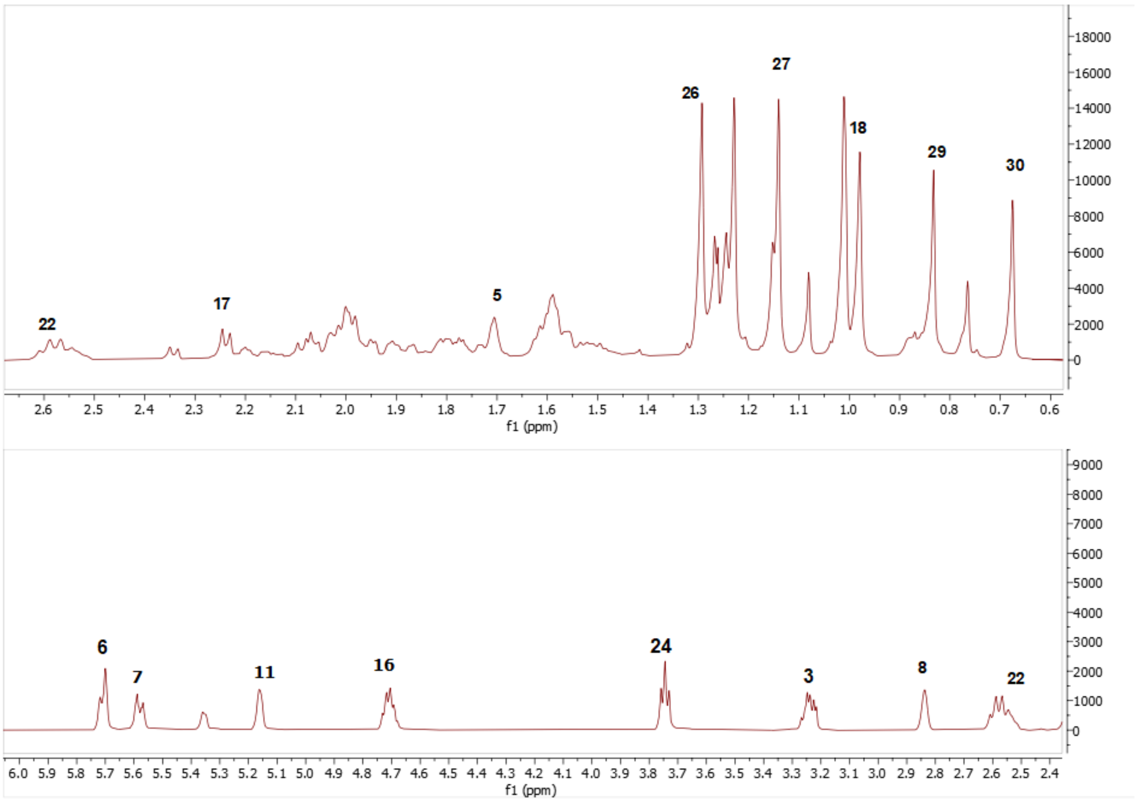

Spectrum 9. 1H NMR Spectrum of AG-03

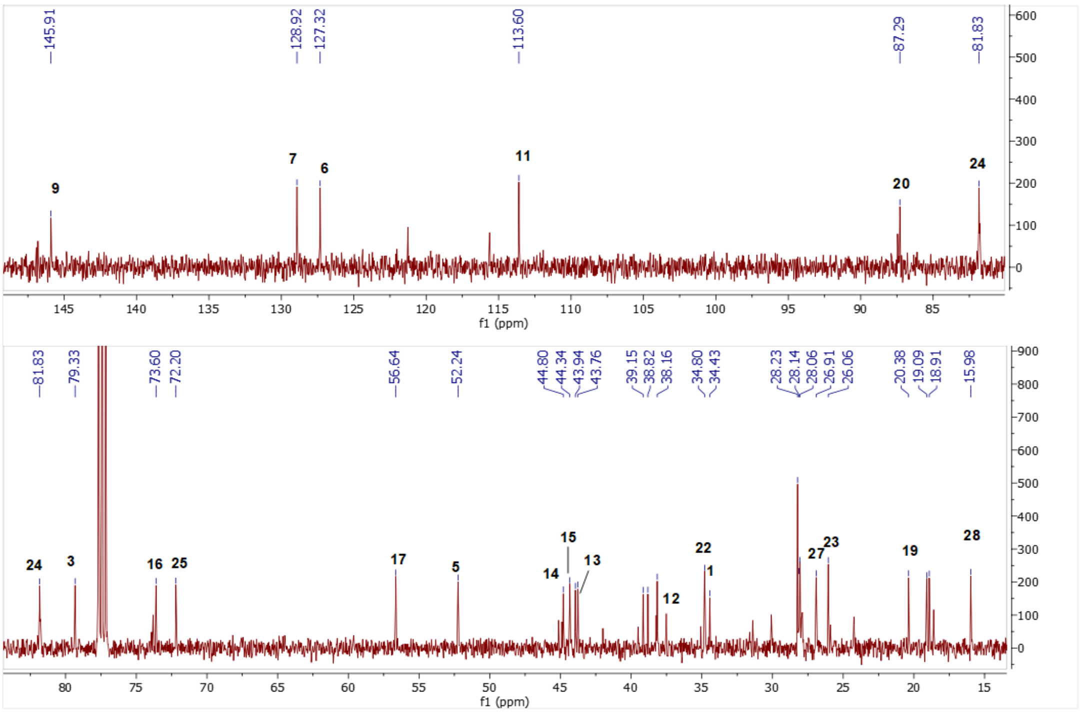

Spectrum 10. $^{13}C$ NMR Spectrum of AG-03.

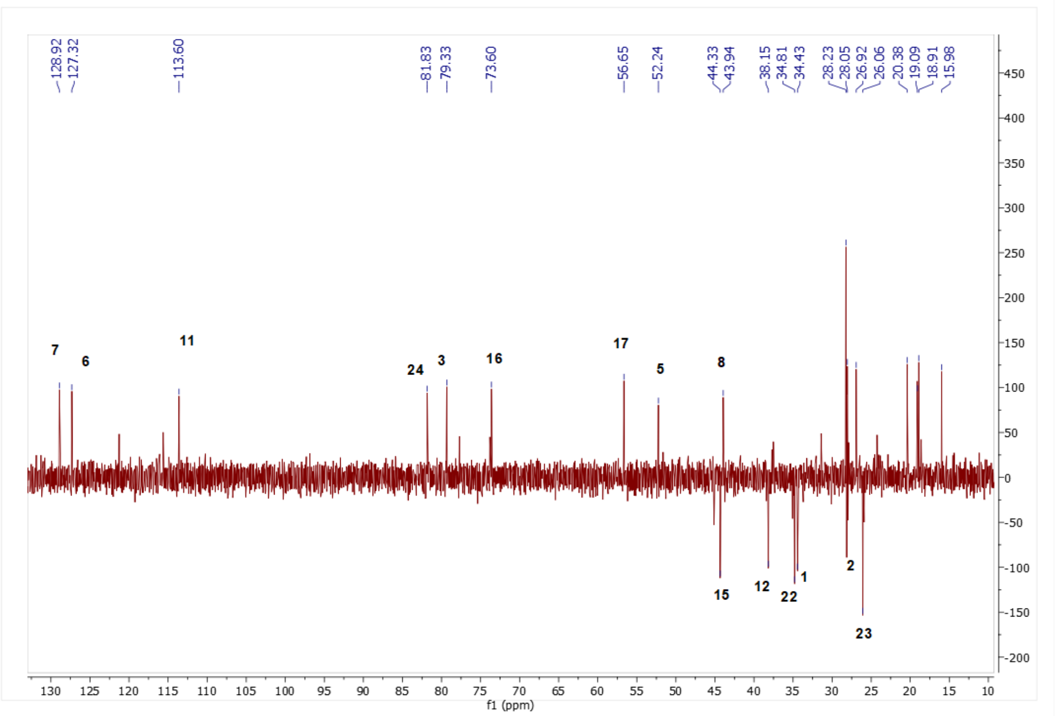

Spectrum 11. DEPT135 spectrum of AG-03.

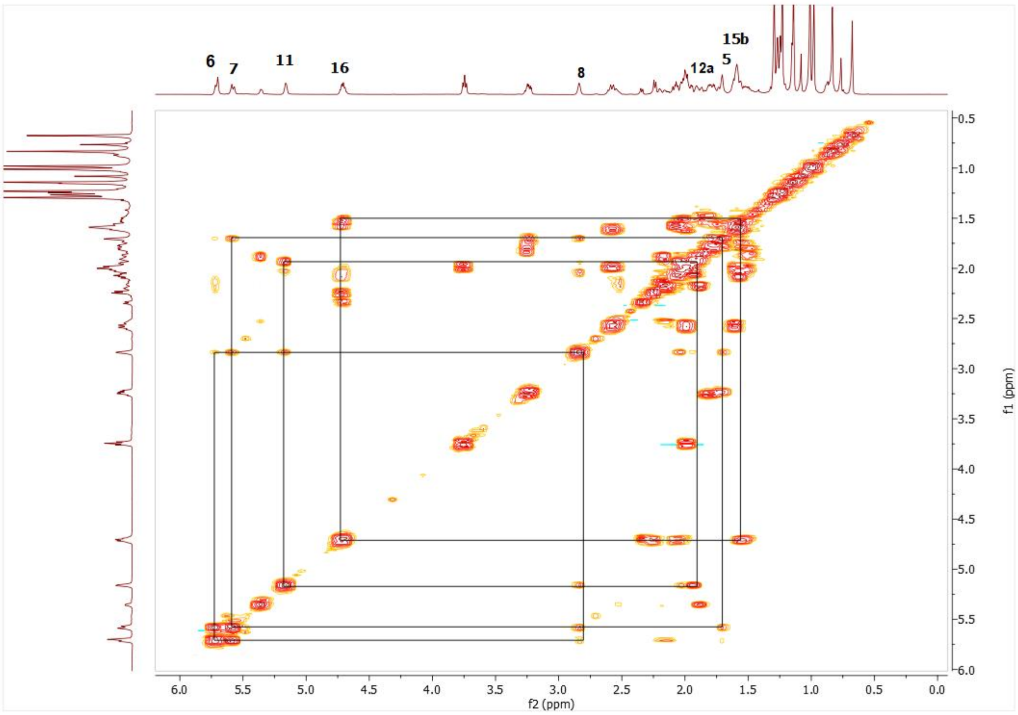

Spectrum 12. COSY spectrum of AG-03.

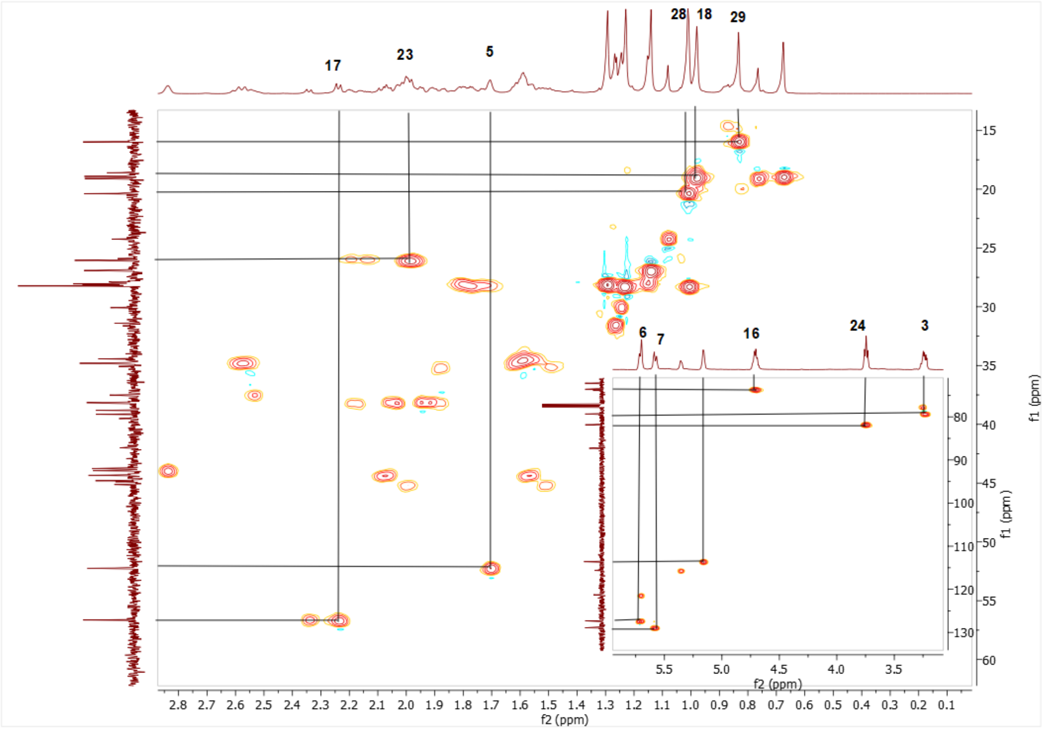

Spectrum 13. HMQC spectrum of AG-03

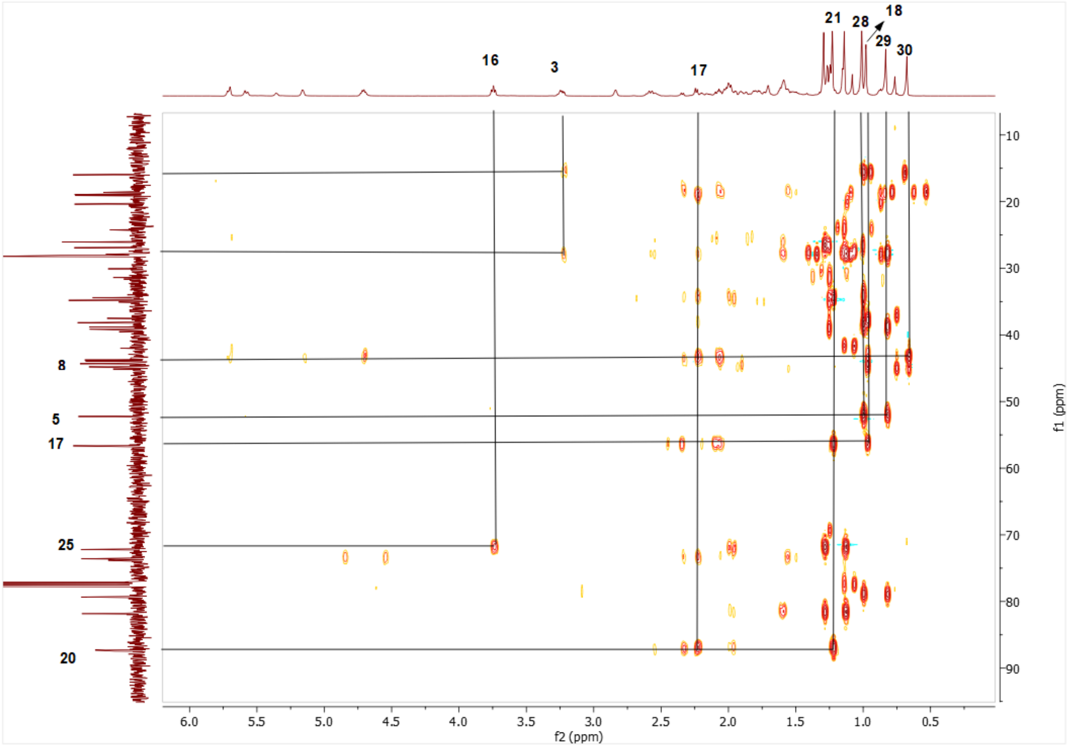

Spectrum 14. HMBC spectrum of AG-03.

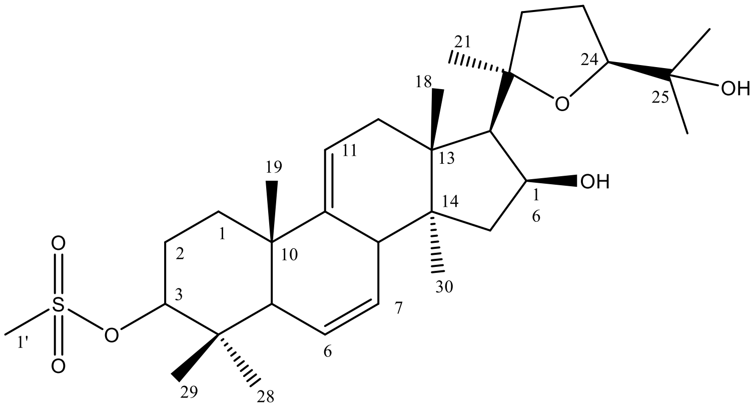

Supplementary Figure 5. Chemical Structure of AG-04

Supplementary Table 4. The $^{13}C$ and $^{1}H$ NMR data of AG-04 (100/400 MHz, δ ppm, in CDCl3).

| H/C | $\delta_C$ (ppm) | $\delta_H$ (ppm), *J* (Hz) |
|---|---|---|
| **1** | 44.0 t | 1.65 m |
| **2** | 25.74 t | 2.06 m, 2.2 m |
| **3** | 90.2 d | 4.35 dd (11.6, 4.9) |
| **4** | 38.6 s | - |
| **5** | 52.2 d | 1.81 brs |
| **6** | 126.1 d | 5.67 dt (10.1, 2.1) |
| **7** | 128.4 d | 5.63 m |
| **8** | 43.7 d | 2.85 brs |
| **9** | 144.8 s | - |
| **10** | 38.3 s | - |
| **11** | 113.9 d | 5.17 dt (5.3, 2.5) |
| **12** | 37.9 t | 1.96 m, 2.05 m |
| **13** | 44.6 s | - |
| **14** | 43.5 s | - |
| **15** | 44.0 t | 1.57 m, 2.08 m |
| **16** | 73.3 d | 4.71 ddd (7.7, 7.7, 5.9) |
| **17** | 56.4 d | 2.24 d (7.6) |
| **18** | 18.7 q | 0.67 s |
| **19** | 20.1 q | 1.05 s |
| **20** | 87.0 s | - |
| **21** | 27.9 q | 1.24 s |
| **22** | 34.6 t | 1.64 m, 2.57 q (10.6) |
| **23** | 25.81 t | 2.0 m |
| **24** | 81.6 d | 3.76 t (7.2) |
| **25** | 72 s | - |
| **26** | 27.8 q | 1.3 s |
| **27** | 26.7 q | 1.15 s |
| **28** | 28.1 q | 1.05 s |
| **29** | 16.5 q | 0.92 s |
| **30** | 18.84 q | 0.98 s |
| **1'** | 39 q | 3.03 s |

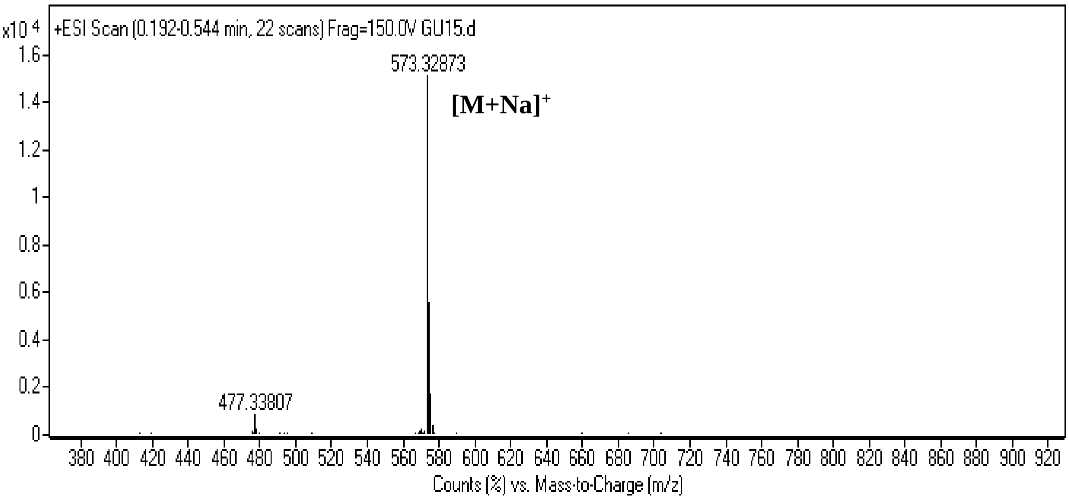

Spectrum 15. HR-ESI-MS Spectrum of AG-04 (positive mode).

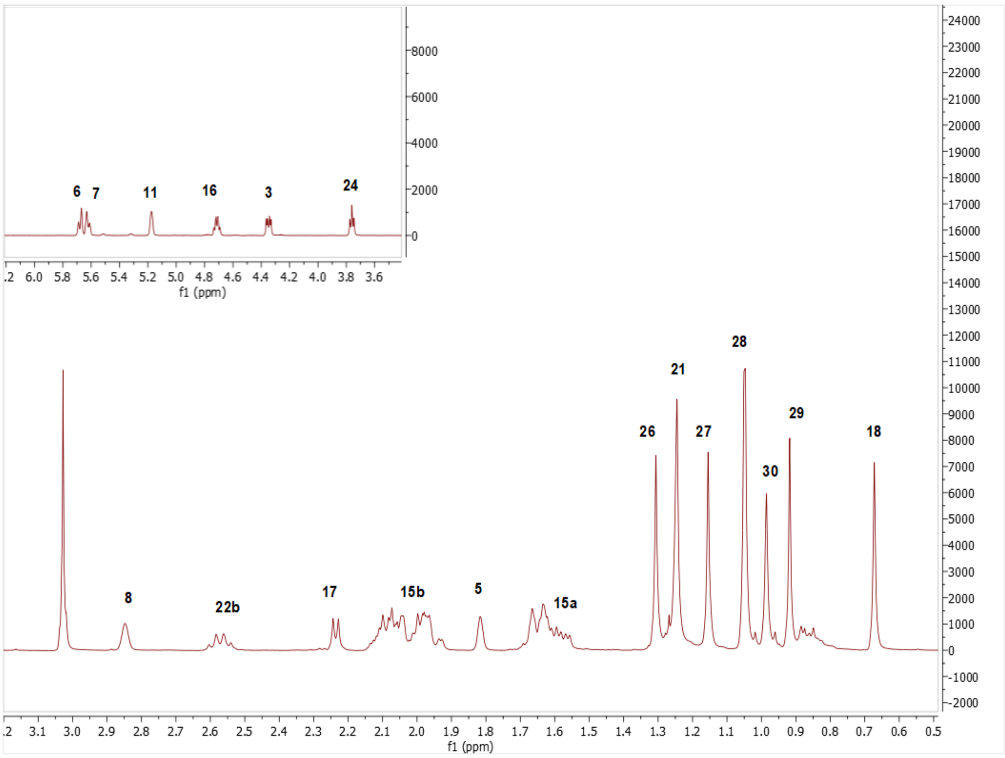

Spectrum 16. $^{1}H$ NMR Spectrum of AG-04.

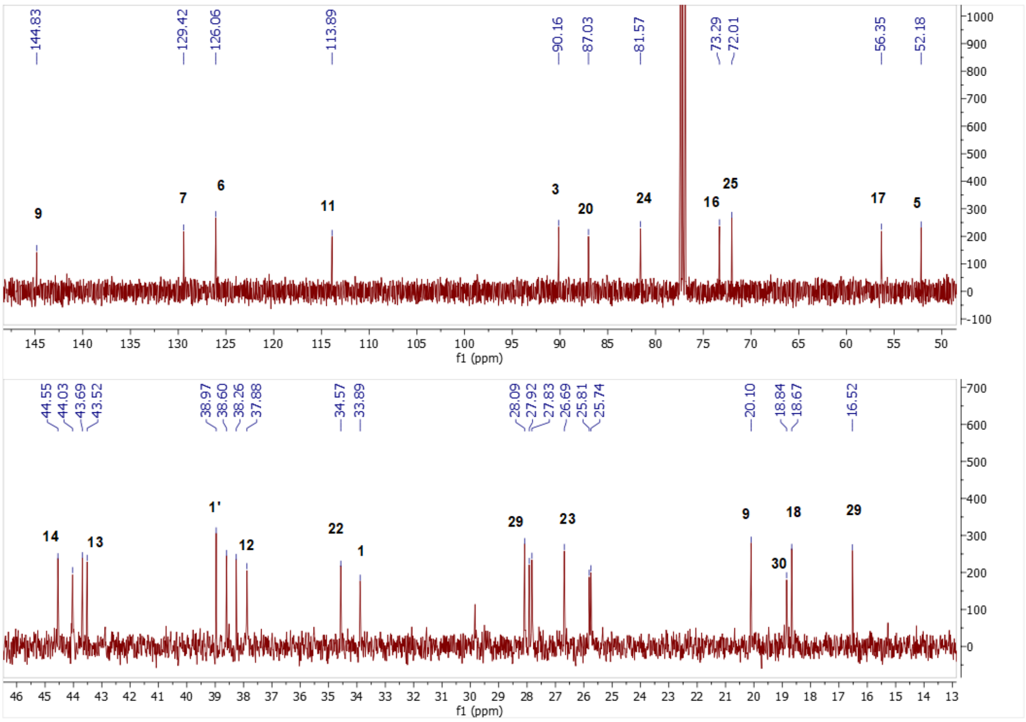

Spectrum 17. $^{13}C$ NMR Spectrum of AG-04.

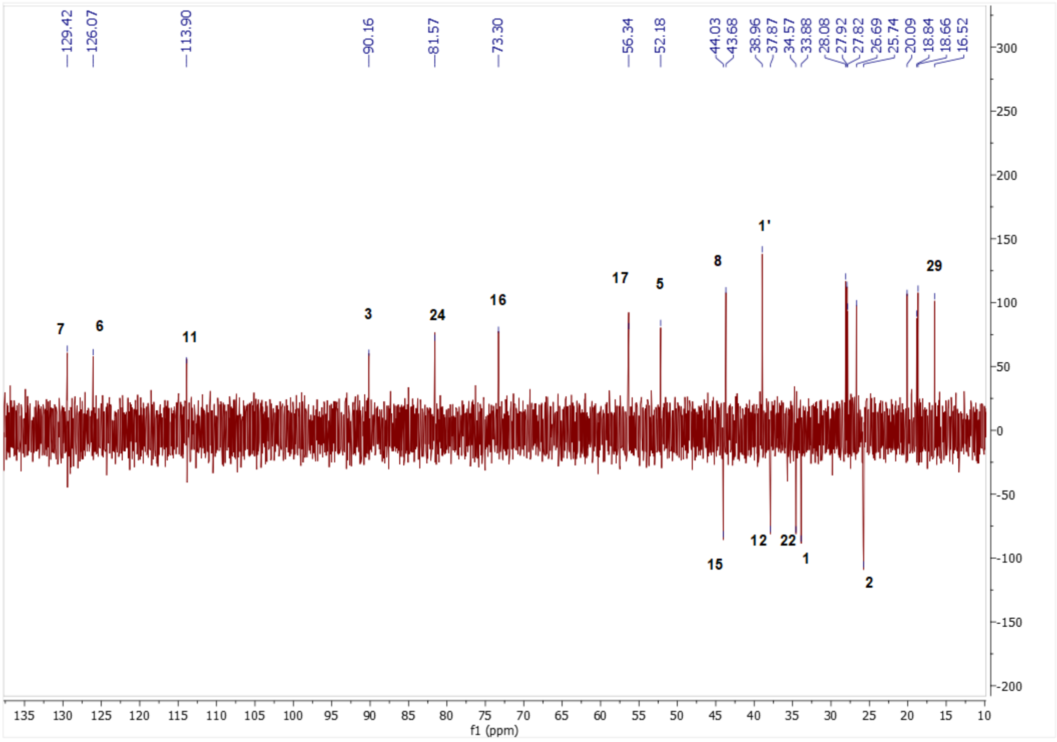

Spectrum 18. DEPT135 spectrum of AG-04

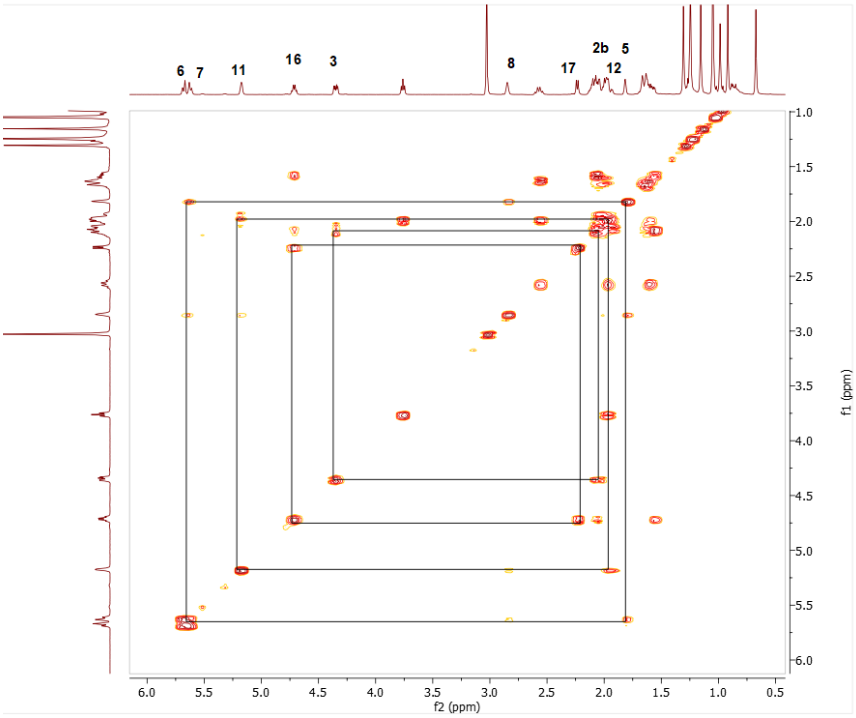

Spectrum 19. COSY spectrum of AG-04.

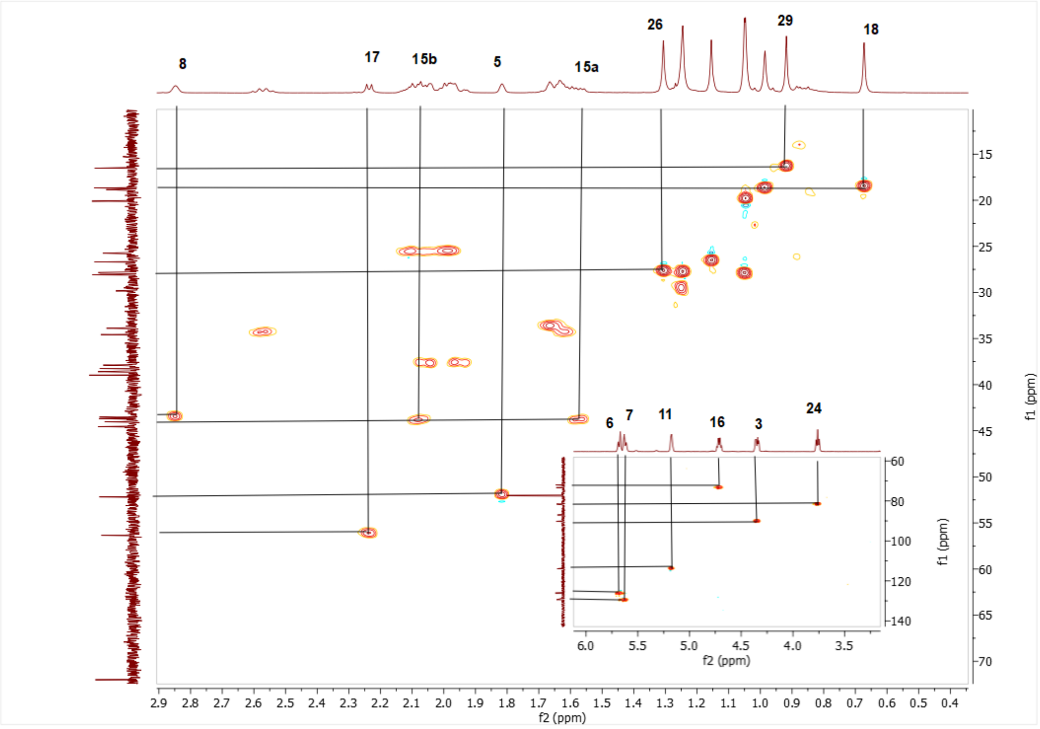

Spectrum 20. HMQC spectrum of AG-04

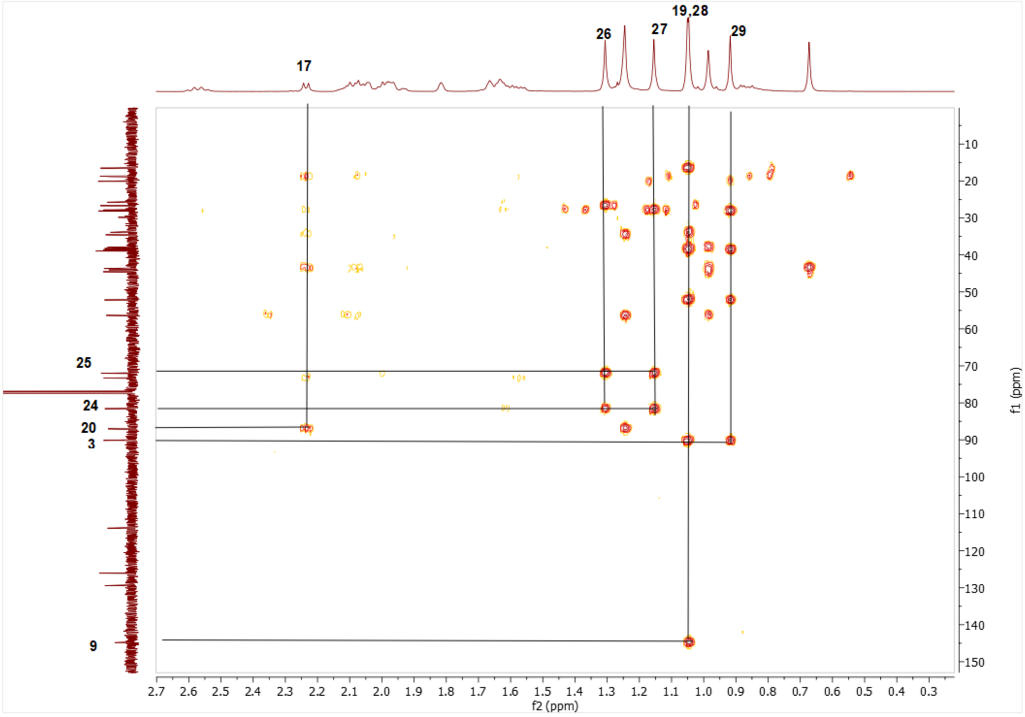

Spectrum 21. HMBC spectrum of AG-04

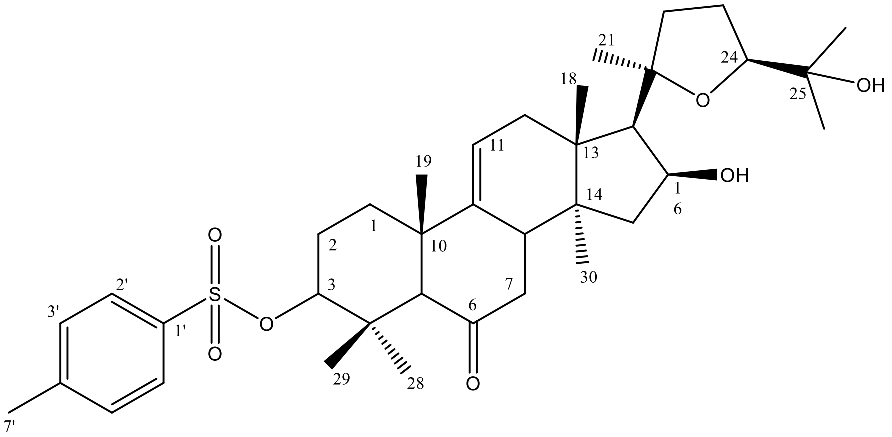

Supplementary Figure 6. Chemical Structure of AG-05

Supplementary Table 5. The $^{13}C$ and $^{1}H$ NMR data of AG-05 (100/400 MHz, δ ppm, in CDCl3).

| H/C | $\delta_C$ (ppm) | $\delta_H$ (ppm), *J* (Hz) |
|---|---|---|
| **1** | 35.9 t | 1.68 m, 1.85 d (1,5) |
| **2** | 25.1 t | 1.91 m |
| **3** | 89.4 d | 4.07 m |
| **4** | 37.7 s | - |
| **5** | 61.7 d | 2.17 brs |
| **6** | 210.4 s | - |
| **7** | 43.98 t | 2.19 m, 2.34 d (5) |
| **8** | 43.8 d | 2.69 brs |
| **9** | 144.5 s | - |
| **10** | 44.3 s | - |

| | | |
|---|---|---|
| **11** | 118.1 d | 5.44 dd (6.3, 1.7) |
| **12** | 44.2 t | 1.43 dd (12.9, 6.3), 1.88 d (4.24) |
| **13** | 44.2 s | - |
| **14** | 44.3 s | - |
| **15** | 37.4 t | 1.95 m, 2.15 m |
| **16** | 73.0 d | 4.68 q (6.8) |
| **17** | 56.3 d | 2.32 brs |
| **18** | 18.2 q | 0.9 s |
| **19** | 23.8 q | 1.04 s |
| **20** | 86.9 s | - |
| **21** | 28.0 q | 1.22 s |
| **22** | 34.5 t | 1.56 dt (12,6), 2.56 q (10.4) |
| **23** | 25.8 d | 1,96 m |
| **24** | 81.4 d | 3.75 td (7.4, 1.6) |
| **25** | 72.0 s | - |
| **26** | 26.8 q | 1.14 s |
| **27** | 27.8 q | 1.29 s |
| **28** | 27.8 q | 0.84 s |
| **29** | 16.1 q | 1.27 d (1.3) |
| **30** | 18.9 q | 0.84 s |
| **1’** | 134.6 s | - |
| **2’** | 127.8 d | 7.77 dd (8.2, 1.7) |
| **3’** | 129.8 d | 7.32 dd (8.4, 1.8) |
| **4’** | 144.6 s | - |
| **5’** | 129.8 d | 7.32 dd (8.4, 1.8) |
| **6’** | 127.8 d | 7.77 dd (8.2, 1.7) |
| **7’** | 21.78 q | 2.43 s |

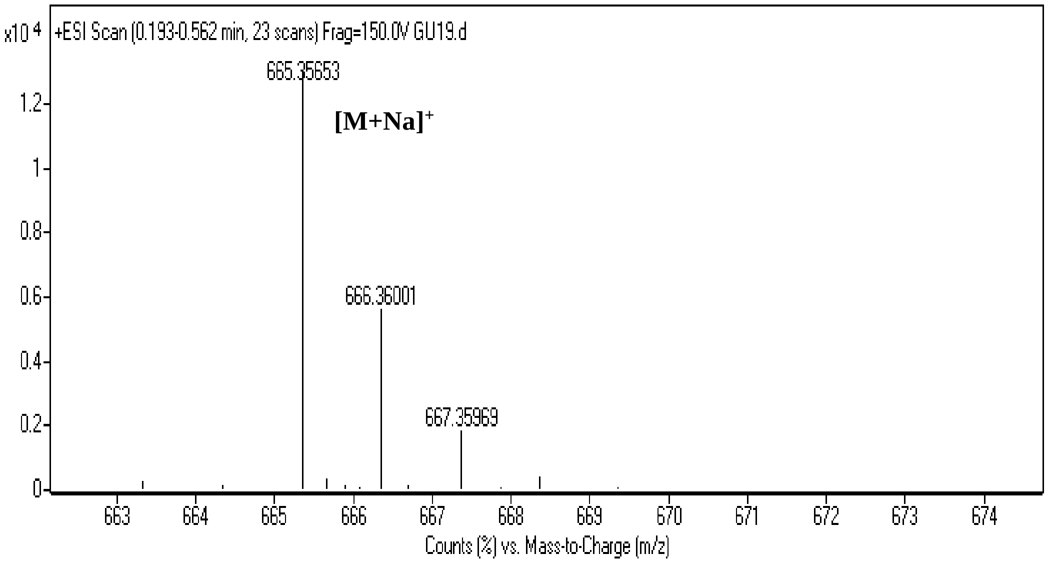

Spectrum 22. HR-ESI-MS Spectrum of AG-05 (positive mode).

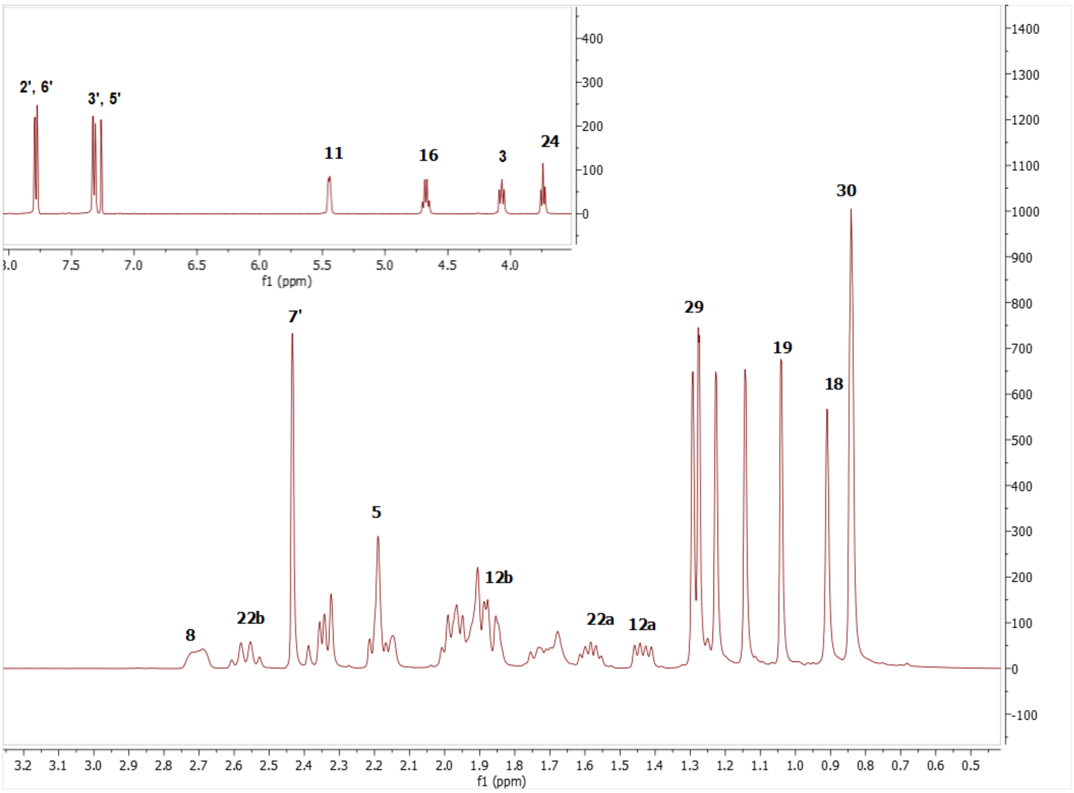

Spectrum 23. $^{1}H$ NMR Spectrum of AG-05.

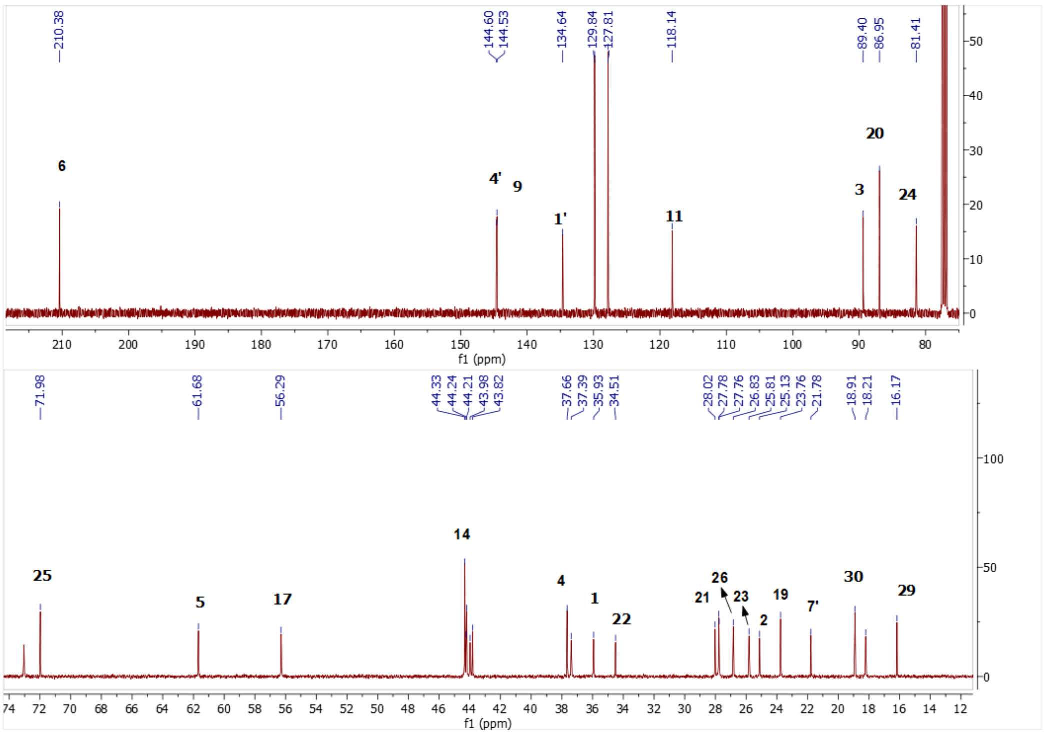

Spectrum 24. $^{13}C$ NMR Spectrum of AG-05.

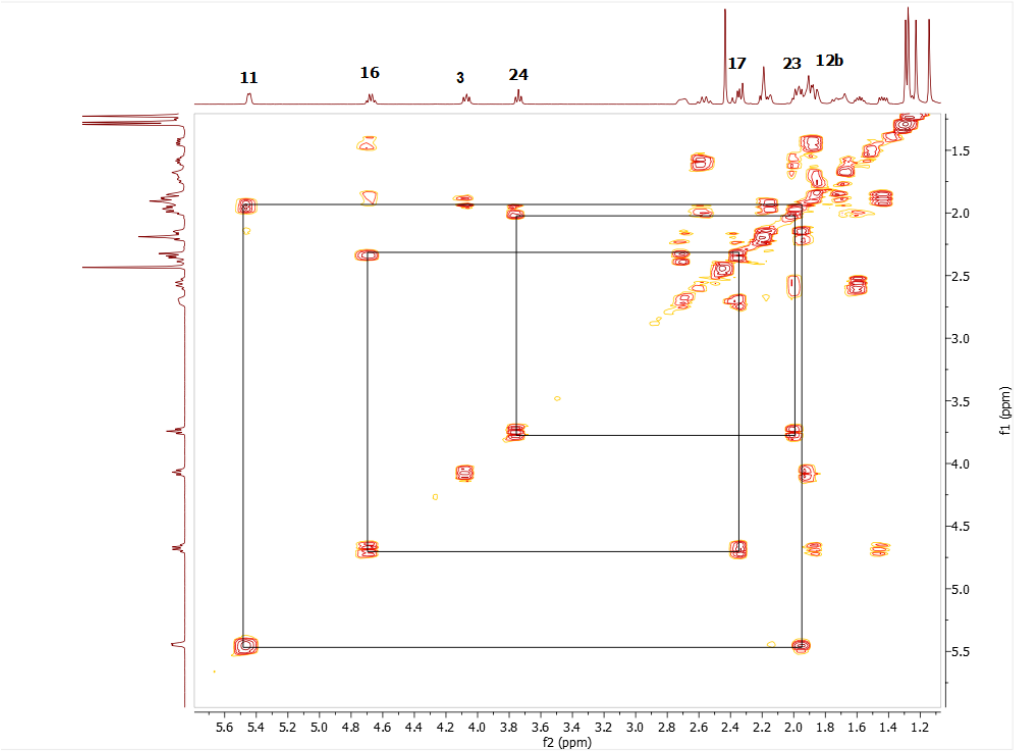

Spectrum 25. COSY spectrum of AG-05

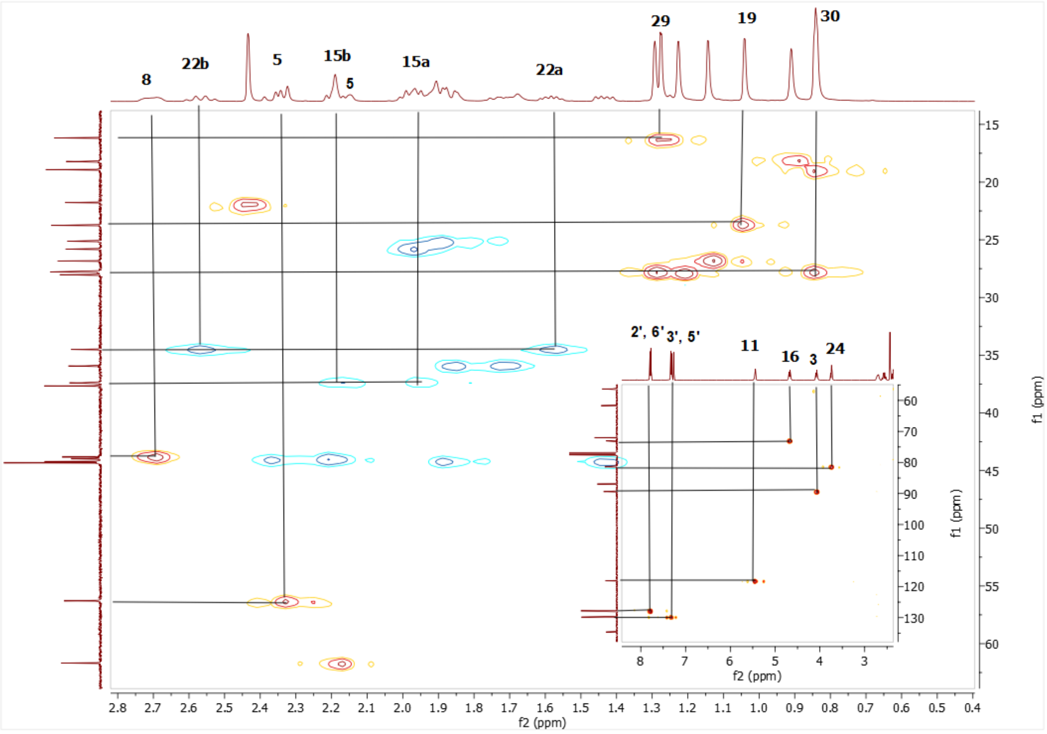

Spectrum 26. HSQC spectrum of AG-05.

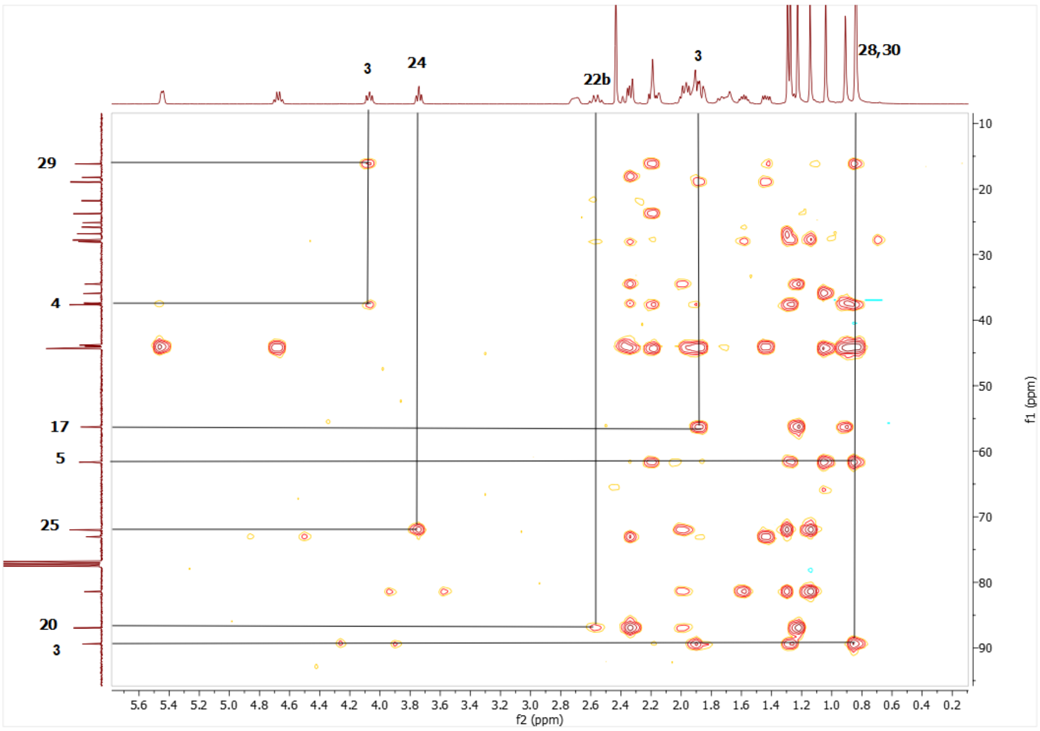

Spectrum 27. HMBC spectrum of AG-05.

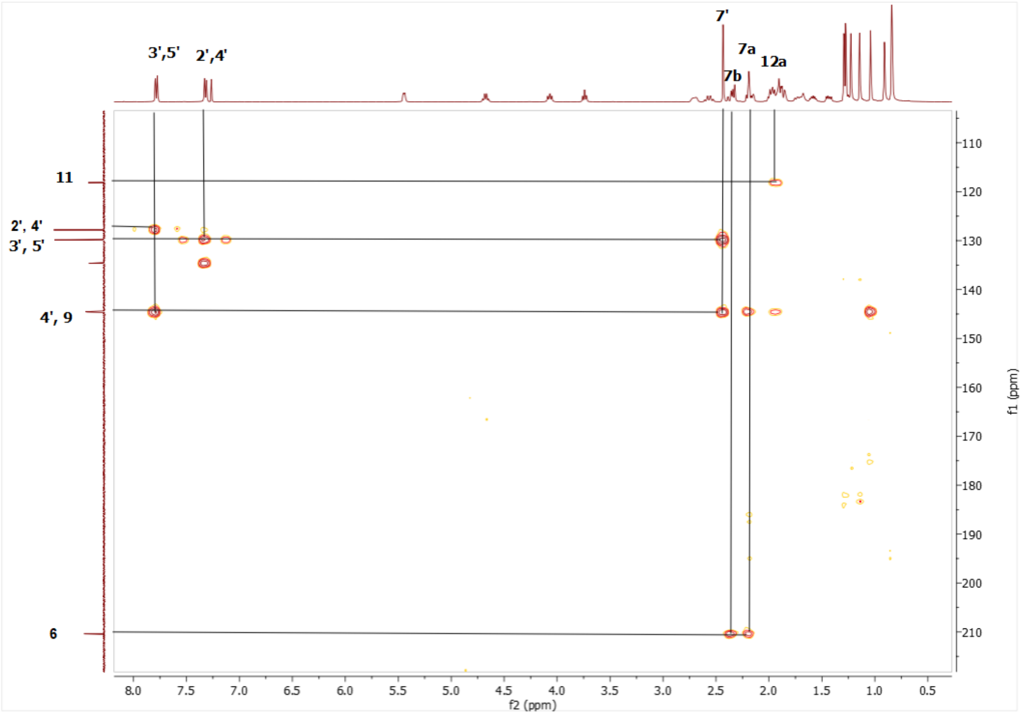

Spectrum 28. HMBC spectrum of AG-05.

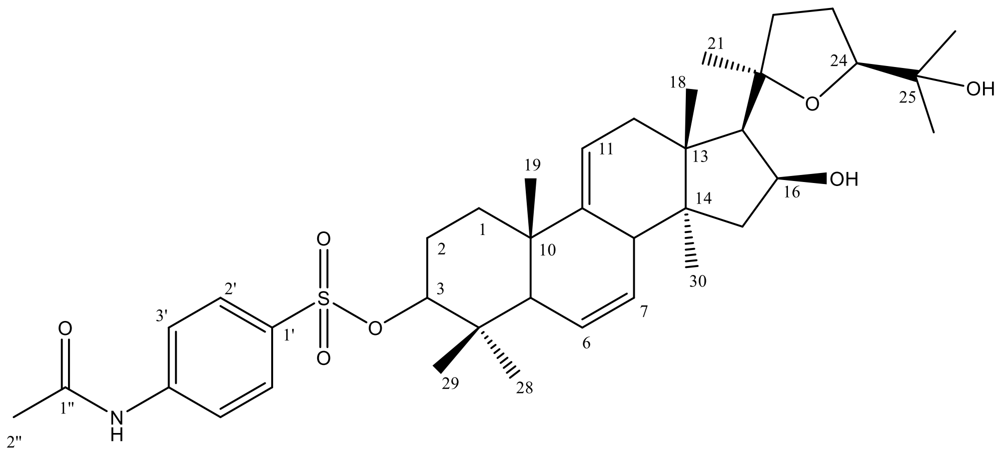

Supplementary Figure 7. Chemical Structure of AG-06

Supplementary Table 6. The $^{13}C$ and $^{1}H$ NMR data of AG-06 (100/400 MHz, δ ppm, in CDCl3).

| H/C | $\delta_C$ (ppm) | $\delta_H$ (ppm), *J* (Hz) |
|---|---|---|
| **1** | 33.9 t | 1.55 m |
| **2** | 25.4 t | 1.91 m |
| **3** | 90.9 d | 4.15 dd (9.8, 6.6) |
| **4** | 38.6 s | - |
| **5** | 52.1 d | 1.69 d (4.2) |
| **6** | 126.1 d | 5.58 d (10.4) |
| **7** | 129.2 d | 5.53 dt (10.4, 3) |
| **8** | 43.6 d | 2.8 brs |
| **9** | 144.8 s | |
| **10** | 38.2 s | - |
| **11** | 113.8 d | 5.13 t (5.3) |
| **12** | 37.8 d | 1.9 m, 2.0 m |
| **13** | 44.5 s | - |
| **14** | 43.4 s | - |
| **15** | 43.8 t | 1.53 m, 2.0 m |
| **16** | 73.3 d | 4.69 ddd (6.5, 6.5, 6) |
| **17** | 56.3 d | 2.12 d (7.7) |
| **18** | 18.6 q | 0.62 s |
| **19** | 20.0 q | 0.97 s |
| **20** | 86.9 s | - |
| **21** | 28.0 q | 1.22 s |
| **22** | 34.5 t | 1.59 m, 2.55 q (10.5) |
| **23** | 25.8 t | 1.99 m |
| **24** | 81.5 d | 3.74 t (7.1, 7.1) |
| **25** | 72.0 s | - |
| **26** | 27.6 q | 1.28 s |
| **27** | 26.7 q | 1.14 s |
| **28** | 27.7 q | 0.73 s |

| | | |
|---|---|---|
| **29** | 16.5 q | 0.83 s |
| **30** | 18.8 q | 0.95 s |
| **1’** | 131.8 s | |
| **2’** | 129.0 d | 7.8 d (8.4) |
| **3’** | 119.3 d | 7.7 d (8.8) |
| **4’** | 143.2 s | |
| **5’** | 119.3 d | 7.7 d (8.8) |
| **6’** | 129.0 d | 7.8 d (8.4) |
| **1’’** | 169.3 | - |
| **2’’** | 24.8 q | 2.18 s |

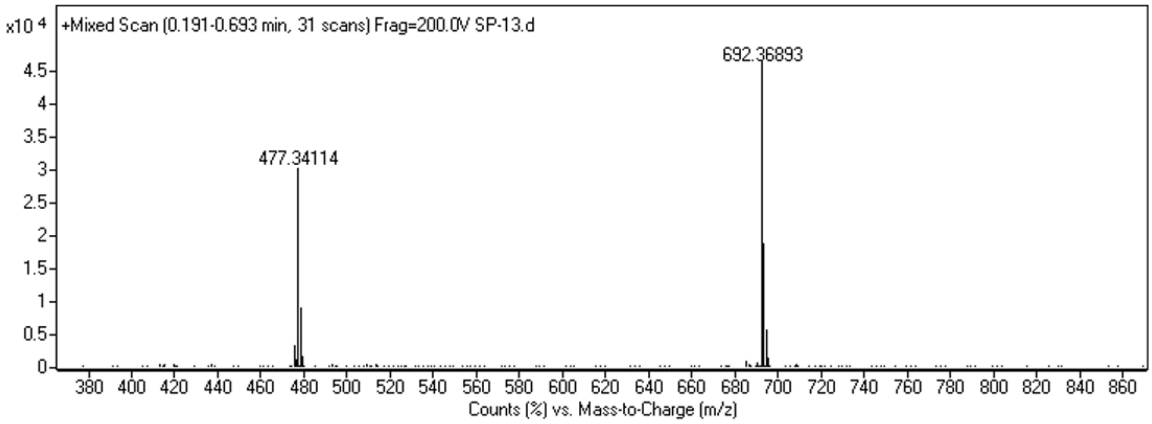

Spectrum 29. HR-ESI-MS Spectrum of AG-06 (positive mode).

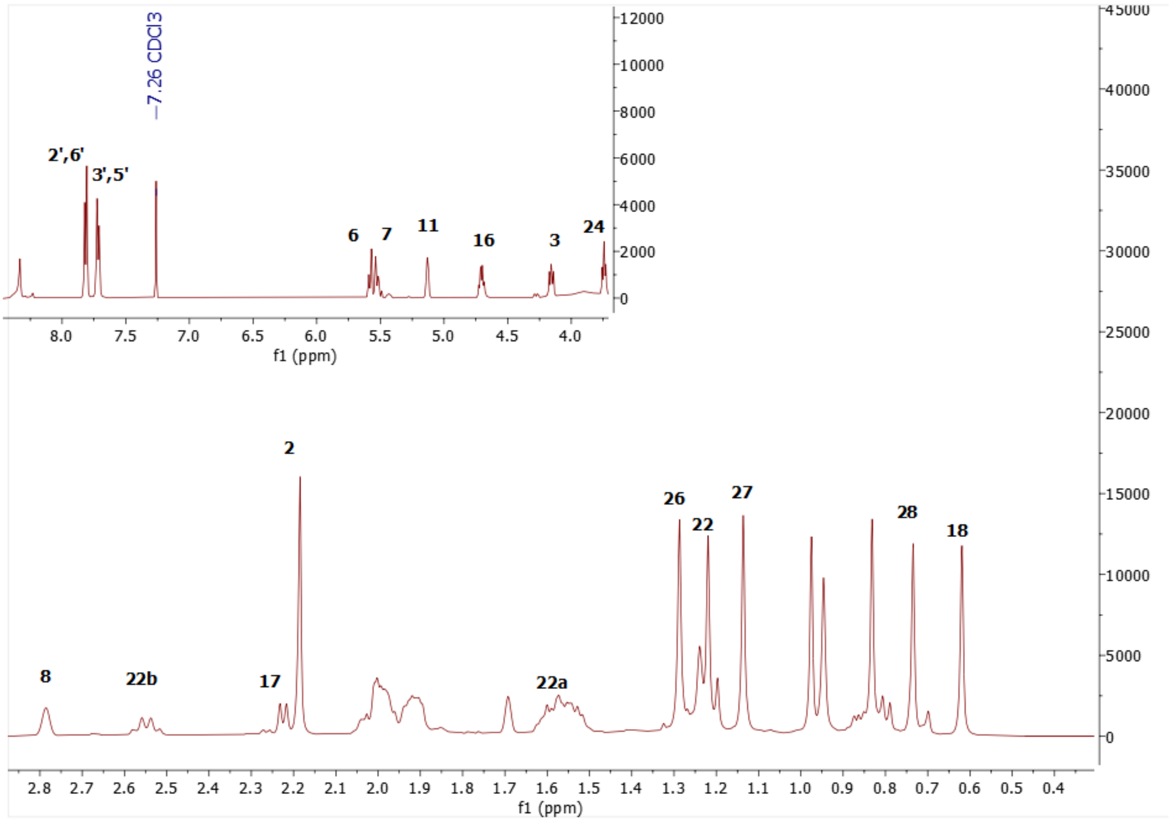

Spectrum 30. $^{1}H$ NMR Spectrum of AG-06.

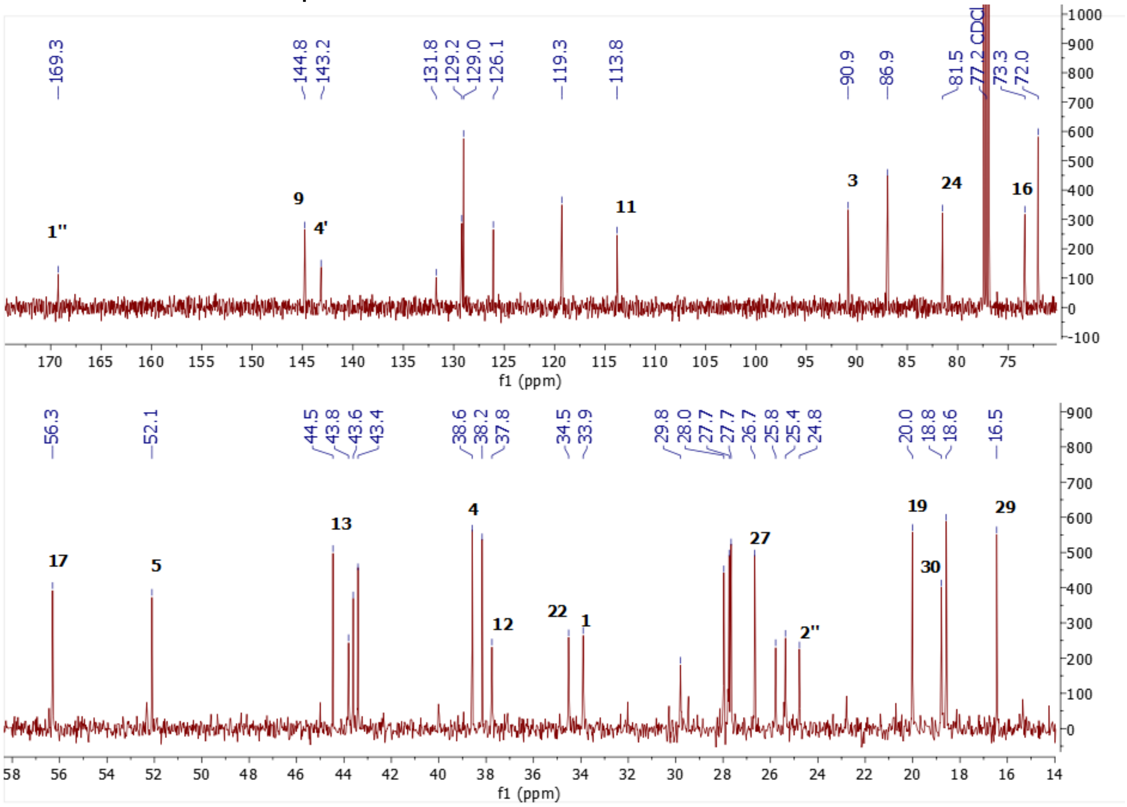

Spectrum 31. $^{13}C$ NMR Spectrum of AG-06.

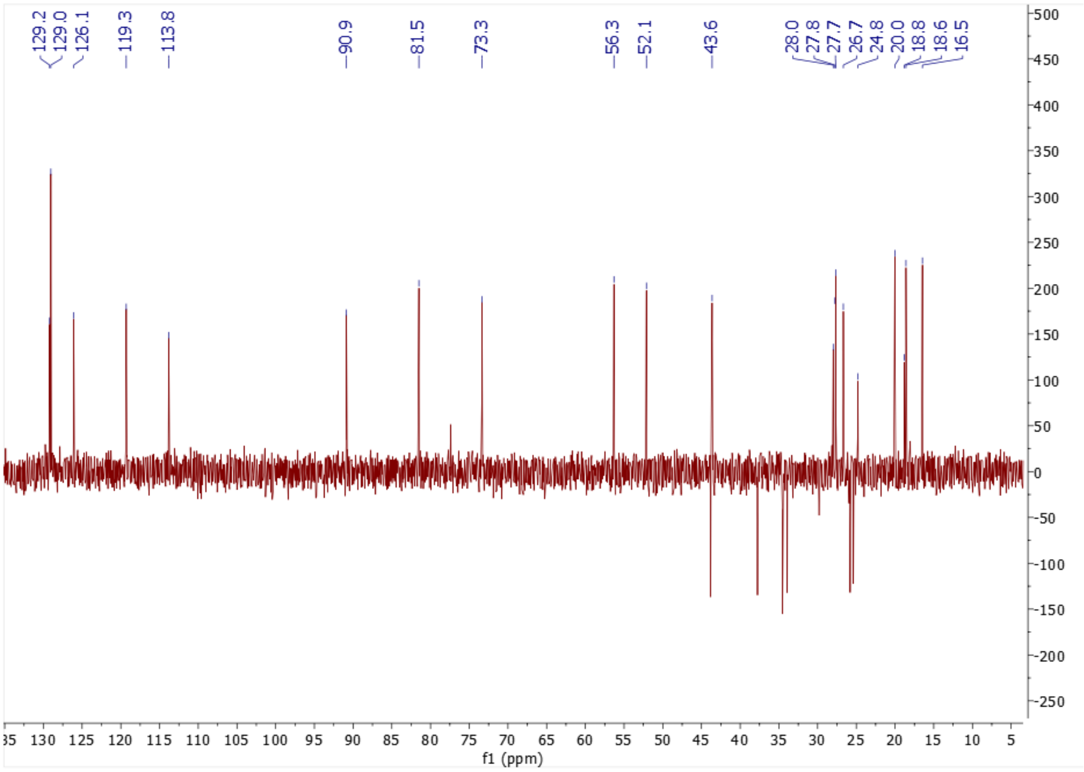

Spectrum 32. DEPT135 spectrum of AG-06

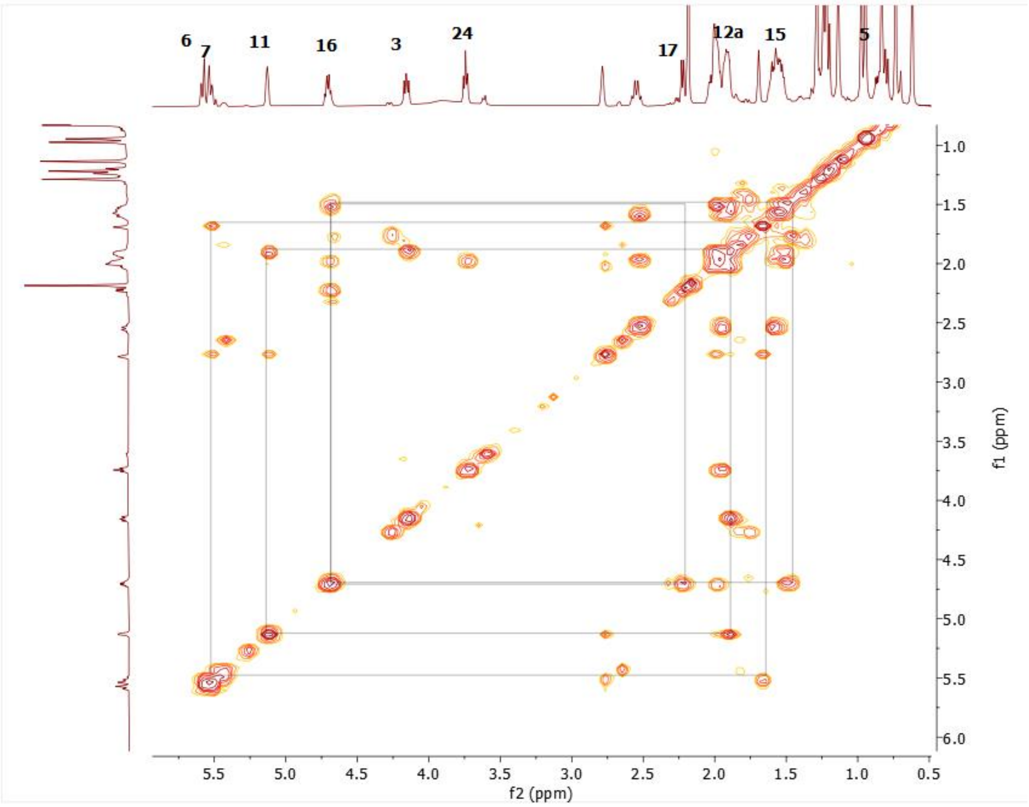

Spectrum 33. COSY spectrum of AG-06

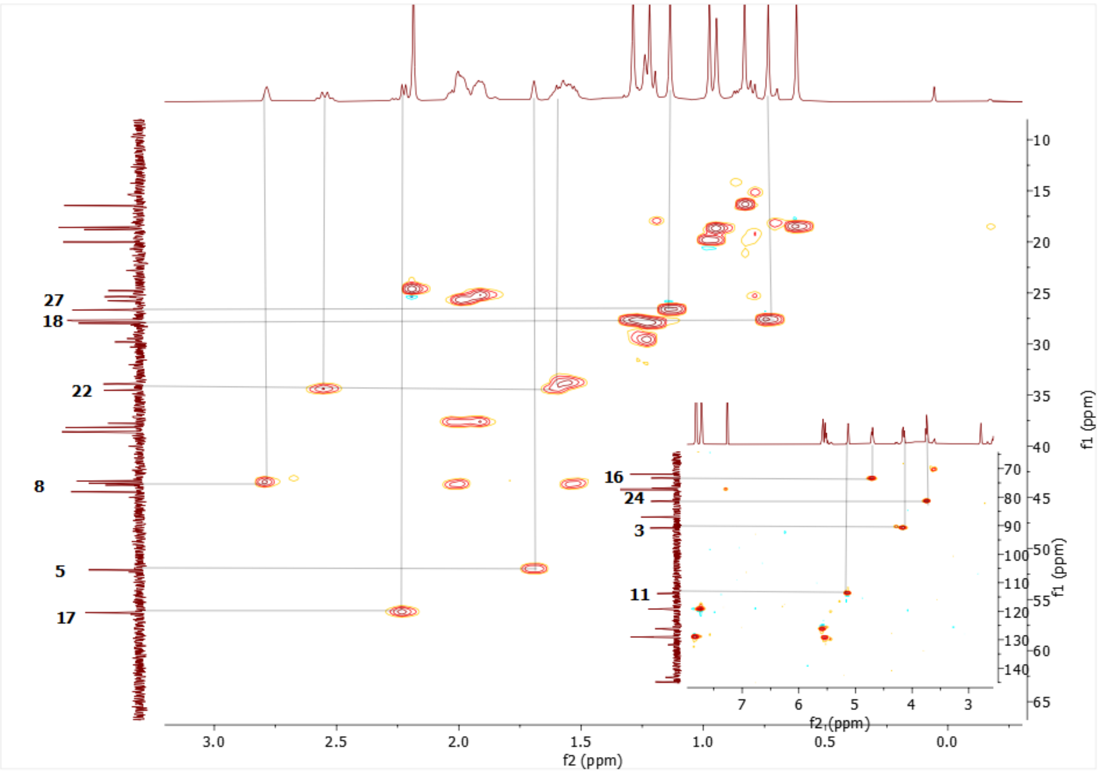

Spectrum 34. HSQC spectrum of AG-06.

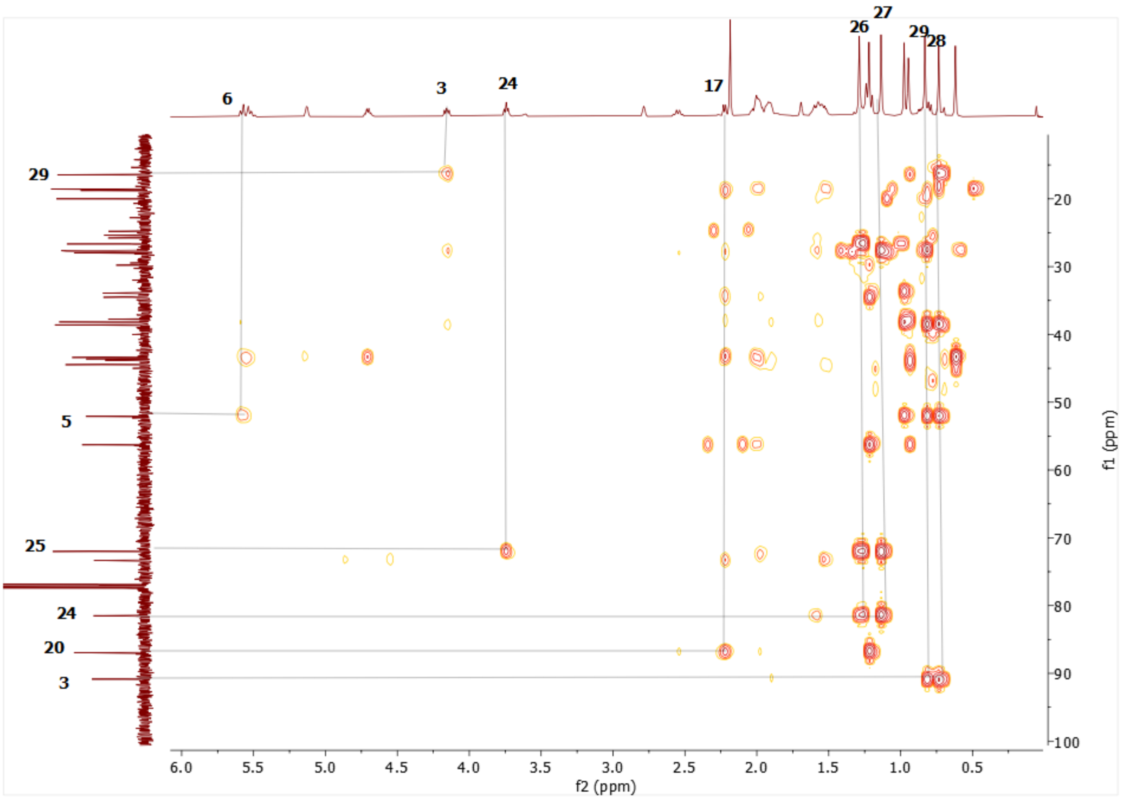

Spectrum 35. HMBC spectrum of AG-06.

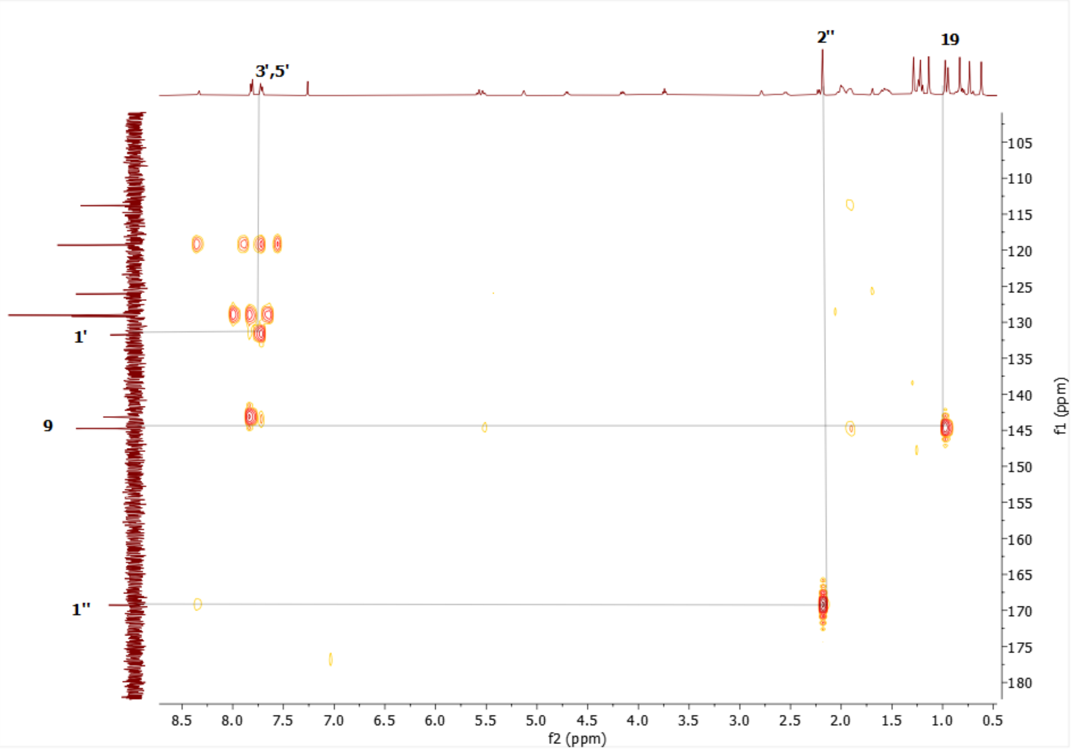

Spectrum 36. HMBC spectrum of AG-06

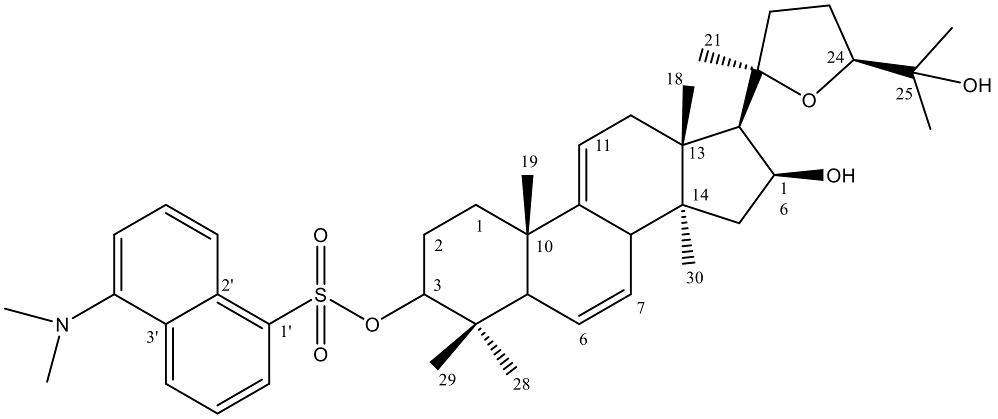

Supplementary Figure 8. Chemical Structure of AG-07

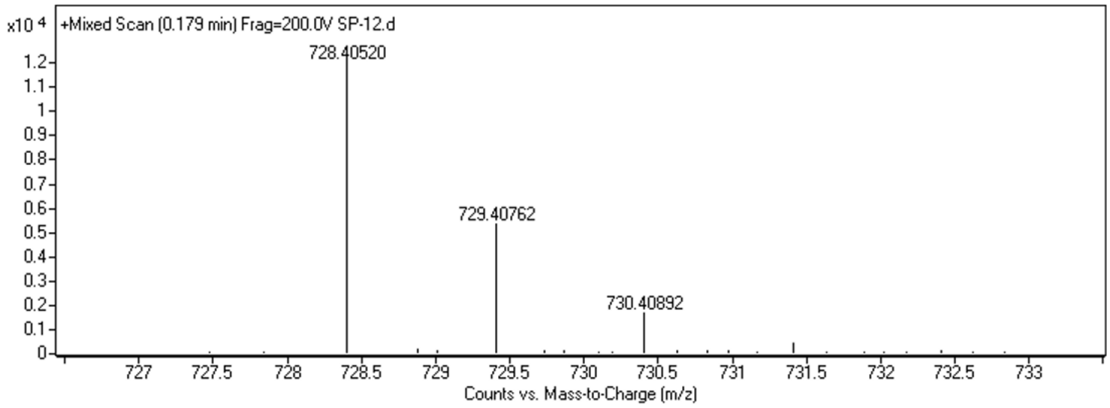

Spectrum 37. HR-ESI-MS Spectrum of AG-07 (positive mode).

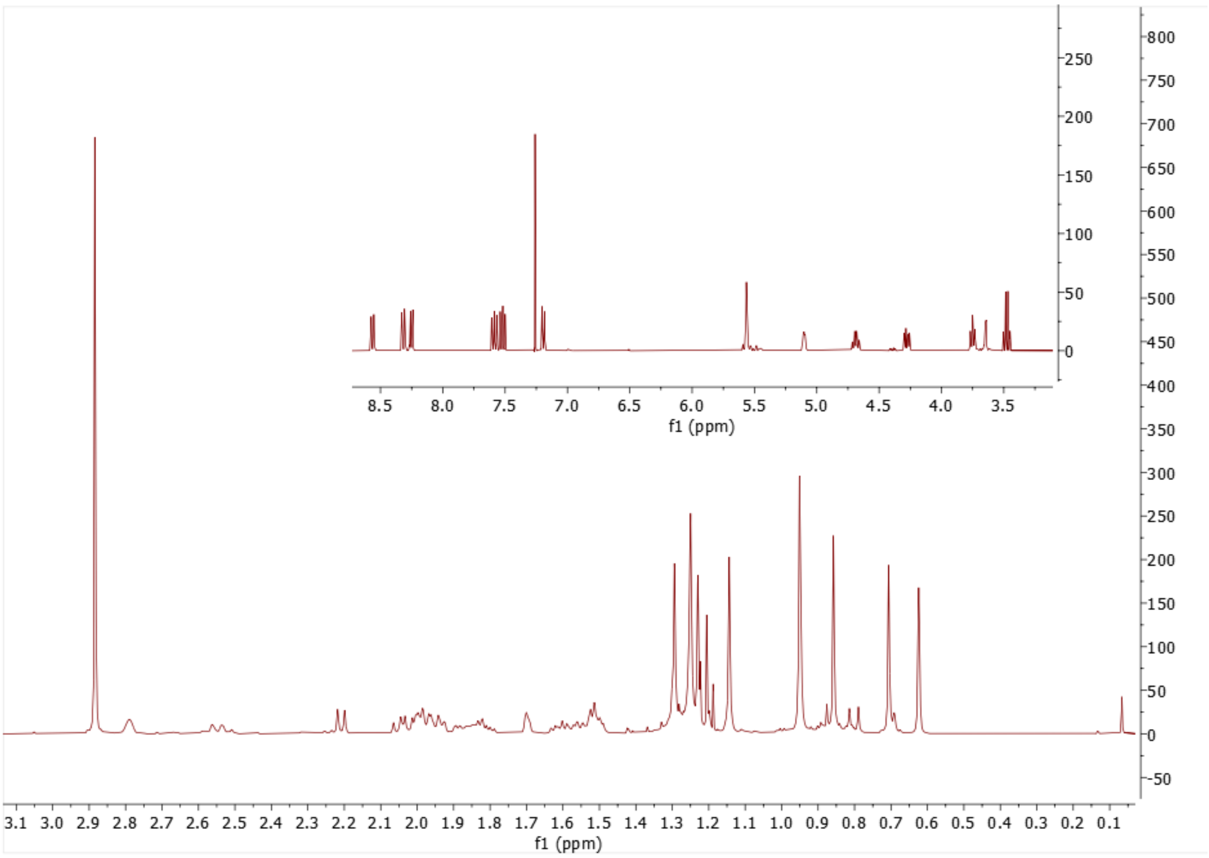

Spectrum 38. $^{1}H$ NMR Spectrum of AG-07.

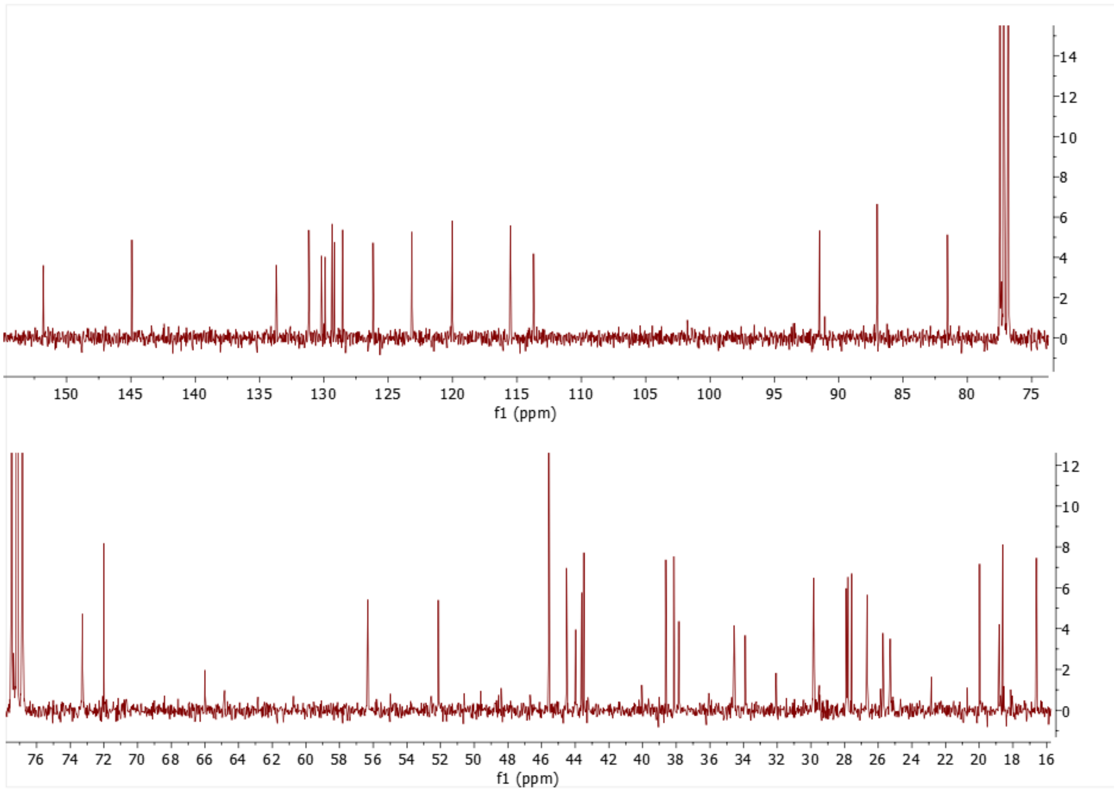

Spectrum 39. $^{13}C$ NMR Spectrum of AG-07.

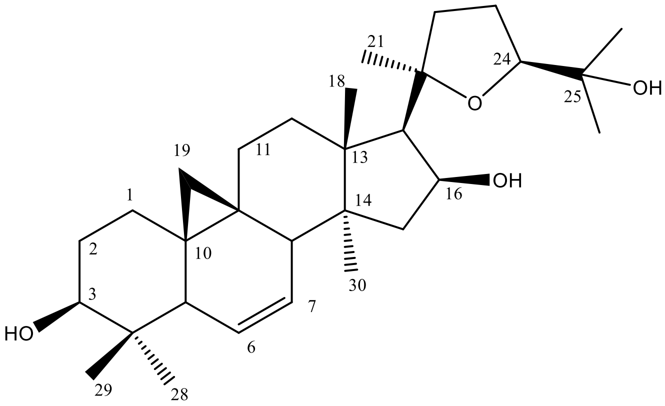

Supplementary Figure 9. Chemical Structure of CG-02

Supplementary Table 7. The $^{13}C$ and $^{1}H$ NMR data of CG-02 (100/400 MHz, δ ppm, in CDCl3).

| **H/C** | **$\delta_C$ (ppm)** | **$\delta_H$ (ppm), *J* (Hz)** |
|---|---|---|
| **1** | 29.8 t | 1.25 s, 1.41 m |

| | | |
|---|---|---|
| **2** | 30.1 t | 1.82 m, 1.59 m |
| **3** | 78.5 d | 3.32 dd (11.4, 4.6) |
| **4** | 40.3 s | - |
| **5** | 46.6 d | 1.85 m |
| **6** | 126.6 d | 5.63 d (10.6) |
| **7** | 129.2 d | 5.48 brs |
| **8** | 43.4 d | 2.71 d (7.9) |
| **9** | 20.6 s | - |
| **10** | 28.4 s | - |
| **11** | 25.2 t | 1.40 m, 1.88 m |
| **12** | 33.4 t | 1.62 m, 1.42 m |
| **13** | 45.2 s | - |
| **14** | 48.5 s | - |
| **15** | 43.6 t | 1.84 m, 1.52 m |
| **16** | 73.5 d | 4.69 q (7.2) |
| **17** | 56.4 d | 2.27 d (7.6) |
| **18** | 18.1 q | 1.23 s |
| **19** | 18.7 t | -0.16 d (4.3), 0.74 m |
| **20** | 87.3 s | - |
| **21** | 28.1 q | 1.22 s |
| **22** | 34.6 t | 1.6 m, 2.59 q (10.6) |
| **23** | 25.9 t | 2 m |
| **24** | 81.4 d | 3.74 t (7.1) |
| **25** | 72.1 s | - |
| **26** | 27.9 q | 1.31 s |
| **27** | 26.7 q | 1.15 s |
| **28** | 14.5 q | 0.78 s |
| **29** | 25.6 | 1.05 s |
| **30** | 18.2 | 1.23 s |

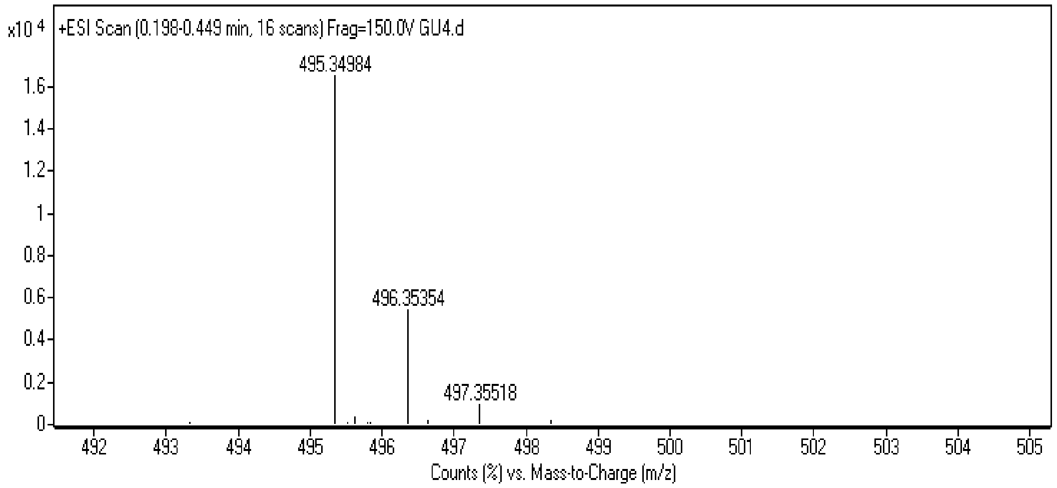

Spectrum 40. HR-ESI-MS Spectrum of CG-02 (positive mode).

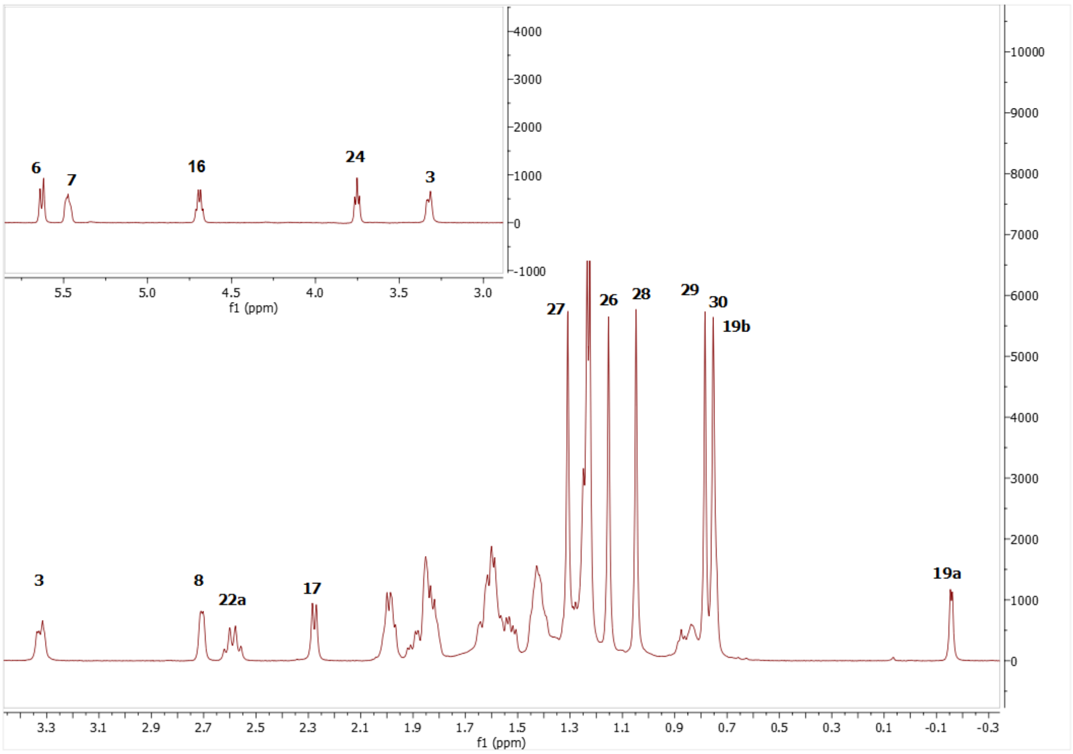

Spectrum 41. $^{1}$H NMR Spectrum of CG-02.

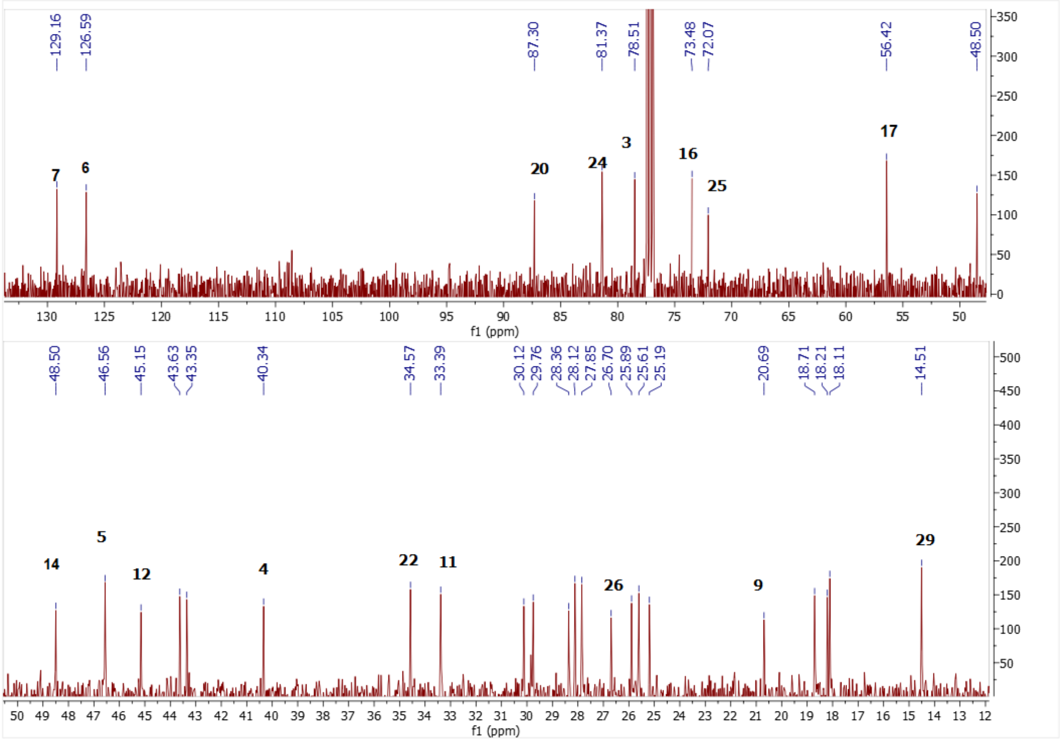

Spectrum 42. $^{13}$C NMR Spectrum of CG-02.

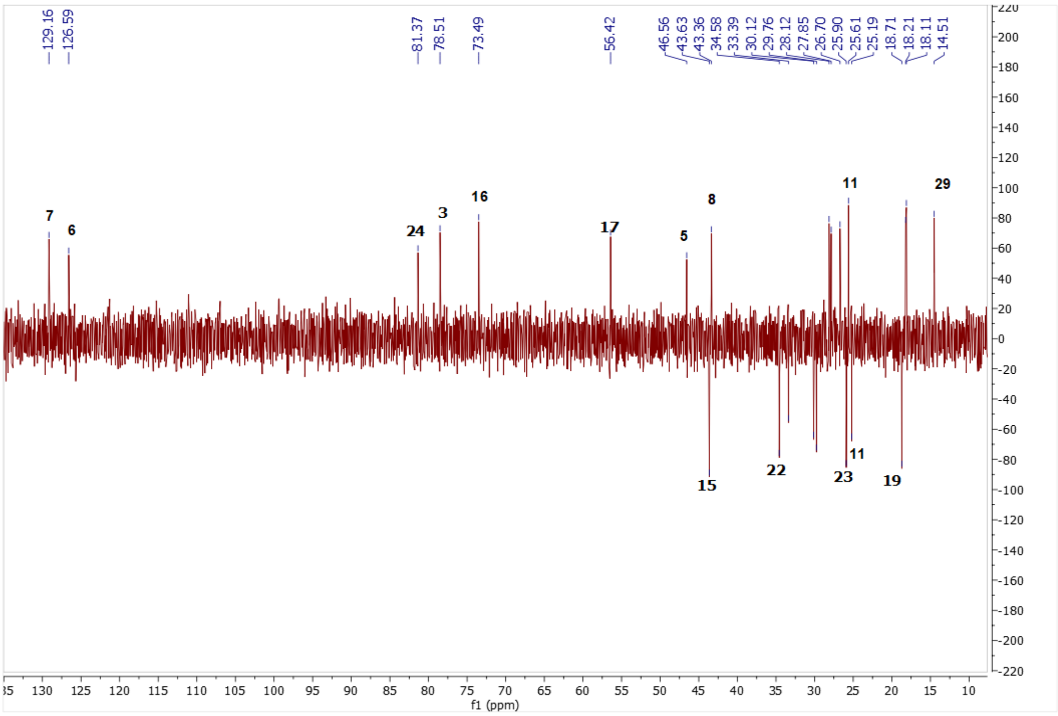

Spectrum 43. DEPT135 spectrum of CG-02.

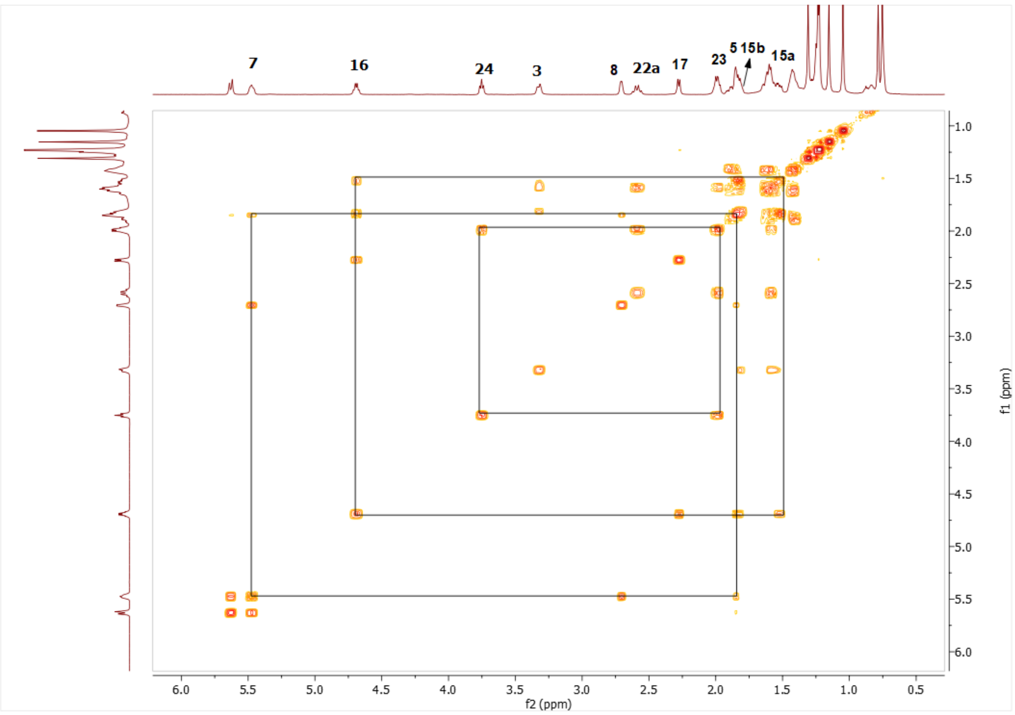

Spectrum 44. COSY spectrum of CG-02.

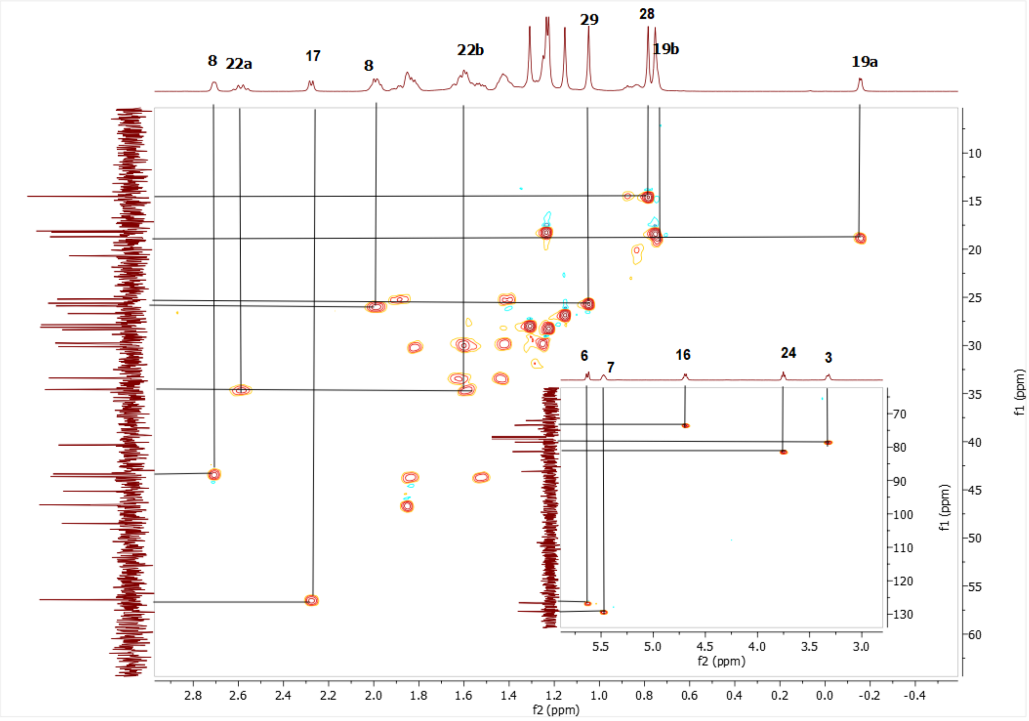

Spectrum 45. HMQC spectrum of CG-02.

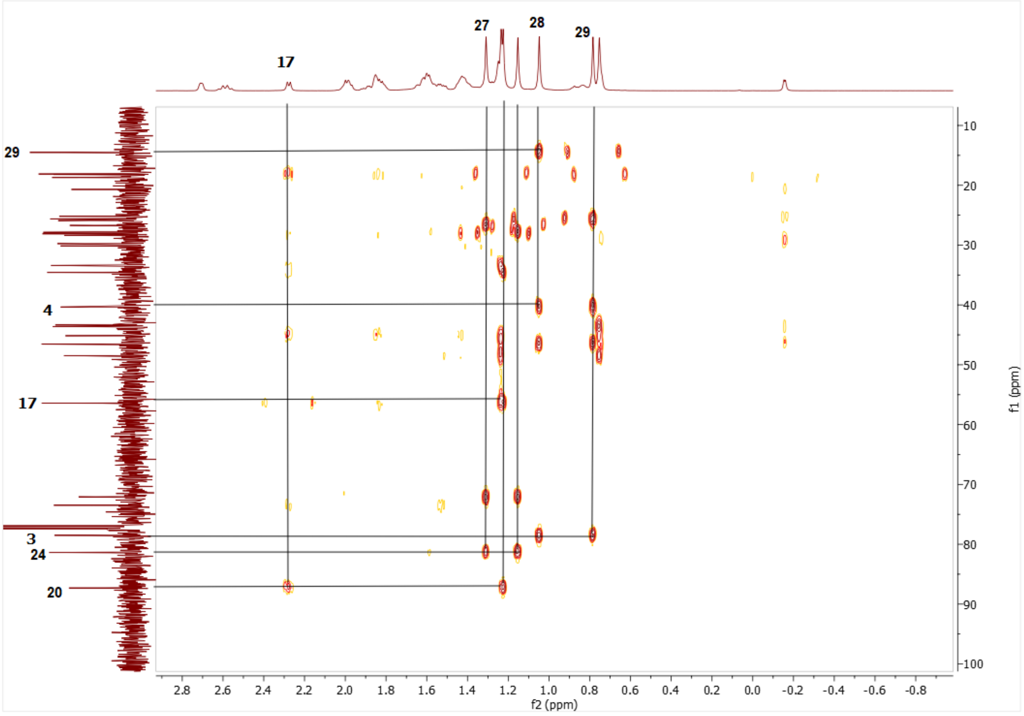

Spectrum 46. HMBC spectrum of CG-02.

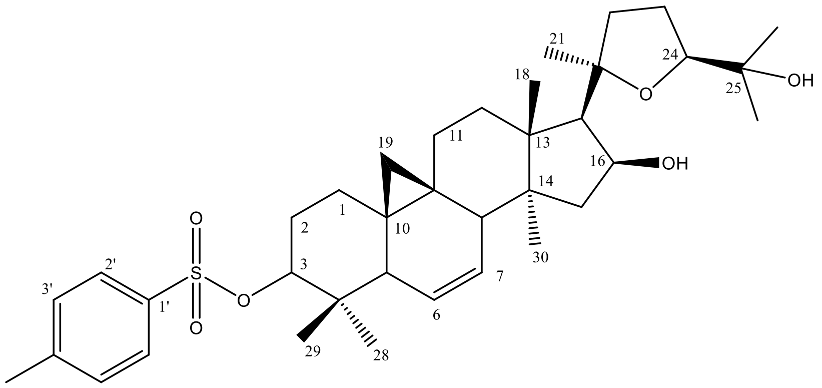

Supplementary Figure 10. Chemical Structure of CG-03

Supplementary Table 8. The $^{13}C$ and $^{1}H$ NMR data of CG-03 (100/400 MHz, δ ppm, in CDCl3).

| H/C | $\delta_C$ (ppm) | $\delta_H$ (ppm), *J* (Hz) |
|---|---|---|
| **1** | 29.5 t | 1.38 m, 1.58 m |
| **2** | 27.8 t | 1.77 m, 1.89 m |
| **3** | 90.2 s | 4.31 dd (11.8, 4.6) |
| **4** | 40 s | - |
| **5** | 46.7 d | 1.86 m |
| **6** | 125.6 d | 5.51 d (10.7) |
| **7** | 129.8 d | 5.46 ddd (10.7 ,6, 3) |
| **8** | 43.3 d | 2.68 dd (6, 2.5) |
| **9** | 20.7 s | - |
| **10** | 27.9 s | - |
| **11** | 25.2 t | 1.34 m, 1.85 m |
| **12** | 33.3 t | 1.41 m,1.57 m |
| **13** | 48.6 s | - |
| **14** | 45.1 s | - |
| **15** | 43.5 t | 1.48 m, 1.79 m |
| **16** | 73.4 d | 4.68 ddd (7.7, 7.7, 6.1) |
| **17** | 56.4 d | 2.26 d (7.7) |
| **18** | 18.1 q | 1.21 s |
| **19** | 18.6 t | -0.17, 0.72 d (4.2) |
| **20** | 87.2 s | - |
| **21** | 28.2 q | 1.19 s |
| **22** | 34.5 t | 1.56 m, 2.58 q (10.5) |
| **23** | 25.9 t | 2 td (10.5, 9, 5) |
| **24** | 81.3 d | 3.74 (7.1) |
| **25** | 72.0 s | - |
| **26** | 28.8 q | 1.29 s |
| **27** | 26.7 q | 1.13 s |
| **28** | 25.4 q | 0.81 s |
| **29** | 15.4 q | 0.8 s |
| **30** | 18.2 q | 0.71 s |
| **1’** | 132.9 s | - |
| **2’** | 127.8 d | 7.79 d (8.2) |
| **3’** | 129.8 d | 7.32 d (8.2) |
| **4’** | 144.5 s | - |
| **5’** | 129.8 d | 7.32 d (8.2) |
| **6’** | 127.8 d | 7.79 d (8.2) |
| **7’** | 21.2 q | 2.43 s |

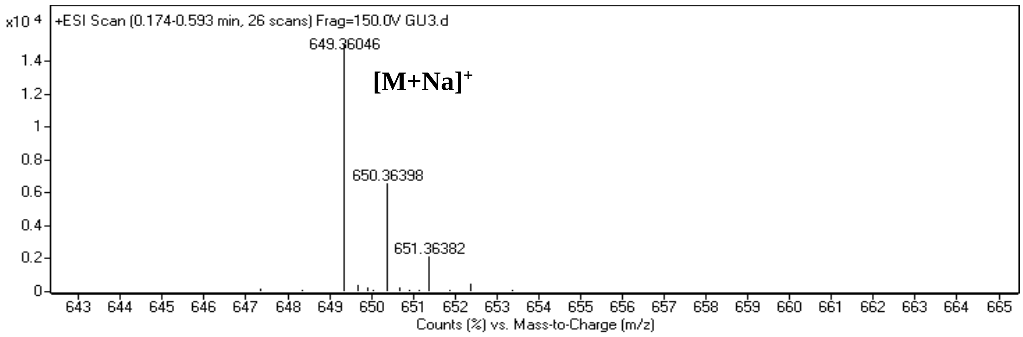

Spectrum 47. HR-ESI-MS Spectrum of CG-03 (positive mode).

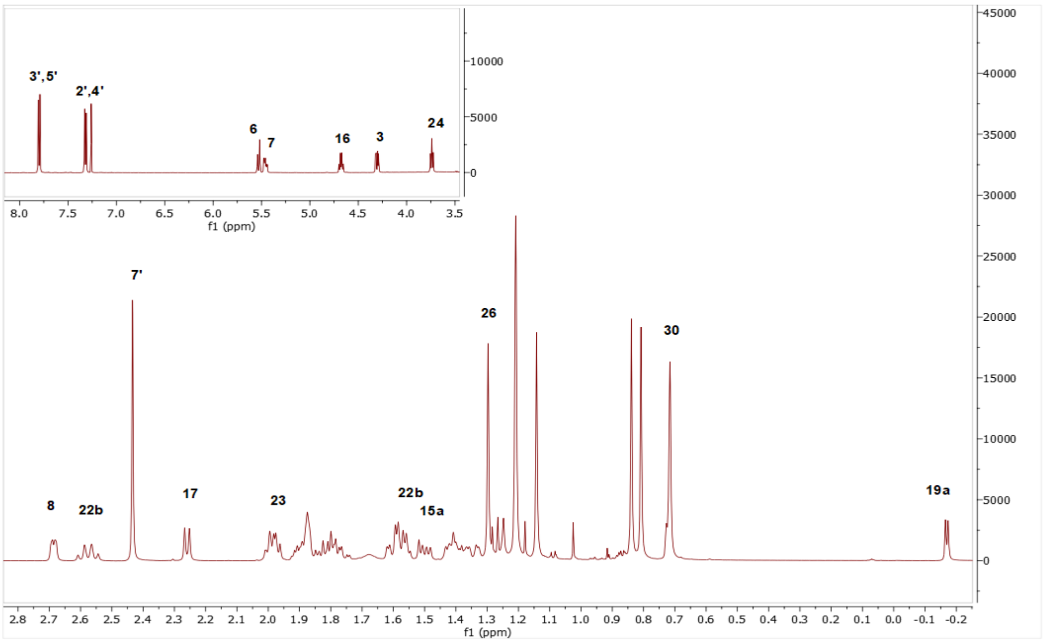

Spectrum 48. $^{1}$H NMR Spectrum of CG-03.

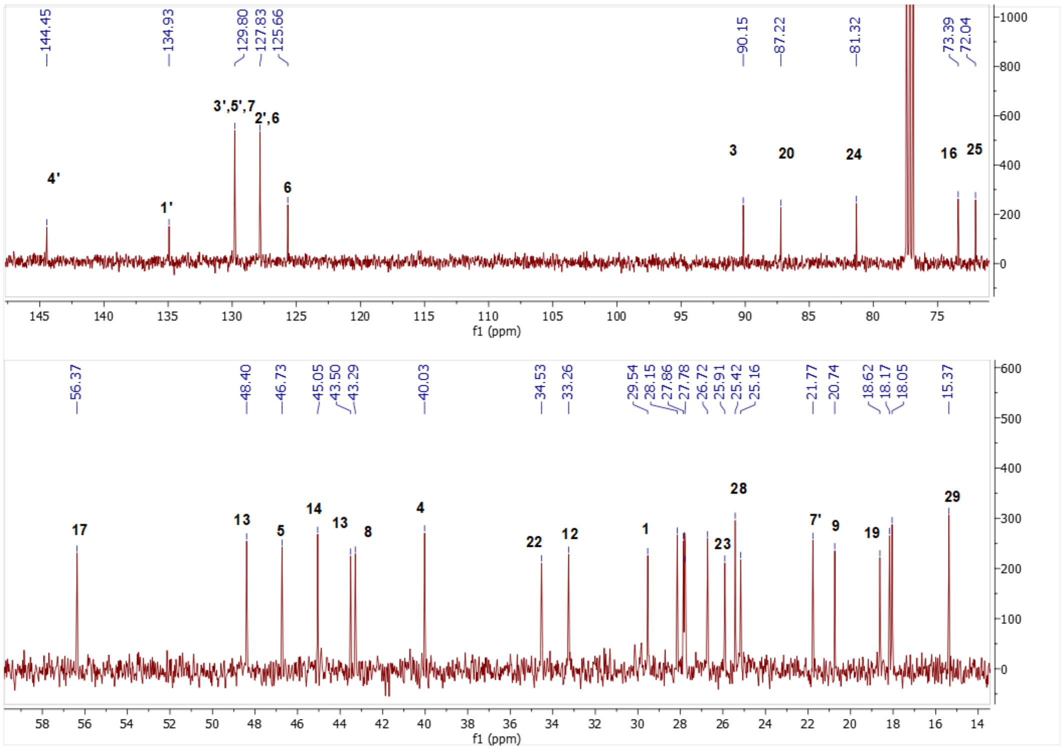

Spectrum 49. $^{13}$C NMR Spectrum of CG-03.

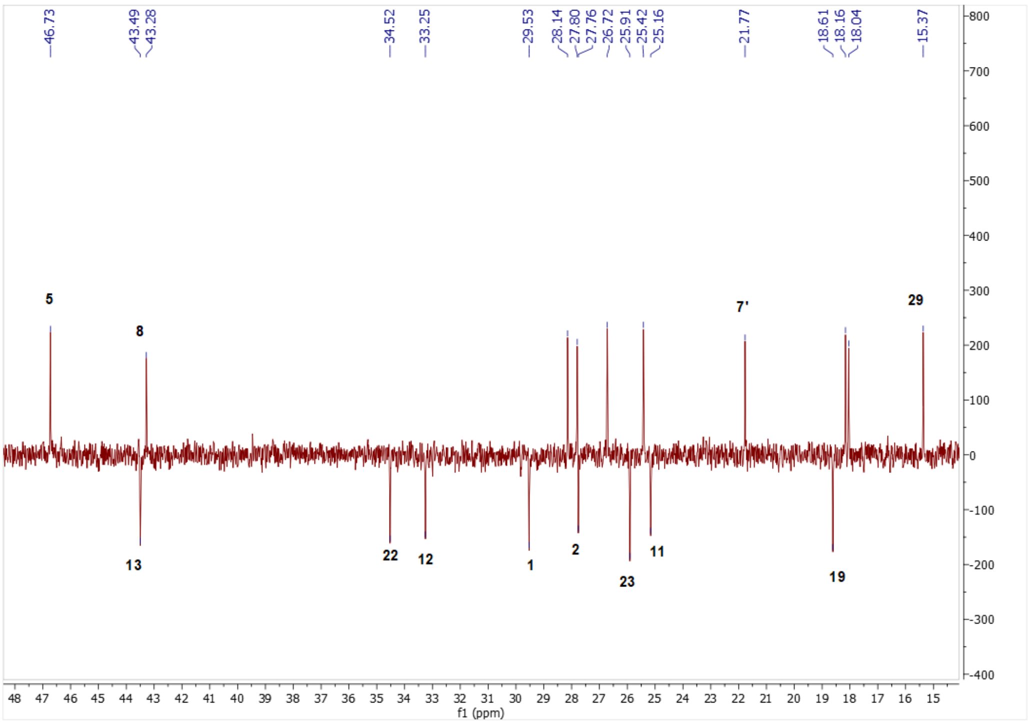

Spectrum 50. DEPT135 spectrum of CG-03.

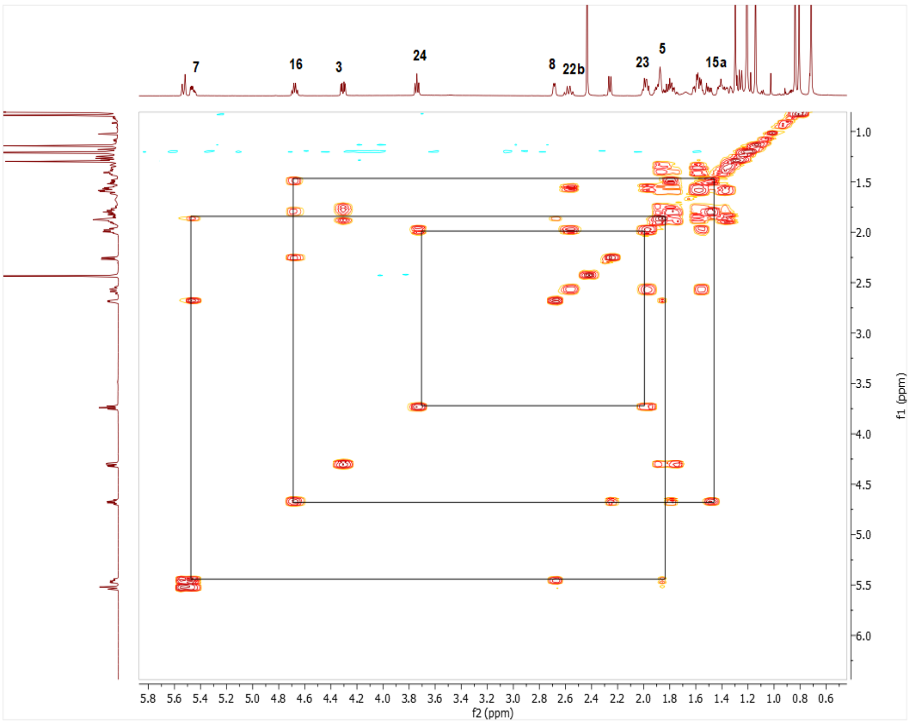

Spectrum 51. COSY spectrum of CG-03.

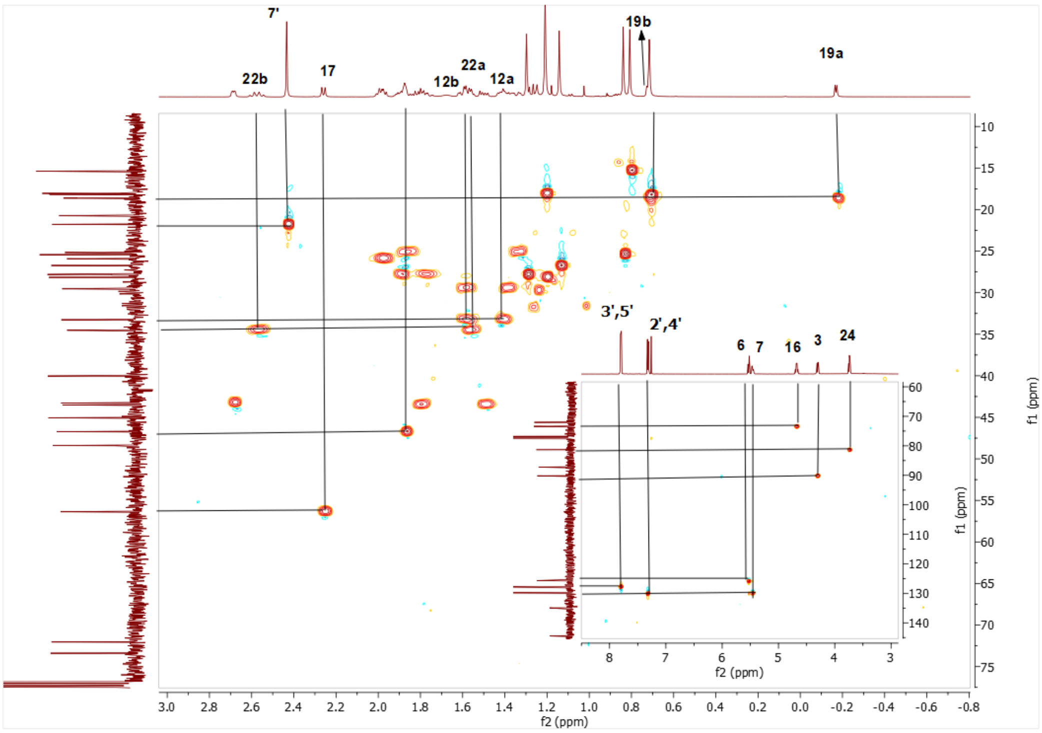

Spectrum 52. HMQC spectrum of CG-03.

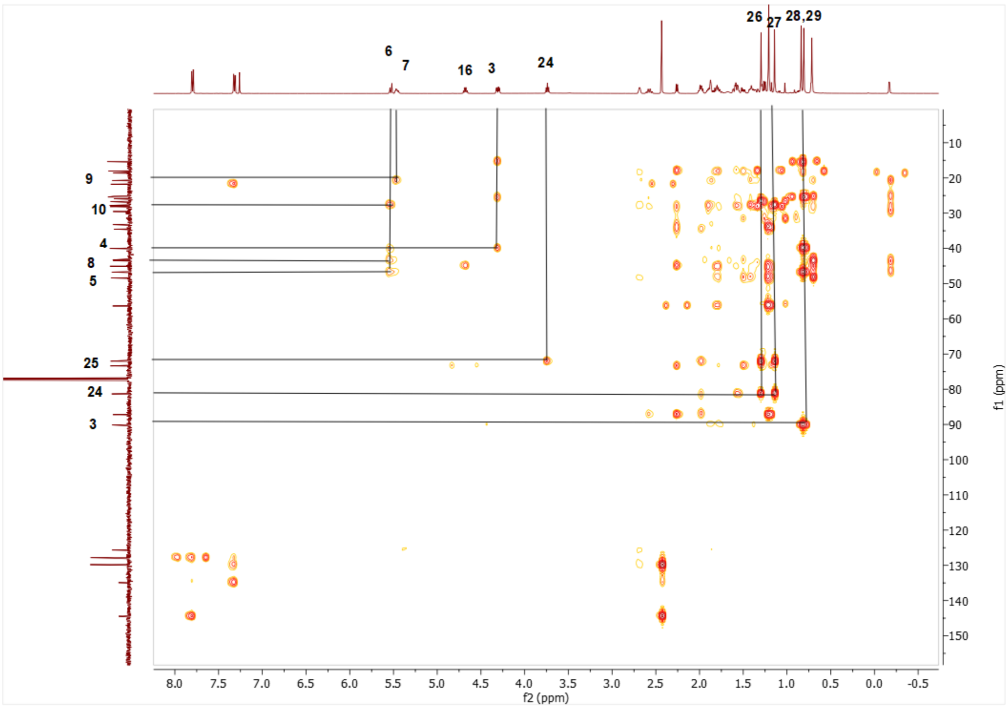

Spectrum 53. HMBC spectrum of CG-03.

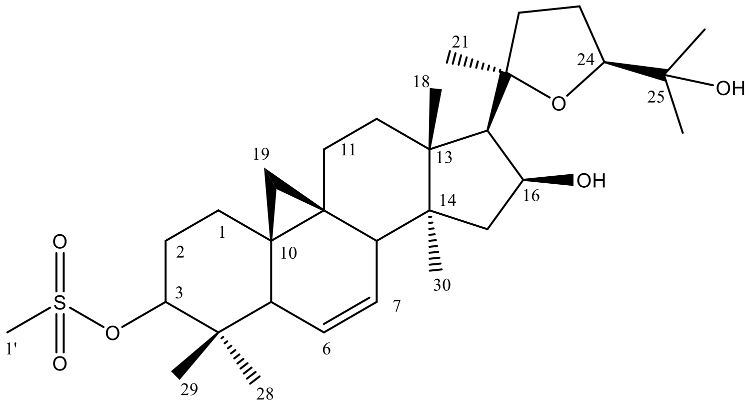

Supplementary Figure 11. Chemical Structure of CG-04

Supplementary Table 9. The $^{13}C$ and $^{1}H$ NMR data of CG-04 (100/400 MHz, δ ppm, in CDCl3).

| **H/C** | **$\delta_C$ (ppm)** | **$\delta_H$ (ppm), *J* (Hz)** |
|---|---|---|
| **1** | 29.5 d | 1.47 m, 1.68 m |
| **2** | 28.2 t | 1.9 m, 2.12 dd (12.5, 3.9) |
| **3** | 89.7 d | 4.43 dd (11.9, 4.6) |
| **4** | 40.1 s | - |
| **5** | 46.7 d | 1.96 m |
| **6** | 125.6 d | 5.58 d (10.6) |
| **7** | 129.9 d | 5.49 ddd (10.6, 6.1, 3.1) |
| **8** | 43.3 d | 2.71 dd (6.2, 2.6) |
| **9** | 20.8 s | - |
| **10** | 27.9 s | - |
| **11** | 25.2 t | 1.38 m, 1.89 m |
| **12** | 33.3 t | 1.43 m, 1.61m |
| **13** | 48.4 s | - |
| **14** | 45.1 s | - |
| **15** | 43.5 t | 1.51m, 1.81m |
| **16** | 73.4 d | 4.7 ddd (7.7, 7.7, 6) |
| **17** | 56.4 d | 2.27 d (7.6) |
| **18** | 18.1 q | 1.22 s |
| **19** | 18.6 t | -0.12 d (4.3), 0.77 d (3.6) |
| **20** | 87.2 s | - |
| **21** | 28.2 q | 1.2 s |
| **22** | 34.5 t | 1.6 m, 2.58 d (10.6) |
| **23** | 25.8 t | 2.0 m |
| **24** | 81.3 d | 3.73 t (7.1) |
| **25** | 72.0 s | - |
| **26** | 27.8 q | 1.29 s |
| **27** | 26.7 q | 1.14 s |
| **28** | 25.7 q | 1.05 s |
| **29** | 15.4 q | 0.86 s |
| **30** | 18.2 q | 0.74 s |
| **1’** | 38.9 q | 3.01 s |

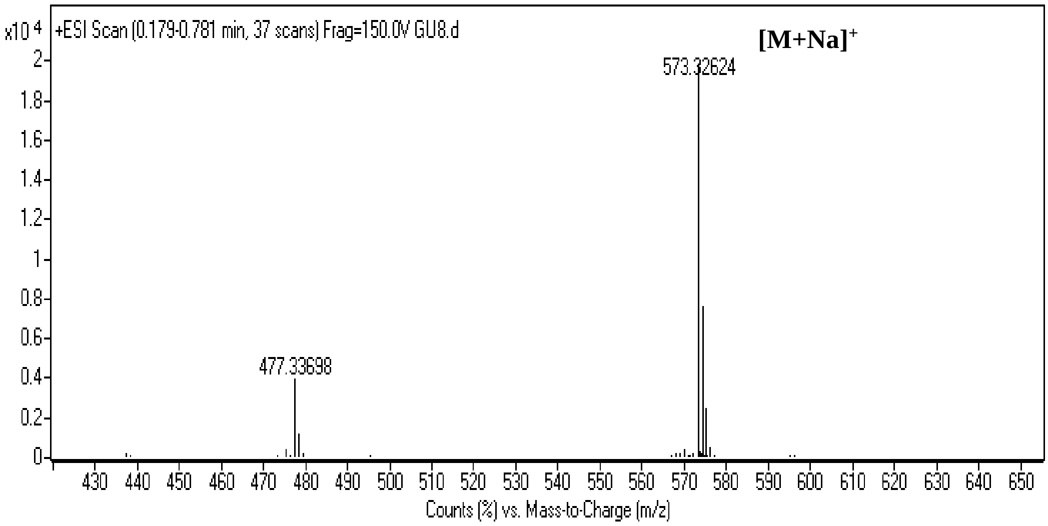

Spectrum 54. HR-ESI-MS Spectrum of CG-04 (positive mode).

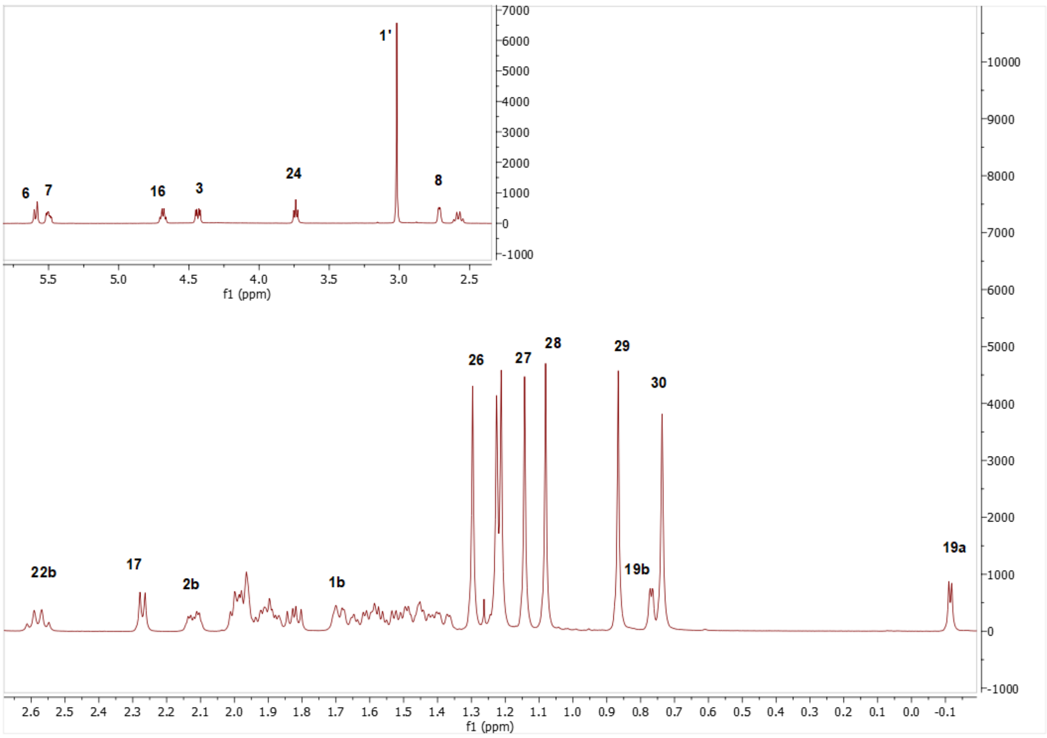

Spectrum 55. $^{1}$H NMR Spectrum of CG-04.

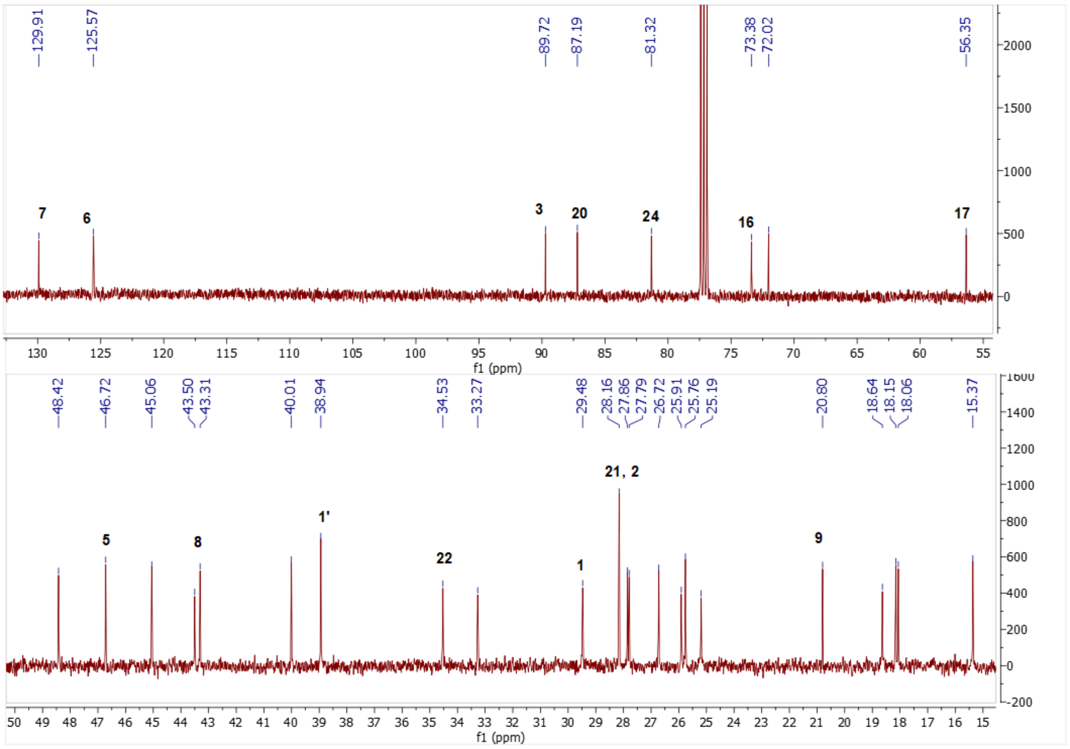

Spectrum 56. $^{13}$C NMR Spectrum of CG-04.

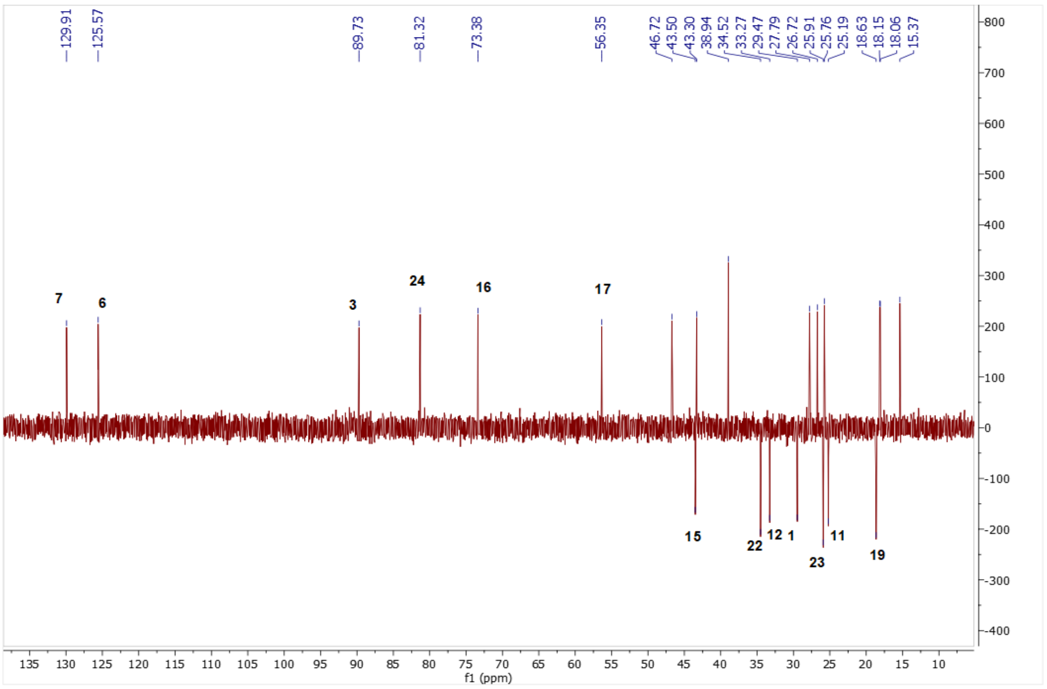

Spectrum 57. DEPT135 spectrum of CG-04.

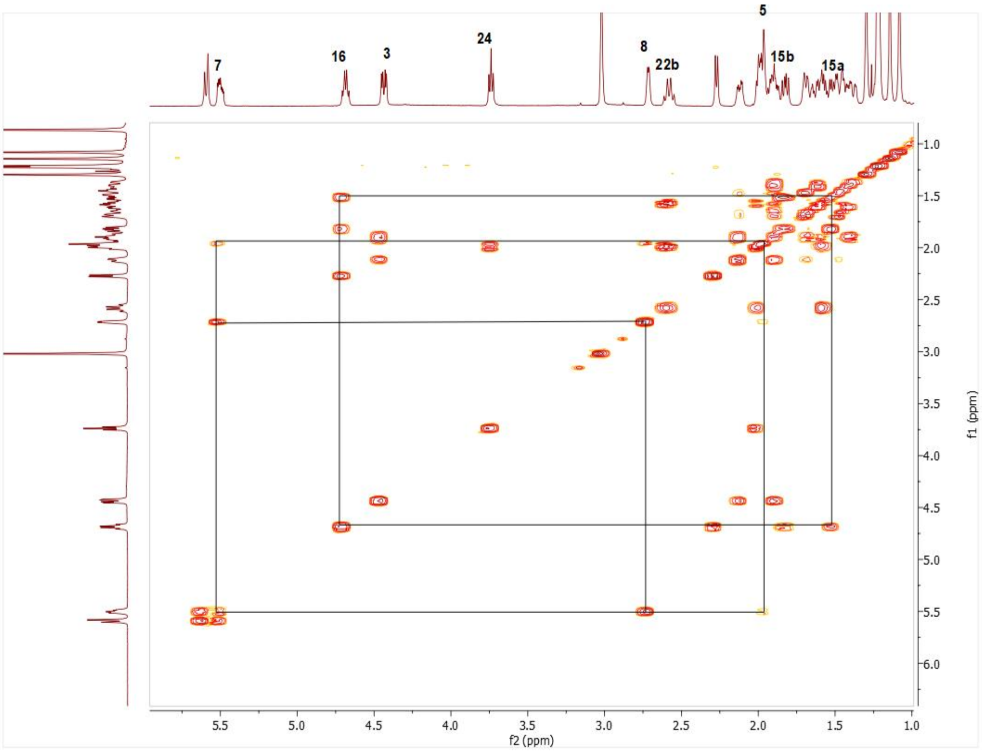

Spectrum 58. COSY spectrum of CG-04.

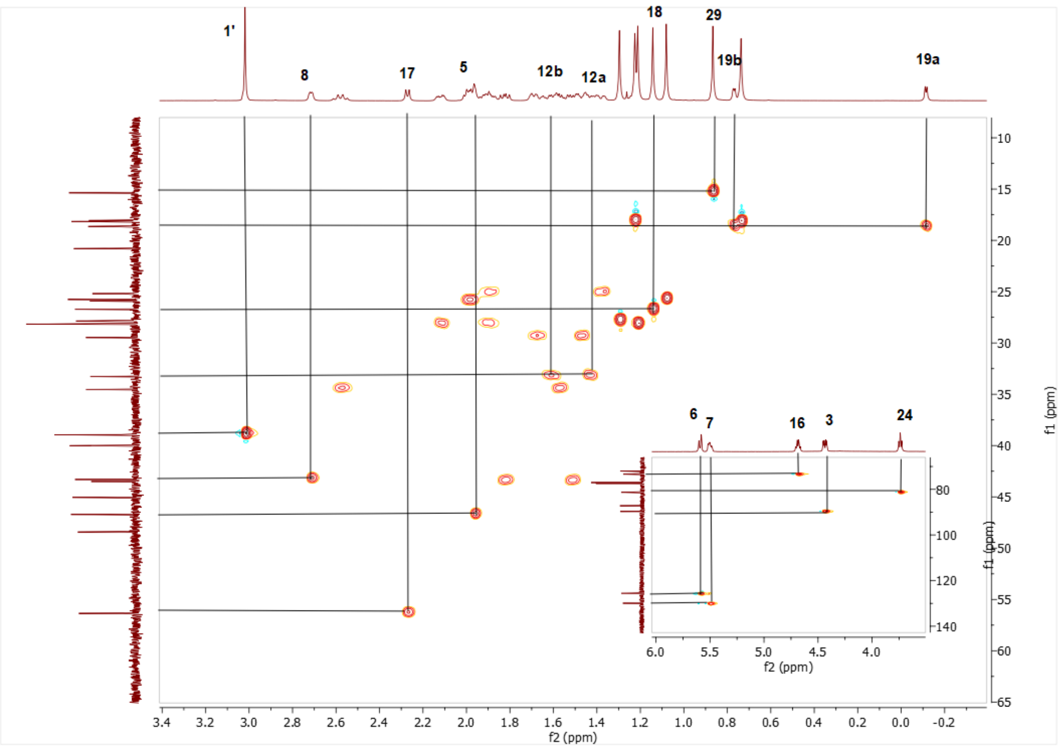

Spectrum 59. HMQC spectrum of CG-04.

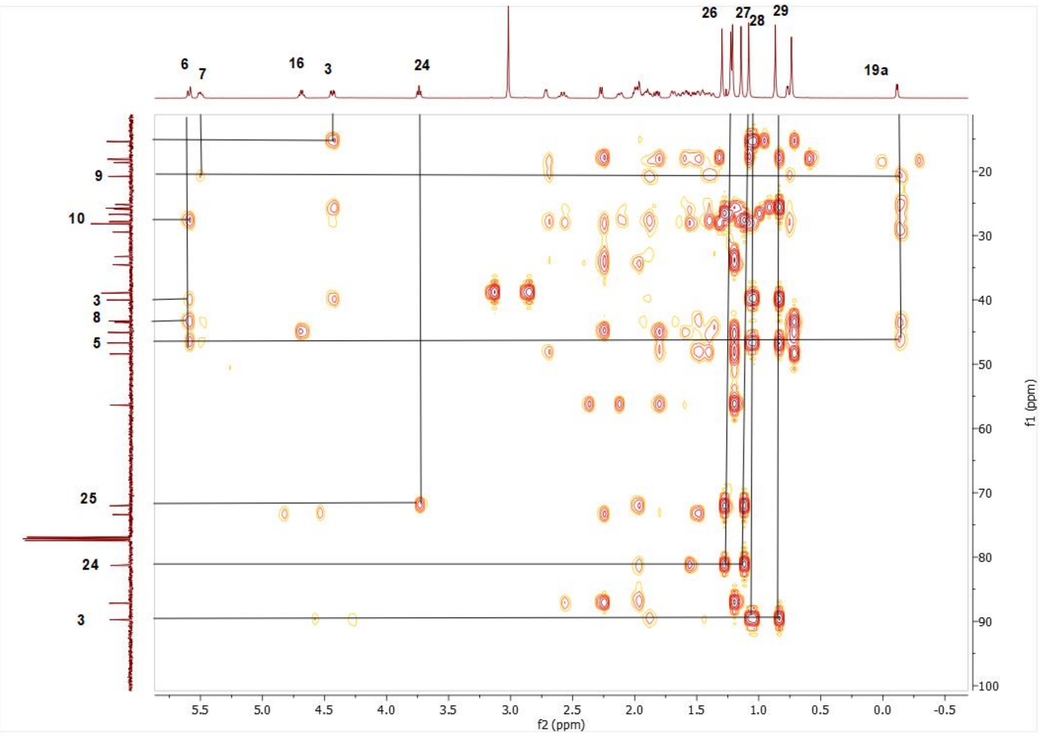

Spectrum 60. HMBC spectrum of CG-04.

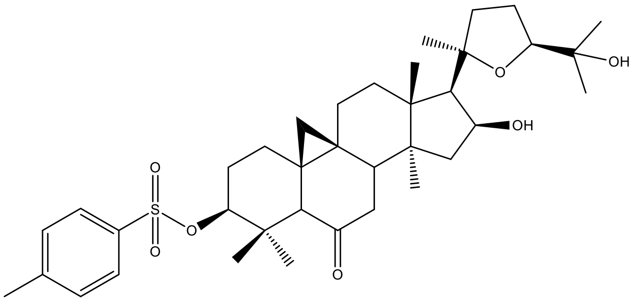

Supplementary Figure 12. Chemical Structure of CG-05

Supplementary Table 10. The $^{13}C$ and $^{1}H$ NMR data of CG-05 (100/400 MHz, δ ppm, in CDCl3).

| H/C | $\delta_C$ (ppm) | $\delta_H$ (ppm), *J* (Hz) |
|---|---|---|
| **1** | 30.0 t | 1.42 m, 1.76 m |
| **2** | 27.3 t | 1.79 m, 1.94 m |
| **3** | 89.3 | 4.21 m |
| **4** | 40.0 s | - |
| **5** | 57.2 d | 2.3 brs |
| **6** | 210 s | - |
| **7** | 41.2 t | 2.12 m, 2.17 m |

| | | |
|---|---|---|
| **8** | 42.4 d | 2.66 dd (8.5, 4) |
| **9** | 21.7 s | - |
| **10** | 29.7 s | - |
| **11** | 26.5 t | 1.46 m, 1.85 m |
| **12** | 33.0 | 1.47 m, 1.6 m |
| **13** | 47.1 s | - |
| **14** | 45.3 s | - |
| **15** | 43.8 t | 1.37 m, 1.84 m |
| **16** | 72.9 d | 4.69 ddd (7.8, 7.8, 6.1) |
| **17** | 56.9 d | 2.32 d (7.6) |
| **18** | 18.4 q | 1.21 m |
| **19** | 22.2 t | 0.21, 0.6 d (5.5) |
| **20** | 87.1 s | - |
| **21** | 28.1 | 1.21 s |
| **22** | 34.5 t | 1.57 m, 2.57 q (10.8) |
| **23** | 25.91 t | 1.98 m |
| **24** | 81.26 d | 3.75 dd (8.3, 6.1) |
| **25** | 72.1 s | - |
| **26** | 26.8 q | 1.14 s |
| **27** | 27.8 q | 1.3 s |
| **28** | 26.3 q | 1.02 s |
| **29** | 14.8 q | 1.0 s |
| **30** | 19.1 q | 0.89 s |
| **1’** | 134.7 s | - |
| **2’** | 127.8 d | 7.79 d (8.3) |
| **3’** | 129.8 d | 7.3 d (8.1) |
| **4’** | 144.6 s | - |
| **5’** | 129.8 d | 7.3 d (8.1) |
| **6’** | 127.8 d | 7.79 d (8.3) |
| **7’** | 21.8 q | 2.43 s |

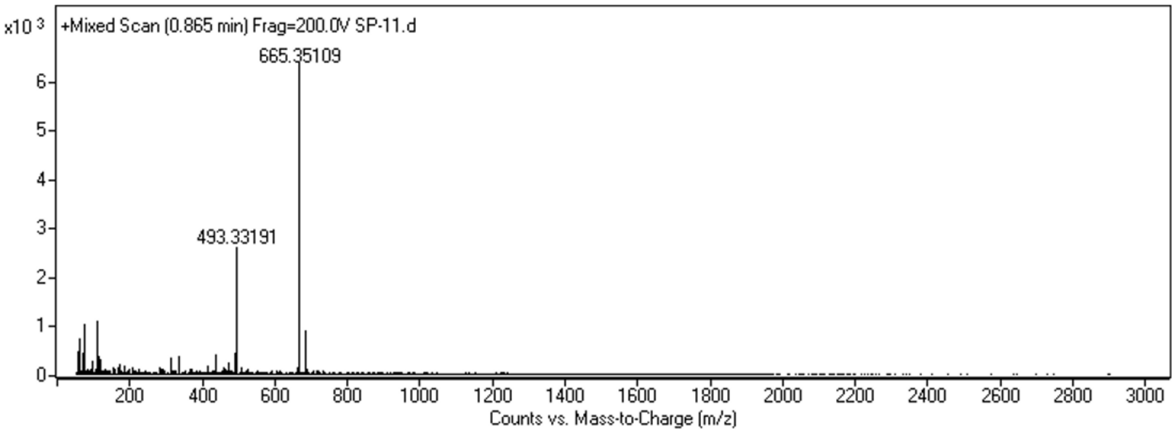

Spectrum 61. HR-ESI-MS Spectrum of CG-05 (positive mode).

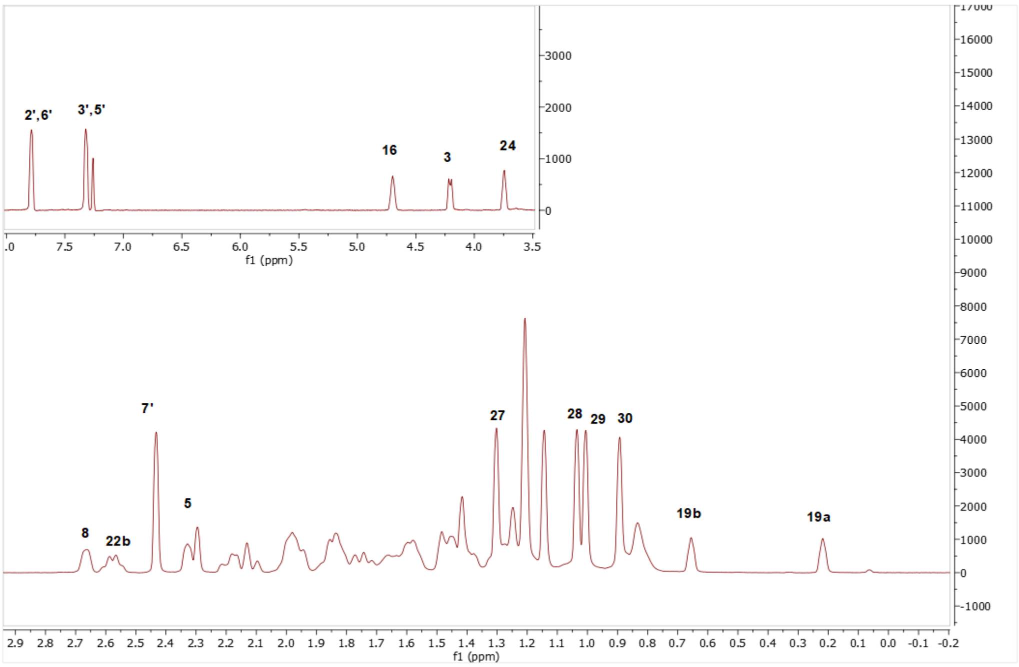

Spectrum 62. $^{1}H$ NMR Spectrum of CG-05.

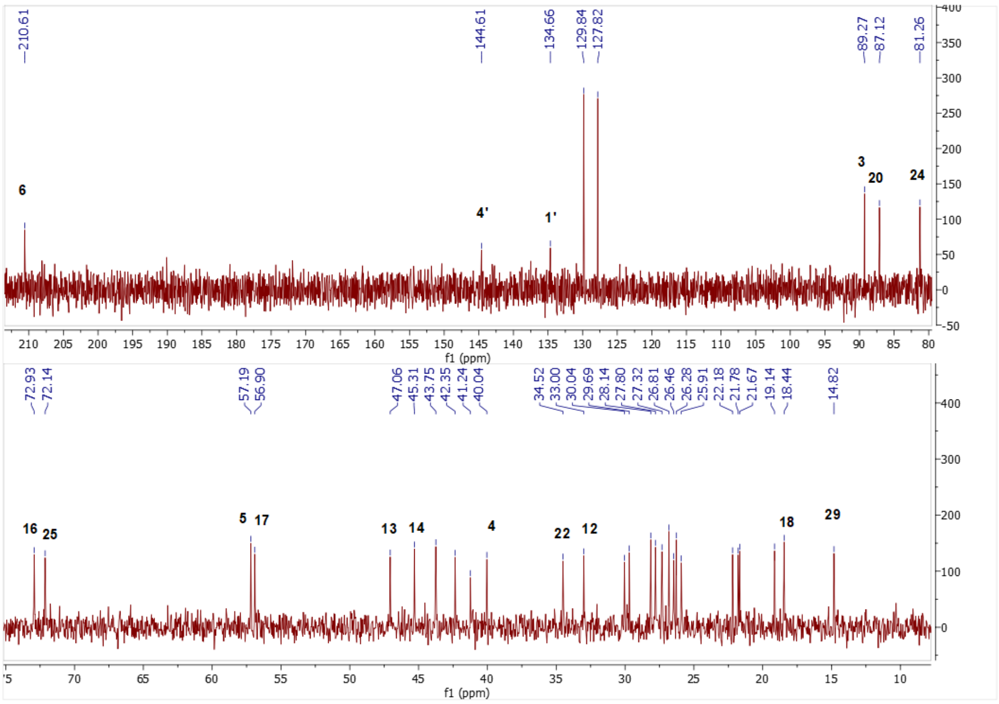

Spectrum 63. $^{13}C$ NMR Spectrum of CG-05.

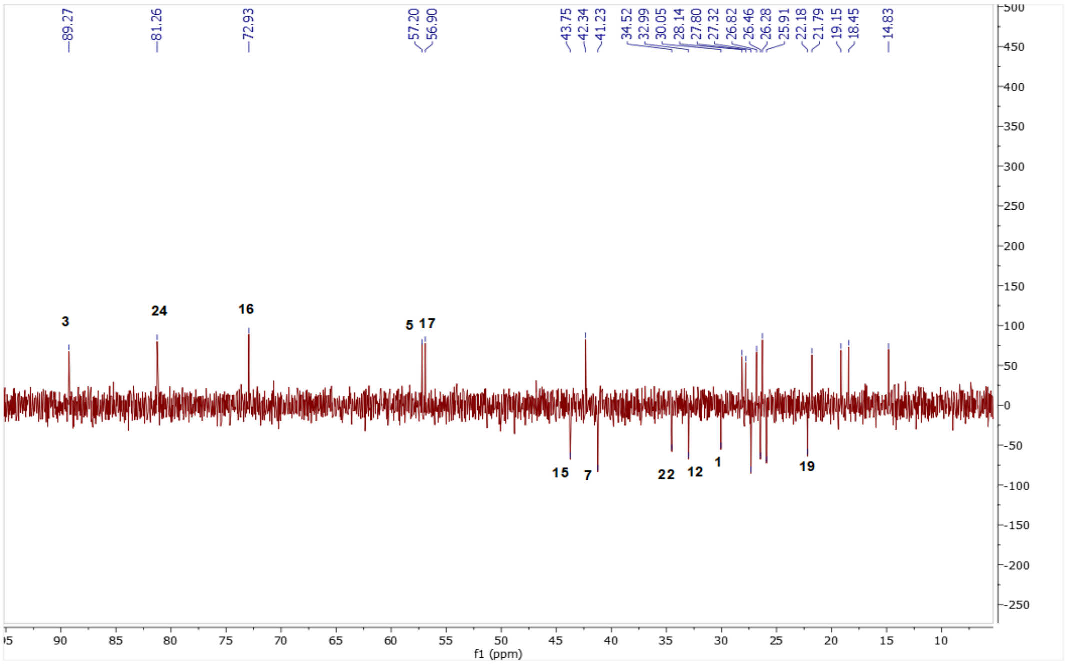

Spectrum 64. DEPT135 spectrum of CG-05.

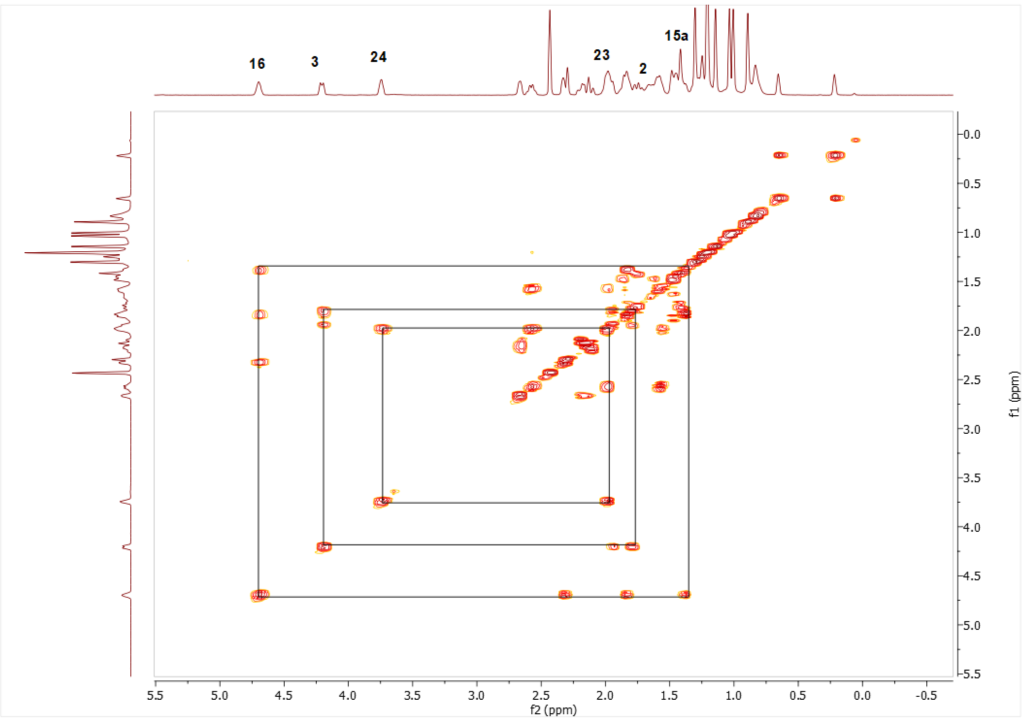

Spectrum 65. COSY spectrum of CG-05.

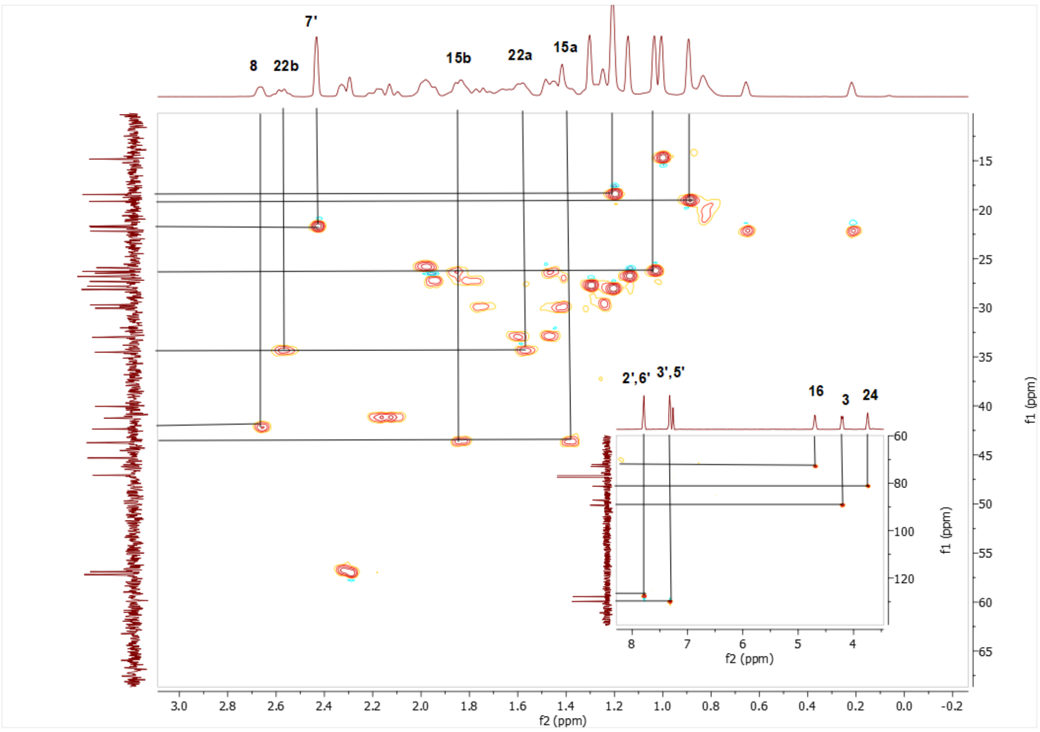

Spectrum 66. HMQC spectrum of CG-05.

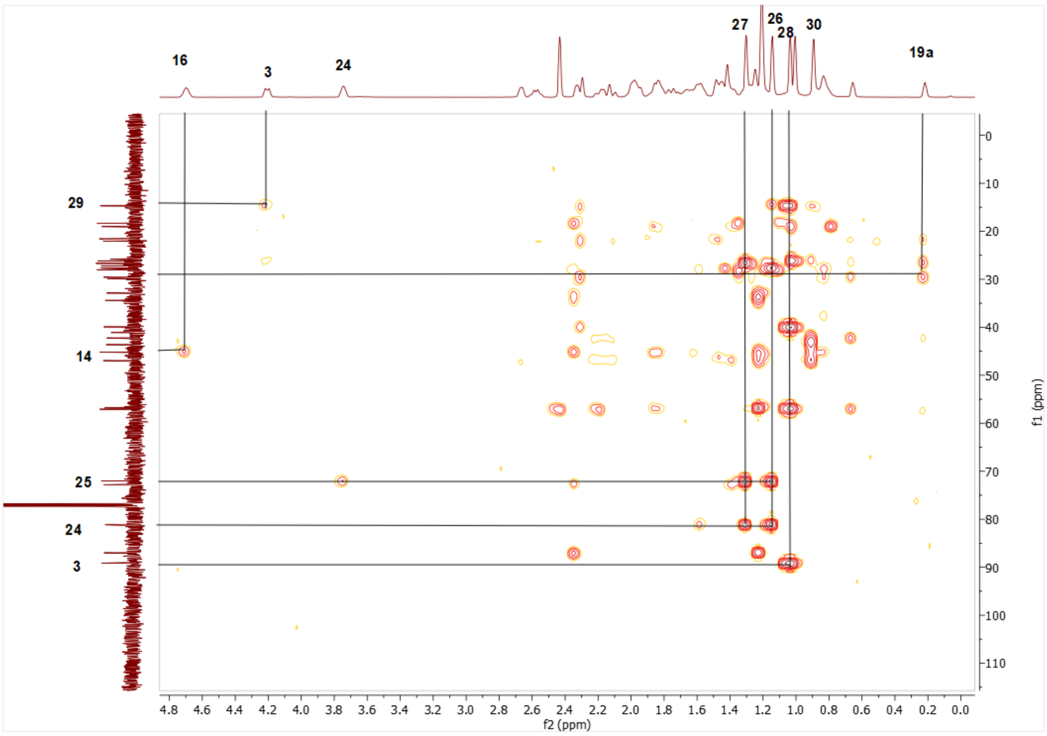

Spectrum 67. HMBC spectrum of CG-05.

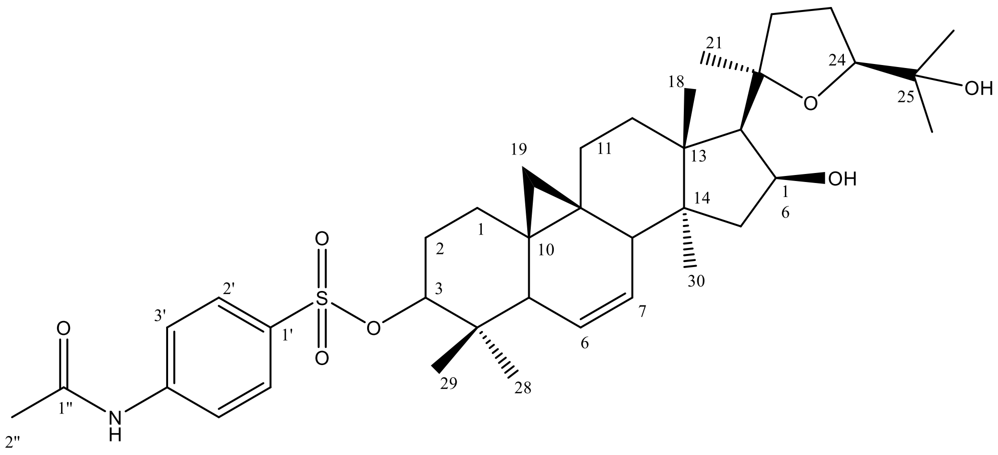

Supplementary Figure 13. Chemical Structure of CG-06

Supplementary Table 11. The $^{13}C$ and $^{1}H$ NMR data of CG-06 (100/400 MHz, δ ppm, in CDCl3).

| H/C | $\delta_C$ (ppm) | $\delta_H$ (ppm), *J* (Hz) |
|---|---|---|
| **1** | 29.5 t | 1.37 m, 1.56 m |
| **2** | 27.7 t | 1.77 m, 1.87 m |
| **3** | 90.4 d | 4.26 dd (11.8, 4.7) |
| **4** | 40.0 s | - |
| **5** | 46.6 d | 1.85 m |
| **6** | 125.6 d | 5.5 d (10.7) |
| **7** | 129.7 d | 5.43 ddd (0.2, 6.2, 2.9) |
| **8** | 43.2 d | 2.66 m |
| **9** | 20.7 s | - |
| **10** | 27.8 s | - |
| **11** | 25.1 t | 1.33 m, 1.85 m |
| **12** | 33.2 t | 1.40 m, 1.57 m |
| **13** | 48.3 s | - |
| **14** | 45.0 s | - |
| **15** | 43.4 t | 1.48 m, 1.78 m |
| **16** | 73.4 d | 4.67 q (4.2) |
| **17** | 56.3 d | 2.26 d (7.5) |
| **18** | 18.1 q | 1.18 s |
| **19** | 18.6 t | -0.18, 0.69 d (4.1) |
| **20** | 87.1 s | - |
| **21** | 28.2 q | 1.18 s |
| **22** | 34.5 t | 1.56 m, 2.57 q (10.8) |
| **23** | 25.9 t | 1.98 m |
| **24** | 81.3 d | 3.73 t (6.9) |
| **25** | 72.0 s | - |
| **26** | 27.7 q | 1.27 s |
| **27** | 26.7 q | 1.13 s |
| **28** | 25.4 q | 0.8 s |
| **29** | 15.3 q | 0.78 s |
| **30** | 18.1 q | 0.69 s |
| **1’** | 131.8 s | - |

| | | |
|---|---|---|
| **2’** | 129.0 d | 7.82 d (8.5) |
| **3’** | 119.3 d | 7.3 d (8.5) |
| **4’** | 143.2 s | - |
| **5’** | 119.3 d | 7.3 d (8.5) |
| **6’** | 129.0 d | 7.82 d (8.5) |
| **1’’** | 169.3 | - |
| **2’’** | 24.7 s | 2.17 s |

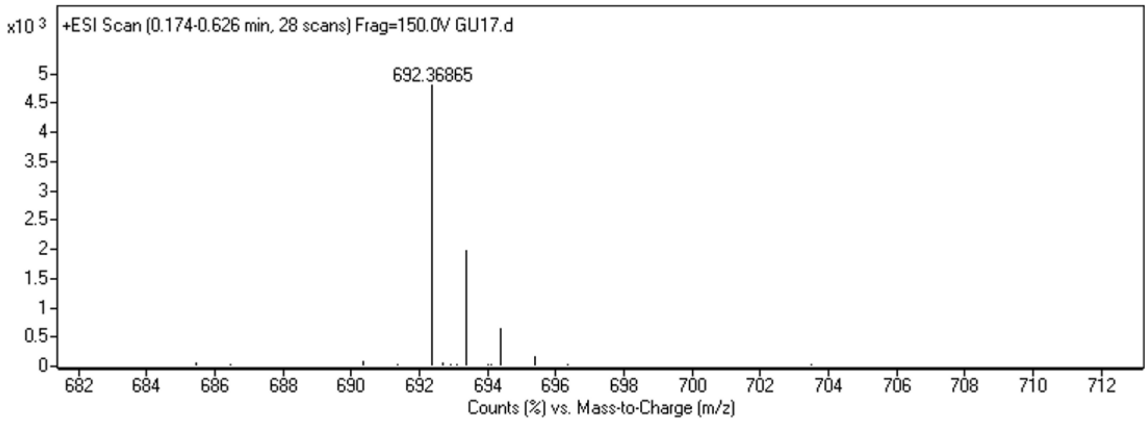

Spectrum 68. HR-ESI-MS Spectrum of CG-0*6* (positive mode).

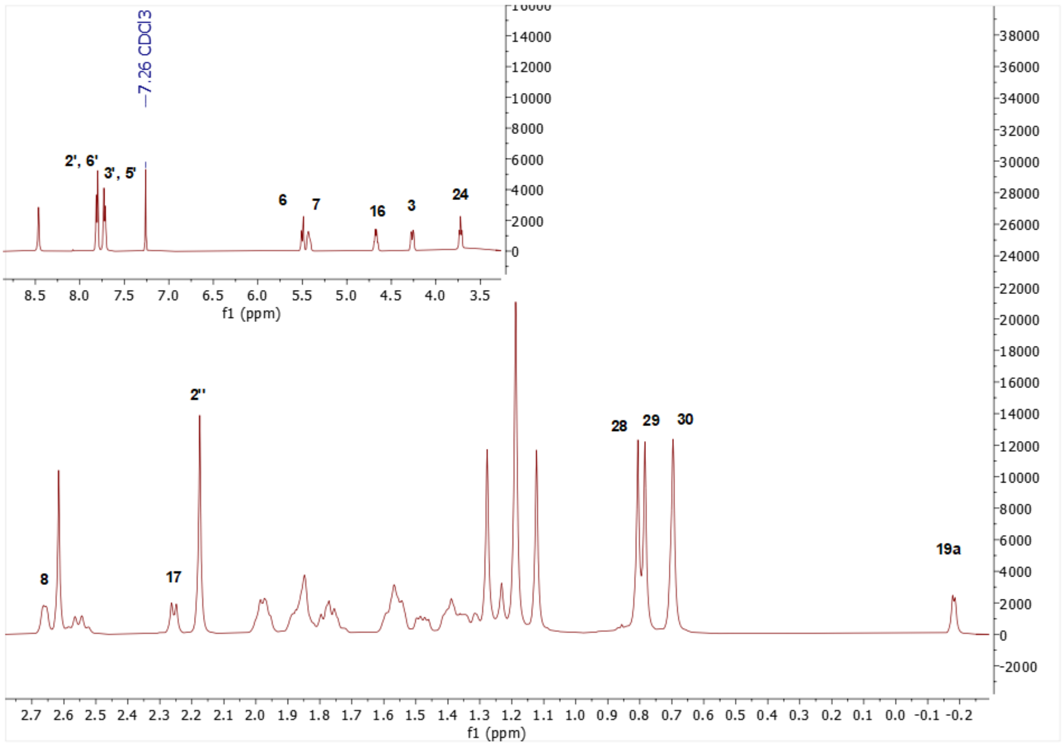

Spectrum 69. $^{1}$H NMR Spectrum of CG-06.

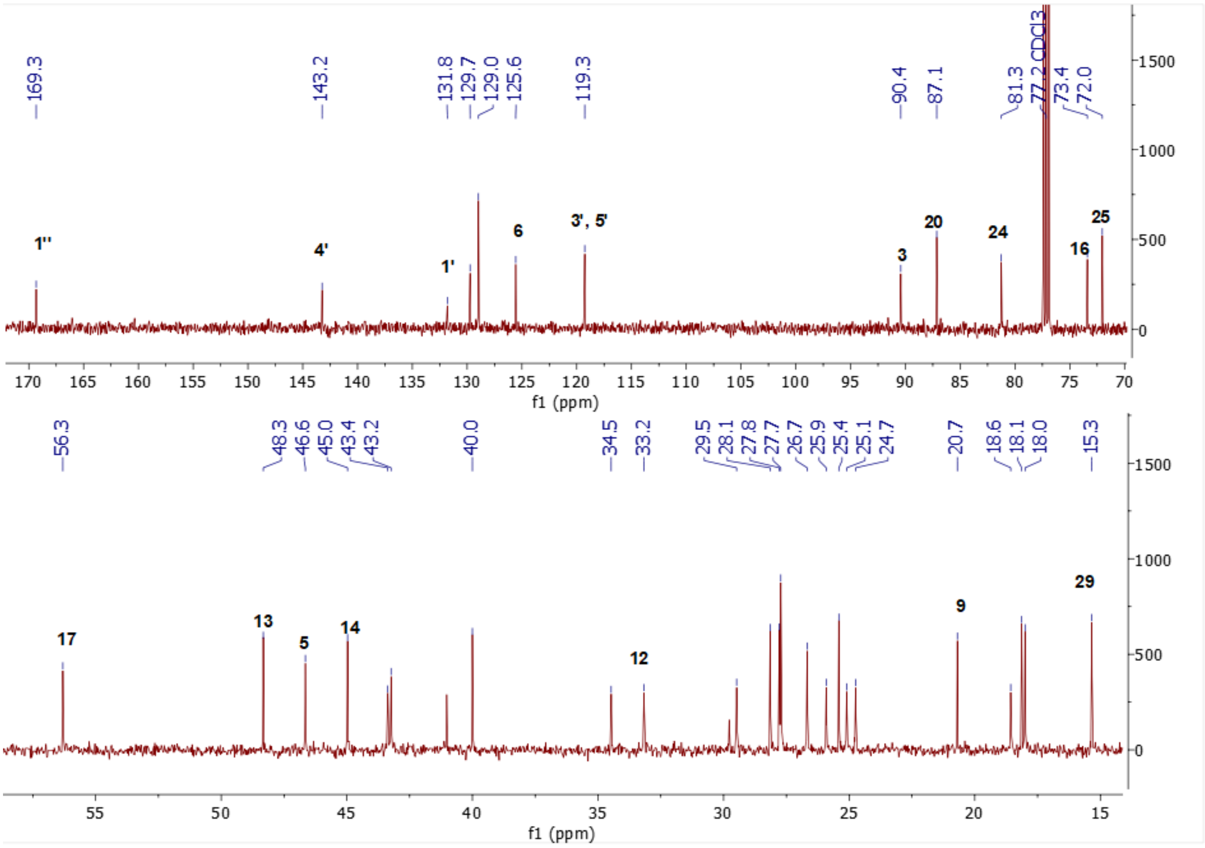

Spectrum 70. $^{13}$C NMR Spectrum of CG-06.

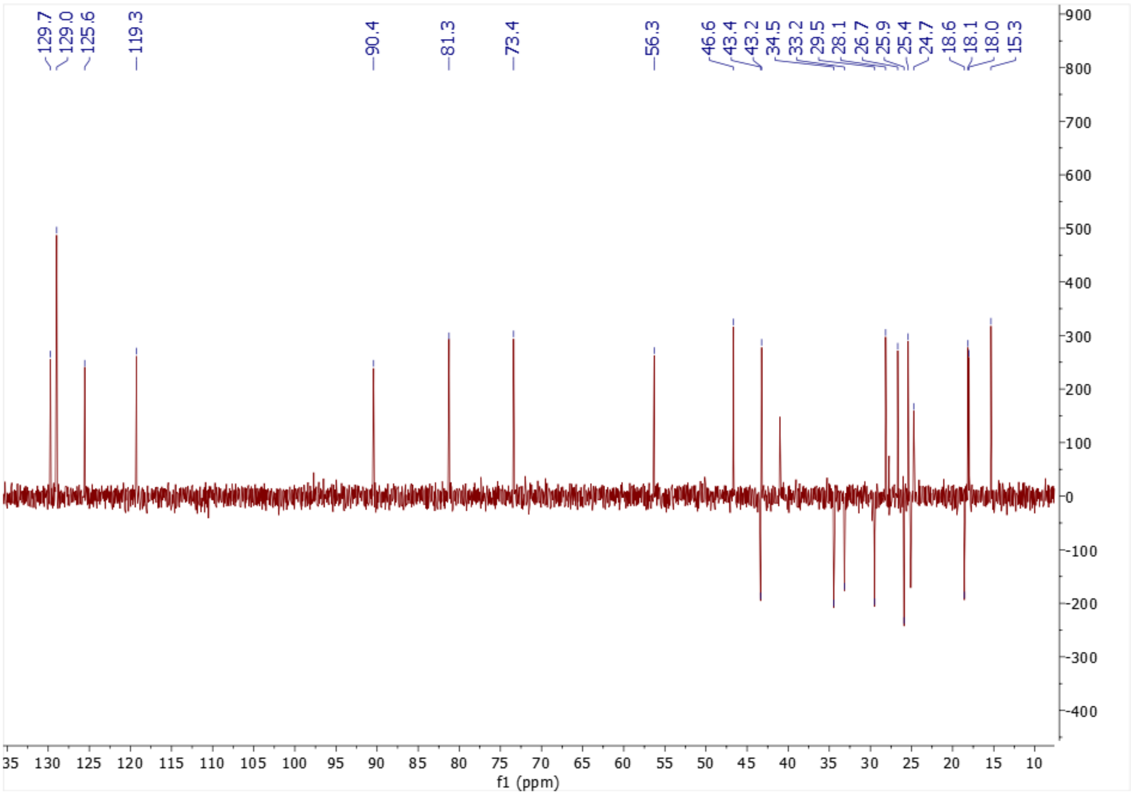

Spectrum 71. DEPT135 spectrum of CG-06

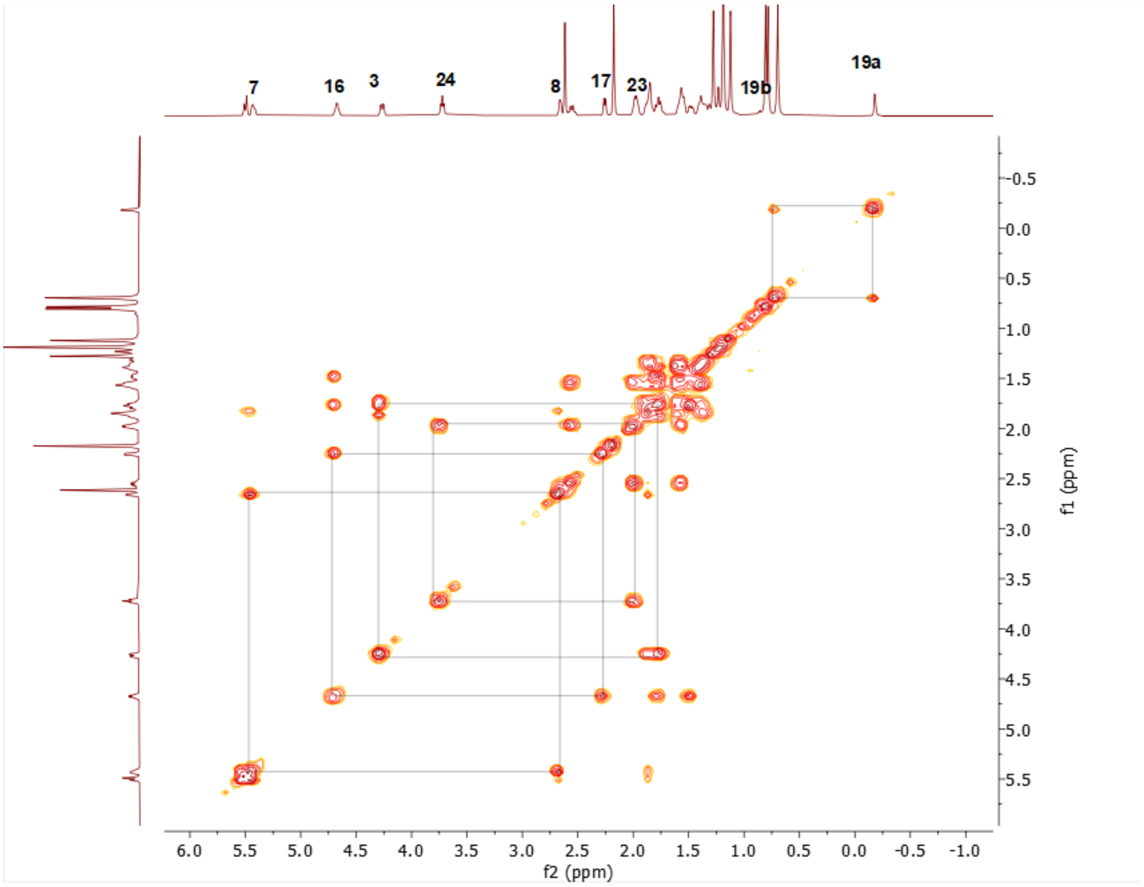

Spectrum 72. COSY spectrum of CG-06

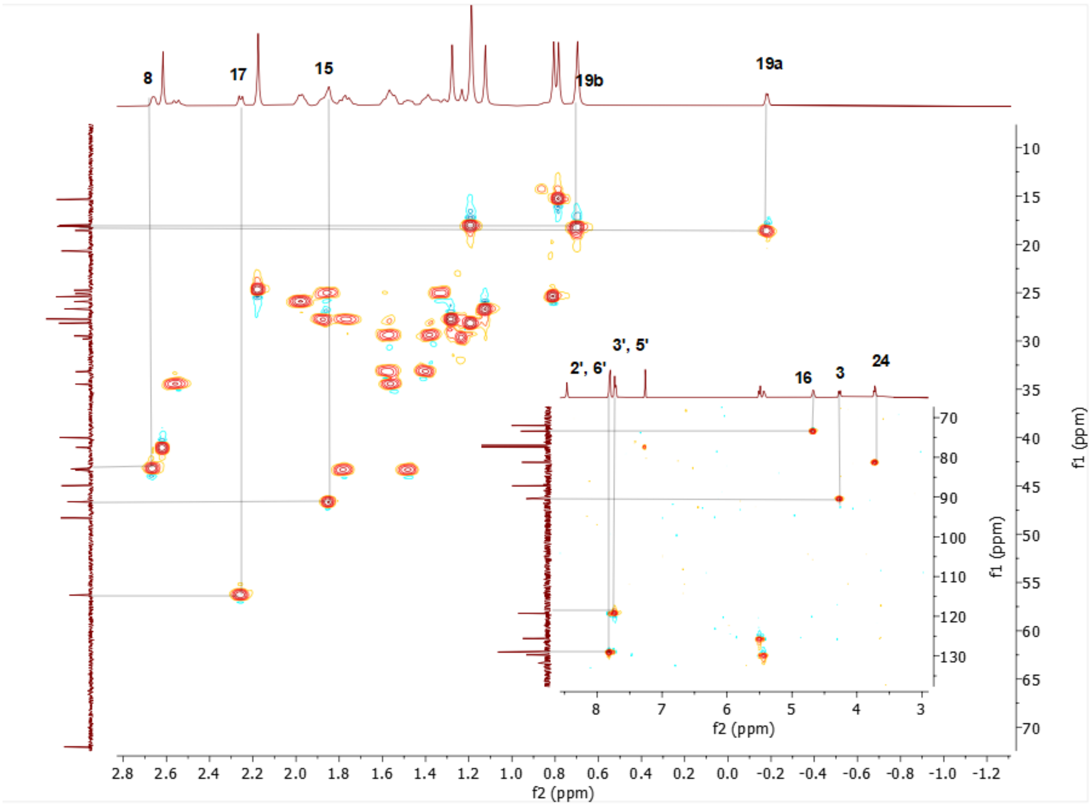

Spectrum 73. HMQC spectrum of CG-06.

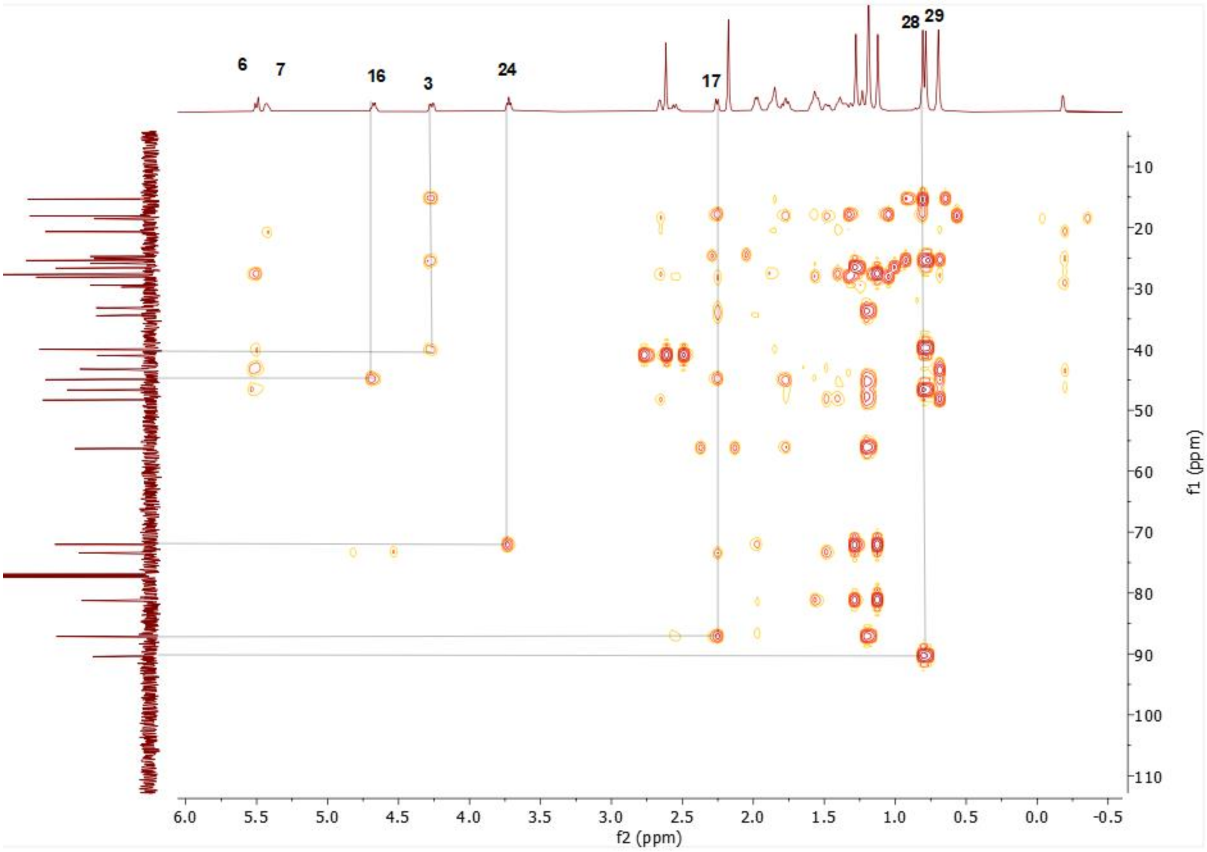

Spectrum 74. HMBC spectrum of CG-05.

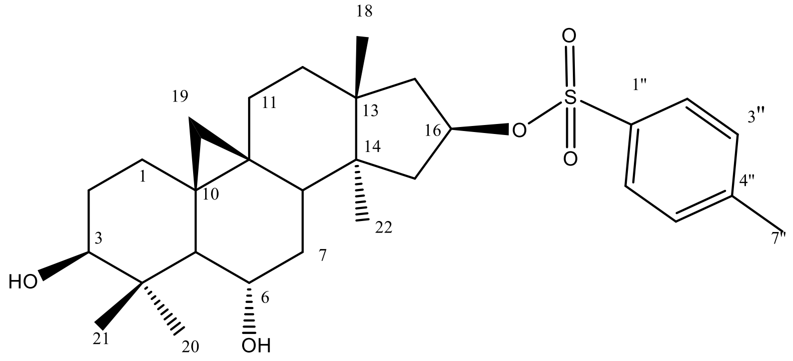

Supplementary Figure 14. Chemical Structure of SCG-01

Supplementary Table 12. The $^{13}$C and $^{1}$H NMR data of SCG-01 (100/400 MHz, δ ppm, in CDCl3).

| H/C | $\delta_C$ (ppm) | $\delta_H$ (ppm), *J* (Hz) |
|---|---|---|
| **1** | 32.0 t | 1.2 d (3.2), 1.57 m |
| **2** | 30.3 t | 1.56 m, 1.78 m |

| | | |
|---|---|---|
| **3** | 78.4 d | 3.29 dd (11.3, 4.6) |
| **4** | 41.6 s | - |
| **5** | 53.5 d | 1.34 d (1.97) |
| **6** | 68.5 d | 3.51 ddd (9.1, 9.1, 4.2) |
| **7** | 37.5 t | 1.3 m, 1.44 m |
| **8** | 45.8 d | 1.6 m |
| **9** | 20.8 s | - |
| **10** | 29.8 s | - |
| **11** | 26.1 t | 1.25 m, 1.92 m |
| **12** | 30.3 t | 1.4 m, 1.6 m |
| **13** | 44.5 s | - |
| **14** | 45.9 s | - |
| **15** | 46.0 t | 1.88 m |
| **16** | 82.9 d | 5.06 ddd (15.4, 7.9, 1.4) |
| **17** | 44.4 t | 1.86 m, 1.64 d (1.43) |
| **18** | 25.0 q | 0.95 s |
| **19** | 30.0 t | 0.28 d (4.6), 0.45 d (4.6) |
| **28** | 28.0 q | 1.22 s |
| **29** | 15.3 q | 0.92 s |
| **30** | 19.8 q | 1.04 s |
| **1’** | 134.4 s | - |
| **2’** | 127.8 d | 7.75 d (8.2) |
| **3’** | 129.9 d | 7.31 d (8.2) |
| **4’** | 144.6 s | - |
| **5’** | 129.9 d | 7.31 d (8.2) |
| **6’** | 127.8 d | 7.75 d (8.2) |
| **7’** | 21.8 q | 2.44 s |

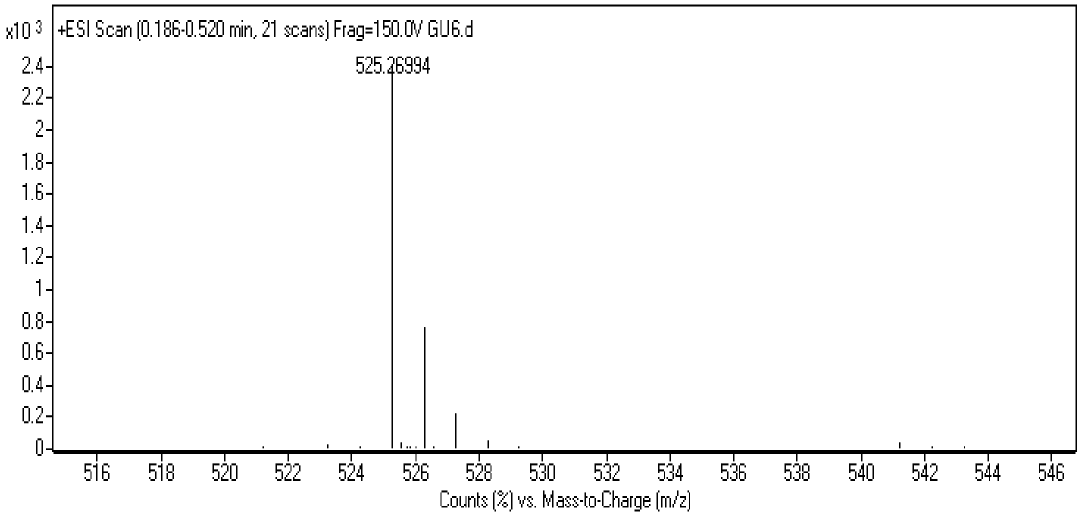

Spectrum 75.HR-ESI-MS Spectrum of SCG-01 (positive mode).

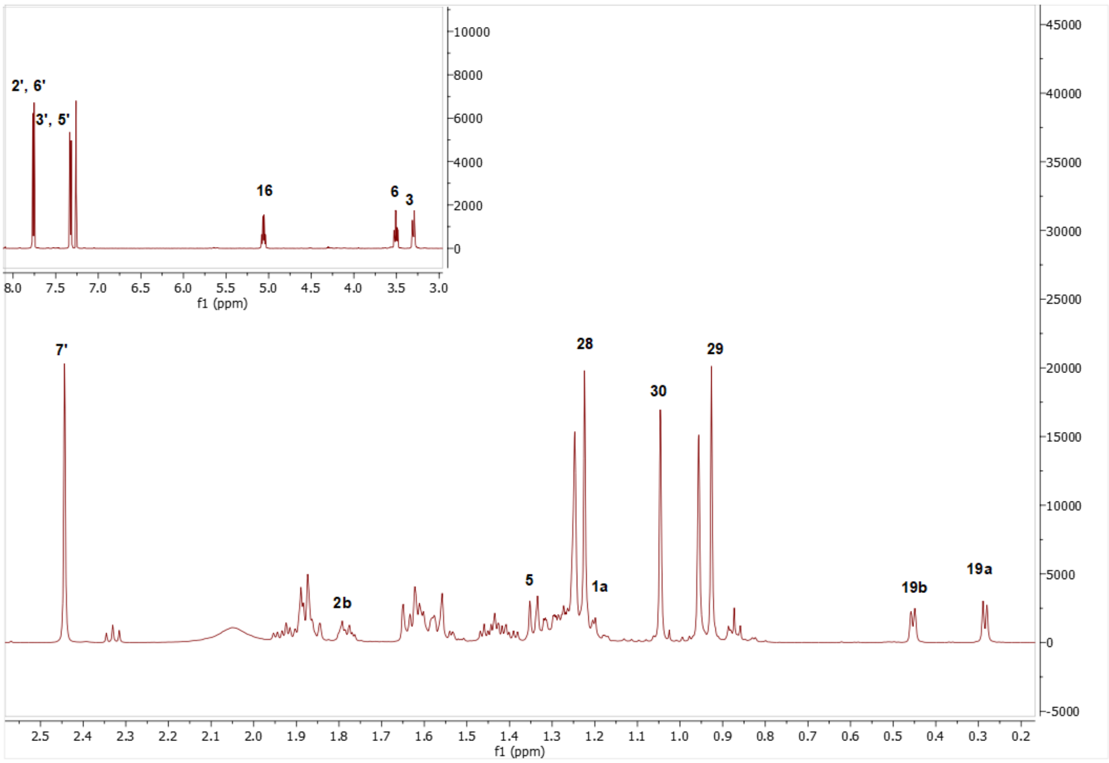

Spectrum 761. $^{1}$H NMR Spectrum of SCG-01

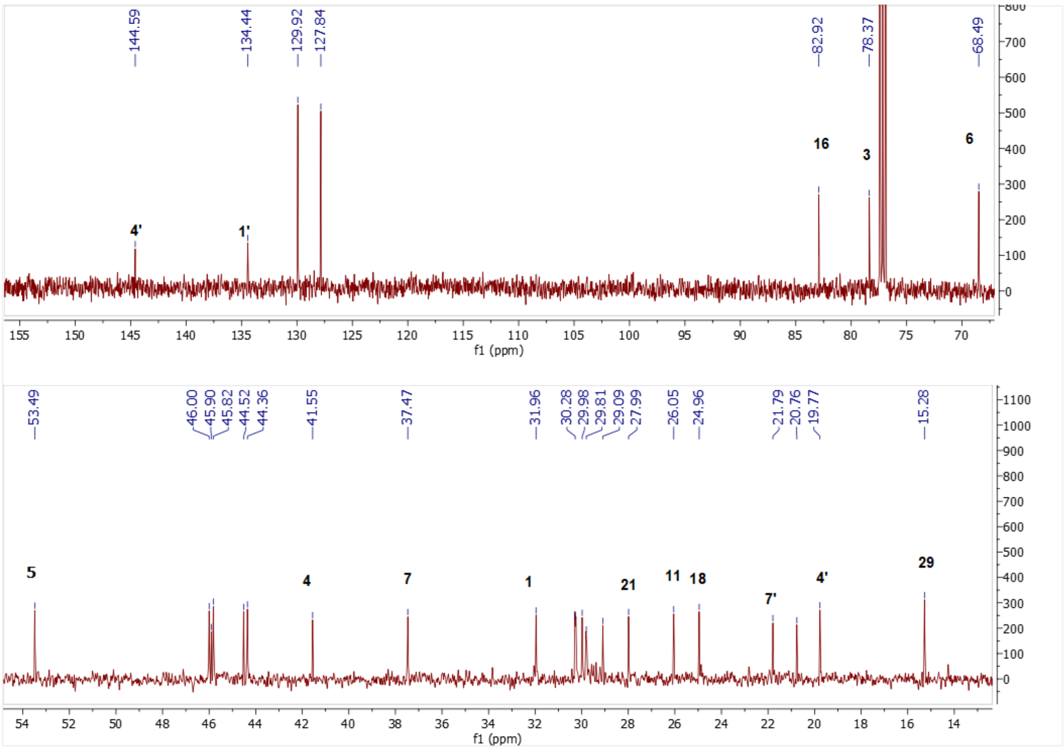

Spectrum 77. $^{13}$C NMR Spectrum of SCG-01.

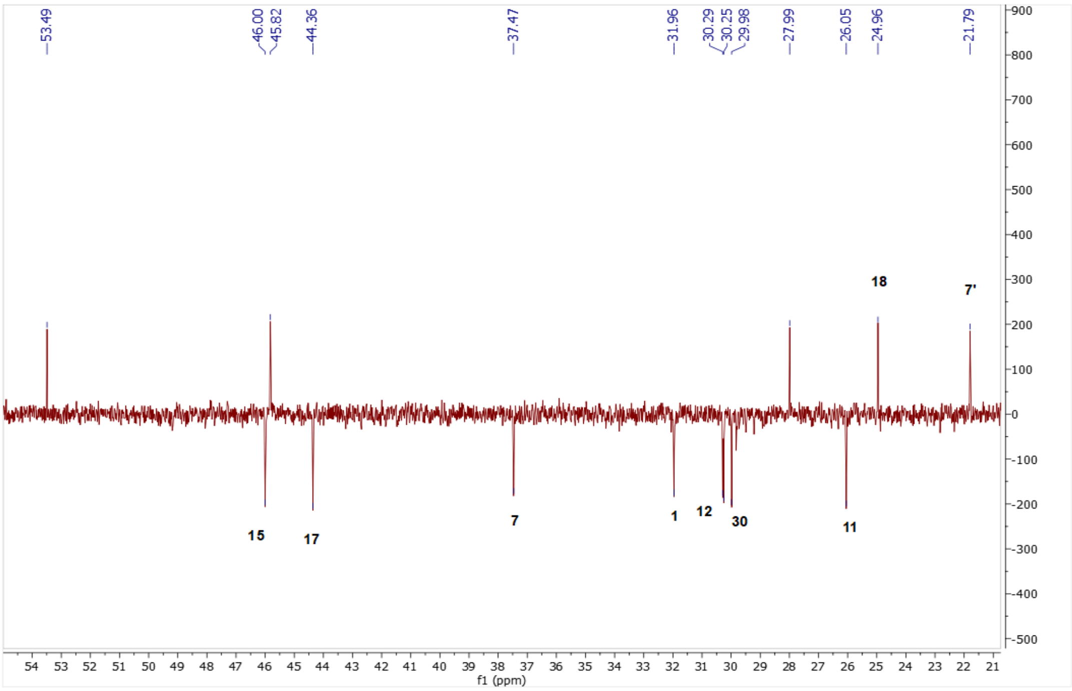

Spectrum 78. DEPT135 spectrum of SCG-01.

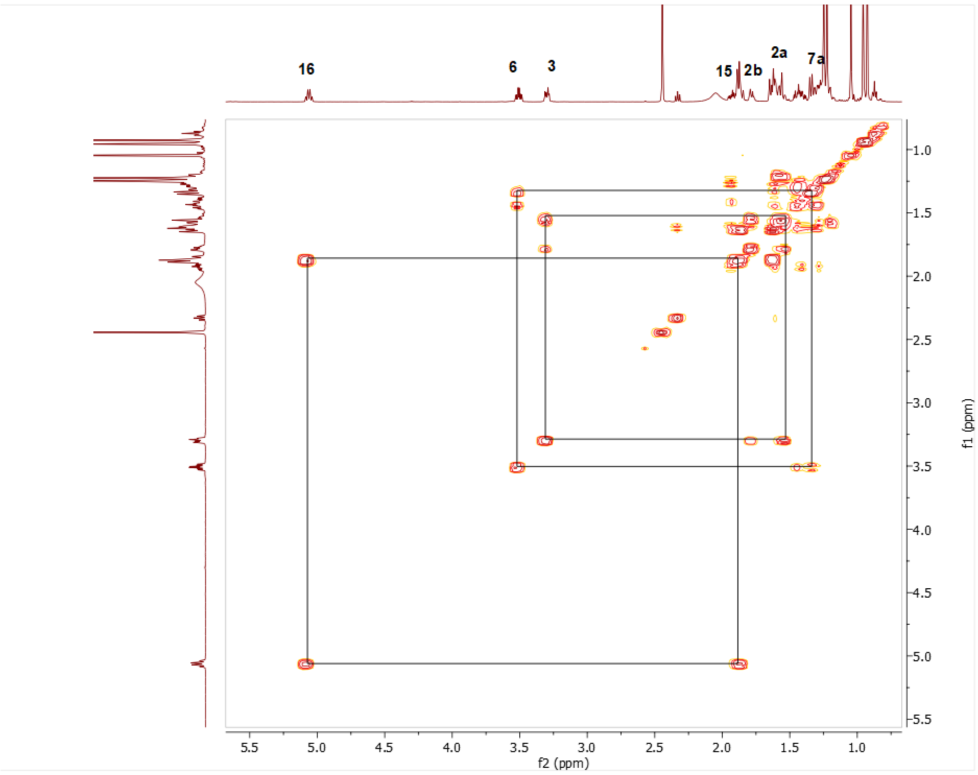

Spectrum 79. COSY spectrum of SCG-01.

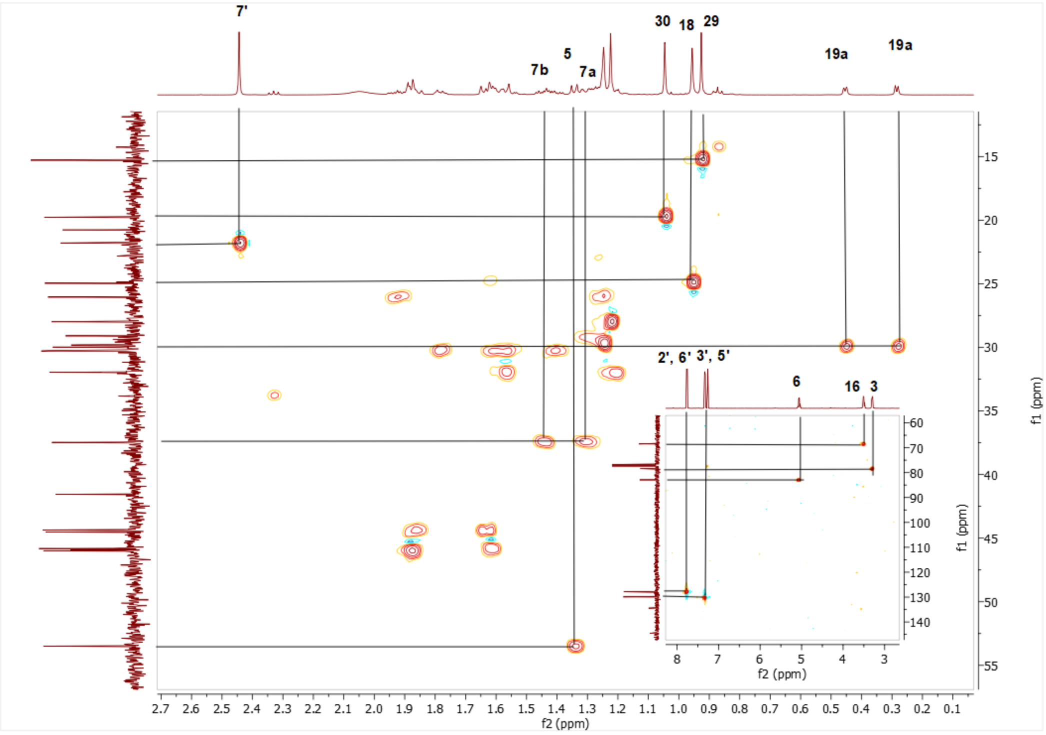

Spectrum 80. HMQC spectrum of SCG-01.

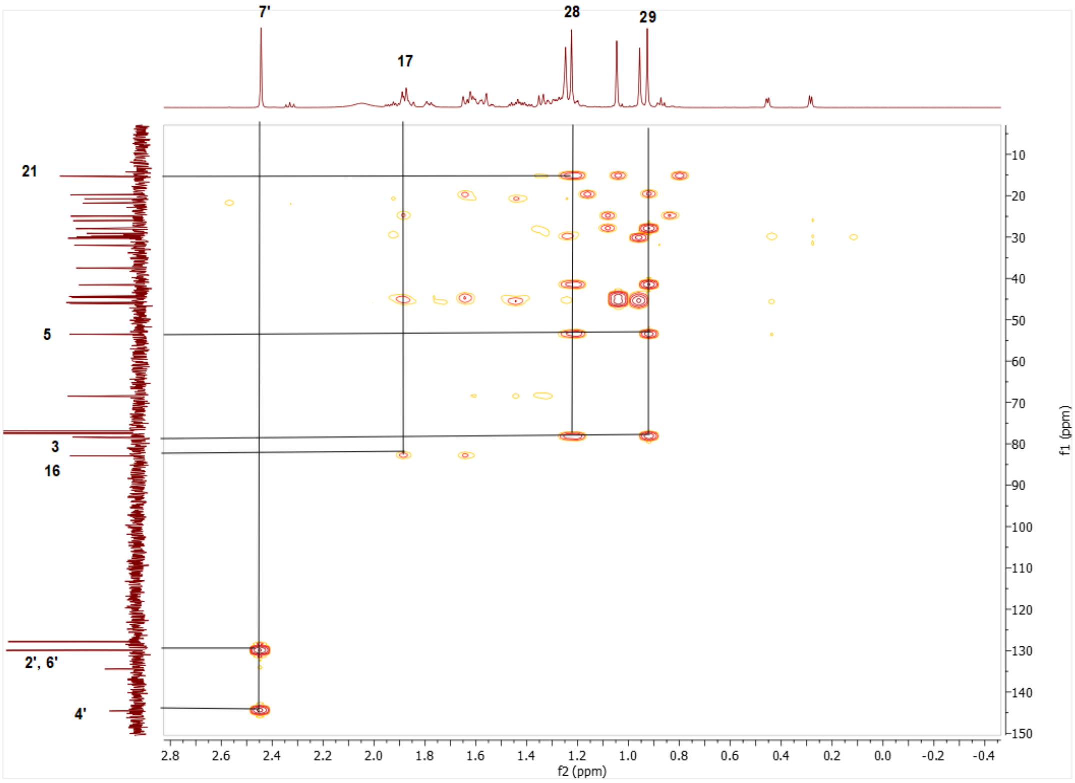

Spectrum 81. HMBC spectrum of SCG-01.

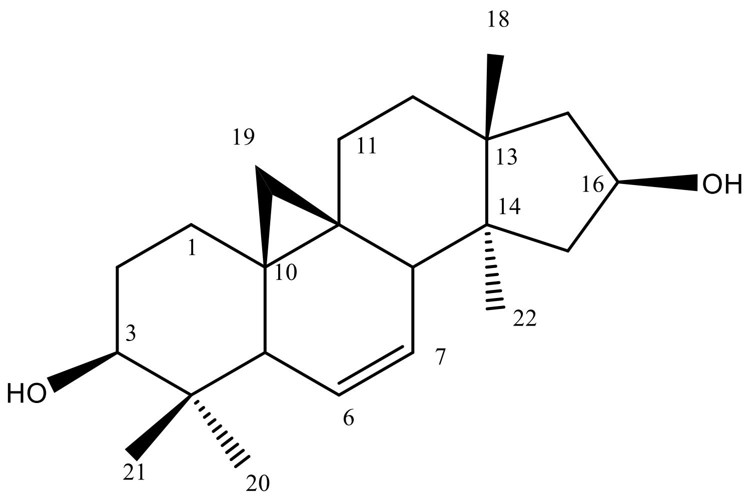

Supplementary Figure 15. Chemical Structure of SCG-02

Supplementary Table 13. The $^{13}C$ and $^{1}H$ NMR data of SCG-02 (100/400 MHz, δ ppm, in CDCl3).

| H/C | $\delta_C$ (ppm) | $\delta_H$ (ppm), *J* (Hz) |
|---|---|---|
| **1** | 29.8 t | 1.25 m, 1.46 m |
| **2** | 30.1 t | 1.6 m, 1.82 m |
| **3** | 78.5 d | 3.3 dd (11.2, 4.4) |
| **4** | 40.3 s | - |
| **5** | 46.5 d | 1.89 m |
| **6** | 126.3 d | 5.62 d (10.5) |
| **7** | 129.1 d | 5.44 ddd (10.6, 6.1, 3.2) |
| **8** | 43.7 d | 2.48 dd (6.2, 2.6) |
| **9** | 21.2 s | - |
| **10** | 28.3 s | - |
| **11** | 25.2 t | 1.42 m, 1.85 m |
| **12** | 31.4 t | 1.22 m, 1.69 m |
| **13** | | |
| | 48.5 s | - |
| **14** | 45.3 s | - |
| **15** | 45.4 t | 1.27 m, 2.05 dd (13.7, 8.2) |
| **16** | 72.2 d | 4.55 ddd (14.5, 7.7, 1.4) |
| **17** | 47.8 t | 1.86 m, 1.62 m |
| **18** | 22.2 q | 0.96 s |
| **19** | 18.5 t | -0.15 d (4.1), 0.73 d (4.5) |
| **28** | 25.6 q | 1.05 s |
| **29** | 14.5 q | 0.77 s |
| **30** | 18.4 q | 0.92 s |

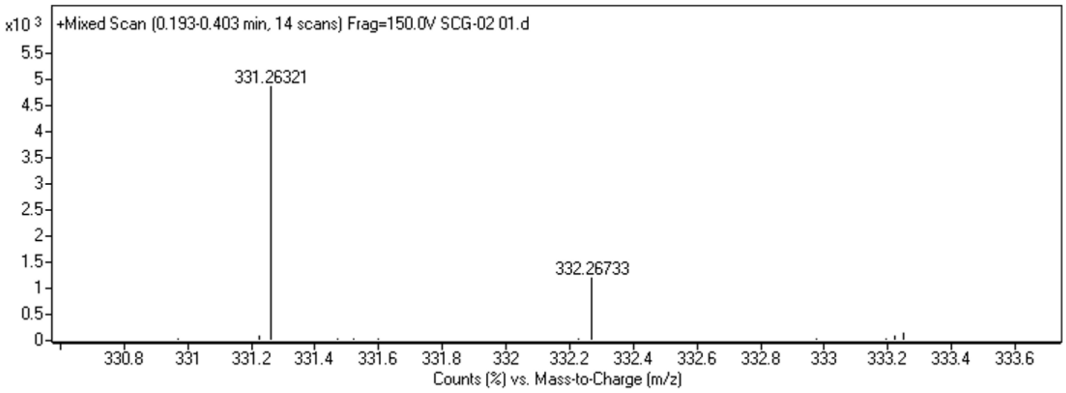

Spectrum 82. HR-ESI-MS Spectrum of SCG-02 (positive mode).

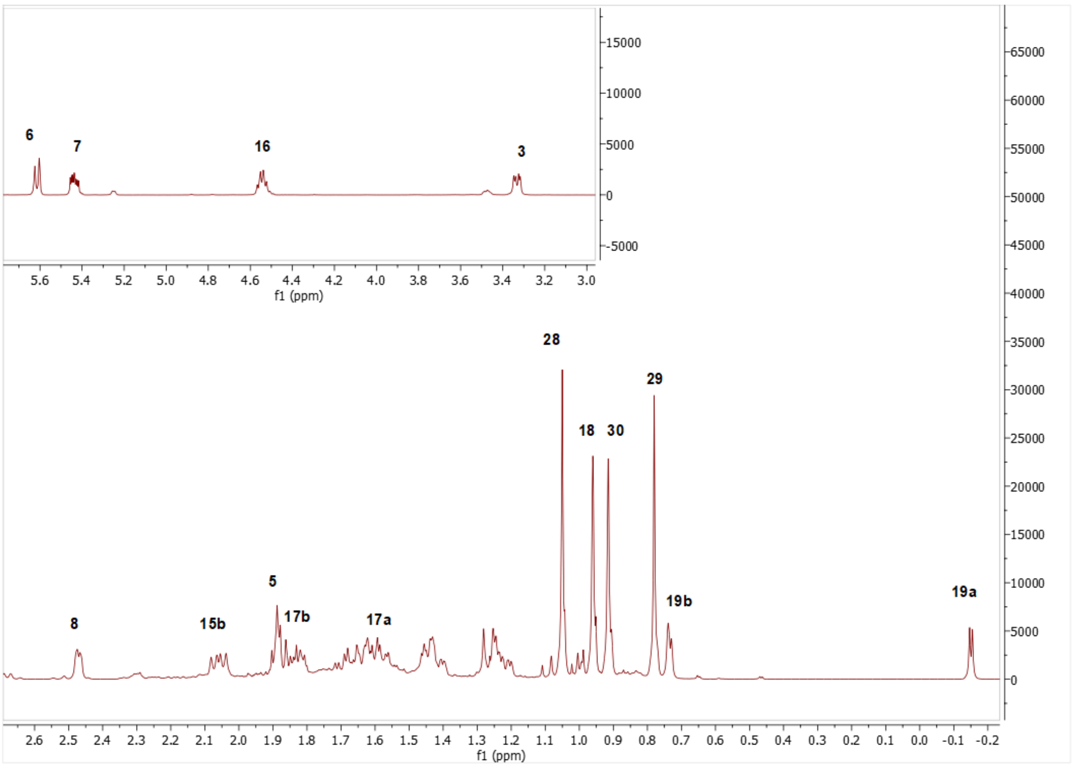

Spectrum 83. $^{1}$H NMR Spectrum of SCG-02.

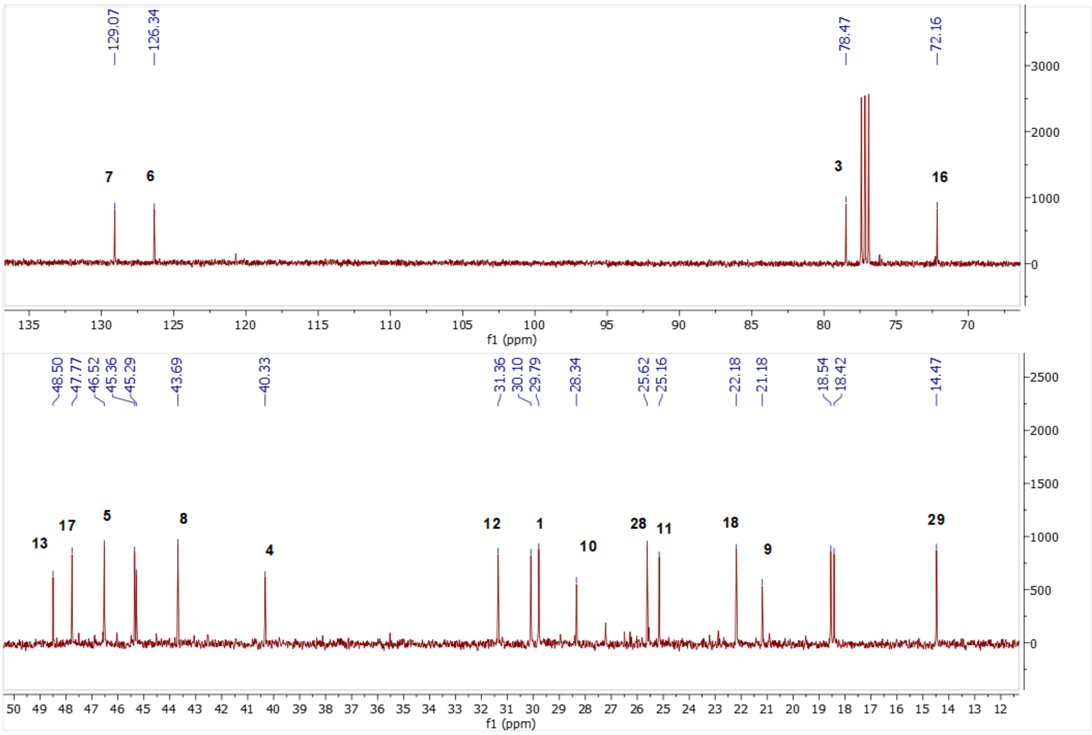

Spectrum 84. $^{13}$C NMR Spectrum of SCG-02.

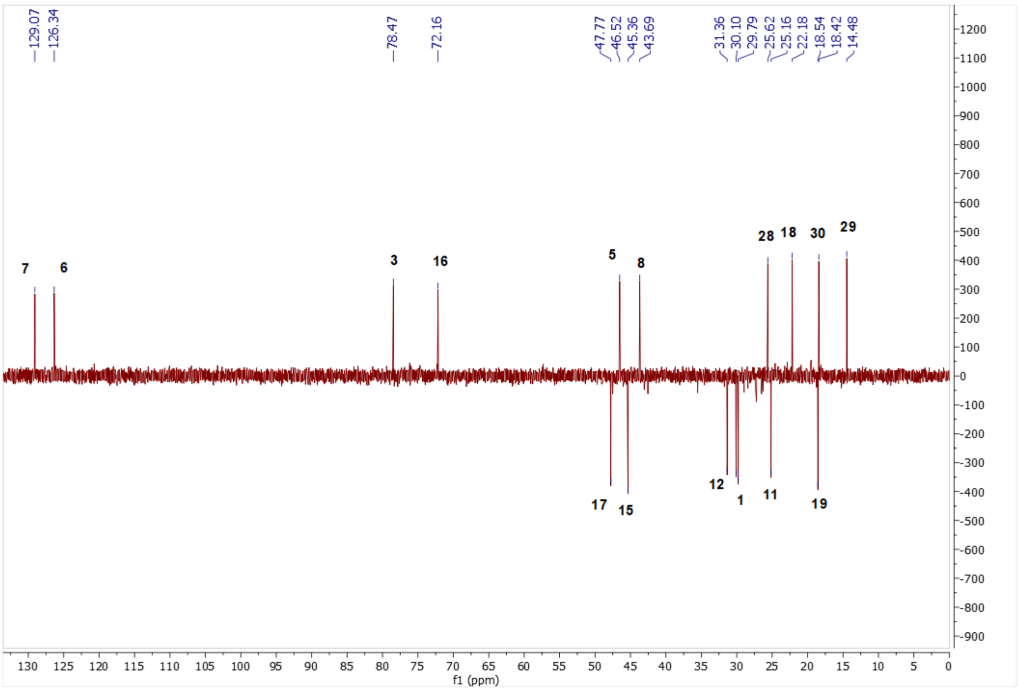

Spectrum 85. DEPT135 spectrum of SCG-02.

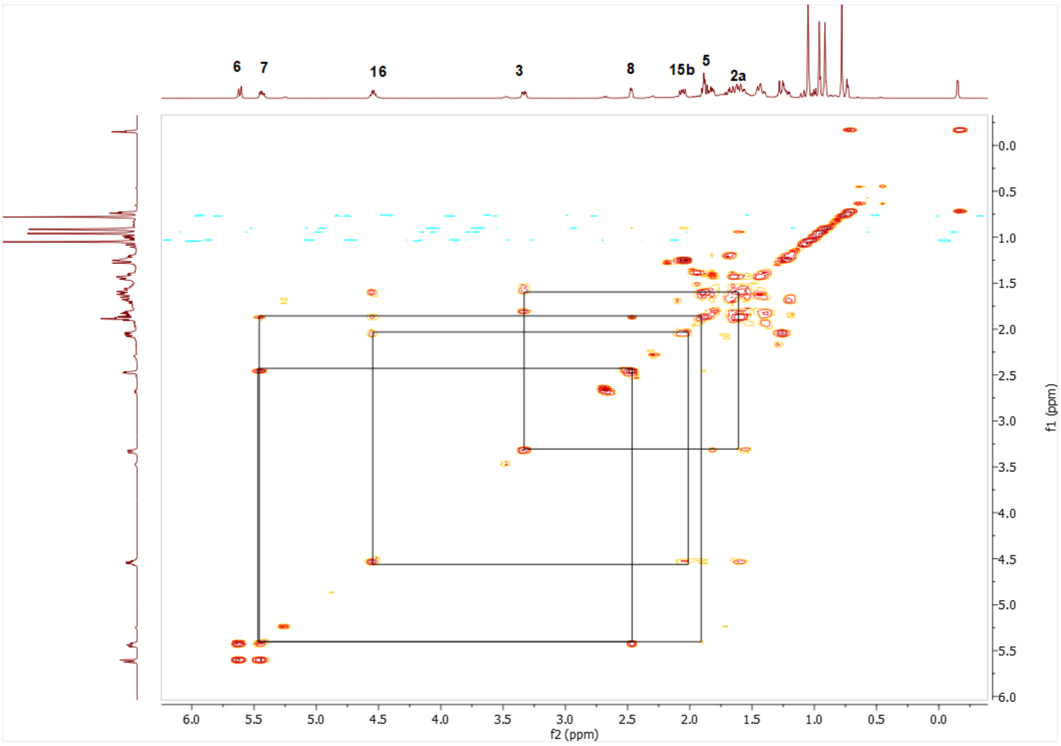

Spectrum 86. COSY spectrum of SCG-02.

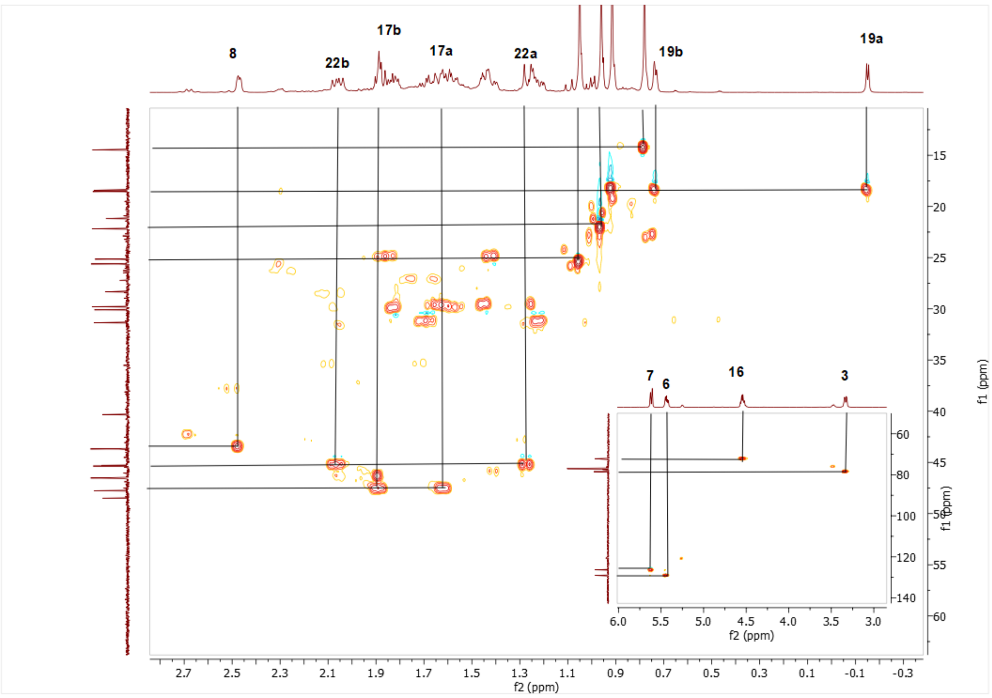

Spectrum 87. HSQC spectrum of SCG-02.

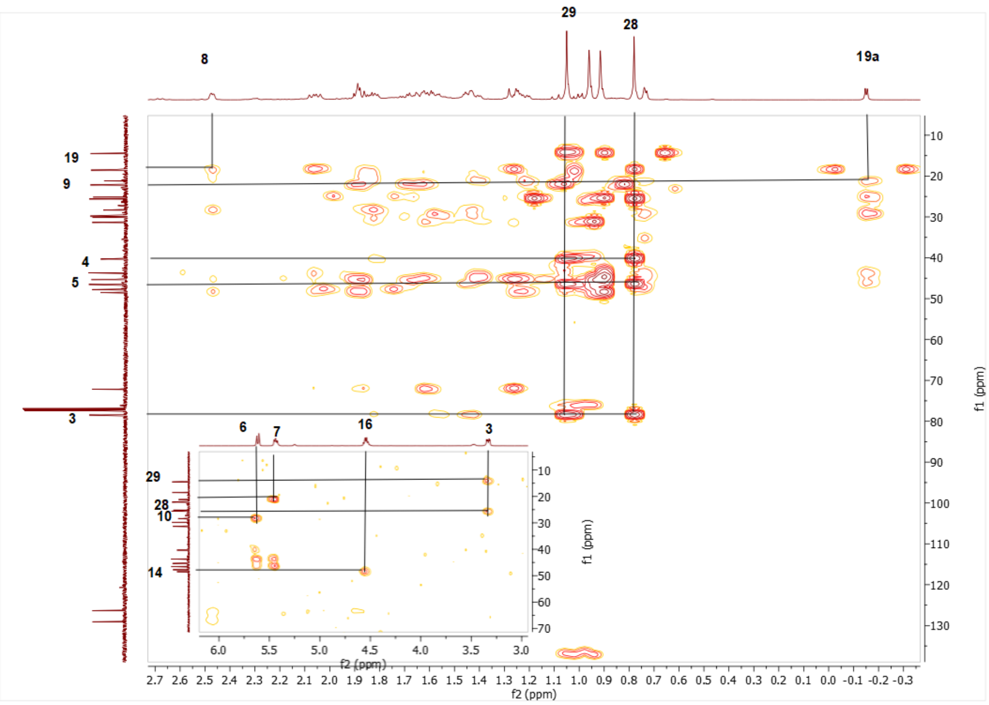

Spectrum 882. HMBC spectrum of SCG-02.

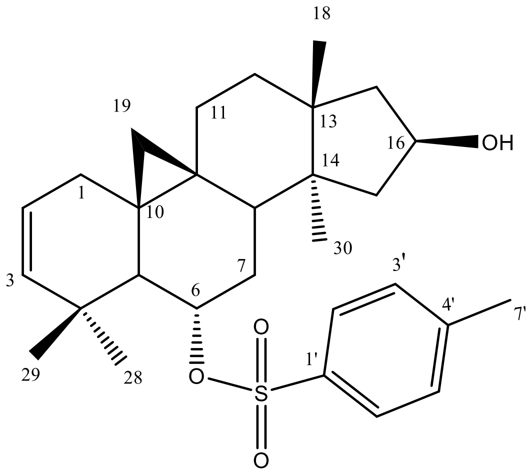

Supplementary Figure 16. Chemical Structure of SCG-03

Supplementary Table 14. The $^{13}$C and $^{1}$H NMR data of SCG-03 (100/400 MHz, δ ppm, in CDCl3).

| H/C | δC (ppm) | δH (ppm), *J* (Hz) |
|---|---|---|
| **1** | 34.9 t | 1.39 m, 2.30 m |
| **2** | 122.8 d | 5.49 ddd (9.9, 5.8, 2) |
| **3** | 140.7 | 5.3 dd (9.8, 2.7) |
| **4** | 38.0 s | - |
| **5** | 52.6 d | 1.58 m |
| **6** | 83 d | 5.04 q (7.4) |

| | | |
|---|---|---|
| **7** | 28.4 t | 1.34 m, 1.90 m |
| **8** | 47.9 d | 1.54 m |
| **9** | 19.2 s | - |
| **10** | 28.2 s | - |
| **11** | 25.8 t | 1.1 m, 2.07 m |
| **12** | 30.16 | 1.47 m, 1.62 m |
| **13** | 44.4 s | - |
| **14** | 45.9 s | - |
| **15** | 44.9 t | 1.65 m, 1.87 m |
| **16** | 70.7 d | 3.45 td (9.8, 4.7) |
| **17** | 46.1 t | 1.88 m |
| **18** | 25.7 q | 0.99 s |
| **19** | 31.7 q | 0.34 d (4.5), 0.51 d (5) |
| **28** | 33.1 q | 1.23 s |
| **29** | 23.5 q | 1.05 s |
| **30** | 20.2 q | 1.03 s |
| **1’** | 134.2 s | - |
| **2’** | 127.8 d | 7.74 dd (8.2, 3.6) |
| **3’** | 129.9 d | 7.31 dd (8.2, 3.6) |
| **4’** | 144.6 s | - |
| **5’** | 129.9 d | 7.31 dd (8.2, 3.6) |
| **6’** | 127.8 d | 7.74 dd (8.2, 3.6) |
| **7’** | 21.8 q | 2.44 s |

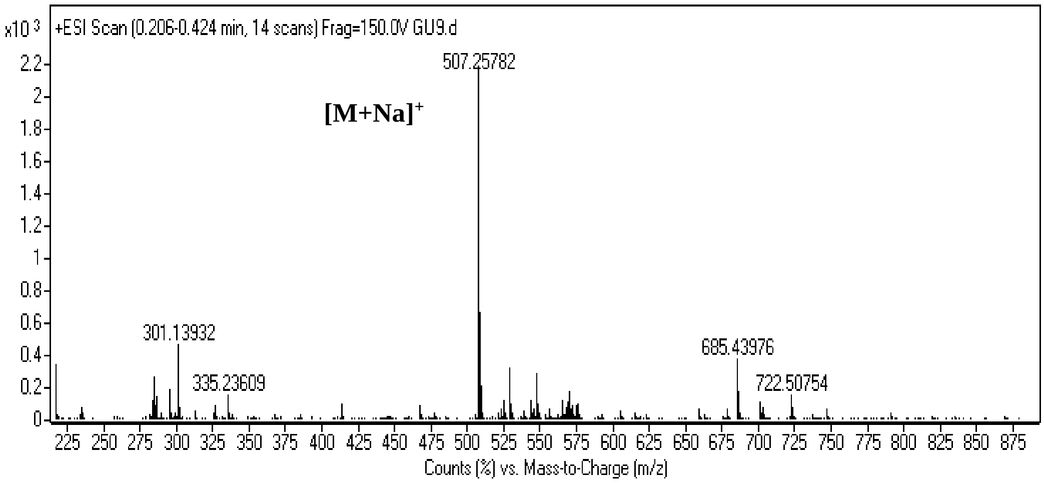

Spectrum 89. HR-ESI-MS Spectrum of SCG-03 (positive mode).

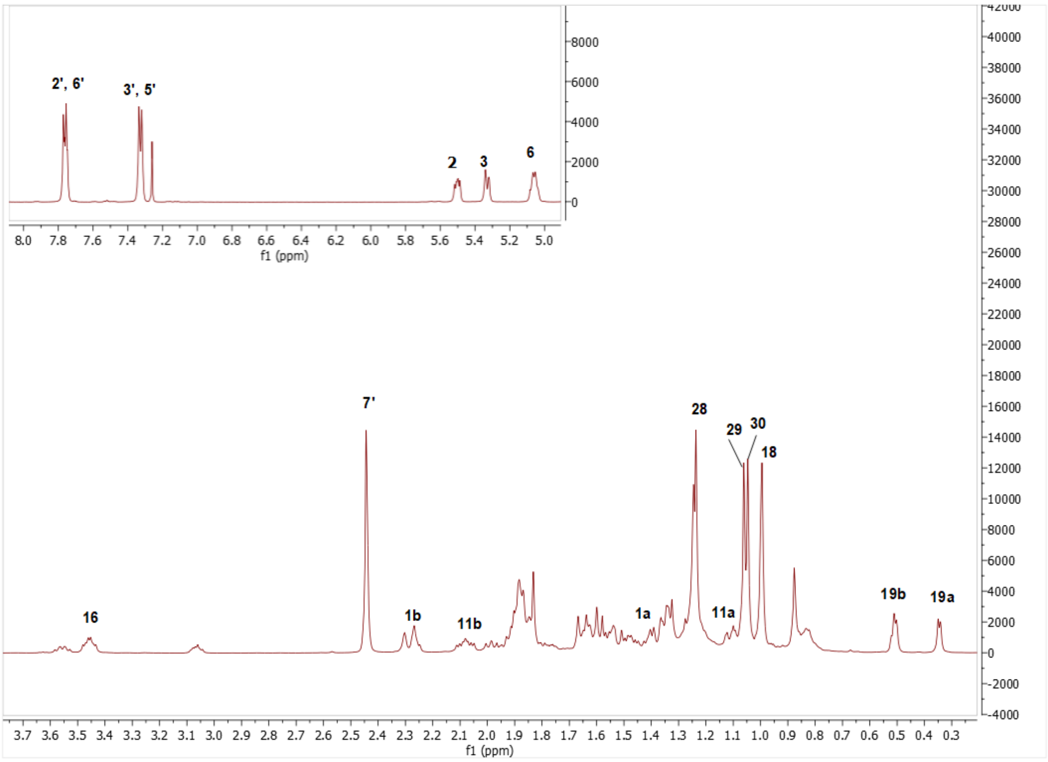

Spectrum 90. $^{1}H$ NMR Spectrum of SCG-03.

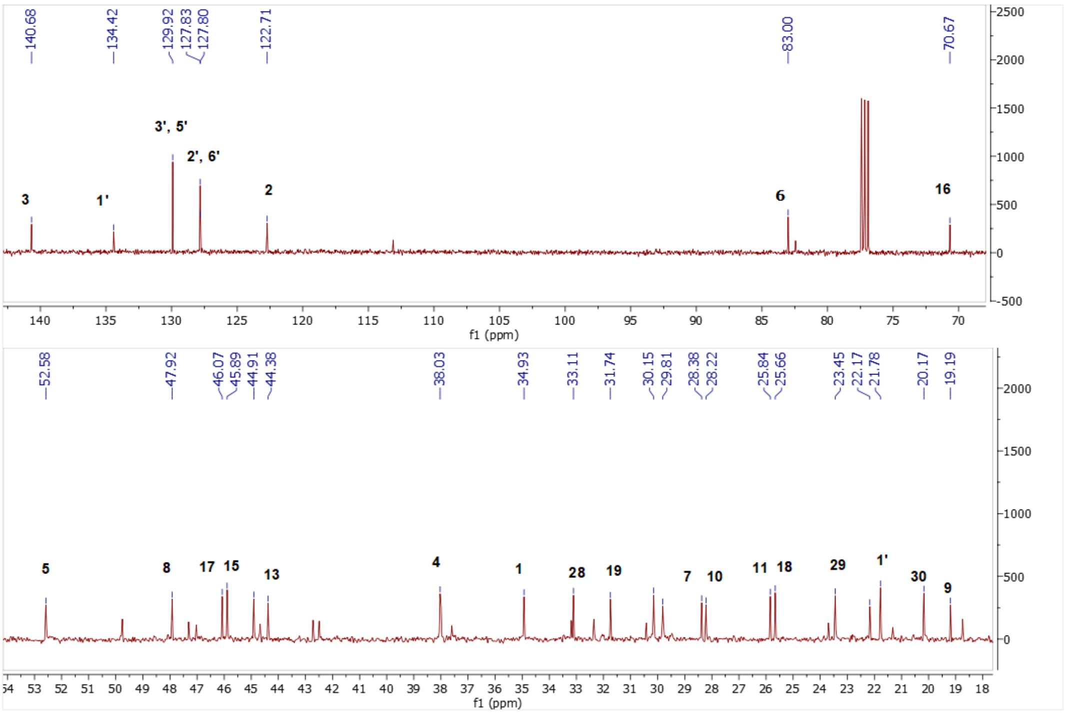

Spectrum 91. $^{13}C$ NMR Spectrum of SCG-03.

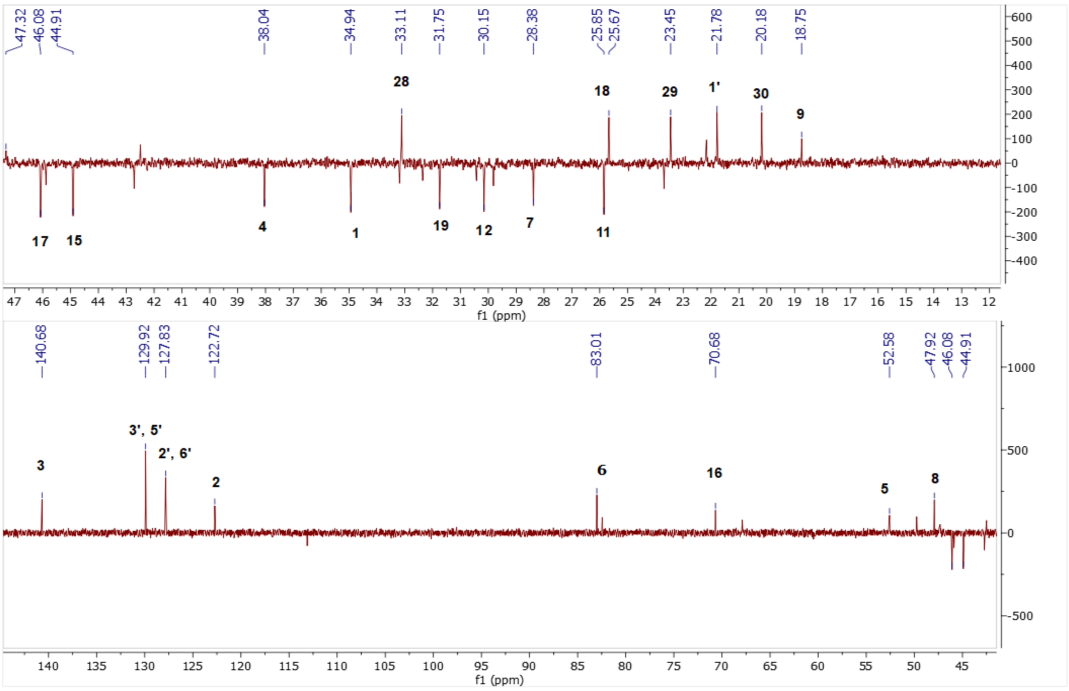

Spectrum 92. DEPT135 spectrum of SCG-03.

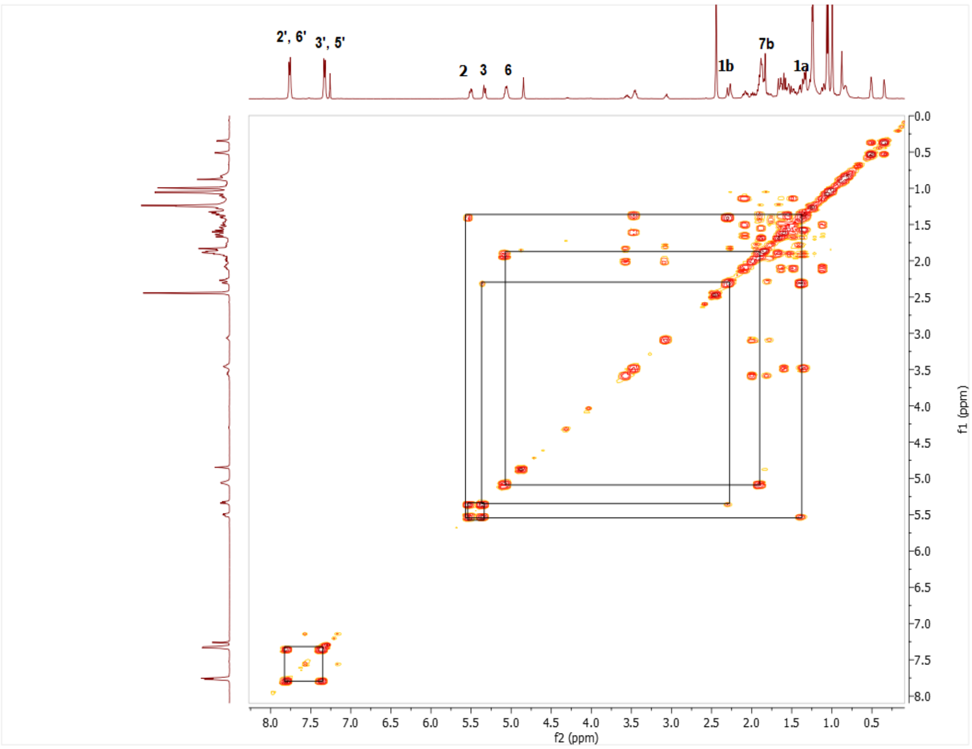

Spectrum 93. COSY spectrum of SCG-03.

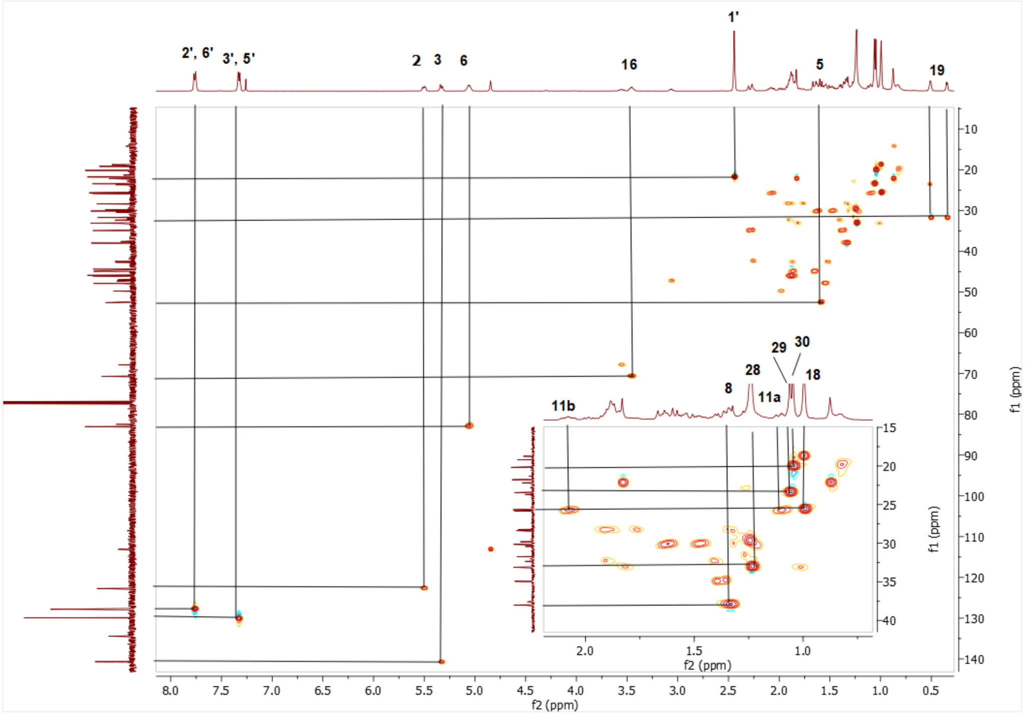

Spectrum 94. HMQC spectrum of SCG-03.

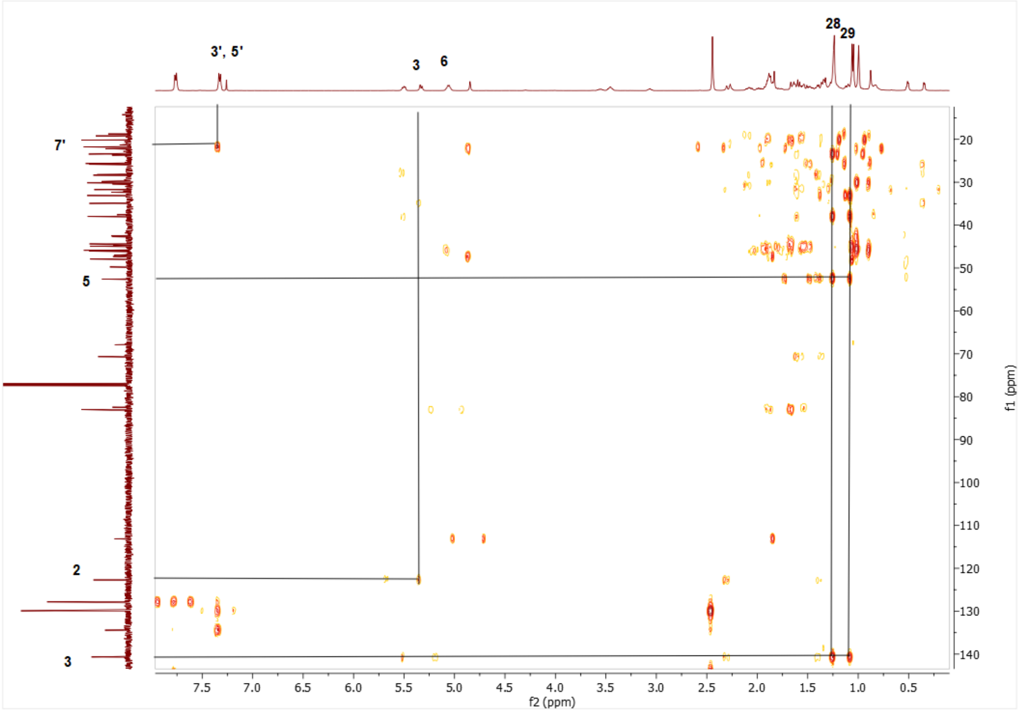

Spectrum 95. HMBC spectrum of SCG-03.

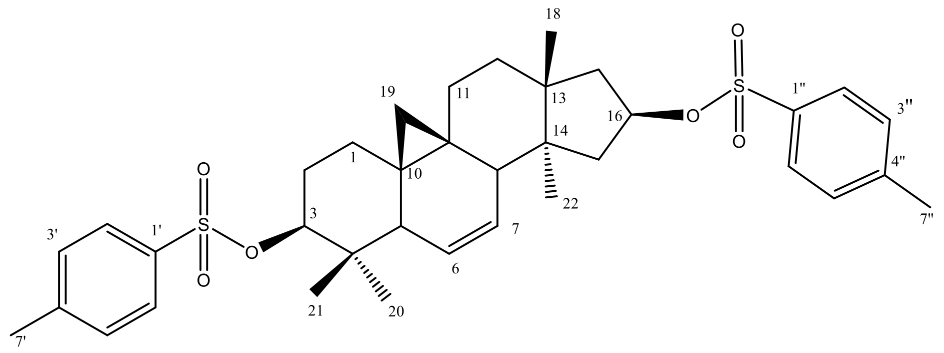

Supplementary Figure 17. Chemical Structure of SCG-04

Supplementary Table 15. The $^{13}$C and $^{1}$H NMR data of SCG-04 (100/400 MHz, δ ppm, in CDCl3).

| **H/C** | **$\delta_C$ (ppm)** | **$\delta_H$ (ppm), *J* (Hz)** |
|---|---|---|
| **1** | 29.4 t | 1.4 m,1.6 m |
| **2** | 27.6 t | 1.8 m, 1.9 m |
| **3** | 89.9 d | 4.3 dd (11.6, 4.5) |
| **4** | 39.8 s | - |
| **5** | 46.5 d | 1.89 m |
| **6** | 125,7 d | 5.52 d (10.7) |
| **7** | 128.9 d | 5.36 ddd (9.6, 5.9, 2.9) |
| **8** | 43.1 d | 2.41 m |
| **9** | 20.9 s | - |
| **10** | 27.7 s | - |
| **11** | 24.79 | 1.8 m, 1.39 m |
| **12** | 30.7 | 1.16 m, 1.61 |
| **13** | 44.4 s | - |
| **14** | 48.1 s | - |
| **15** | 44.9 t | 1.81 m |
| **16** | 82.4 d | 5.06 q (7.5) |
| **17** | 41.9 t | 2 m, 1.5 m |
| **18** | 21.8 q | 0.88 s |
| **19** | 18.3 t | -0.16 d (4.1), 0.71 d (4.1) |
| **28** | 25.3 q | 0.83 s |
| **29** | 15.2 q | 0.79 s |
| **30** | 17.6 q | 0.82 s |
| **1’** | 134.7* s | - |
| **2’** | 127.7 d | 7.75+ d (7.8) |
| **3’** | 129.7’ d | 7.32 d (7.8) |
| **4’** | 144.4- s | - |
| **5’** | 129.7’ d | 7.32 d (7.8) |
| **6’** | 127.7 d | 7.75+ d (7.8) |
| **7’** | 21.6 q | 2.44 s |
| **1’’** | 134.2* s | - |
| **2’’** | 127.7 d | 7.8+ d (7.7) |

| | | |
|---|---|---|
| **3''** | 129.7' d | 7.32 d (7.8) |
| **4''** | 144.7- s | - |
| **5''** | 129.7' d | 7.32 d (7.8) |
| **6''** | 127.7 d | 7.8+ d (7.7) |
| **7''** | 21.6 q | 2.44 s |

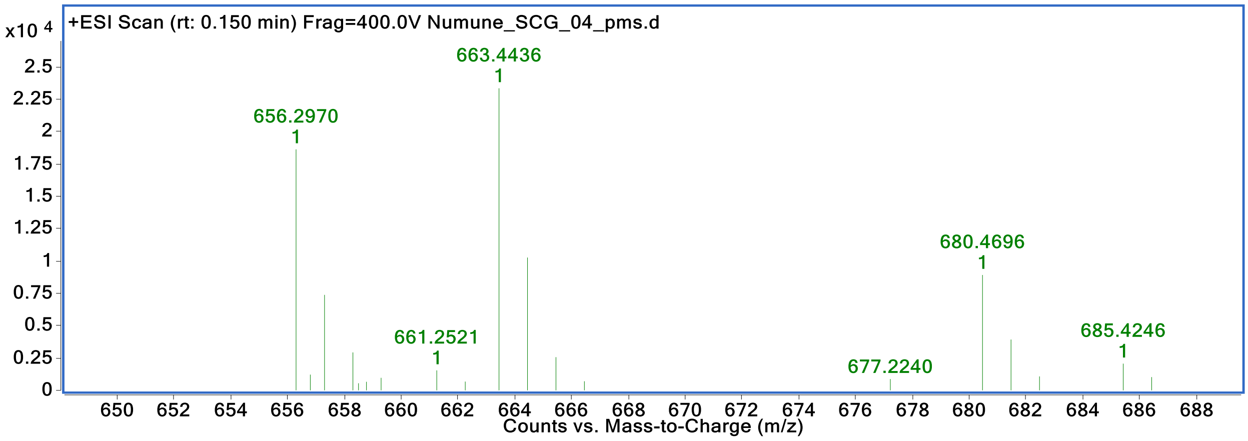

Spectrum 96. HR-ESI-MS Spectrum of SCG-04 (positive mode).

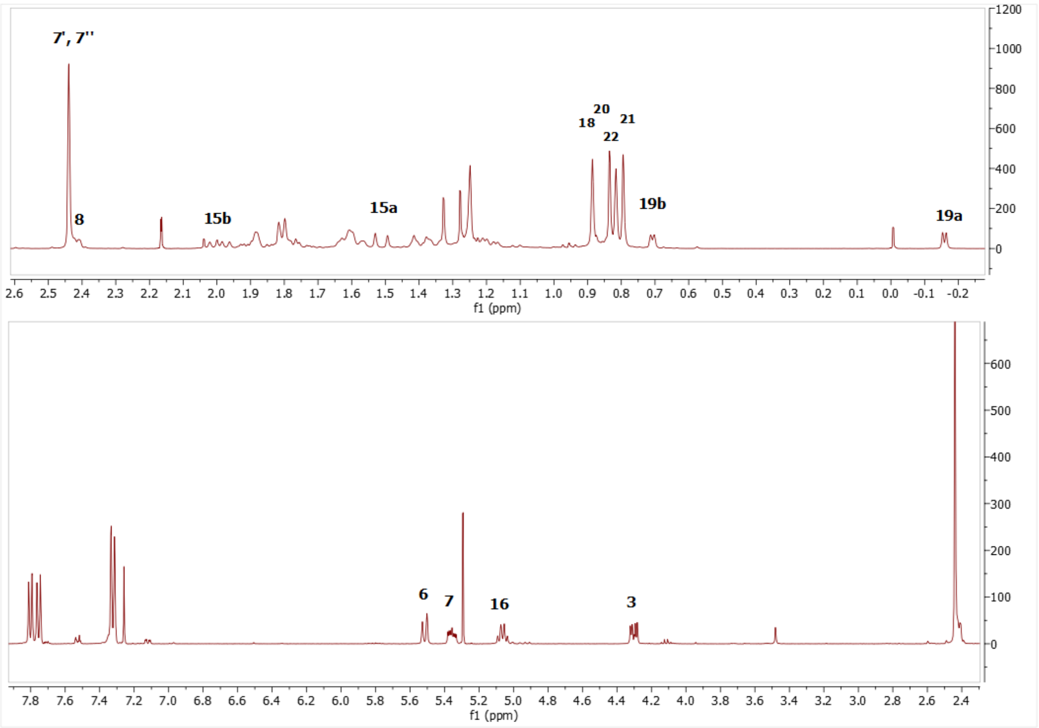

Spectrum 97. $^1$H NMR Spectrum of SCG-04.

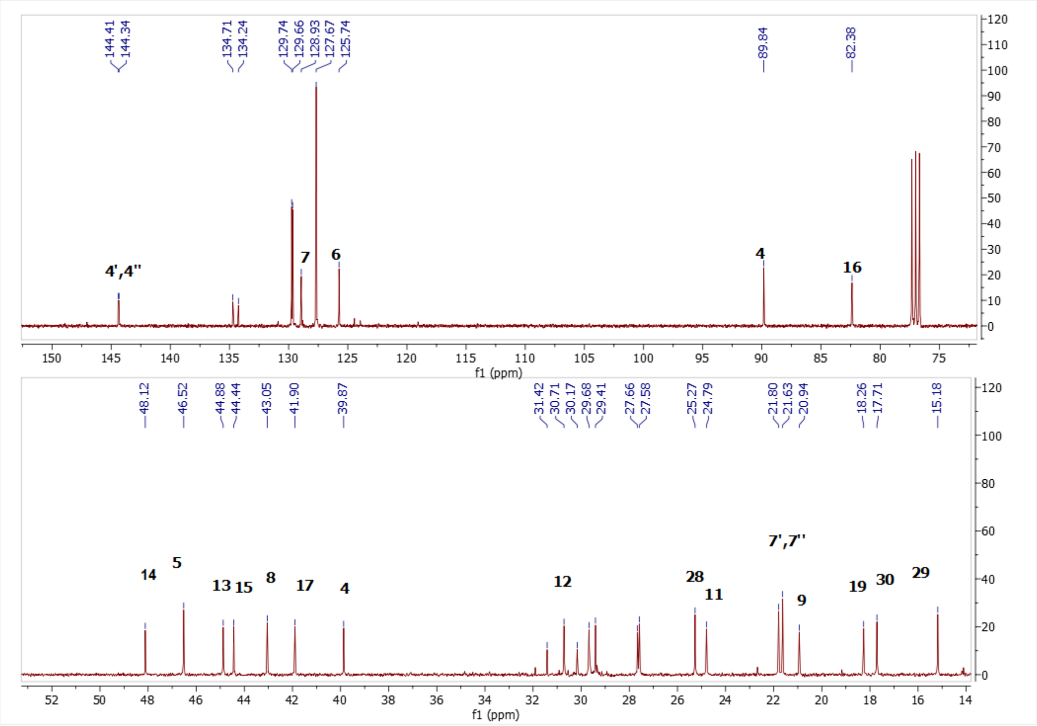

Spectrum 98. $^{13}$C NMR Spectrum of SCG-04.

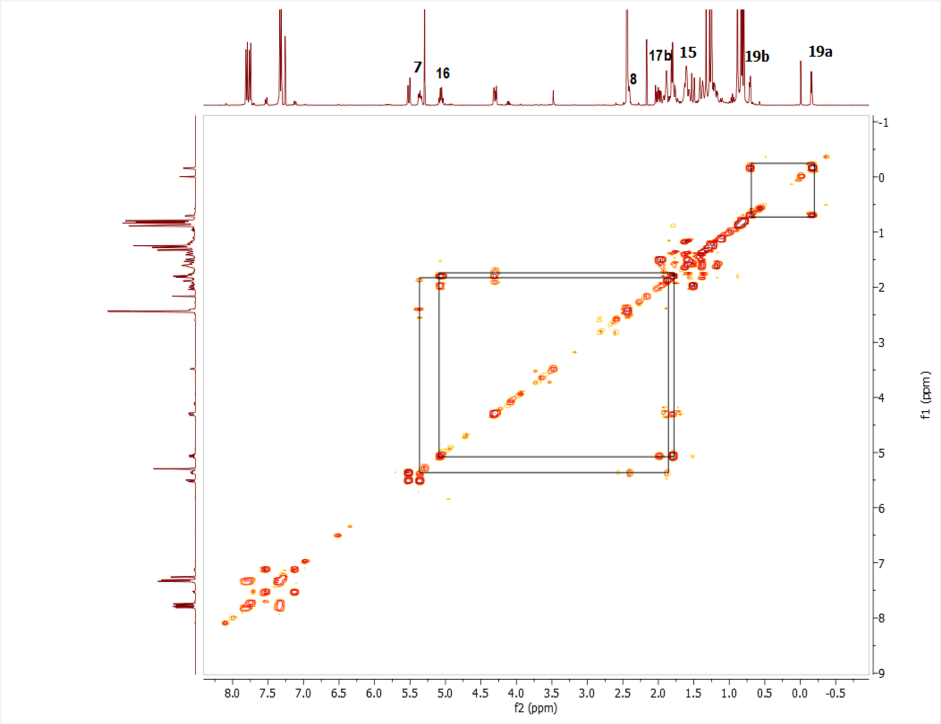

Spectrum 99. COSY spectrum of SCG-04.

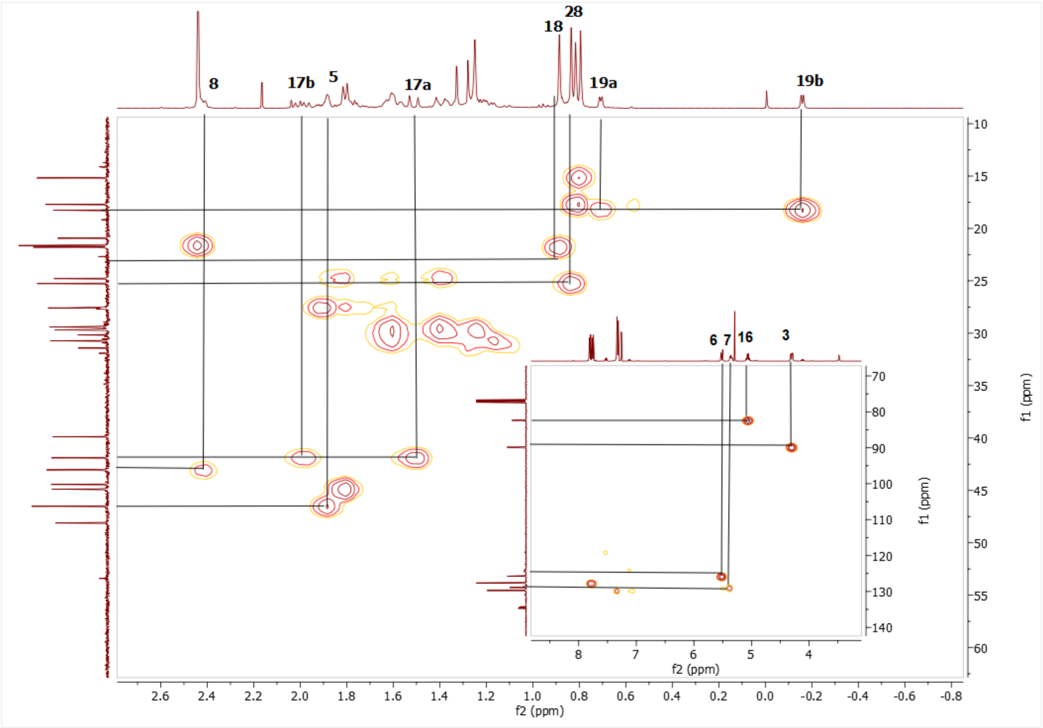

Spectrum 100. HMQC spectrum of SCG-04.

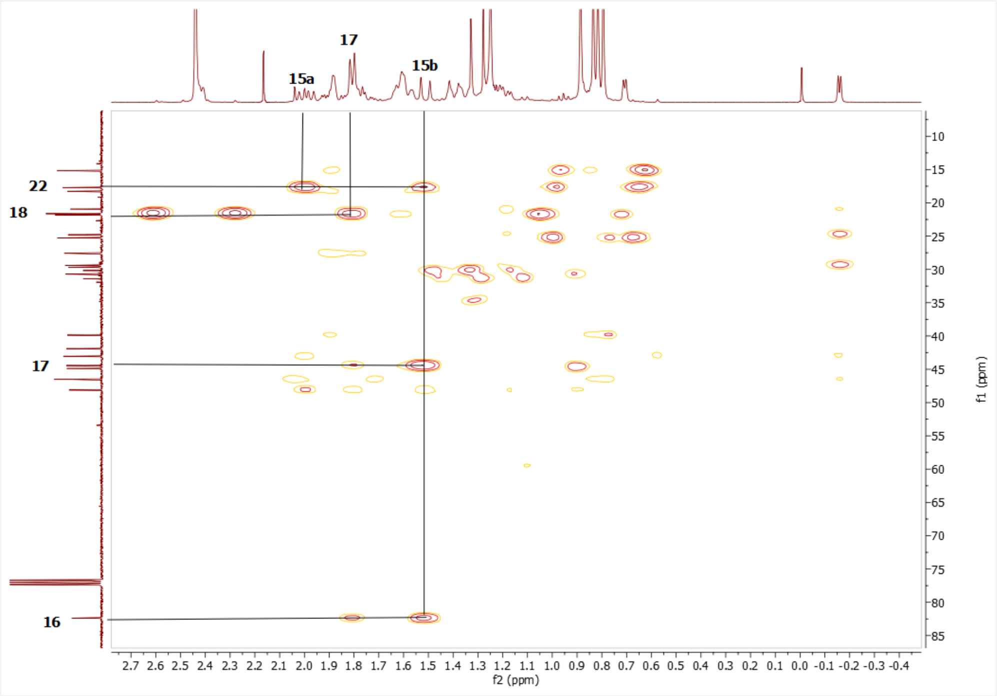

Spectrum 1013. HMBC spectrum of SCG-04.

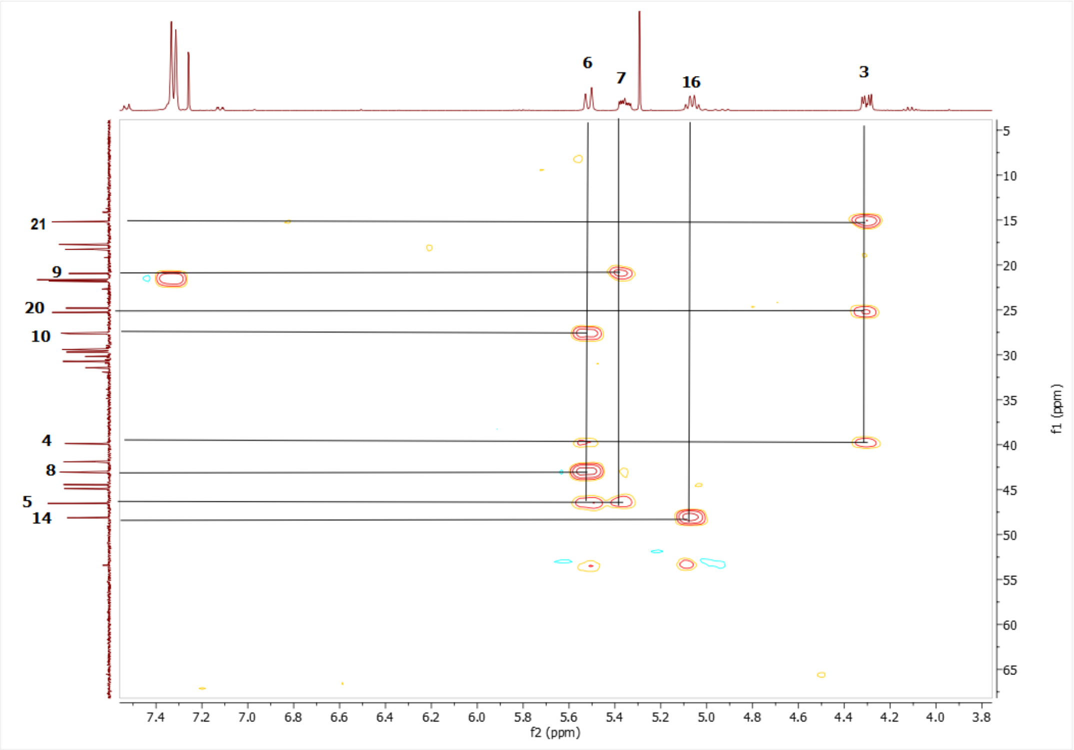

Spectrum 1024. HMBC spectrum of SCG-04.

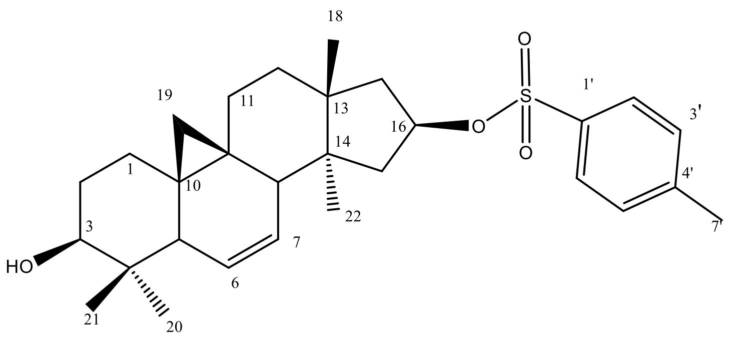

Supplementary Figure 18. Chemical Structure of SCG-05

Supplementary Table 16. The $^{13}C$ and $^{1}H$ NMR data of SCG-05 (100/500 MHz, δ ppm, in CDCl3).

| H/C | $\delta_C$ (ppm) | $\delta_H$ (ppm), *J* (Hz) |
|---|---|---|
| **1** | 29.8 t | 1.23 s, 1.43 s |
| **2** | 30.1 t | 1.62 m, 1.82 s |
| **3** | 78.5 d | 3.33 dd (10.8, 3.9) |
| **4** | 40.4 s | - |
| **5** | 46.5 d | 1.86 d (2.4) |
| **6** | 126.9 d | 5.61 d (10.5) |
| **7** | 128.5 d | 5.37 m |
| **8** | 43.3 d | 2.44 m |
| **9** | 21.1 s | - |
| **10** | 28.3 s | - |
| **11** | 25.0 t | 1.81 d (5.6), 1.42 m |
| **12** | 31.0 t | 1.63 d (12.9), 1.2 dd (12.7, 4.3) |
| **13** | 48.4 s | - |
| **14** | 44.6 s | - |
| **15** | 45.2 t | 1.83 s |
| **16** | 82.7 | 5.07 dd (14.7, 7.2) |
| **17** | 42.1 t | 1.53 d (14.7), 2.01 dt (23.2, 11.6) |
| **18** | 21.8 q | 0.9 s |
| **19** | 18.5 t | 0.72d (3.1), -0.15d (3.9) |
| **28** | 25.6 q | 1.05 s |
| **29** | 14.5 q | 0.77 s |
| **30** | 17.9 q | 0.85 s |
| **1’** | 134.5 s | - |
| **2’** | 127.8 d | 7.76 d (8.1) |
| **3’** | 129.9 d | 7.32 d (7.9) |
| **4’** | 127.8 d | 7.76 d (8.1) |
| **5’** | 129.9 d | 7.32 d (7.9) |
| **6’** | 144.5 s | - |
| **7’** | 21.6 q | 2.44 s |

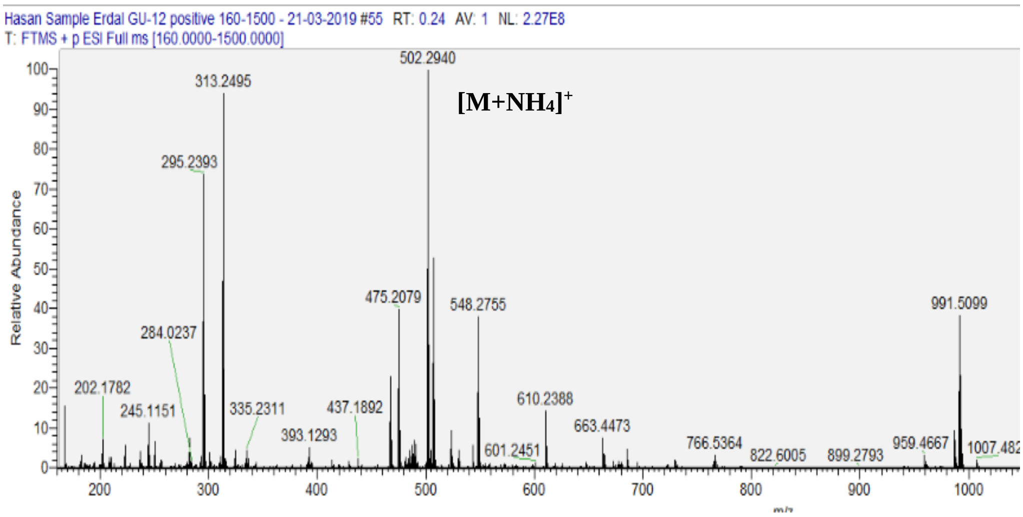

Spectrum 103. HR-ESI-MS Spectrum of SCG-05 (positive mode).

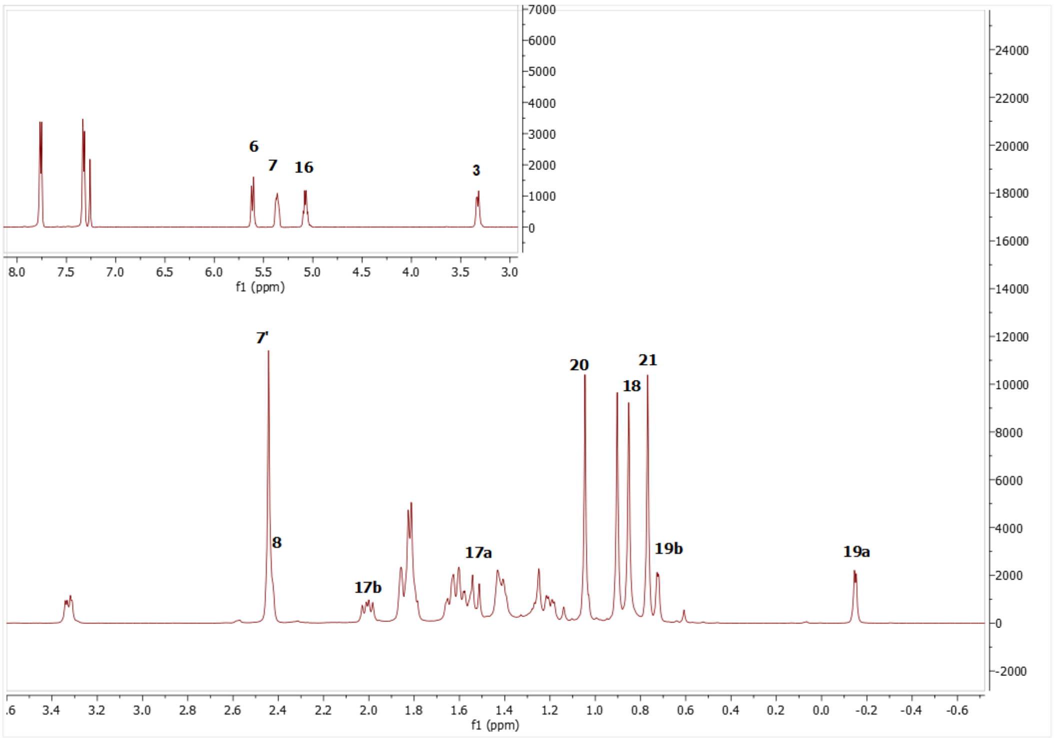

Spectrum 104. $^{1}$H NMR Spectrum of SCG-05.

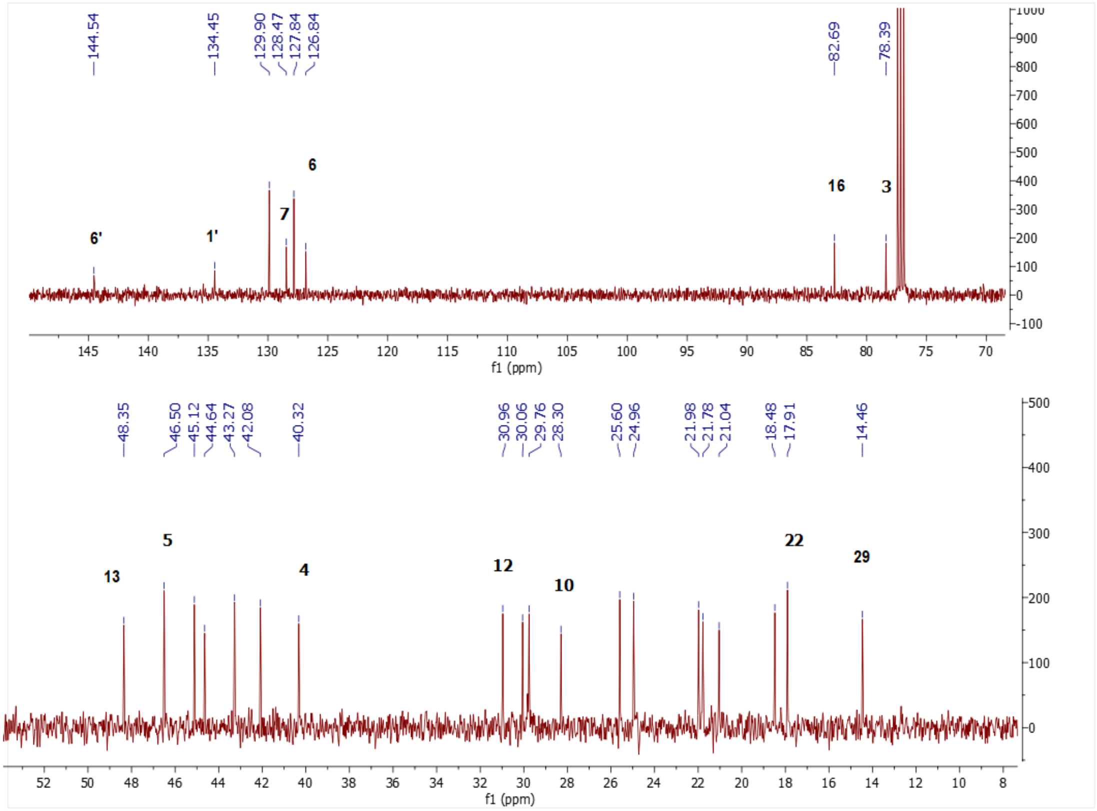

Spectrum 105. $^{13}$C NMR Spectrum of SCG-05.

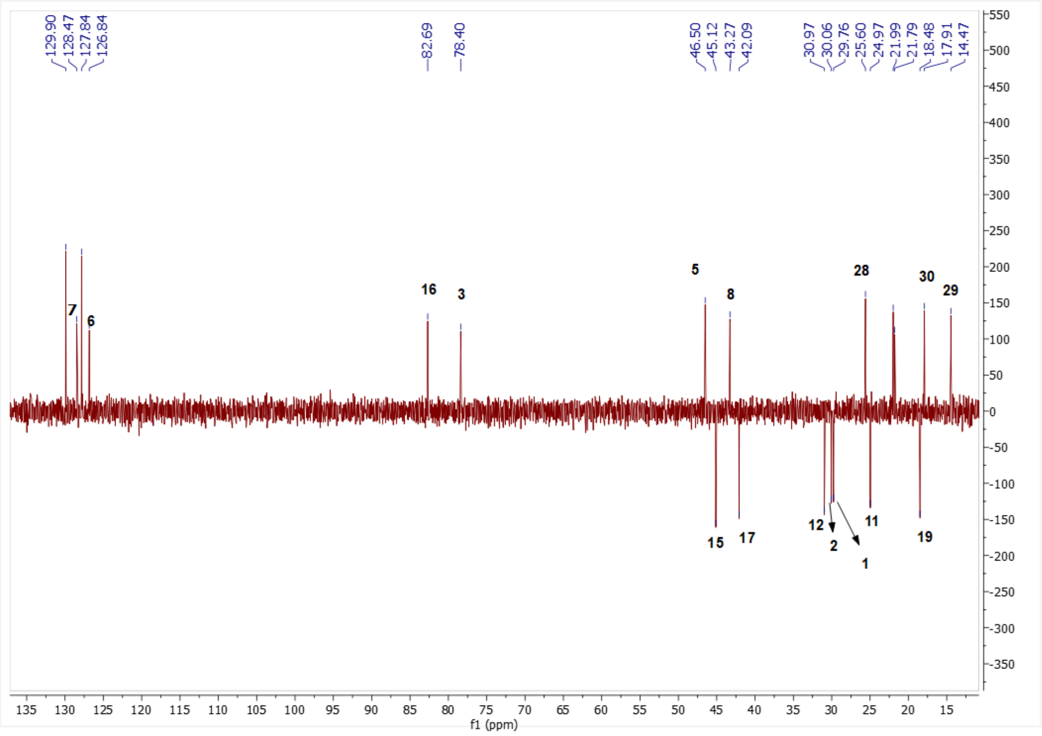

Spectrum 106. DEPT135 spectrum of SCG-05.

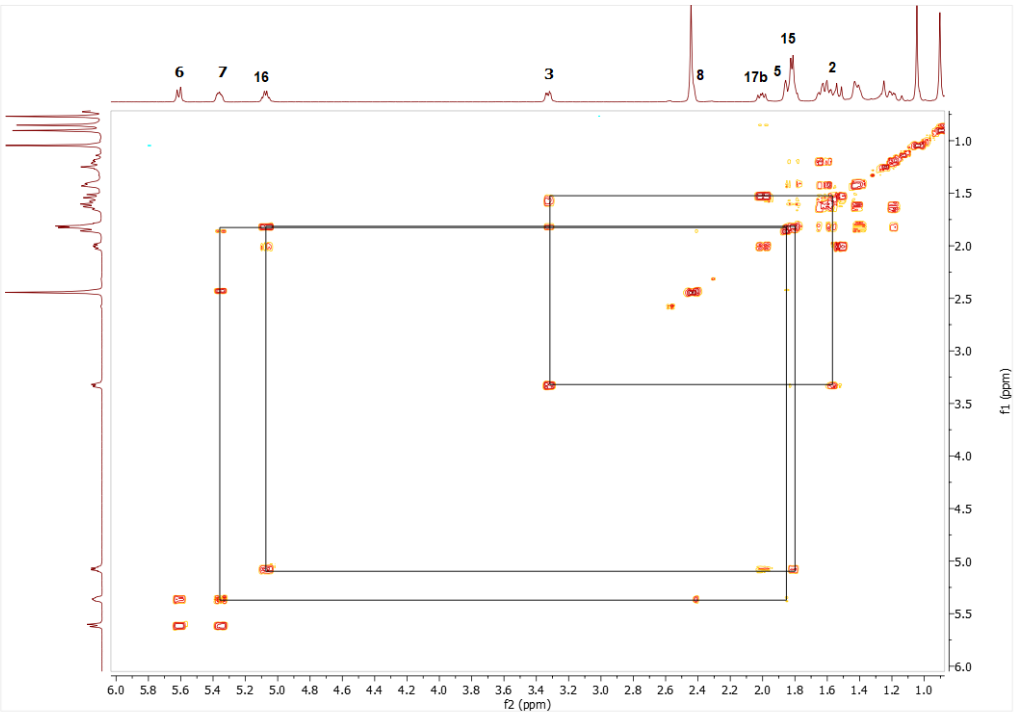

Spectrum 1075. COSY spectrum of SCG-05.

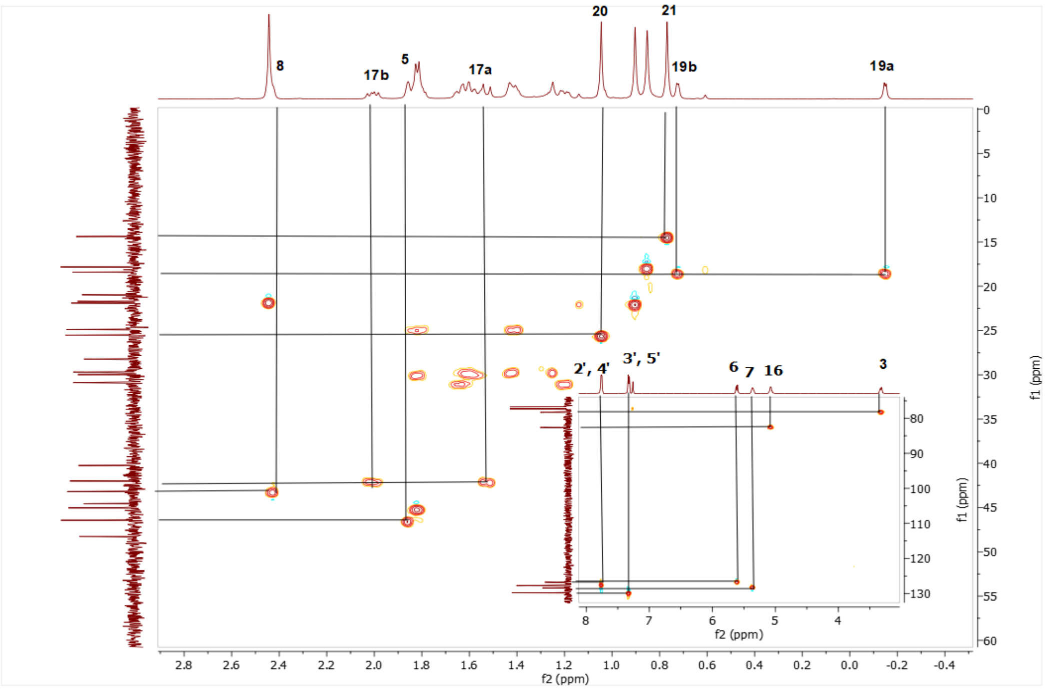

Spectrum 1086. HMQC spectrum of SCG-05.

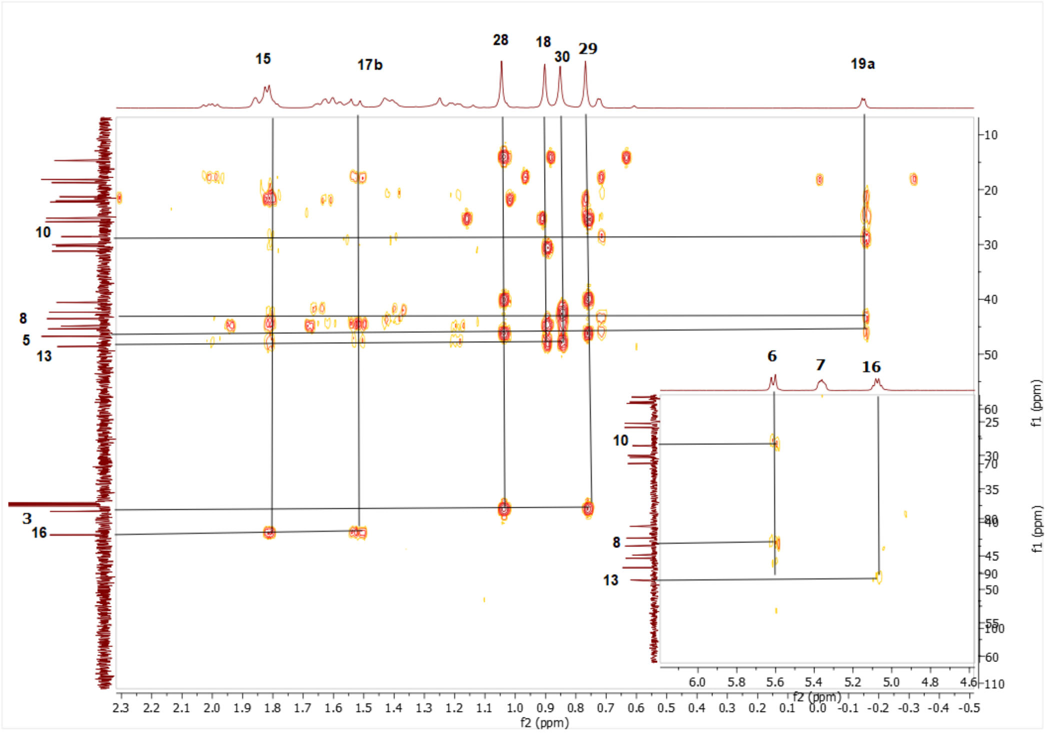

Spectrum 109. HMBC spectrum of SCG-05.

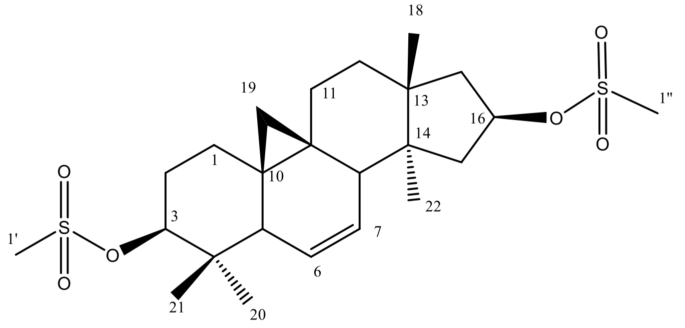

Supplementary Figure 19. Chemical Structure of SCG-06

Supplementary Table 17. The $^{13}C$ and $^{1}H$ NMR data of SCG-06 (100/500 MHz, δ ppm, in CDCl3).

| **H/C** | **$\delta_C$ (ppm)** | **$\delta_H$ (ppm), *J* (Hz)** |
|---|---|---|
| **1** | 29.5 t | 1.7 m, 1.92 m |
| **2** | 28.1 t | 1.92 m, 2.14 m |
| **3** | 89.5 d | 4.45 dd (12, 4.6) |
| **4** | 40.0 s | - |
| **5** | 46.7 d | 2.0 m |
| **6** | 125.9 d | 5.61 d (10.5) |
| **7** | 129.2 d | 5.45 ddd (10.3, 6.1, 3.0) |
| **8** | 43.2 d | 2.51 dd (6, 2.6) |
| **9** | 21.2 s | - |
| **10** | 27. 8 s | - |
| **11** | 25.0 t | 1.43 m, 1.89 m |
| **12** | 30.9 t | 1.27 dd (13, 5.1), 1.69 m |
| **13** | 44.7 s | - |
| **14** | 48.4 s | - |
| **15** | 42.3 t | 1.63 m, 2.2 m |
| **16** | 82.0 d | 5.29 q (7.6) |
| **17** | 45.2 t | 1.93 m, 2.04 m |
| **18** | 22.0 q | 0.98 s |
| **19** | 18.4 t | -0.07 d (4.3), 0.77 d (4.3) |
| **28** | 15.3 q | 0.86 s |
| **29** | 25.8 q | 1.1 s |
| **30** | 17.9 q | 0.87 s |
| **1'** | 39.0 q* | 3.03 s' |
| **1''** | 38.5 q* | 2.97 s' |

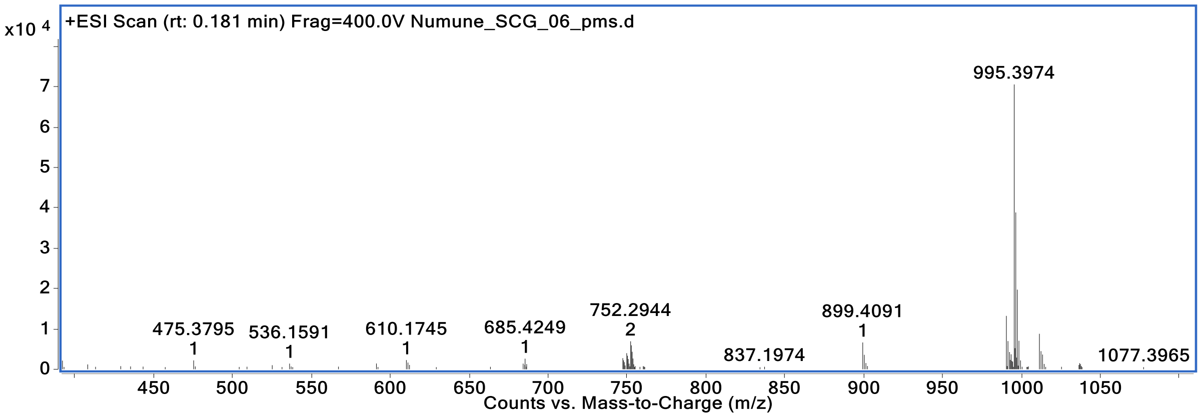

Spectrum 110. HR-ESI-MS Spectrum of SCG-06 (positive mode).

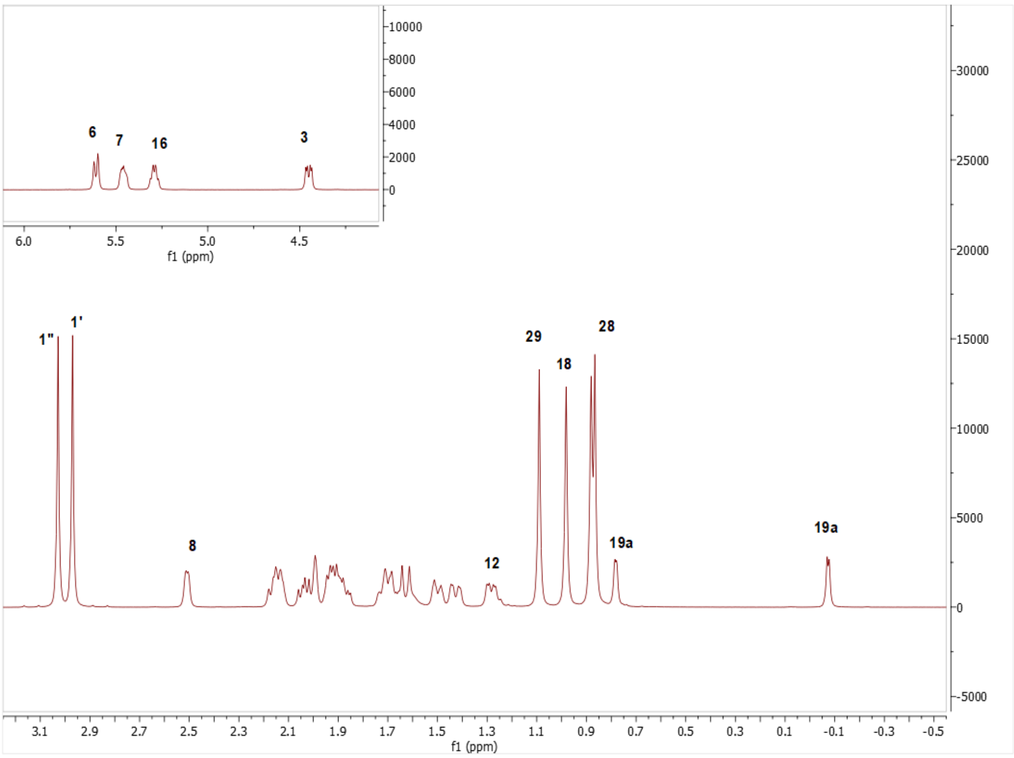

Spectrum 111. $^{1}H$ NMR Spectrum of SCG-06.

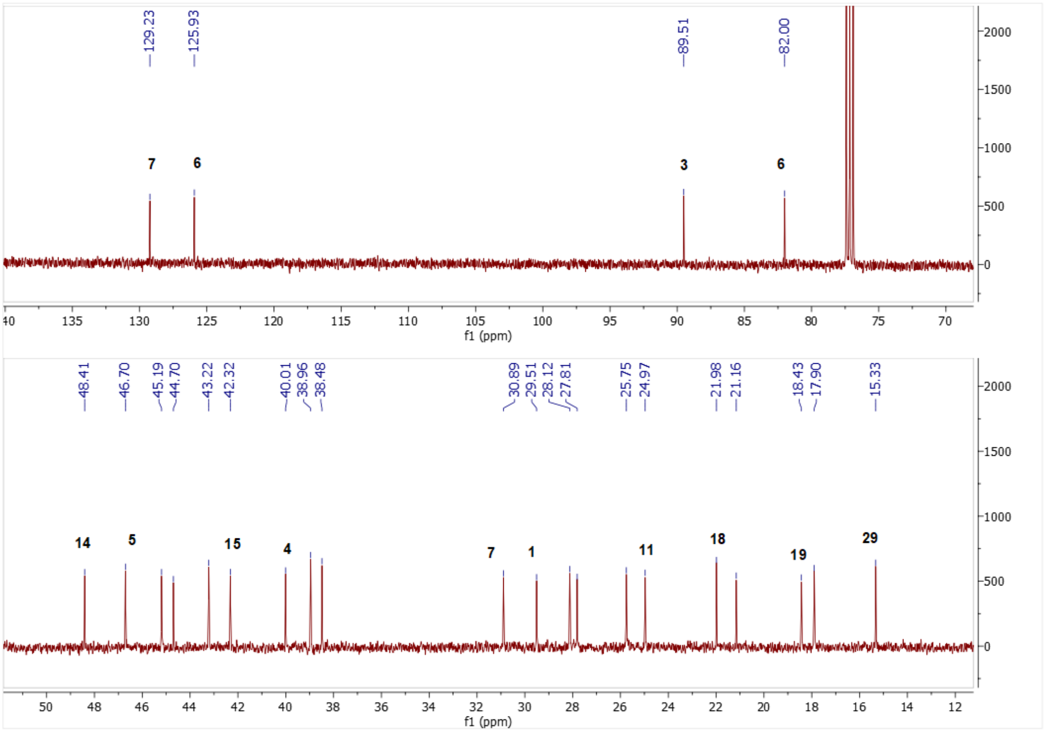

Spectrum 1127. $^{13}C$ NMR Spectrum of SCG-06.

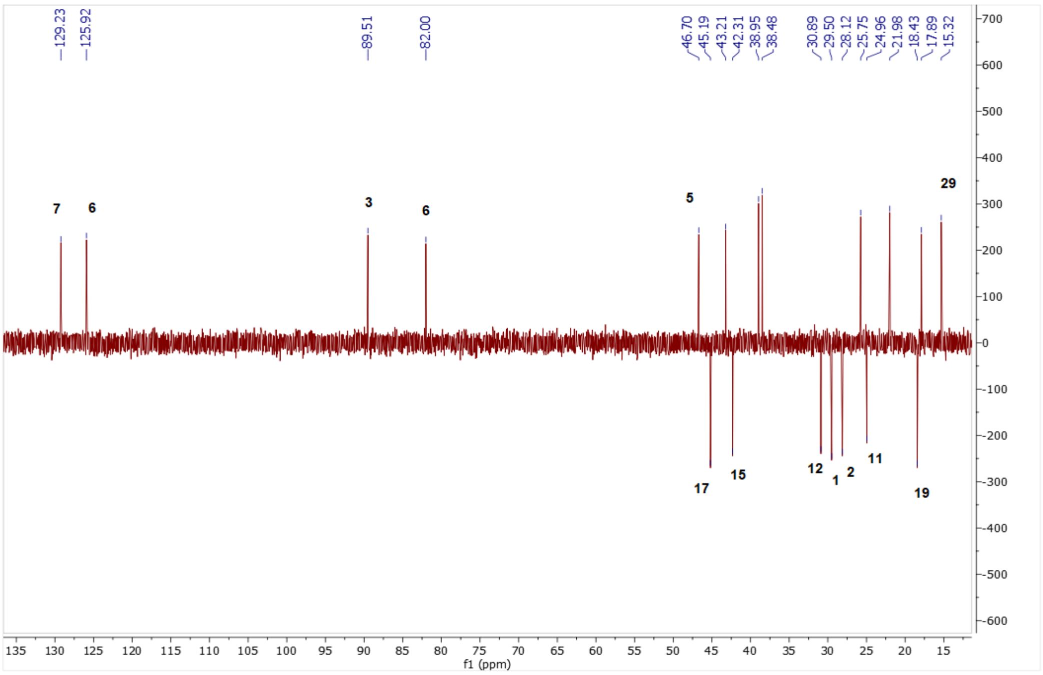

Spectrum 1138. DEPT135 spectrum of SCG-06.

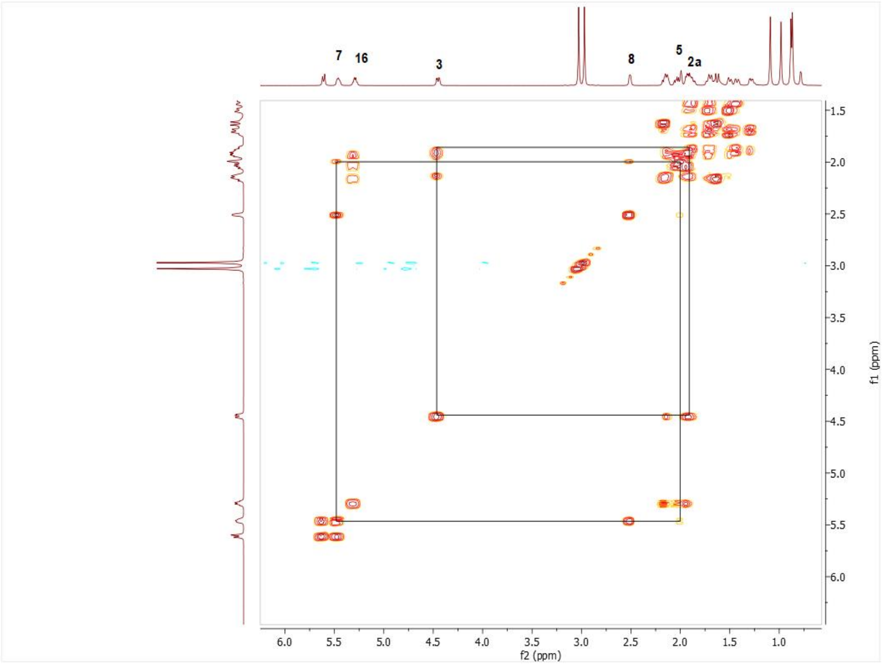

Spectrum 114. COSY spectrum of SCG-06.

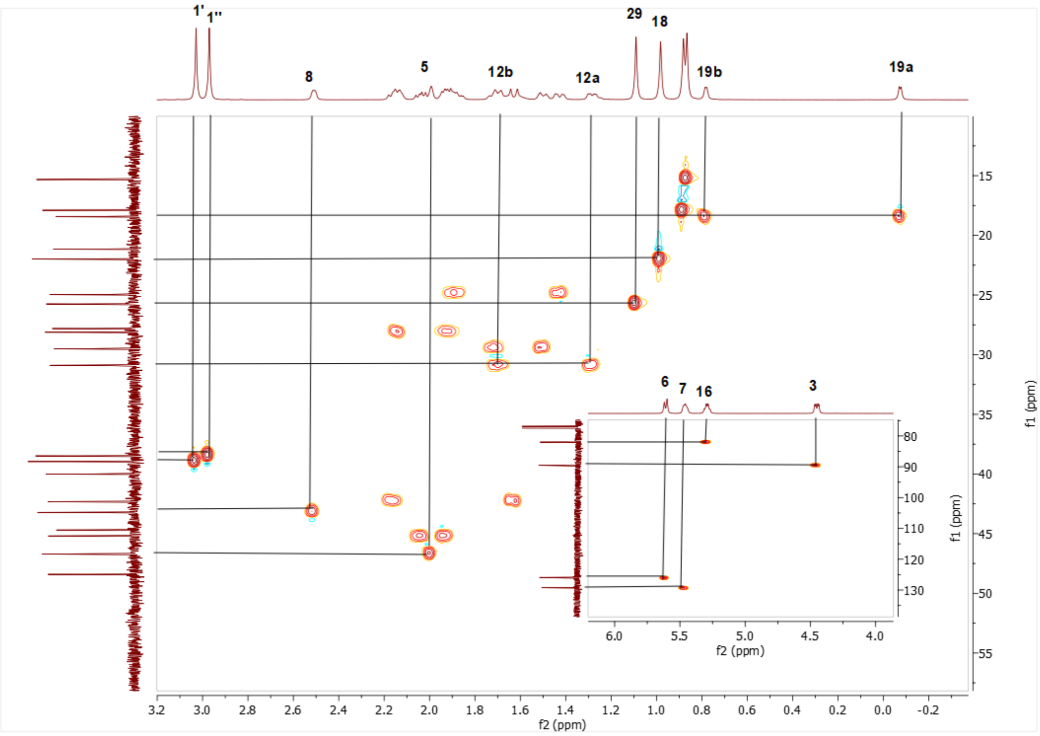

Spectrum 115. HMQC spectrum of SCG-06.

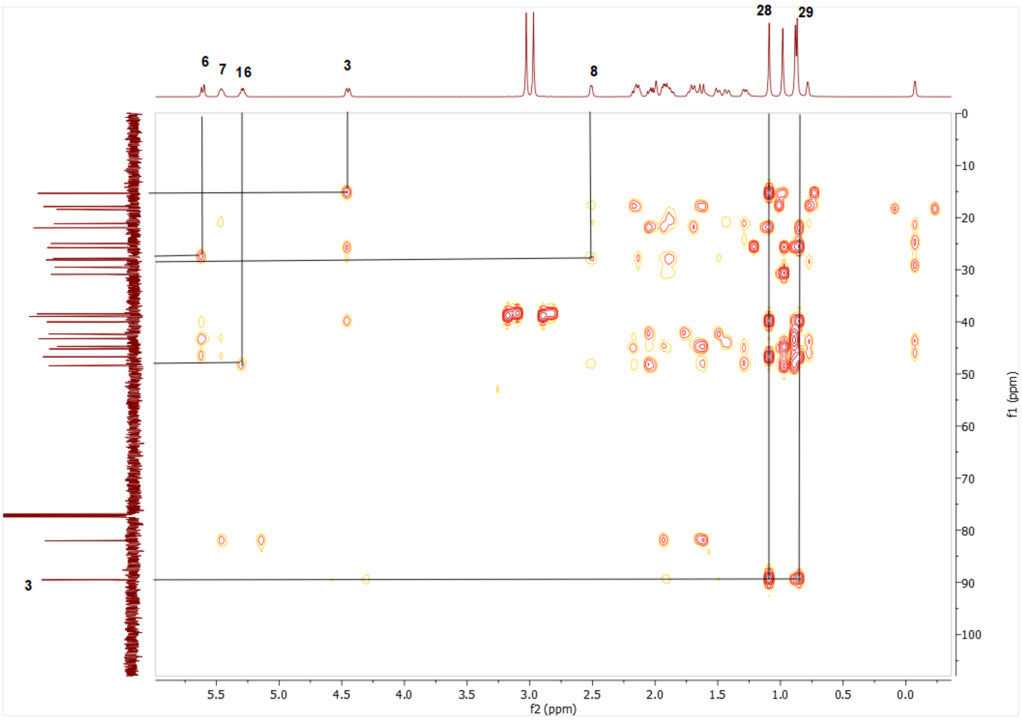

Spectrum 116. HMBC spectrum of SCG-06.

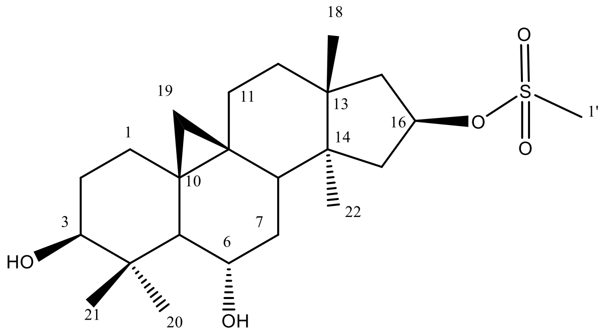

Supplementary Figure 20. Chemical Structure of SCG-07

Supplementary Table 18. The $^{13}C$ and $^{1}H$ NMR data of SCG-07 (100/500 MHz, δ ppm, in CDCl3).

| H/C | $\delta_C$ (ppm) | $\delta_H$ (ppm), *J* (Hz) |
|---|---|---|
| **1** | 31.3 t | 1.26 m, 1.63 m |
| **2** | 30.2 t | 1.58 m, 1.8 m |
| **3** | 78.48 d | 3.22 dd (11.2, 4.2) |
| **4** | 41.6 s | - |
| **5** | 53.4 d | 1.37 m |
| **6** | 68.4 d | 3.56 ddd (9.2, 9.2, 4.1) |
| **7** | 37.6 t | 1.38 m, 1.55 m |
| **8** | 45.6 d | 1.71 m |
| **9** | 20.7 s | - |
| **10** | 29.0 s | - |
| **11** | 26.1 t | 1.31 m, 1.97 m |
| **12** | 30.3 t | 1.48 m |
| **13** | 44.7 s | - |
| **14** | 46.0 s | - |
| **15** | 44.6 t | 1.79 m, 2.06 m |
| **16** | 82.39 d | 5.26 q (8) |
| **17** | 46.2 t | 2.01 m, 2.11 m |
| **18** | 24.9 q | 1.03 s |
| **19** | 26.7 t | 0.31 d, 0.5 d(4.7) |
| **28** | 15.3 q | 0.94 s |
| **29** | 27.9 q | 1.23 s |
| **30** | 19.8 q | 1.1 s |
| **1'** | 38.5 q | 2.97 s |

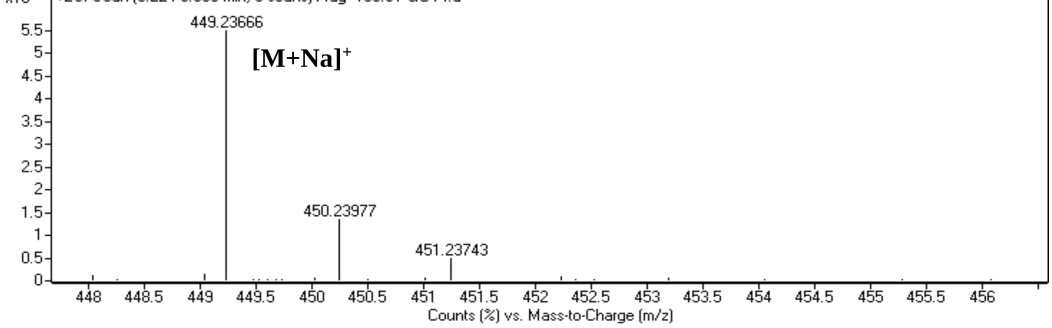

Spectrum 117. HR-ESI-MS Spectrum of SCG-07 (positive mode).

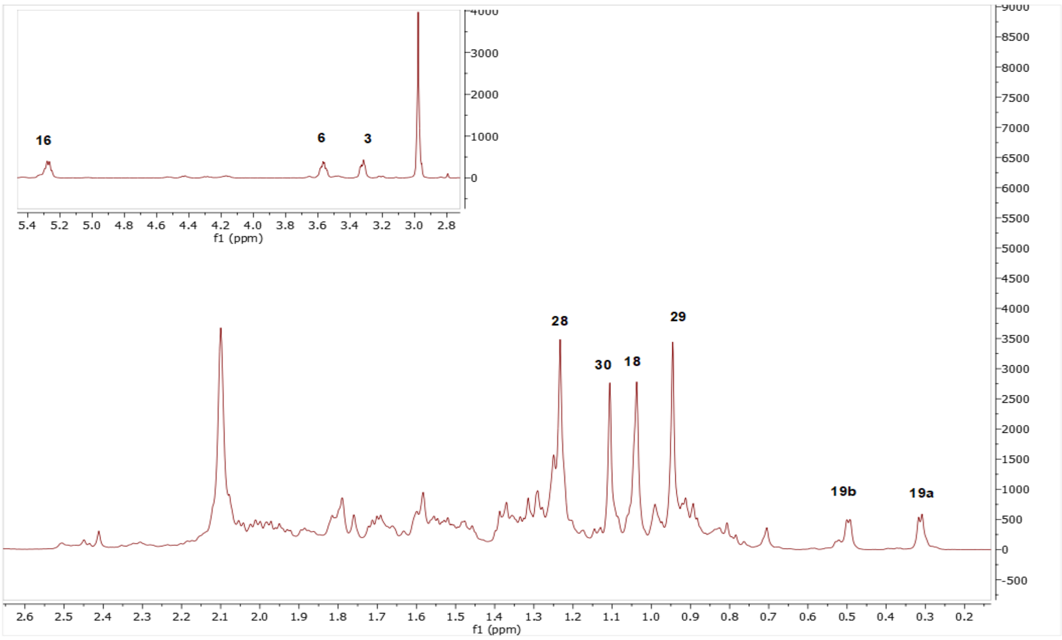

Spectrum 118. $^{1}H$ NMR Spectrum of SCG-07.

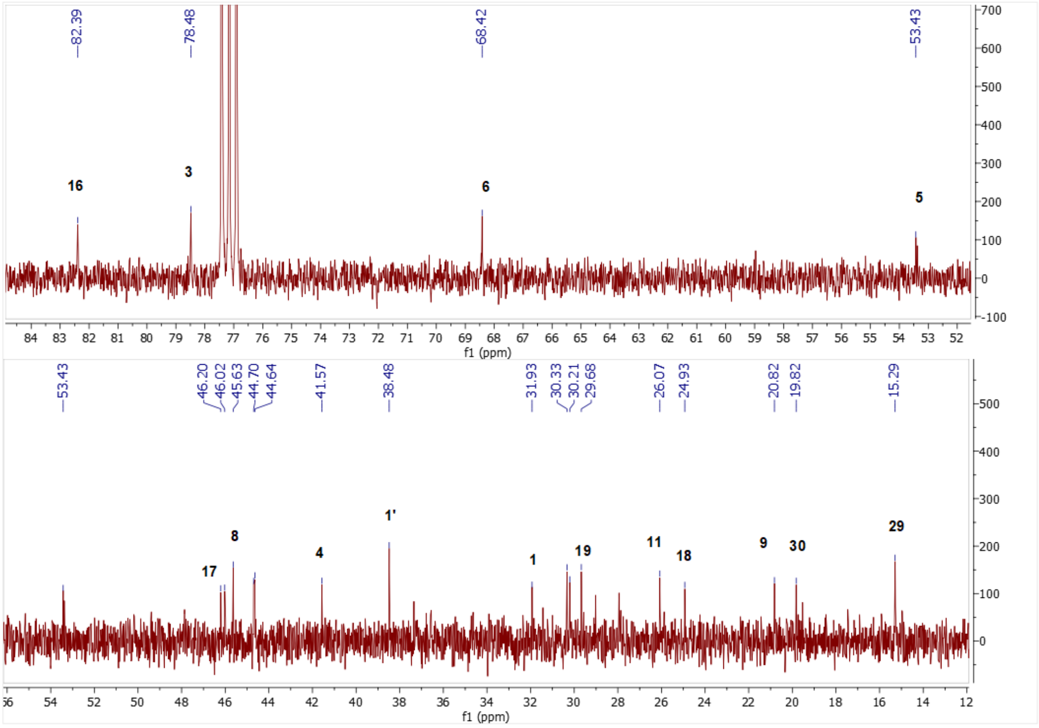

Spectrum 119. $^{13}C$ NMR Spectrum of SCG-07.

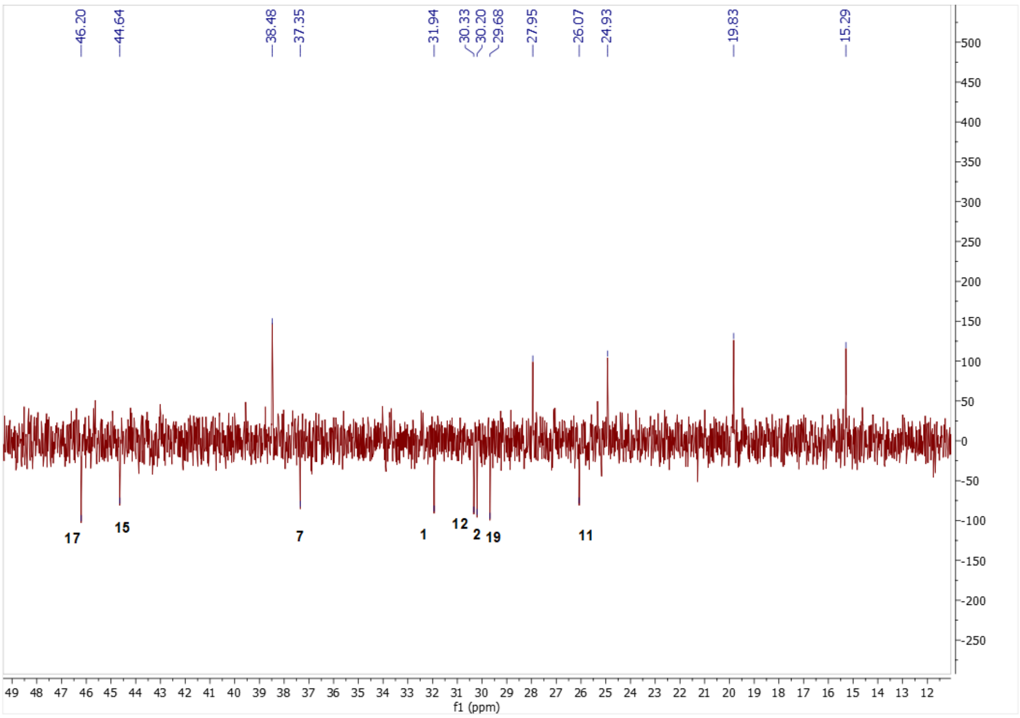

Spectrum 120. DEPT135 spectrum of SCG-07.

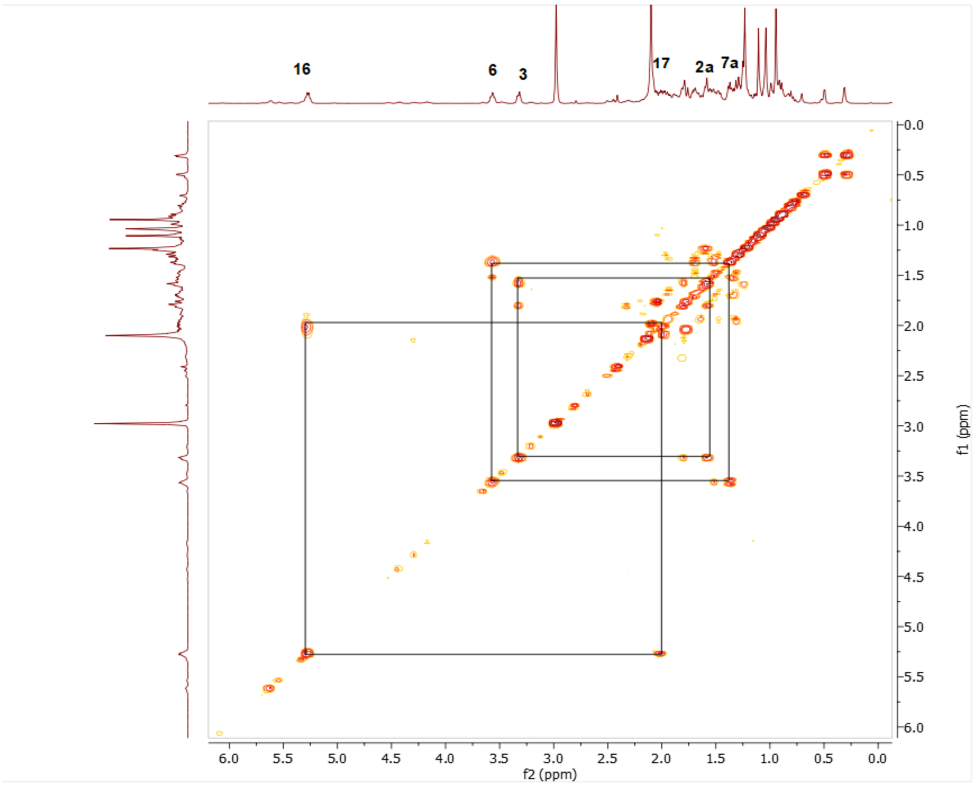

Spectrum 121. COSY spectrum of SCG-07.

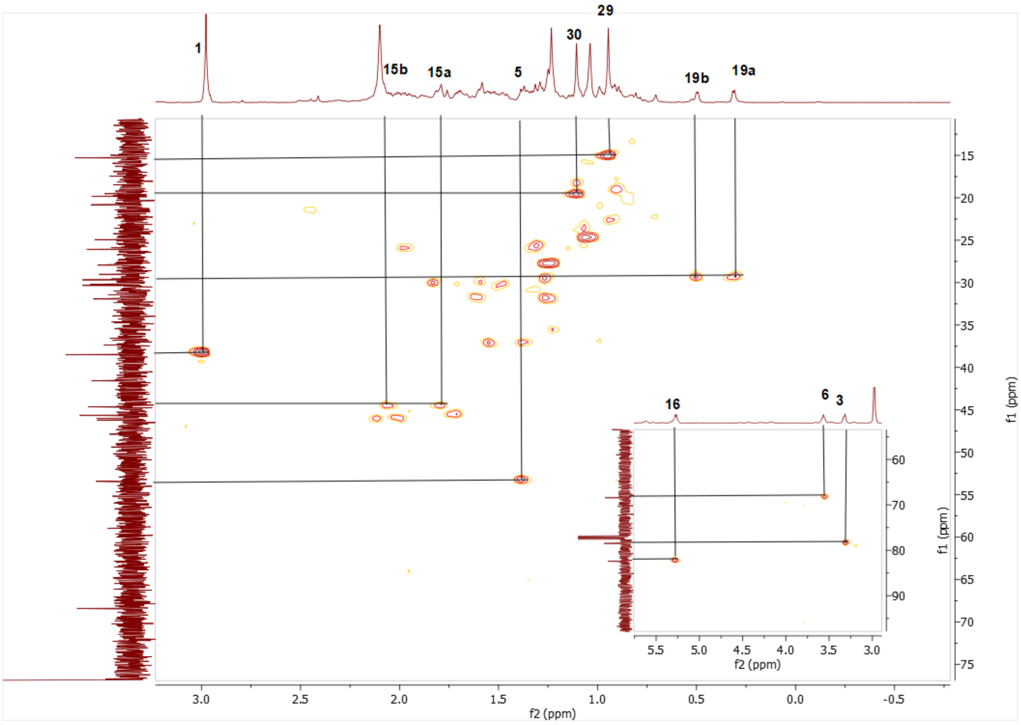

Spectrum 122. HMQC spectrum of SCG-07.

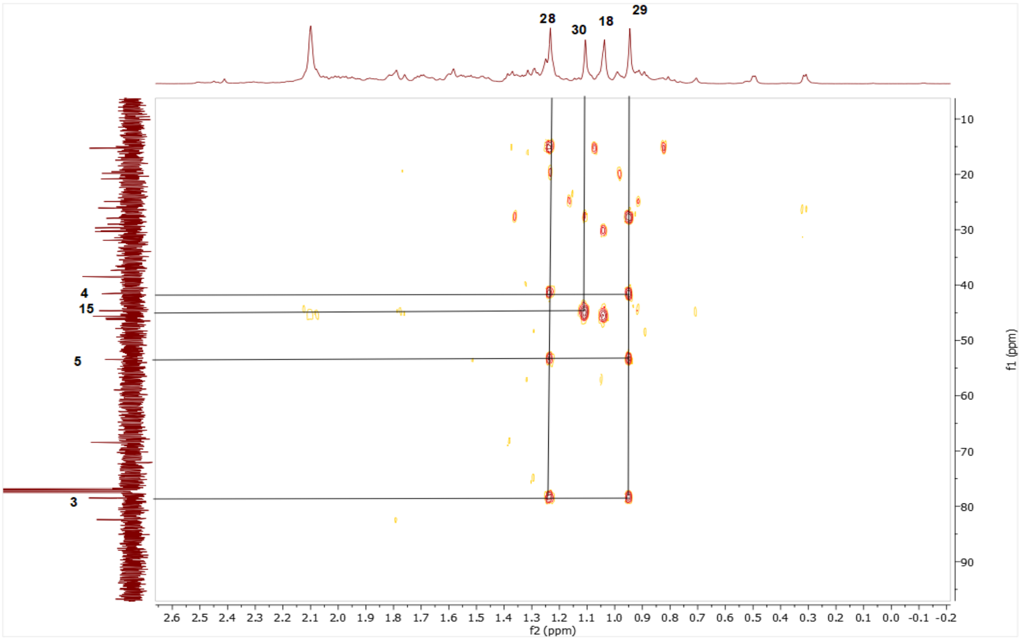

Spectrum 123. HMBC spectrum of SCG-07.
